# Evolutionarily diverse caveolins share a common structural framework built around amphipathic discs

**DOI:** 10.1101/2022.11.15.516482

**Authors:** Bing Han, Sarah Connolly, Louis F. L. Wilson, Darrin T. Schultz, Alican Gulsevin, Jens Meiler, Erkan Karakas, Melanie D. Ohi, Anne K. Kenworthy

## Abstract

Caveolins are a unique family of membrane-remodeling proteins present broadly across animals (Metazoa), and in vertebrates form flask-shaped invaginations known as caveolae. While human caveolin-1 assembles into an amphipathic disc composed of 11 spirally packed protomers, the structural basis underlying caveolin function across animals remains elusive. Here, we predicted structures for 73 caveolins spanning animal diversity, as well as a newly identified choanoflagellate caveolin from *Salpingoeca rosetta,* a unicellular relative to animals. This analysis revealed seven conserved structural elements and a propensity to assemble into amphipathic discs. Despite extreme sequence variability, new cryo-EM structures of caveolins from the choanoflagellate and the purple sea urchin *Strongylocentrotus purpuratus* exhibit striking structural similarities to human caveolin-1, validating the structural predictions. Lastly, tracing the chromosomal evolutionary history of caveolins revealed evolutionary branches where caveolins translocated and expanded, including a parahoxozoan ancestral chromosome as the origin of most caveolin diversity. These results show that caveolins possess an ancient structural framework predating Metazoa and provide a new structural paradigm to explore the molecular basis of caveolin function across diverse evolutionary lineages.

## Introduction

Eukaryotes contain elaborate endomembrane systems consisting of morphologically and functionally distinct membrane-bound compartments. The construction and maintenance of these compartments relies on the actions of ancient families of proteins capable of remodeling membranes (*1, 2*). To support enhanced requirements for cell-cell adhesion, communication, and signaling during the transition from single-celled eukaryotes to animals (Metazoa), a dramatic expansion in membrane-associated proteins occurred ^1^. Among the membrane proteins thought to have first emerged in Metazoa is the caveolin family of membrane remodeling proteins ^1, 2, 3^. Best recognized for their role in vertebrates as structural components of caveolae, flask-shaped invaginations of the plasma membrane, caveolins have been identified in most metazoan clades ^2, 3, 4^. Thus they have fulfilled essential biological roles since the ancestor of the clade existed approximately 800 million years ago ^5^. In humans, caveolins and caveolae are distributed throughout the body serving as important regulators of multiple organ systems ^6, 7, 8, 9^. Furthermore, caveolins and caveolae are broadly implicated in regulation of cell signaling, lipid metabolism, and sensing and responding to stress ^8, 10, 11, 12, 13^.

Unlike vesicle coat proteins such as clathrin, COPI, and COPII that cycle on and off membranes and share evolutionary origins and structural features ^14^, caveolins are unrelated in sequence to other vesicle coat proteins and remain integrated in membranes throughout their life cycle. Most studies of caveolins have focused on their roles in mammalian cells where caveolae are often abundant ^15, 16, 17^. However, in some cell types lacking cavins, a second family of proteins required for caveolae biogenesis, caveolins can function independently of caveolae ^18, 19, 20, 21^. Since the cavin family appears to be found only in vertebrates ^22^, this suggests that in most organisms caveolins function independently of classically defined caveolae. To date, however, only a handful of examples of caveolins from non-vertebrate organisms have been studied ^3, 23, 24, 25, 26^. It is also unclear whether caveolins exist in ctenophores, the sister clade to all other animals ^27^, and whether caveolins’ provenance can be traced back to known ancestral chromosomes ^28^. Thus, our knowledge of the existence and functions of caveolins across evolutionary space is limited.

The molecular architecture of caveolins was unknown until the discovery of the structure of the homo-oligomeric 8S complex of human caveolin-1 (Cav1), an essential component of caveolin in non-muscle cells ^29, 30^. Cryo-electron microscopy (cryo-EM) showed the complex is composed of 11 Cav1 protomers symmetrically arranged into an unusual amphipathic disc-like structure predicted to fully insert into the cytoplasmic leaflet of the plasma membrane ^29, 30, 31^. However, whether related proteins from other organisms (including distantly related caveolin homologues) behave in the same way is completely unknown.

Here, we report that in addition to being expressed in metazoans, caveolin homologs exist in choanoflagellates, free-swimming unicellular protists and the closest relatives of animals, suggesting that caveolins are of pre-metazoan and pre-choanozoan origin. We also find that ctenophores lack caveolins, and that most of caveolin diversity in animals can be traced to a single ancestral chromosome in the ancestor of the Parahoxozoa. Using a combination of computational, phylogenetic, and structural approaches, we show that despite extreme sequence variability, even the most distantly related caveolin homologues share a surprisingly conserved set of structural elements. These findings suggest that caveolins share an ancient and conserved structural framework that diverse organisms co-opted to fulfill distinct physiological roles. They also provide a new framework to probe the structural basis for the function of caveolins across evolution.

## Results

### Consistent evidence for the existence of choanoflagellate caveolins

Caveolins have typically been treated as a metazoan-specific family ^1, 2, 3, 32^. However, one sequence located on chromosome 10 in the choanoflagellate *Salpingaeca rosetta* (strain: ATCC 50818) is currently annotated in the UniProt database as a caveolin (UniProt: F2U793) ^27^. Surprisingly, this sequence shares almost no recognizable sequence identity with human Cav1 (13%); however, if truly related, it would represent the most evolutionarily divergent caveolin found to date. To verify this annotation, we built a caveolin HMM (hidden Markov model) profile from an alignment based on a previously identified region of high conservation ^3^. Encouragingly, HMMER searches against the *S. rosetta* proteomes retrieved the F2U793 sequences with an E-value of 8.8 x 10^−11^, indicating a confident prediction of the homology. By contrast, a sequence from the filasterean protist *Capsaspora owczarzaki* (strain: ATCC 30864) annotated in UniProt as a caveolin (A0A0D2VH37) yielded an E-value > 0.05, suggesting this annotation is spurious. Moreover, genomic frame-shifted HMM profile searches of the *C. owczarzaki* and ichthyosporean *Creolimax fragrantissima* genomes ^27, 33, 34^ did not yield identifiable caveolin proteins (E-value > 0.05). These findings support the idea that caveolins are in fact not limited to Metazoa but are also present in some choanoflagellates, the closest relatives of Metazoa. Furthermore, *S. rosetta* can exist both as a free-living single cell and in a colonial pseudomulticellular state, highlighting its unique position in the evolution from single-celled organisms to multicellular animals.

### Ancestral Linkage Group Analysis reveals caveolin chromosomal origins

The unexpected finding of a putative caveolin in *S. rosetta* led us to revisit the evolutionary history of caveolins. Current evidence suggests all vertebrate caveolins descend from three ancestral sequences: CavX, CavY, and CavZ. CavX and CavZ were colocalized to the same chromosome in the ancestral vertebrate genome ^3^. Whereas CavY appears to have been lost in most vertebrates, CavX is proposed to have given rise to Cav1 and Cav3, the ‘canonical’ caveola-forming family members, which also has closely related homologs in cnidarians ^3^. CavZ appears to have given rise to the Cav2 ^3^.

To understand the chromosomal origins of caveolins and their emergence in animals, we analyzed ancestral linkage groups (ALGs) to identify the ancestral chromosome on which caveolin originated. ALGs are collections of sequences representing whole or partial chromosomes in a clade’s ancestor. There were 29 ALGs in the ancestor of myriazoans (all animals except ctenophores) and are highly conserved on single chromosomes in several animal clades ^28^, with some partially preserved on single chromosomes in the choanoflagellate *S. rosetta* ^27^.

To perform this analysis, we collected chromosome-scale genomes from across animal diversity with minimal chromosomal changes since their divergence from the Myriazoan ancestor ^35^ (**Fig. 1A, Supplementary File 1**). With these genomes, we identified caveolin orthologs on chromosome-scale scaffolds and identified the myriazoan ALGs associated with those chromosomes. No credible caveolin orthologs were found in the filasterean amoeba *Capsaspora owczarzaki,* in ctenophores, or in scyphozoan or hydrozoan cnidarians (**Fig 1B**). However, genes encoding caveolins or caveolin-like proteins were detected in the chromosome-scale genomes of two 400 million-year-diverged sponges, and in anthozoan cnidarians (corals, anemones) (**Fig 1B**). Corroborating previous results ^3^, we found that genomes within the Parahoxozoa, a clade of animals that consists of Bilateria, Placozoa, and Cnidaria, contained many caveolin paralogs (**Fig 1B)**.

**Fig. 1.**
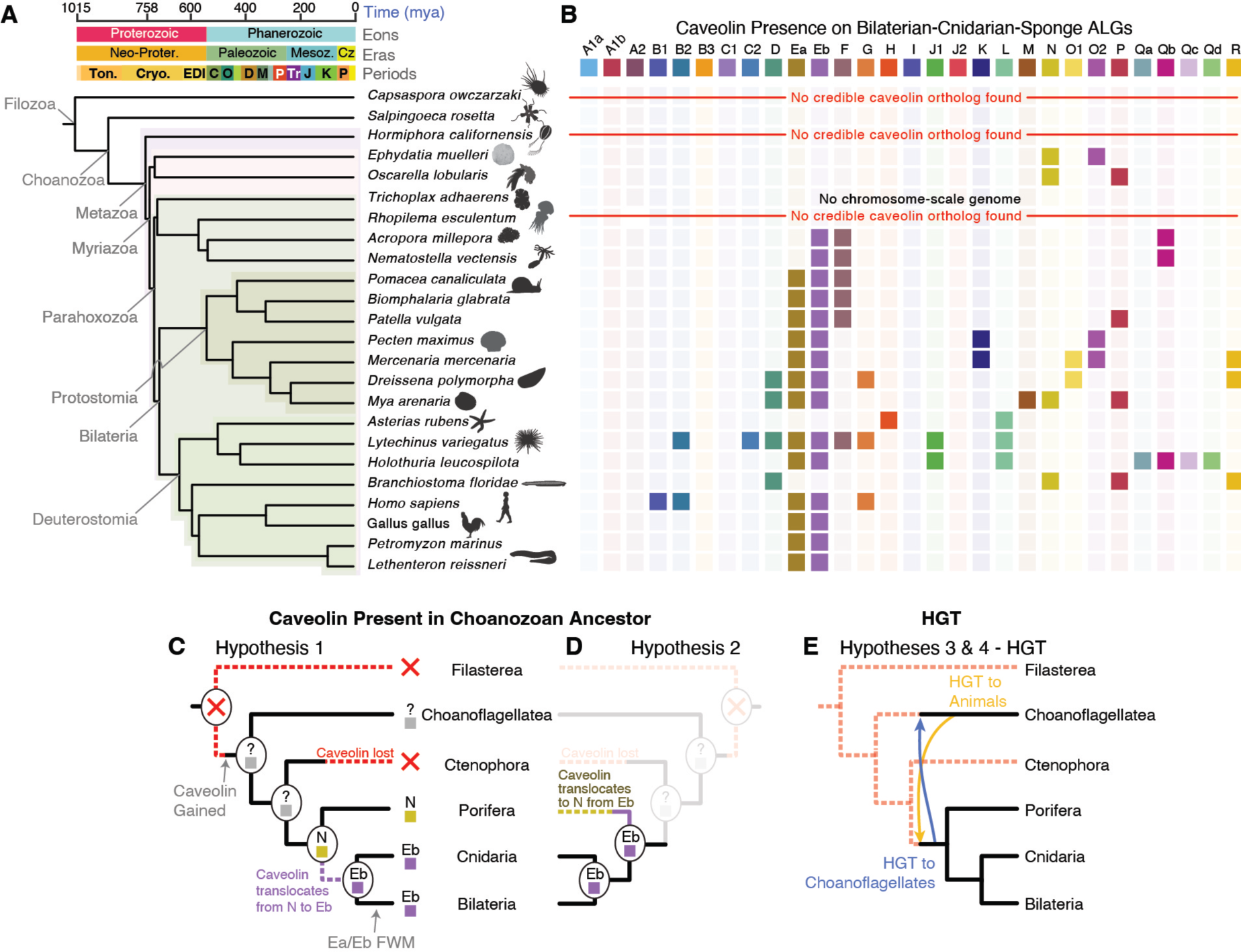
Chromosomal origins of caveolin. (**A**) Samples across animal diversity as well as in two unicellular relatives of animals were used to determine at the ALG identities of chromosomes on which caveolins were present. **(B)** Caveolins are present on Bilaterian-Cnidarian-Sponge (BCnS) ALG Eb-bearing chromosomes in all but two species observed within the Parahoxozoa. In sponges, caveolins are present on ALG N in species from two anciently diverged sponge clades. The BCnS ALG identity of *S. rosetta* chromosome 10, on which caveolin is present, does not have a significant ALG identity. **(C-E)** Possible models for the chromosomal origins of caveolin. (C, D) Assuming that caveolin was present in the choanozoan ancestor, it must have been lost in the ctenophore lineage. Caveolin’s chromosomal provenance is parsimoniously equally likely to have been ALG N, followed by translocation to ALG Eb in the parahoxozoan ancestor (C), or ALG Eb in the ancestral myriazoan genome, followed by translocation to ALG N in the common ancestor of Demospongia and Homoschleromorpha sponges. (E) Another potential explanation of caveolin’s provenance that accounts for its presence in both animals and in choanoflagellates is horizontal gene transfer (HGT). This could have occurred in either direction after acquisition of the gene (i.e. from choanoflagellates to animals, or from animals to choanoflagellates).

Using the myriazoan ALGs ^28^, we identified a pattern of caveolin-ALG colocalization within the Parahoxozoa — that caveolin genes were located on chromosomes containing the ALG Eb (**Fig 1B**). Exceptions to this are that caveolins were absent on Eb-bearing chromosomes in the lancelet *Branchiostoma floridae* and the sea star *Asterias rubens* (**Fig 1B**). As the ALGs Ea and Eb fused before the Bilaterian ancestor (*34*), this indicates that caveolin was present on this ancestral chromosome (**Fig. 1B**). With regards to caveolin’s pre-bilaterian origins, we note that ALG Ea is not fused with ALG Eb in cnidarians. In cnidarians, ALG Ea and Eb remain on separate chromosomes (*34)*, but we found putative caveolins on Eb-containing chromosomes in the two cnidarians that diverged between 300 and 600 million years ago ^36^, suggesting that caveolin was present on the ancestral Eb-bearing chromosome before Anthozoa diversified. Similarly, caveolin-like sequences were found on ALG N-containing chromosomes in two sponge species, despite their divergence over 500 million years ago ^37^ (**Fig 1B**).

These findings suggest two equally parsimonious scenarios for the chromosomal origin of caveolins in myriazoans (**Fig 1C,D**). In one, caveolin originated on ALG Eb in the myriazoan ancestor and later translocated to ALG N in sponges. In the other, caveolin was originally on ALG N in the myriazoan ancestor and translocated to ALG Eb in the lineage leading to Parahoxozoa. Due to the absence of caveolins in ctenophores and the extensive rearrangements between animal and choanoflagellate genomes, we are not able to determine the chromosome on which caveolin originated in the animal ancestor, or whether caveolin was present in the Choanozoa ancestor. Additionally, we cannot formally exclude the possibility of horizontal gene transfer between the ancestors of myriazoans and choanoflagellates (**Fig 1E**).

### Phylogenetic analysis provides further insights into the evolutionary history of caveolins

To gain further insight into the specific evolutionary relationship between caveolins, we set out to conduct an updated phylogenetic analysis of the caveolin family since many new metazoan species have been sequenced since the last analysis was performed ^3^. Although genome information has been obtained for over 3,200 animal species so far, vertebrates, which account for only 3.9% of all species, represent over 50% of the species that have been sequenced ^38^, which can skew this type of analysis. To gain a more balanced and comprehensive view of how caveolin evolves across different evolutionary branches, we selected caveolins from one representative species from each phylum or superphylum of Metazoa, along with the previously mentioned caveolin from *S. rosetta* for further study.

For this analysis, we included caveolins from *Amphimedon queenslandica* (Porifera, Metazoa), a species of evolutionary interest due to its early divergence with respect to vertebrates ^39, 40^. In the *A. queenslandica* reference proteome, we found five proteins with caveolin Pfam annotations (UniProt: A0A1X7UHP5, A0A1X7UGA1, A0A1X7TMH4, A0A1X7VPY7, A0A1X7VRV8). As with the *S. rosetta* sequence, the *A. queenslandica* sequences were highly dissimilar to human CAV1 (11–19% identity). A HMMER search against the *A. queenslandica* genome retrieved hits with significant E-values (ranging from 5.1 × 10^−15^ to 2.5 × 10^−11^). This suggests that the putative sponge caveolins may be true homologs to the caveolins found in bilaterians, a major metazoan clade characterized by bilateral symmetry.

Next, we inferred a maximum-likelihood phylogeny using 74 protein sequences with caveolin Pfam annotations from 15 distantly related holozoans (**Table S1**), in addition to a set of previously categorized caveolin and caveolin-like sequences ^3^. Though achieving only weak bootstrap support values, the resulting tree largely replicated the previously proposed clades ^3^, consisting of Cav1/3, Cav2/2R, CavY extended, Protostomia Group 1, Protostomia Group 2, and Cav-like (**Fig S1**). Many of the sequences of interest could be tentatively assigned to one of these previously defined groups (**Fig S1**). However, the caveolin homologs from *A. queenslandica* also could not be placed in any of these pre-established clades. Instead, these sequences appeared in a monophyletic group that appears to be a sister to all other metazoan sequences, suggesting an early divergence with respect to the other metazoan sequences.

Given the long branch length connecting the *S. rosetta* sequence to the rest of the tree and the known evolutionary relationship between choanolagellates and Metazoa, we designated this sequence as a provisional outgroup, named Choa-CAV (**Fig S1**). Correspondingly, we named all metazoan caveolins as Meta-CAV. We chose to designate the clade containing caveolin homologs from *A. queenslandica* group as “atypical caveolins” and the remaining Meta-CAV groups as “typical caveolins”. We further broke down the typical caveolins into Type I and Type II-CAV. Almost all the relatively well studied caveolins, such as human Cav1, Cav2, and Cav3, as well as *C. elegans* caveolins and *Apis mellifera* caveolin, belong to Type II-CAV. Type I-CAV corresponds to the CAV-like clade identified in a previous study ^3^. In the following sections, we will use this newly defined framework to trace the evolutionary trajectory and compare structural similarities and differences among caveolins.

### Caveolin protomers are predicted to organize into disc-shaped complexes composed of spiraling amphipathic α-helices

The existence of distantly related caveolin homologues with limited sequence similarity such as Choa caveolin raises the question as to whether they have similar folds and/or functions with human Cav1 ^29^. To investigate whether other metazoan caveolins and the newly identified choanoflagellate caveolin share similar structural features, we used AlphaFold2 (AF2) as a predictive tool ^41^. Despite the unusual features of human Cav1, AF2 is able to predict its overall fold and reproduce its ability to oligomerize to form an amphipathic, disc-shaped structure characteristic of the Cav1 8S complex ^42^.

We predicted the structures of 74 Pfam-annotated caveolin family members from the representative species each phylum or superphylum of Metazoa as well as the Choa caveolin. A single example from each species is shown in **Fig 2**. For our initial analysis, we generated *n*-mers of increasing size with AF2.1 (**Supplementary File 2**) and then generated the 11-mer with AF2.2 (**Supplementary File 3**). In essentially all the examined species, the caveolins were predicted to form closely packed amphipathic discs or rings that spiral in a clockwise direction when viewed from the cytoplasmic face (**Fig 2**). Most were also predicted to contain N-terminal disordered regions located around the outer rim of the complex and central β-barrels formed by parallel β-strands, similar to the structure of human Cav1. Interestingly, C-terminal disordered regions emanating from the central β-barrel were also visible in several caveolins (**Fig 2**). Similar results were obtained when we used AF2.2 to predict the structure of the caveolins used in the ALG analysis, which included five metazoan phyla (Porifera, Cnidaria, Mollusca, Ambulacraria, and Chordata) (**Supplementary File 3**). In contrast, AF2.2 predicts for the sequence A0A0D2VH37 from *Capsaspora owczarzaki* forms a much different structure (**Supplementary File 2**), consistent with the conclusions drawn from the sequence alignment.

**Fig. 2.**
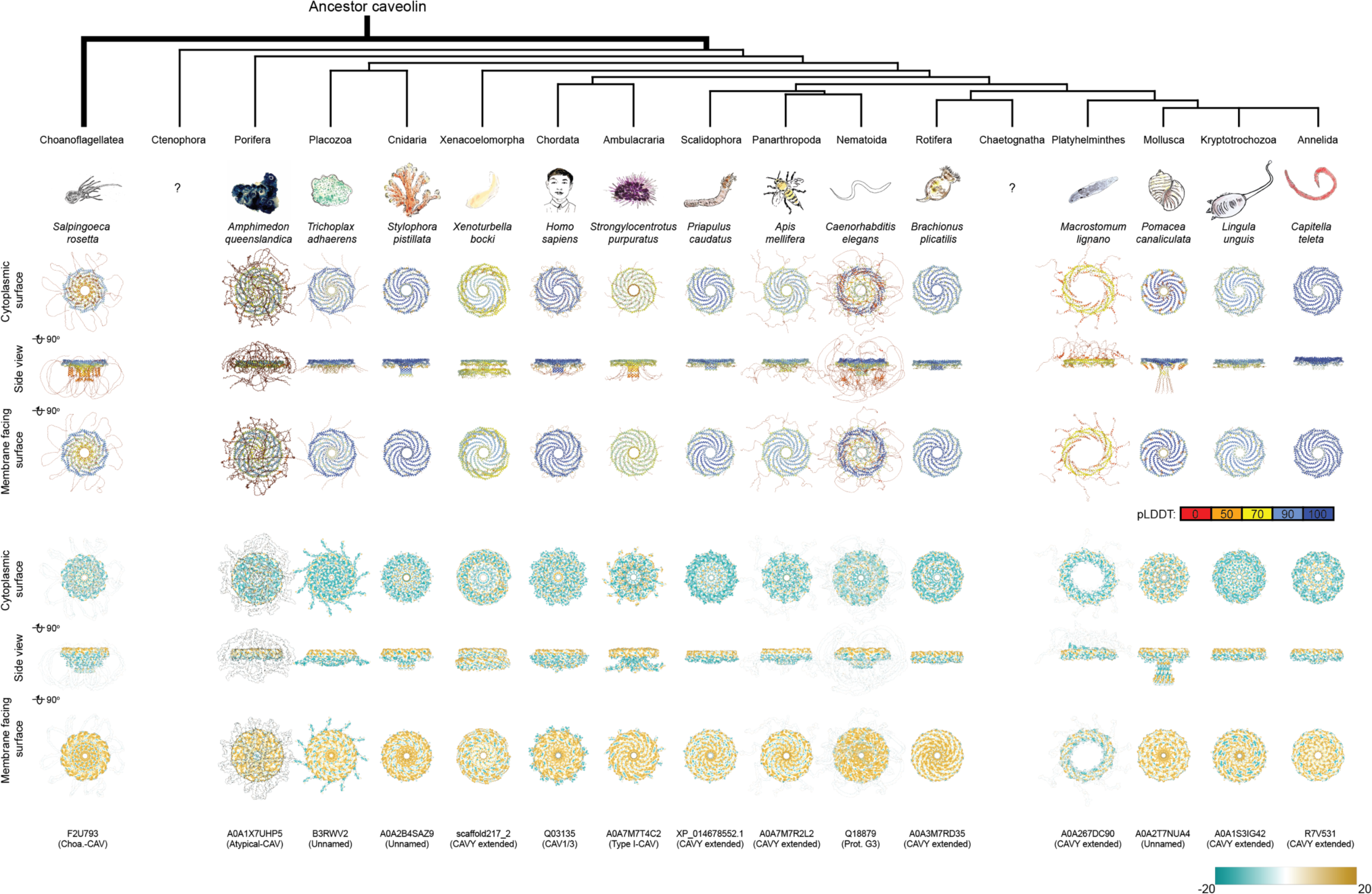
Conserved structural features of metazoan and choanoflagellate caveolins highlighted by computational modeling predictions. AF2.2 models for 11-mers caveolins from representative species of Choanoflagellatea and fourteen different metazoan phyla/superphyla. Models in top three rows are in ribbon mode and colored by pLDDT confidence values. Models in bottom three rows are in surface mode and colored by lipophilicity values. To better demonstrate the distribution pattern of hydrophobicity in surface mode, we applied transparency to the N-terminus of F2U793, A0A1X7UHP5, A0A7M7R2L2, Q18879, and the residues near the C-terminus of A0A267DC90.

Together, these results suggest that even in the most distantly related species, caveolins can homo-oligomerize into amphipathic discs and contain similar structural elements as human Cav1. They also highlight the presence of additional structural elements such as a C-terminal variable region in some caveolins. For simplicity, in the subsequent analyses, we will focus on results obtained from caveolins obtained from the set of 74 Pfam-annotated caveolins.

### Caveolins across evolution consist of seven basic structural units

We next sought to identify common structural motifs shared by these diverse caveolins. Previous studies identified a series of functionally important domains across mammalian caveolins, including the oligomerization, scaffolding, and intramembrane domains ^43, 44, 45, 46, 47^. However, these domains were primarily identified by sequence analysis or truncation studies and do not map in a straightforward way to the experimentally determined structure of human Cav1 ^29^. We thus identified seven basic structural units using the structure of the human CAV1 and a computationally predicted structure of *C. elegans* caveolin (Q94051) as templates (**Fig 3**):

**<u>1) N-terminal variable region</u>**. In human Cav1, residues 1-48 are predicted to be disordered and were not resolved in the cryo-EM structure ^29^. Similarly, many other caveolins, including *C. elegans* caveolin Q94051, are predicted to contain N-terminal disordered regions (**Fig 3, yellow**).
**<u>2) Pin motif.</u>** The pin motif of human Cav1 (residues 49-60) makes critical contacts with each neighboring protomer at the rim of the 8S complex ^29^. A similar motif is predicted to exist in *C. elegans* caveolin Q94051 (**Fig 3, red**).
**<u>3) Hook structure.</u>** Residues 61-81 of human Cav1 consist of a loop that undergoes a 180° turn (**Fig 3, blue**). A similar hook-shaped structural motif is predicted to exist in *C. elegans* caveolin Q94051 (**Fig 3**). This structural element corresponds to the first half of the oligomerization domain (residues 61-101) of human Cav1. Embedded within this same region is the highly conserved signature motif (residues 68-75), consisting of a 3/10 helix followed by a short-coiled structure.
**<u>4) Scaffolding domain.</u>** Residues 82-101 of human Cav1 are traditionally defined as the caveolin scaffolding domain (CSD). This corresponds to the initial α-helix (α-1) of the Cav1 protomer, which forms the periphery of the 8S complex disc ^29^. Importantly however, the α-1 helix extends beyond the classical boundaries of the CSD. Thus, we redefined the entire α-1 region (residues 81-107) as the scaffolding domain, considering its cohesive functional role in the experimental and predicted structures (**Fig 3**).
**<u>5) Spoke region.</u>** Residues 108-169 of human Cav1 consist of a series of α-helices connected in tandem by short loops, forming a semi-circle arc of about 180° ^29^. This region encompasses the previously defined as the intramembrane domain (IMD) (residues 102-134) as well as the helical region we previously designated as the spoke-like region (residues 135-169) (**Fig 3, gray**). To reflect the structural continuity of this region, we here define it as the spoke region.
**<u>6) C-terminal β-strand.</u>** Residues 170-176 of the C-terminal domain of the human Cav1 protomer fold into a β-strand that assembles into an amphipathic parallel β-barrel together with neighboring protomers ^29^. A β-strand is likewise predicted to exist in *C. elegans* caveolin (**Fig 3, cyan**).
**<u>7) C-terminal variable region.</u>** While the structure of human Cav1 essentially ends in a β-strand, a subset of other caveolins contain an additional C-terminal region that differs in length and composition across caveolins (**Fig 3, purple**). Accordingly, we refer to this region as the C-terminal variable domain. The structure of this region is typically predicted by AF2 with low confidence, suggesting it is disordered (**Supplemental Files 2 and 3**).

**Fig. 3.**
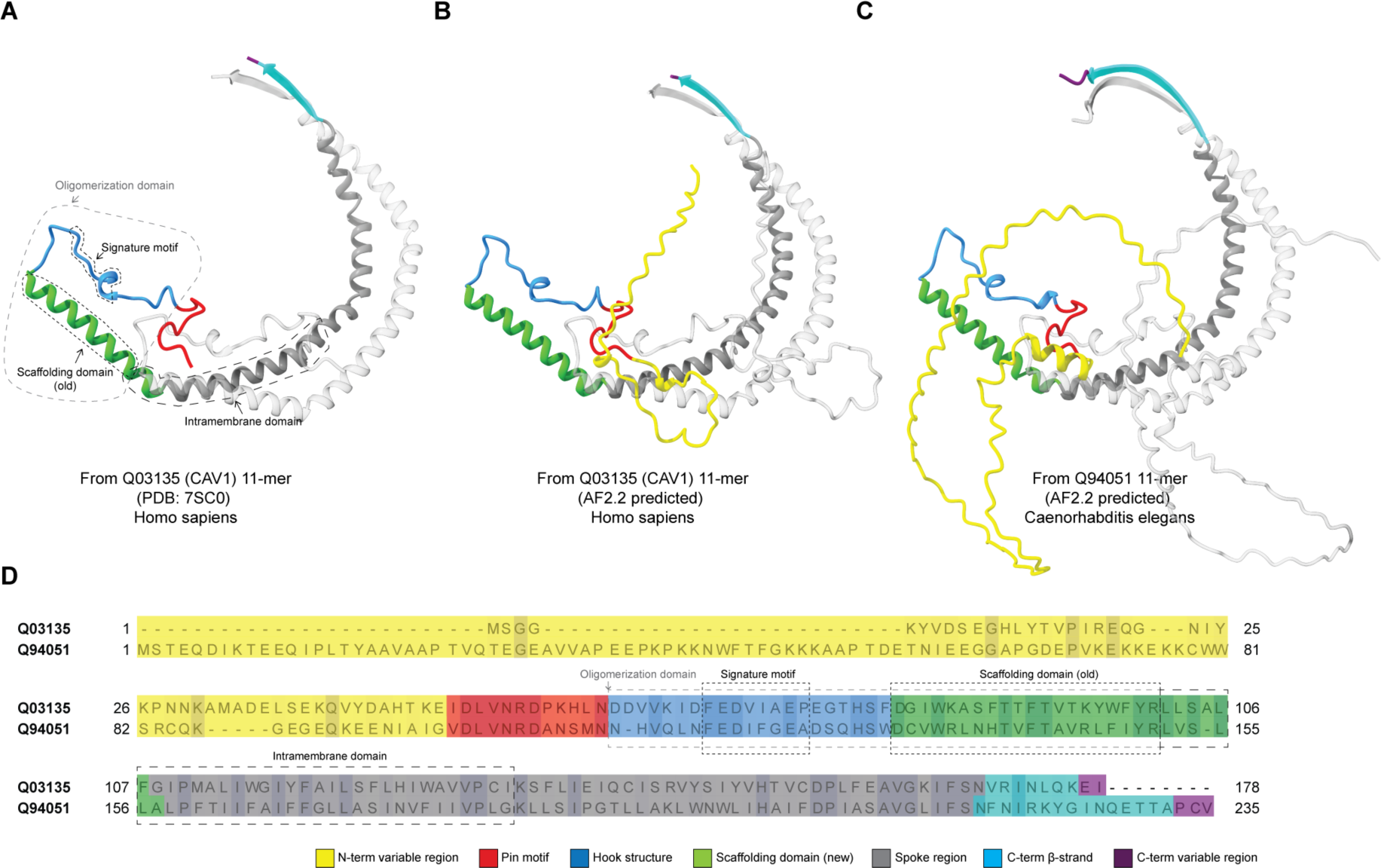
Proposed structural elements of caveolins. **(A-D)** Structural elements include a variable N-terminal variable region (yellow), pin motif (red), hook region (blue), scaffolding domain (green), spoke region (blue), β-strand (cyan), and C-terminal variable region (purple). For illustration purposes, elements are mapped into the structures of **(A)** two neighboring protomers from the cryo-EM based secondary structure model of human Cav1 8S complex (PDB: 7SC0), **(B)** two neighboring protomers from the 11-mer of human Cav1 (Q03135) predicted by AF2.2, and **(C)** two neighboring protomers from the 11-mer of *Caenorhabditis elegans* caveolin (Q94051) predicted by AF2.2. The comparison between the old domain segregation and our proposed segmentation using structural elements can be observed in both the model panel (A) and sequence alignments of human Cav1 and *C. elegans* caveolin Q94051 **(D)**.

Next, we asked how these structural elements are utilized by different caveolins and how they change during evolution (**Fig 4, 5**). To illustrate key similarities and differences across evolutionary distant caveolins we selected four examples taken from the major classes of caveolins: 1) human Cav1, a Type II-CAV; 2) a Type I-CAV from *Strongylocentrotus purpuratus* (A0A7M7T4C2); 3) an atypical-caveolin from *A. queenslandica* (A0A1X7UHP5); and 4) the Choa-caveolin from *S. rosetta* (F2U793) (**Fig 5B-N**). All four caveolin classes are predicted to contain an N-terminal variable region, hook structure, scaffolding domain, and spoke region (**Fig 5B-N**). Despite being essential for the formation of human Cav1 complexes, the pin motif is found only in Type II caveolins (**Fig 4, 5A**). The presence of the C-terminal β-strand also varied across clades. β-strands were predicted to be absent from 60% of the Atypical caveolins, including the representative *A. queenslandica* caveolin (**Figs 4, 5**), but were predicted to exist in an extended form in other clades. The occurrence of the C-terminal variable region was predicted to vary across caveolins, even within the same clade (**Fig 4**).

**Fig. 4.**
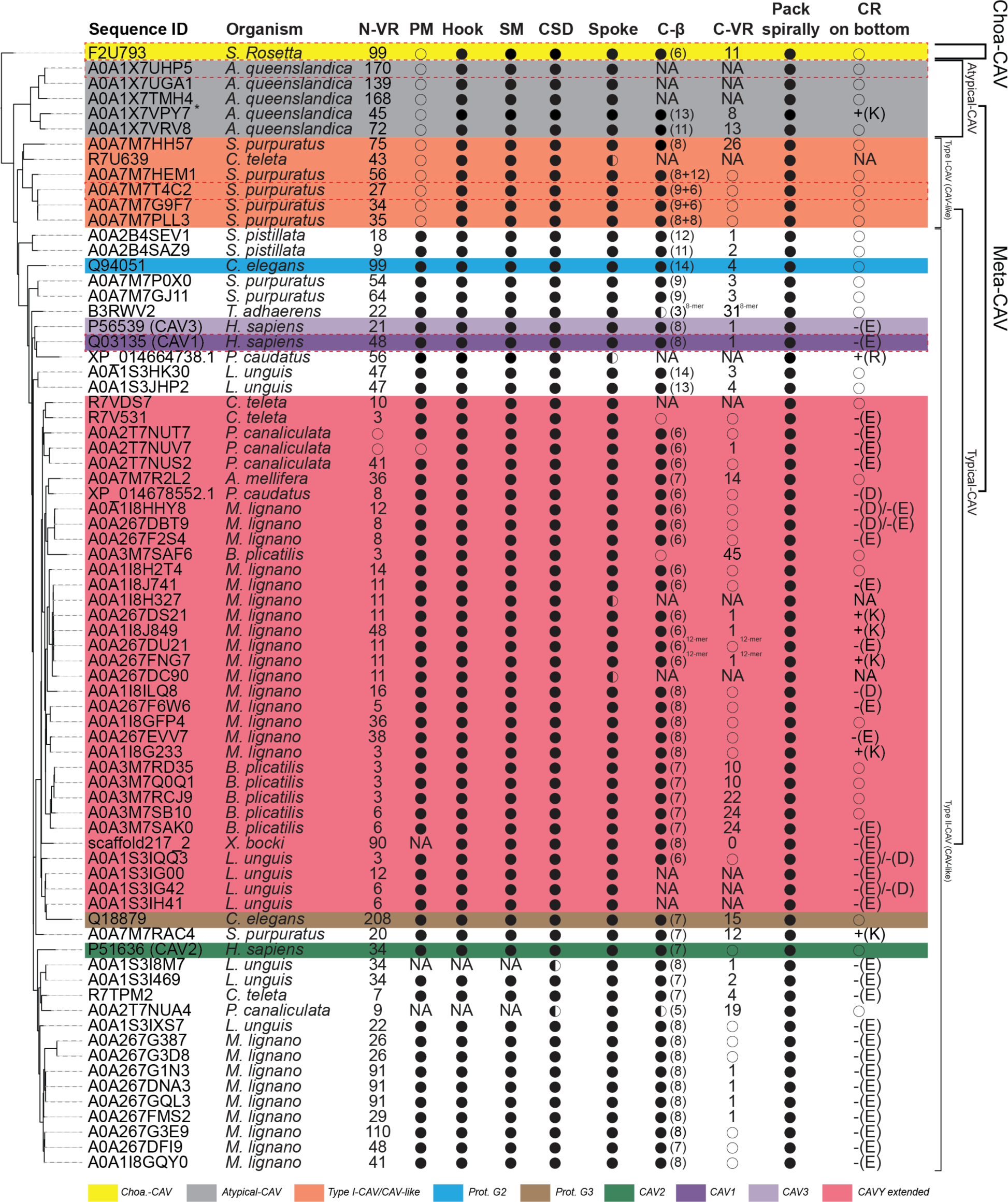
Summary of the structural features of caveolins suggested by computational modeling predictions. Phylogenetic tree shown on the left-hand side of the table is based on Figure S1. N-VR; N-terminus variable region; PM, pin motif; Hook, hook structure; SM, signature motif; CSD, scaffolding domain; Spoke, spoke region; C-β, C-terminus β-strand; C-VR, C-terminus variable region; Pack spirally, whether the protein sequence is predicted to be assembled into a disc-shaped oligomer; CR on bottom, charged residues on hydrophobic side of complexes (“+” represents a positive charge, “-” represents a negative charge, and uppercase English letters represent the corresponding amino acid residue abbreviations). Structural features were summarized mainly based on AF2.2 predicted 11-mers unless otherwise noted in the upper right corner of particular features. 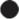, structural feature was predicted to be present; 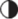, structural feature was predicted to be partially present; 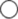, structural feature was not predicted to be present. NA, not applicable (corresponding sequences were either missing or shifted due to sequence missing). *, A0A1X7VPY7 was not predicted by AF2.2 to form any homologous disc-like or ring-like complexes. However, it was predicted to form a hybrid complex structure with A0A1X7VRV8. The structural features listed for A0A1X7VPY7 are summarized from the model of the hybrid complex consisting of A0A1X7VPY7 (1 copy) and A0A1X7VRV8 (10 copies).

**Fig. 5.**
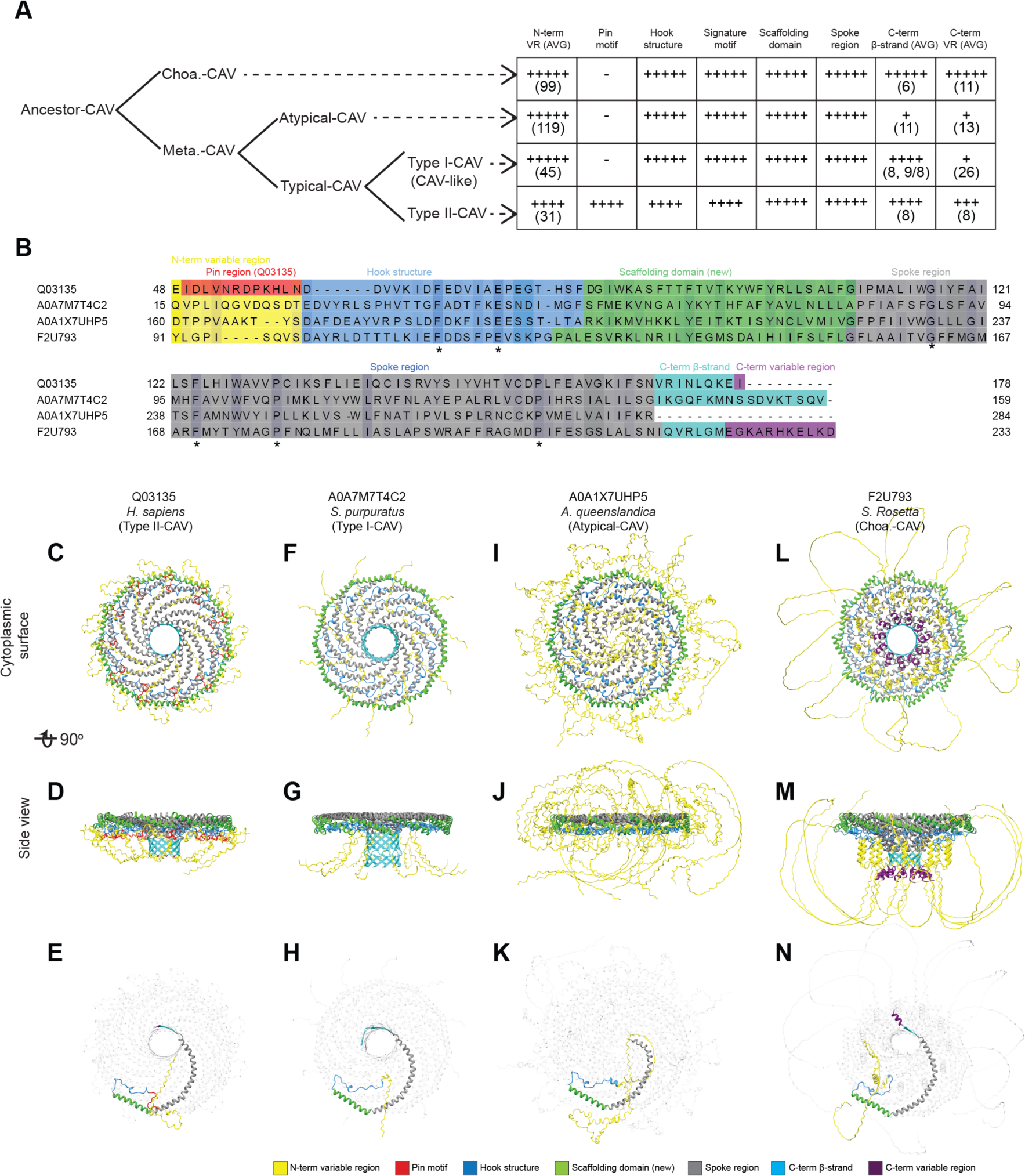
Predicted structural features of representative metazoan and non-metazoan caveolins. **(A)** Model summarizing the key structural similarities and difference in different groups of caveolins based on the phylogenetic analysis and structural comparisons presented in Table 1 and Supplemental File 2. **(B)** Sequence alignment of representative caveolins. Conserved residues are colored with different intensity according to the percent identity. The range of different structure regions were also highlighted with different colors. **(C-N)** Different views of AF2.2 models of representative caveolins**. (C-E)** Human Cav1 (Type II-CAV, Q03135). **(F-H)** *S. purpuratus* (Type I-CAV, A0A7M7T4C2). **(I-K)** *A. queenslandica* (Atypical-CAV, A0A1X7UHP5). **(L-N)** *S. rosetta* (Choa-CAV, F2U793). The range of different structure regions were also highlighted with different colors. To better display the structure of a single protomer within the complex, the ten protomers in the models in panels D, G, J, and M have been made transparent.

Finally, we examined the hydrophobic membrane facing surfaces. Although all the 74 caveolin complexes we examined form a disc with a hydrophobic face, approximately half of the complexes, including *S. rosetta* and *A. queenslandica* caveolins, have no charged residues on this face, whereas others, including human Cav1, contain a few charged residues that because of the symmetry of the complex form a charged ring circling the hydrophobic face (**Fig 4, Fig S2**).

### Electron microscopy (EM) shows diverse caveolins can form disc-shaped oligomers

Computational modeling is useful to generate hypotheses but requires experimental validation ^48^. To test key predictions from our evolutionary and computational modeling analyses, we used a combination of biochemistry and negative stain EM to examine the structure of members of four major classes of caveolins. Human Cav1, *S. purpuratus* (A0A7M7T4C2), *A. queenslandica* (A0A1X7UHP5), and *S. rosetta* (F2U793) sequences were expressed in *Escherichia coli* and purified in detergent using size exclusion chromatography (SEC). Western blotting of fractions eluted from SEC confirmed that these caveolins form high molecular weight complexes (**Fig S3**). To visualize their overall structure, we performed negative stain EM (**Fig 6**).

**Fig. 6.**
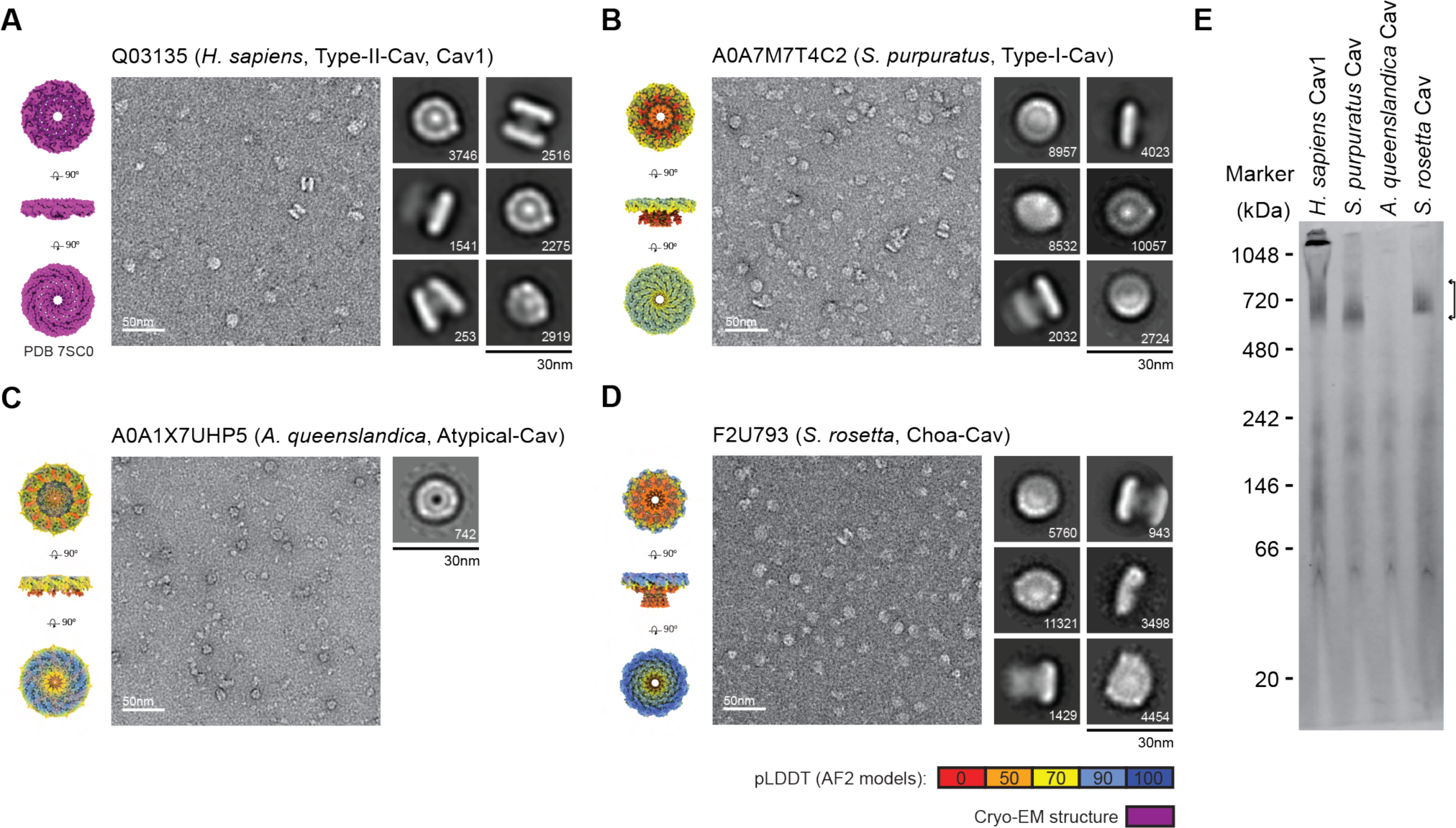
Negative stain EM shows diverse caveolins assemble into disc-shaped complexes. **(A)** *H. sapiens* Cav1, (**B**) *S. purpuratus* caveolin, (**C**) *A. queenslandica* caveolin, and (**D**) *S. rosetta* caveolin. (**A-D**) In each panel, surface-filling models for the cryo-EM structure or AF2.2 11-mer structures are shown on the left, representative images of negatively stained caveolin complexes are shown in the middle, and representative 2D averages of caveolins are shown on the right. The number of particles found in each class average is shown in the bottom right. Scale bar, 30 nm. (**E**) Blue Native gel with arrows highlighting the position of 8S-like complexes. MW markers are indicated.

Negative stain 2D class averages of the complexes from each purification were roughly the same size and most assumed a disc-like geometry although there was some heterogeneity in the shape of the discs (**Fig 6, Fig S4**). *S. purpuratus* and *S. rosetta* caveolins formed uniform discs with distinct inner and outer rings, similar to the appearance of human Cav1 in negative stain (**Fig 6A, B, D**). The *A. queenslandica* caveolin complex, although not as well-ordered, still appeared to form disc-like complexes, but lacked the central density observed in the human, *S. purpuratus,* and *S. rosetta* caveolin classes (**Fig 6C**). The *S. purpuratus* and *S. rosetta* caveolins migrated as stable complexes on blue native gels (**Fig 6E**). In contrast, the *A. queenslandica* caveolin complex was unstable, possibly due to its inability to form a β-barrel (**Fig 6E**). Taken together with our computational models, these results show members of all four classes of caveolins, even though evolutionarily distant, can assemble into disc shaped complexes, suggesting a conserved structural “fingerprint” for the caveolin family of proteins.

### Cryo-EM reveals the molecular basis for the assembly of S. purpuratus caveolin complex

We used single particle cryo-EM to determine a 3.1 Å resolution structure of the *S. purpuratus* caveolin complex and built a model spanning amino acids 29-152 (**Figs 7, S5, and S6**). Residues 1-28 and 153-159, regions predicted to be disordered (**Fig 4**), were not visible in the map. The complex, ∼130 Å in diameter and ∼36 Å in height (**Fig 7E**), is composed of 11 spiraling α-helical protomers organized into a disc with a protruding β-barrel (**Fig 7B, E**). The variable N-terminal region extends along the cytoplasmic face of the complex before making a 180° turn at the hook structure. Amphipathic α-helices form the spoke region, with the α-1 helix forming the rim of the disc that is ∼15 Å in height, and the C-terminus of each protomer is a β-strand that forms a central parallel β-barrel ∼30 Å in diameter (**Fig 7G, H**). An unstructured detergent micelle covers the hydrophobic face of the disc and reaches into the interior of the β-barrel, suggesting both regions interact with membrane (**Fig 7I**). No defined density was detected in the β-barrel even with no applied symmetry. Comparison of the experimental and AF2.2 structures shows the computational model did not correctly predict the length of the β-barrel or the curvature of the disc (**Fig 7J-L**).

**Fig. 7.**
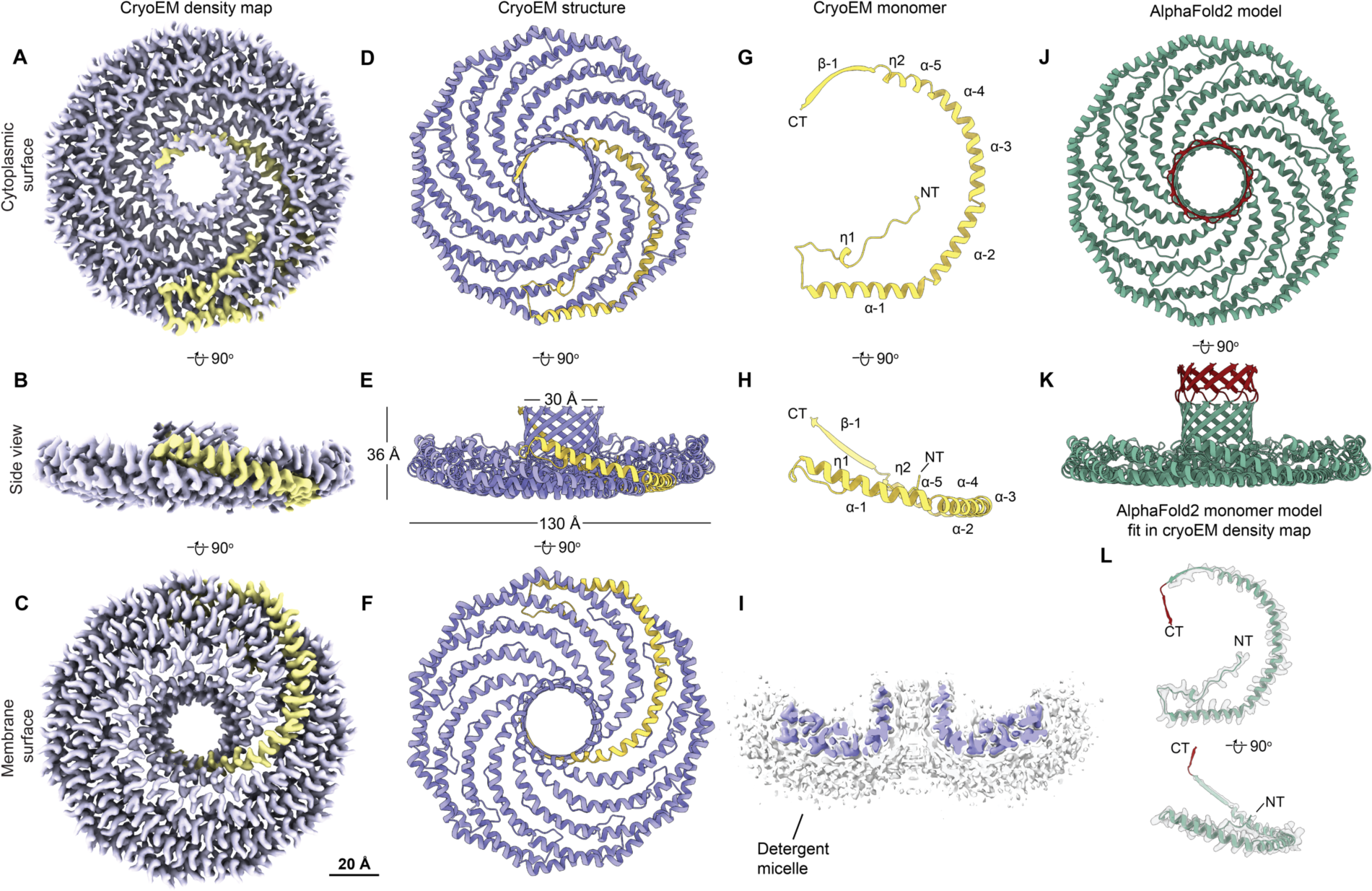
*S. purpuratus* caveolin forms an 11-mer complex. **(A-C)** A 3.1 Å resolution cryo-EM density map of the *S. purpuratus* caveolin complex with 11-fold symmetry. Complex is shown with ninety-degree rotated views displaying the cytoplasmic-facing surface (**A**), side (**B**), and membrane-facing surface (**C**). The complex has an overall disc-like structure with 11 spiraling α-helices and a central β-barrel. A single protomer is highlighted in yellow. Scale bar, 20 Å. (**D to F**) Secondary structure model of the *S. purpuratus* caveolin in the same views as shown in panels A-C. (**G and H**) Secondary structure of *S. purpuratus* caveolin protomer with secondary features and N- and C-termini noted. (**I**) Central slice of the density map (purple) with the detergent micelle (grey). (**J, K**) AF2.2-predicted structure of the *S. purpuratus* caveolin 11-mer showing views of the cytoplasmic-facing surface (J) and side view (K). Structured regions predicted by AF2.2 that are not found in the cryo-EM structure are highlighted in burgundy. (**L**) Protomer from the AF2.2 11-mer model fit into the cryo-EM density map (gray outline) of a protomer.

### Cryo-EM structure of the choanoflagellate S. rosetta caveolin complex

We next determined a 2.9 Å resolution cryo-EM structure of the *S. rosetta* caveolin complex (**Figs 8, S7, and S8**). A model of *S. rosetta* caveolin spanning amino acids 79-23 was built from the density map (**Fig 8D-F**). There was no density for the predicted disordered N-terminal region (a.a. 1-78) and the C-terminal residues (a.a. 232-233). The *S. rosetta* caveolin complex, ∼127 Å in diameter and ∼46 Å in height, is a disc composed of spiraling α-helices with a central β-barrel. The variable N-terminal region forms a short α-helix that is positioned about halfway up the β-barrel before snaking along the cytoplasmic facing surface of the complex. Each protomer makes a 180° turn at the hook structure, which is followed by the spoke region. The α-1 helix forms the rim at the edge of the disc that is ∼16 Å in height (**Fig 8G-H**). Finally, *S. rosetta* caveolin has a C-terminal β-strand with a height of ∼46 Å which forms a parallel β-barrel with a diameter of ∼30 Å (**Fig 8E**). The detergent micelle surrounds the hydrophobic face of the disc and reaches into the β-barrel (**Fig 8I**). As with the other caveolin complexes, no interpretable density was found in the β-barrel, even with no applied symmetry. For this complex, AF2.2 failed to predict the structure of the N- and C-terminal portions of *S. rosetta* caveolin or the correct curvature of the disc (**Fig 8E, J-L**).

**Fig. 8.**
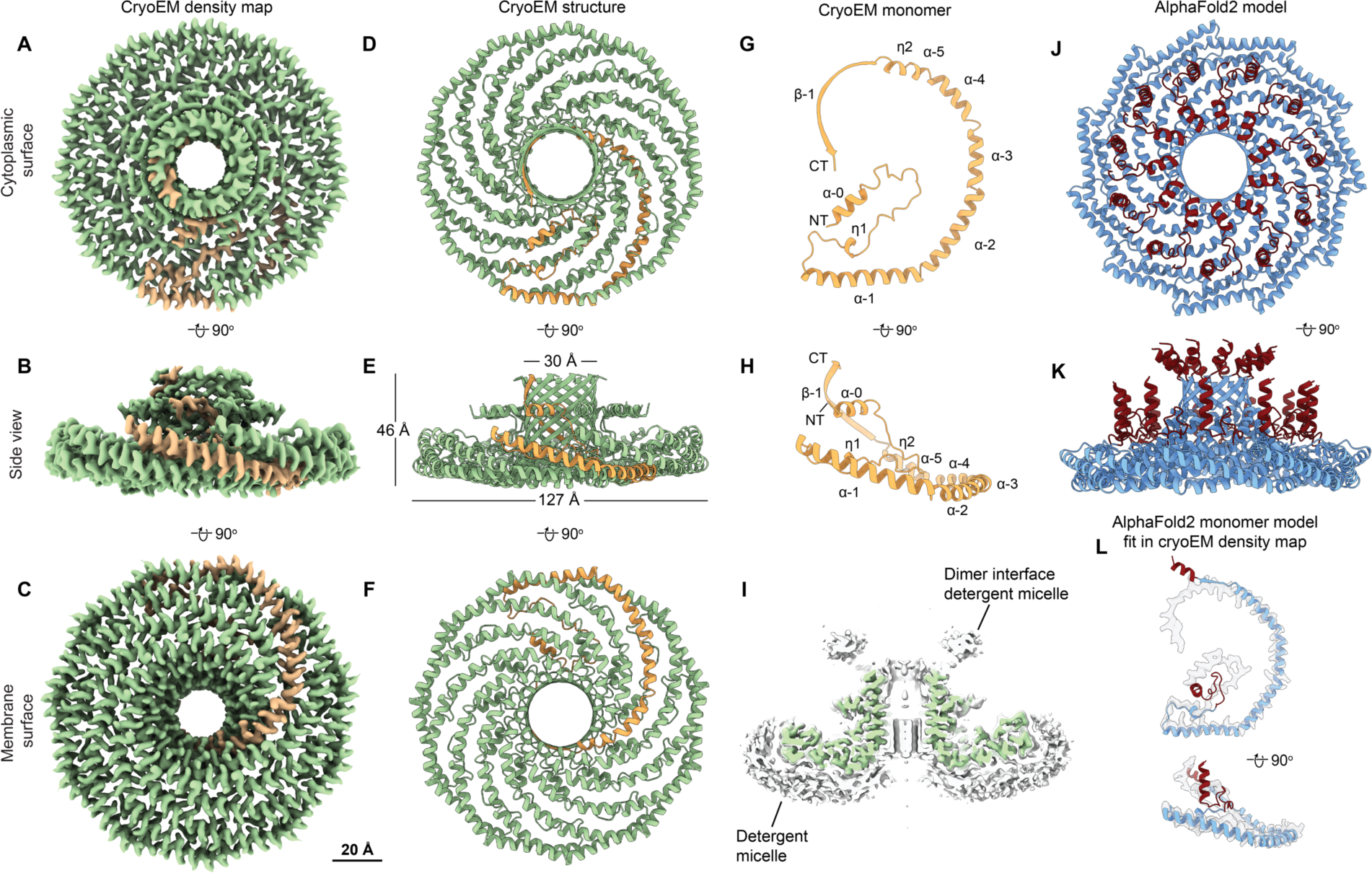
*S. rosetta* caveolin forms an 11-mer complex with elongated β-barrel and extended N-terminal region. (**A-C**) A 2.9 Å resolution cryo-EM density of *S. rosetta* caveolin complex with 11-fold symmetry. The complex is shown with ninety-degree rotated views displaying the cytoplasmic-facing surface (**A**), side (**B**), and membrane-facing surface (**C**). The complex has an overall disc-like structure with 11 spiraling α-helices, a central elongated β-barrel, and extended N-terminal region. A single protomer is highlighted in orange. Scale bar, 20 Å. (**D to F**) Secondary structure model of the *S. rosetta* caveolin complex in the same views as shown in panels A-C. (**G and H**) Secondary structure of *S. rosetta* caveolin protomer with secondary features and N- and C-termini noted. (**I**) Central slice of the density map (green) with the detergent micelle (grey). (**J, K**) AF2.2-predicted structure of the *S. rosetta* caveolin 11-mer showing views of the cytoplasmic-facing surface (J) and side view (K). Structured regions predicted by AF2.2 that are not found in the cryo-EM structure are highlighted in burgundy. (**L**) Protomer from the AF2.2 11-mer model fit into the cryo-EM density map (gray outline) of a protomer.

### Evolutionarily distant caveolins share structural motifs but differ in dimensions

We next directly compared the structures of the *S. purpuratus* and *S. rosetta* caveolin complexes with the previously determined human Cav1 complex (**Fig 9**). Despite sharing no significant sequence similarity with human Cav1 (16%/35% identity/similarity for *S. purpuratus* caveolin and 13%/28% for *S. rosetta* caveolin), the secondary structure of the protomers and organization of all three complexes are similar (**Fig 9**). Each complex forms a disc with 11 spiraling α-helices and a central β-barrel, with each protomer forming contacts with two protomers to the left and two to the right (**Fig S10**). The spoke regions and scaffolding domains that make up the discs contain a similar number of residues (88, 87, and 90 residues for human Cav1, *S. purpuratus* and *S. rosetta* caveolins, respectively). However, both the *S. purpuratus* and *S. rosetta* caveolin complexes are smaller in diameter (∼130 Å and ∼127 Å, respectively) compared to human Cav1 (∼140 Å). Although the *S. purpuratus* and *S. rosetta* caveolin complexes exhibit tighter packing of the η1 and α1 helices than the human Cav1 complex, this packing loosens towards the rim of the complexes. The difference in diameter between the complexes is due to the increased curvature of the *S. purpuratus* and *S. rosetta* caveolin complexes compared to the human Cav1 complex. The three β-barrels have a similar outer diameter of ∼28-30 Å; however, the *S. rosetta* caveolin complex’s central β-barrel is ∼10 Å longer than the other caveolin complexes (**Fig 9B, E, H**).

**Fig. 9.**
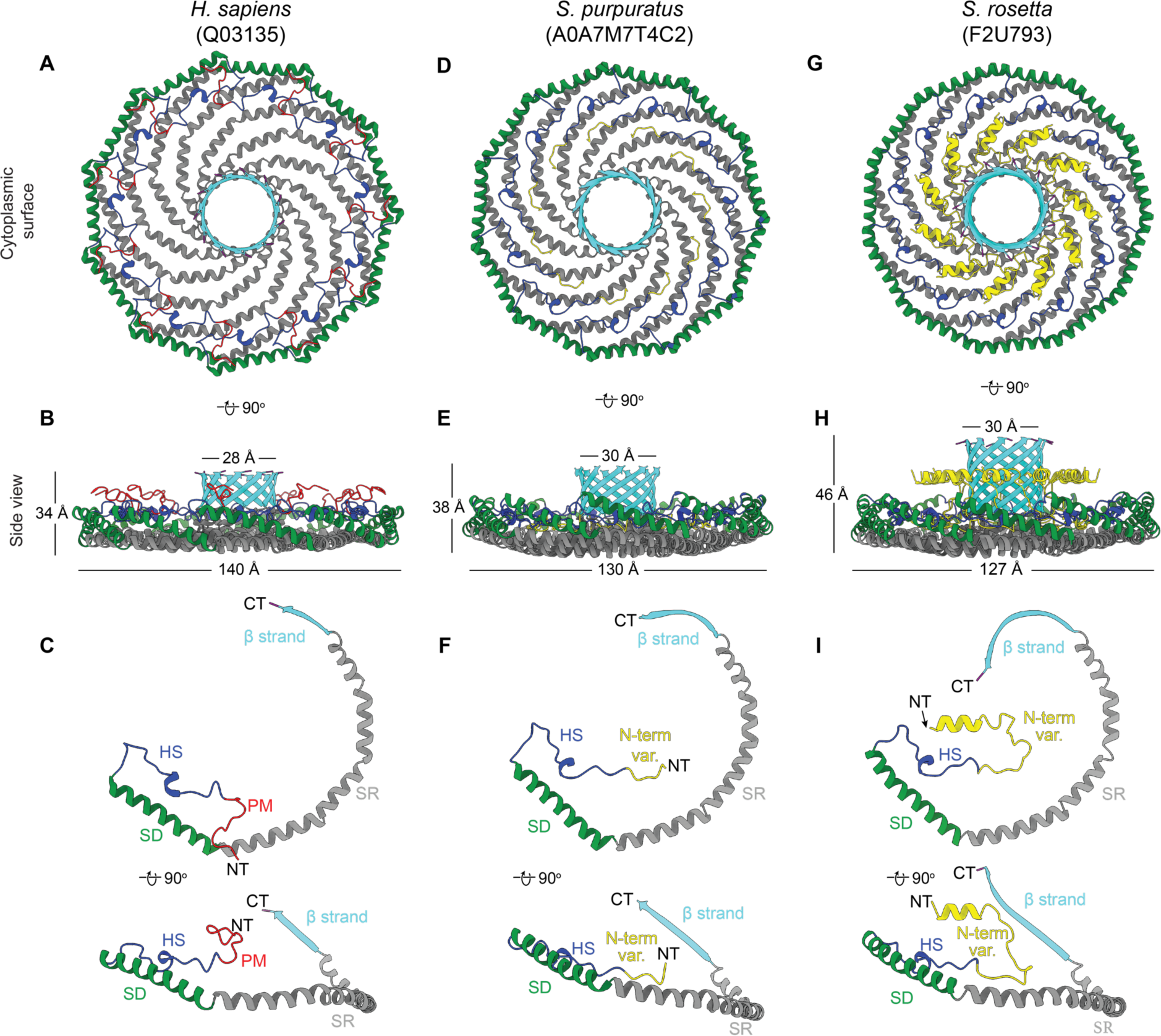
Comparison of cryo-EM structures for *H. sapiens*, *S. rosetta,* and *S. purpuratus* caveolin complexes. **(A, B, D, E)** Cytoplasmic-facing surface and side views of cryo-EM structures are shown for human Cav1 (**A, B**), *S. purpuratus* caveolin (**D, E**), and *S. rosetta* caveolin (**G, H**). (**C, F, I**) Protomer structures extracted from the 11-mers are shown in same views as panel above. Domains are colored as follows: N-terminal variable region (yellow), pin motif (PM, red), hook structure (HS, blue), scaffolding domain (SD, green), spoke region (SR, gray), β-strand (cyan). NT, N-terminus; C-term, C-terminus.

While the overall organization of the three caveolin complexes are similar, their N-terminal structured regions differ significantly. In human Cav1, the pin motif interacts with other protomers along the rim region and then is directly followed by the hook structure. However, the *S. purpuratus and S. rosetta* caveolins do not have pin motifs. In *S. purpuratus* caveolin, the N-terminal region extends outwards from the middle of the hydrophilic α-helical face until it reaches the hook structure (a.a. 35-55) where it makes a ∼180° turn (**Fig 9**). In contrast, for *S. rosetta* caveolin the N-terminal region forms a short α-helix (a.a. 79-88) about halfway up the C-terminal β-barrel and extends parallel to the disc on the presumed cytoplasmic side of the complex (**Fig 9G, H**). As a result, in the *S. rosetta* complex additional contacts are made with the *i* + 5 and *i* - 5 protomers as the variable N-terminal region of *i* rises up the central β-barrel and contacts the C-terminal β strands of *i* + 5 and *i* – 5 (**Fig S10**).

### S. purpuratus and S. rosetta caveolin complexes form amphipathic discs with increased curvature compared to the human Cav1 complex

A key feature of human Cav1 8S complex is the amphipathic nature of the disc ^29^ (**Fig S11A, D**). The *S. purpuratus* and *S. rosetta* caveolin complexes likewise contain distinct hydrophobic and hydrophilic faces (**Fig S11B, C, E, F**). In contrast to the ring of glutamic acid residues on the hydrophobic face of the human Cav1 complex, the hydrophobic surface of *S. purpuratus* and *S. rosetta* caveolins lack any charged residues (**Fig S11B, C**). The hydrophilic faces of the complexes have an array of charged, but not conserved, residues (**Fig S11A-C**). Finally, the interior of the conserved central β-barrel is hydrophobic in all three complexes. Only the human Cav1 β-barrel is capped by a charged residue (Lys176) (**Fig S11D-F**).

The membrane facing surface of the human Cav1 8S complex is flat ^29^ (**Fig S11B**). In contrast, both the *S. purpuratus* and *S. rosetta* caveolin complexes are concave, with curvatures of ∼17° and ∼11°, respectively (**Fig S11E, H**). 2D averages of vitrified *S. purpuratus* caveolin complexes showed variations of curvature, including examples of complexes with either concave or convex curvatures (**Fig S12A-D**). While we were unable to determine high-resolution structures from these averages, 3D variation analysis (3DVA) of the *S. purpuratus* caveolin complex results in components that capture a range of continuous complex conformations (**Movie 1**). Calculated using a filter resolution of 8Å, these low-resolution structures represent the negative and positive values along the reaction coordinate for the components with the largest variance (**Fig S12E**). This analysis shows that the *S. purpuratus* caveolin disc can be concave, similar to the 3.1 Å structure, or can be flat. The differences in curvature are accommodated by the spoke region rising in pitch towards the center of the complex, which leads the β-barrel to be pushed “outwards” ∼4-5 Å towards what would be the cytoplasm (**Movie 1**). The 2D averages from the *S. rosetta* caveolin or human Cav1 samples did not show classes with different curvatures ^29^ and 3DVA did not identify components displaying significant variations in the shape of the discs. Thus, we conclude that the *S. purpuratus* caveolin complex is more flexible than the human or *S. rosetta* caveolin complexes under these experimental conditions.

## Discussion

Using a combination of computational and structural approaches, we now show that caveolins across evolution share the ability to assemble into amphipathic disc-shaped multimers composed of spiraling α-helices and a central protruding parallel β-barrel. Somewhat surprisingly considering the ability of AF2 to build disc-shaped complexes with different numbers of protomers, all three of the experimentally determined cryo-EM structures have 11-fold symmetry. While our analyses reveal many conserved features of caveolin oligomers, they also uncover striking differences such as highly variable N- and C-termini and variations in the curvature of the membrane-facing surface of the disc. These findings suggest that caveolins adopt a conserved structural framework built around an amphipathic disc, but are adaptable enough to accommodate significant molecular variations.

Classically, caveolins have been depicted as consisting of several major domains including a signature motif, scaffolding domain, oligomerization domain, intramembrane domain, and C-terminal region. Based on our structural and modeling results, we propose a new domain nomenclature, consisting of N-terminal variable region, pin motif, hook structure, scaffolding domain, spoke region, β-strand, and C-terminal variable region. Of these elements, the hook structure, scaffolding domain, and spoke region are found in almost every clade that was examined making them the most conserved structural features across caveolins. These regions of the protein contribute to oligomerization as well as help define the hydrophobic membrane facing surfaces of the complex. The β-barrel also contributes to the proper packing of protomers into regular disc-shaped complexes ^49^. Interestingly however, not all caveolins are predicted to have β-strands that contribute to β-barrels, such the *A. queenslandica* caveolin studied here. This could explain why the *A. queenslandica* caveolin complexes are less regular and stable than those formed by caveolins capable of assembling central β-barrels. Whether the β-barrels fulfill additional physiological roles beyond complex organization and structural stability remains to be determined.

Other domains of caveolins are less conserved across clades. While the pin motif appears essential for Type II-CAVs, such as the human Cav1 complex ^29^, it is probably the latest structural unit formed during the evolutionary process and is not essential for caveolins from other evolutionary branches to pack into oligomeric complexes. The length and composition of the N- and C-terminal unstructured regions are also highly variable across caveolins, suggesting that they may be highly tuned for specific functions in various organisms. Because both regions are expected to project into the cytoplasm, we speculate that they could represent a binding site for cytosolic proteins. We also noted that a subset of caveolins, including human Cav1, contain charged residues on the membrane facing surface. The functional significance of these charged residues is not yet clear but could potentially impact lipid packing around the complex ^31^.

We also found that caveolin-related proteins are not limited to Metazoa; they are also found in the choanoflagellate *S. rosetta.* This implies that the proteins evolved independently in the two lineages for the past billion years if the protein was present in the ancestor of Choanozoa, or the past 600 million years in a scenario of horizontal gene transfer to the ancestor of the Myriazoa (*5*). This raises the interesting question of what functional roles caveolins fulfill in choanoflagellates and how these relate to their functions in mammals. A potential clue is that several binding partners and signaling pathways that caveolins have been linked to in mammals are also found in choanoflagellates ^50, 51, 52^. They could also help buffer changes in membrane tension by a caveolae-independent mechanism ^53^. As experimental approaches to study the cell biology of choanoflagellates continue to advance ^54^, it should be possible to test the structure-function relationship of this evolutionarily distant form of the protein in the future.

The unexpected identification of a choanoflagellate caveolin also prompted us to examine the evolutionary history of caveolins more deeply. In addition to performing conventional phylogenetic analysis, we examined the chromosome-scale evolutionary history of caveolins from ALGs. This powerful approach has recently been employed at the chromosome scale to resolve the long-argued question of the evolutionary relationship between sponges, ctenophores, and other animals ^27^. Here, we traced ALG-caveolin colocalizations to track the chromosomal origins of caveolins in animals and infer the relationship between animal and choanoflagellate caveolins. The consistent presence of caveolin orthologs on chromosomes containing the Eb ALG suggests that the ancestral caveolin, which gave rise to all non-sponge caveolins, was present on ALG Eb in the ancestor of Parahoxozoa, dating its critical biological role to before the Cambrian explosion. The persistence of these sequences on homologous chromosomes, even in light of lineage-specific chromosomal changes, is similar to the conservation of biologically critical genes on single ALGs, such as the persistence of the Hox cluster on ALG B2 in parahoxozoans ^35, 55^. Future work to test whether this is the result of cis-regulatory constraints ^56^ may reveal yet-undiscovered loci important to caveolin biology.

There are several limitations to our study. First, while AF2 is able to predict the basic secondary structure and the overall organization of caveolins into spirally packed discs, it does less well in predicting the structure of the N- and C-terminal domains of *S. purpuratus* and *S. rosetta* caveolins. Second, how the oligomeric state of caveolin complexes is controlled is uncertain. The experimental structures of human Cav1, *S. purpuratus* caveolin, and *S. rosetta* caveolin are all 11-mers. However, AF2 predicts additional oligomeric states can exist in addition to the experimentally observed 11-mers (although some are less likely to exist due to energetic strains) ^42^. It is possible that the oligomerization state of the complexes may be influenced by expression and purification conditions of our experiments. Finally, the functional consequences of the structural differences between caveolins remain to be determined. As one example, the differences in curvature in the membrane facing surface observed in the cryo-EM structures of human, *S. purpuratus*, and *S. rosetta* caveolins could indicate they are optimized to bend membranes into different shapes and/or undergo conformational changes in response to changes in membrane tension.

In summary, we conclude that that the ability of caveolins to assemble into amphipathic discs represents an ancient, unique mode of protein-lipid membrane interactions that predates the emergence of metazoans. Given these new insights, it should now become possible to uncover how caveolins control cellular function at a molecular level across an evolutionary scale.

## Materials and Methods

### AlphaFold2 Predictions

Predicted structures of caveolin homo-oligomers were generated using AF2.1 or AF2.2 by systematically increasing the number of monomer input sequences until the upper limit of the residues that could be analyzed was reached. AlphaFold v2.1.0 predictions were performed using a Colab notebook named “AlphaFold2_advanced (<u>AlphaFold2_advanced.ipynb - Colaboratory</u> (google.com)” with default settings. AlphaFold v2.2.0 predictions were performed using default settings via another Colab notebook named “alphafold21_predict_colab” provided by ChimeraX daily builds version (ChimeraX 1.4.0). Version v2.2.0 includes updated AlphaFold-Multimer model parameters. See https://github.com/deepmind/alphafold/releases for a description of other differences in AF2.1.0 versus AF2.2.0. Due to the upper limit in the number of residues that could be analyzed by AF2.1, where indicated, caveolin sequences were truncated to exclude the predicted N-terminal disordered regions. Unless otherwise stated, the rank model 1 of the 5 models output for each prediction is shown. Confidence levels of the predictions are rendered on the models using pLDDT (predicted local-distance difference test) values ^41^.

### Conservation analysis

Conservation analysis was based on protein sequences alignments collected from UniProt of 72 metazoan caveolin sequences from 13 representative species of different phylum/superphylum via EMBL-EBI online tool MAFFT (Multiple Alignment using Fast Fourier Transform) (<u>MAFFT <</u> <u>Multiple Sequence Alignment < EMBL-EBI</u>) and displayed on 3D structure by using Chimera v1.15. The figures were analyzed, rendered, and exported with Chimera v1.15, ChimeraX 1.4.0 (daily builds version), or ChimeraX1.3.

### Sequence alignments

Clustal Omega was used to perform sequence alignments. Jalview 2.11.2.4 ^57^ was used for alignment image typesetting and exporting.

### HMMER searches

For HMMER searches, a hidden Markov model (HMM) profile was built directly from a MAFFT alignment of previously reported truncated caveolin sequences ^3^ using hmmbuild from the HMMER package v 3.1b2 ^58^. The profile was then searched against the *S. rosetta* ATCC 50818 NCBI reference genome and *A. queenslandica* genome v1.1 from EnsemblMetazoa using hmmsearch. To identify putative ctenophore caveolins we used blastp v2.10.0+ ^59^ with the human protein sequences as queries, and HMMer with the above models to query the *Hormiphora californensis* ^60^ and *Bolinopsis microptera* ^27^ genomes (GCA_020137815.1 and GCF_026151205.1) and transcriptomes (TSA GHXS00000000 and GCF_026151205.1 annotation).

### Phylogenetic analyses

The selected caveolin homologues were aligned with MAFFT v7.310 ^61, 62^. The alignment was then truncated to a region corresponding to the residues 54–158 in human CAV1 using a simple Python script (https://doi.org/10.5281/zenodo.6562402). Gaps were removed before combining the sequences with those from the supplementary information of Kirkham et al (2008) ^3^ and re-aligning with MAFFT. ProtTest3 (version 3.4.2) ^63^ was then used to determine the best model of evolution (LG+I+Γ). Finally, a maximum likelihood phylogeny was inferred using RAxML (version 8.2.11; ^64, 65^ with 1000 rapid bootstraps. The trimmed tree was produced in RStudio using the *ape* package ^66^. Trees were rendered using FigTree v1.4.4 and prepared for publication using Inkscape (version 1.0.1).

### Caveolin Ancestral Linkage Group Analysis

Ancestral linkage analysis (ALG) was carried out using previously described methods ^27^ using chromosome-scale animal genomes obtained from *Acropora millepora* (GCF_013753865.1), *Asterias rubens* (GCF_902459465.1), *Biomphalaria glabrata* (GCF_947242115.1), *Branchiostoma floridae* (GCF_000003815.2), *Dreissena polymorpha* (GCF_020536995.1), *Ephydatia muelleri* ^67^, *Gallus gallus* (GCF_016699485.2), *Holothuria leucospilota* (GCA_029531755.1), *Homo sapiens* (GCF_000001405.40), *Lethenteron reissneri* (GCF_015708825.1), *Lytechinus pictus* (GCF_015342785.2), *Lytechinus variegatus* (GCF_018143015.1), *Mercenaria mercenaria* (GCF_021730395.1), *Mya arenaria* (GCF_026914265.1), *Nematostella vectensis* (GCF_932526225.1), *Oscarella lobularis* (GCF_947507565.1), *Patella vulgata* (GCF_932274485.2), *Pecten maximus* (GCF_902652985.1), *Petromyzon marinus* (GCF_010993605.1), *Pomacea canaliculata* (GCF_003073045.1), *Rhopilema esculentum* ^68^, and *Salpingoeca rosetta* (GCA_033442325.1) (**Supplementary File 1**). To assess the ALG homology with these chromosomal sequences, we used the snakefile ‘odp’ from the odp software v.0.3.3 ^27^ with the option ‘ignore_autobreaks: True’, ‘diamond_or_blastp: “diamond”’, ‘duplicate_proteins: “pass”’, ‘plot_LGs: True’, and ‘plot_sp_sp: False’. We used the resulting .rbh files and .pdfs to identify significant (*p* ≤ 0.05, one-sided Bonferroni-corrected Fisher’s exact test) ALG homologies. To identify caveolin orthologs in each genome we searched the annotation keywords for caveolin, and verified the putative identities using blastp v.2.16.0+ ^59^ and hmmer v.3.4 ^69^. TimeTree v.5 ^70^ was used to generate a figure of species divergence times, although the animal topology was adjusted after recent studies ^5, 27^.

### Expression and purification of recombinant caveolins in *E. coli*

Protein expression and purification were performed following previously described protocols with minor modifications ^49^. Genes encoding the entire caveolin polypeptide of *A. queenslandica* (UniProt Accession A0A1X7UHP5), *S. purpuratus* (UniProt Accession A0A7M7T4C2) and *S. Rosetta* (UniProt Accession F2U793) were synthesized by Genscript, NJ and subcloned into the pET20b(+) vector (Novagen) using NdeI and XhoI (New England Biolabs, MA) restriction sites. The primers used (Integrated DNA Technologies, IA) were as follows: *A. queenslandica* forward gcggccCATATGCCTCCACCCCCTCCCCCG and reverse gcggccCTCGAGtcttttgaagataatggccacag; *S. purpuratus* forward gcggccCATATGGAACTGATCCATCCTG and reverse gcggccCTCGAGcacctggctggtcttaacatcagagc and *S. rosetta* forward gcggccCATATGAGCTACCAC and reverse gcggccCTCGAGgtccttcagttccttgtgg. Resulting plasmids were verified by Sanger sequencing (Genewiz/Azenta Life Sciences, NJ). Caveolin proteins were expressed in *E. coli* BL21 using the auto-induction expression system ^49^. Initially, an MDG starter culture of bacteria was incubated at 37°C and 250 rpm for 20 hours. Subsequently, the culture was expanded using auto-inducing ZYM-5052 media at 25°C and 300 rpm for 24 hours. The *E. coli* cells were then washed with 0.9% NaCl and resuspended in a buffer containing 200 mM NaCl and 20 mM Tris–HCl, pH 8.0. Bacterial cells were homogenized using a French press pressure homogenizer, with 1 mM PMSF and DTT added immediately before homogenization. To remove large cell debris, the homogenate was centrifuged at 9000 rpm for 15 minutes at 4°C. Total membranes were subsequently pelleted by centrifugation at 40,000 rpm (Ti-45 rotor, Beckman Coulter) for 1 hour at 4°C. The membrane pellets were then homogenized using a Dounce tissue grinder in a buffer consisting of 200 mM NaCl, 20 mM Tris–HCl (pH 8.0), and 1 mM DTT. To solubilize caveolin proteins from the membranes, a 10% n-Dodecyl-β-D-Maltopyranoside (C12M) (Anatrace) stock solution was added to the membrane homogenate to a final concentration of 2%, and the mixture was gently stirred for 2 hours at 4°C. Insoluble material was removed by centrifugation at 42,000 rpm (Ti-50.2 rotor) for 35 minutes, and the supernatant was subjected to Nickel Sepharose affinity purification. After washing the resin using 8 column volumes of a buffer composed of 200 mM NaCl, 20 mM Tris–HCl, pH 8.0, 1 mM DTT, 0.05% C12M, and 60 mM imidazole, the protein was eluted by increasing the imidazole concentration to 300 mM. The eluate containing caveolin was concentrated and further purified by size-exclusion chromatography using a Superose 6 Increase 10/300 GL column (GE Healthcare) in a buffer containing 200 mM NaCl, 20 mM Tris–HCl (pH 8.0), 1 mM DTT, and 0.05% C12 M.

### Negative stain EM and data processing

Caveolin samples were prepared for negative stain EM using established methods ^71^. Briefly, a PELCO easiGlow™ glow discharge unit (Fresno, CA, USA) was used to glow discharge 200-mesh copper grids covered with carbon-coated collodion film (EMS, Hatfield, PA, USA) for 30 s at 10 mA. Evolutionary caveolin samples (3.5µL) were adsorbed to the grids and incubated for 1 minute at room temperature. Samples were first washed with two drops of water and then stained with two drops of 0.7% (w/v) uranyl formate (EMS, Hatfield, PA, USA). The samples were then blotted until dry. A Morgagni transmission electron microscope operated at an accelerating voltage of 100 kV (Thermo Fisher Scientific, Waltham, MA, USA) was used to image the samples at a nominal magnification of 22,000x (2.1 Å per pixel).

Negative stain EM datasets were collected with a Tecnai Spirit T12 transmission electron microscope operated at 120 kV (Thermo Fisher Scientific, Waltham, MA, USA). Images were collected at a nominal magnification of 30,000x (2.086 Å per pixel). Data were collected using SerialEM v4.0.8 software ^72^ on a 4 k x 4 k Rio camera (Gatan, Pleasanton, CA) with a −2.2 µm defocus value. All image processing was performed in Relion 4.0.0 ^73^. Approximately 1,000 particles were manually selected and 2D classified, and clear resulting classes were selected and used as references for particle picking on all micrographs. Particles were extracted with a 144-pixel box size (30 nm x 30 nm boxes). The extracted particles were 2D classified into 20 classes. The human Cav1, *S. purpuratus* caveolin, *A. queenslandica* caveolin, and *S. rosetta* caveolin datasets had 39,786, 124,021, 66,856, and 94,597 particles respectively.

### Cryo-EM sample preparation

For single particle cryo-EM, 4 µL of the protein sample (*S. purpuratus* caveolin: ∼0.02 mg/mL; *S. rosetta* caveolin: ∼0.04 mg/mL) was applied to a Quantifoil R 2/2, 200-mesh Cu grid with an ultrathin carbon layer (Electron Microscopy Services) that was glow-discharged for 30 s at 5 mA. Following a 30 s incubation, the sample was removed, and another 4 µL of protein sample was applied to the same grid. After a second 30 s incubation, the sample was blotted for 5 s with a blot force of 10 and was then plunge frozen in a slurry of liquid ethane using a Vitrobot Mark IV (Thermo Fisher Scientific). The chamber was kept at 4°C with 100% humidity.

### Cryo-EM data collection

Micrographs of the *S. purpuratus* caveolin sample were collected on a Titan Krios transmission electron microscope (Thermo Fisher Scientific) operated at 300 kV and equipped with a K3 direct detection camera with a BioQuantum energy filter used with a slit width of 20 eV (Gatan). Images were collected at a nominal magnification of 81,000 x (1.11 Å/ pixel). Two datasets were collected of the *S. purpuratus* caveolin sample with 10,127 and 12,803 micrographs using SerialEM v4.0.3 software with a total dose of 60 e-/Å2 and a defocus range of −0.5 µm to −3 µm ^72^.

Images of the *S. rosetta* caveolin sample were collected on a Titan Krios transmission electron microscope (Thermo Fisher Scientific). The microscope was operated at 300 kV and equipped with a K3 direct detection camera with a BioQuantum energy filter used with a slit width of 20 eV (Gatan). Images were collected at a nominal magnification of 105,000 x (0.87Å/ pixel). A total of 18,417 micrographs were collected using SerialEM v4.0.3 software with a total dose of 59.17 e-/Å2 and a defocus range of −0.5 µm to −3 µm ^72^.

### Image processing of *S. purpuratus* caveolin complex

All image processing, 2D classification, and 3D refinements were performed in cryoSPARC v4.2.1 and v4.3.1 ^74^. Two independently collected datasets of 10,127 and 12,803 movies respectively (22,930 total movies) were corrected for local-beam induced drift using patch motion correction. Patch CTF estimation was used to estimate the local CTF parameters. Following exposure curation to keep only micrographs with a CTF estimation of ≤5Å, 22,845 total micrographs were subjected to circular blob picking. After blob picking, 13,044,185 initial picks were extracted in 352-pixel^2^ boxes (390.72 Å x 390.72 Å). Iterative 2D classification resulted in 200,411 particles. These particles were input into a two-class *ab initio* 3D reconstruction. The 135,462 particles that contributed to the reconstruction yielding a better-defined caveolin complex were subjected to a further round of 2D classification, resulting in 80,245 particles. These particles were used for another two-class *ab initio* 3D reconstruction. The 44,388 particles contributing to the better resolved *ab initio* reconstruction were input into a single class *ab initio* 3D reconstruction with higher initial and maximum resolution parameters. The resulting *ab initio* 3D reconstruction and the 44,388 particles contributing to it were used as inputs for C1 and C11 non-uniform refinements. The C1 non-uniform refinement resulted in a map with a resolution of 6.6Å, and the C11 non-uniform refinement produced a map with a resolution of 3.2Å. Local masks were generated in ChimeraX and Relion 4.0 to mask out the detergent micelle ^73, 75^. These masks were then used to run local refinements with C1 and C11 symmetry. The C1 refinement resulted in a map of 6.2 Å resolution, and the C11 refinement reached a final resolution of 3.1 Å. All 3DVA was carried out in cryoSPARC v4.3.1 ^76^.

### Image processing of *S. rosetta* caveolin complex

18,417 movies were motion corrected and CTF estimated using Warp ^77^. Particles were picked using the BoxNet2 centralized neural network (CNN) in Warp, and 1,488,129 initial particles were extracted with a box size of 300-pixel^2^ (261Å x 261Å). Iterative 2D classification of particles in cryoSPARC v4.2.1 resulted in 66,202 particles ^74^. These particles were then input to a single-class *ab initio* 3D reconstruction that resulted in a reconstruction exhibiting 11 spiraling α-helices. The 66,202 particles contributing to the *ab initio* reconstruction were used for a non-uniform refinement with C1 symmetry that resulted in a map with a resolution of 3.8 Å. Imposing C11 symmetry for non-uniform refinement of the same particles resulted in a 3.4 Å map. Using ChimeraX and Relion 4.0, masks were generated to mask out the detergent micelle ^73, 75^. Local refinement using these masks resulted in a C1 refinement with a final resolution of 3.0 Å and a C11 refinement with a final resolution of 2.9 Å.

### Model building, refinement, and validation

Models of the *S. purpuratus* and *S. rosetta* caveolin proteins were built using ModelAngelo with the respective sequences input into the program ^78^. The output models resulting from ModelAngelo were then further refined using ISOLDE within ChimeraX and Phenix real-space refinement ^75, 79, 80^. The final model of *S. purpuratus* caveolin included residues 29-152, and the model of *S. rosetta* caveolin contained residues 79-231. Both models were validated within Phenix (**Table S2**) ^80^. Maps and models are deposited in the EMDB and PDB (*S. purpuratus* caveolin complex*: EMD-XXXXX, PDB-XXXXX; S. rosetta* caveolin complex*: EMD-XXXXX, PDB-XXXXX)*.

## Acknowledgments

We thank Mr. Fuchang Han from Kaifeng City for the schematic diagrams of different species for Figure 1, Yelena Peskova for expert technical assistance, and Drs. Hassane Mchaourab, Itay Budin, and Steven Haddock for helpful discussions. The U-M Cryo-EM Facility has received generous support from the U-M Life Sciences Institute, the U-M Biosciences Initiative, and Beckman Foundation. Computational resources provided by the Life Science Compute Cluster (LiSC) of the University of Vienna are also gratefully acknowledged. The content is solely the responsibility of the authors and does not necessarily represent the official views of the National Institutes of Health.

## Funding

National Institutes of Health grant R01 GM151635 (AKK)

National Institutes of Health grant R01 HL144131 (AKK, MDO)

National Institutes of Health grant R01GM080403 (JM)

National Institutes of Health grant R01HL122010 (JM)

National Institutes of Health grant R01GM129261 (JM)

National institute of Health grant S10OD030275 (MDO)

National Institute of Health T32GM007315 (SC)

European Research Council’s Horizon 2020 grant No. 945026 (DTS)

American Heart Association grant 905705 (SC)

University of Michigan Rackham Predoctoral Fellowship (SC)

Humboldt Professorship of the Alexander von Humboldt Foundation (JM)

HHMI (to Jochen Zimmer)

## Author contributions

Conceptualization: BH, AKK

Methodology: BH, AG, LFLW, SC, DTS, MDO

Investigation: BH, LFLW, SC, DTS

Data curation: BH, SC, LFLW, DTS

Formal analysis: BH, LFLW, SC, DTS, MDO

Writing—Original draft: BH, AKK, LFLW, SC, MDO

Writing—Review and editing: BH, LFLW, SC, DTS, AG, JM, EK, AKK, MDO

Visualization: BH, LFLW, SC, DTS

Supervision: AKK, JM, EK, MDO

Project administration: AKK

Funding acquisition: AKK, JM, MDO

## Data availability statement

Maps and models are deposited in the EMDB and PDB (*S. purpuratus* caveolin complex*: EMD-XXXXX, PDB-XXXXX; S. rosetta* caveolin complex*: EMD-XXXXX, PDB-XXXXX)* and will be made available upon publications. All other data needed to evaluate the conclusions in the paper are present in the paper and/or the Supplementary Materials.

