## Supplementary Figures for "Evolutionarily diverse caveolins share a common structural framework built around amphipathic discs"

**This PDF file includes:**

Figs. S1 to S12

Tables S1 to S2

**Other Supplementary Materials for this manuscript include the following:**

Movie S1

Data files S1 to S3

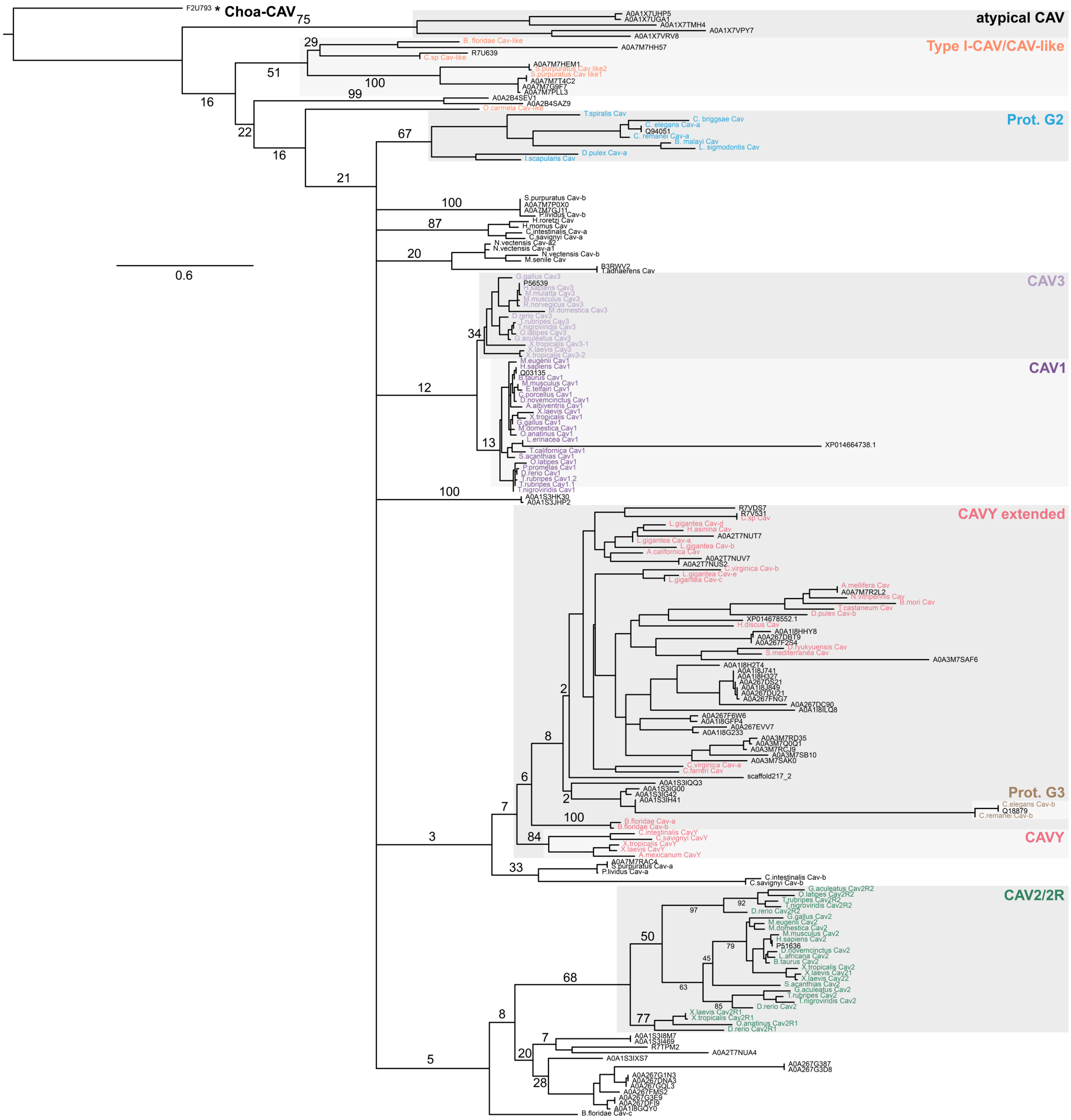

Choa-CAV
atypical CAV
Type I-CAV/CAV-like
Prot. G2
CAV3
CAV1
CAVY extended
Prot. G3
CAVY
CAV2/2R
0.6

**Fig. S1. Phylogenetic relationships of caveolin sequences.** A maximum likelihood phylogeny was inferred for representative caveolins from the current study (black) in combination with caveolin sequences previously analyzed by (*3*). Previously analyzed caveolins are color-coded according to their classifications (*3*). Rooting was performed under the assumption that the choanoflagellate sequence constitutes an outgroup. Support values (percentage replication in 1000 rapid bootstrap pseudo-replicates) are shown for the major splits. Branch lengths are proportional to average number of substitutions per site (refer to scale). Splits denoting the higher order relationship between the apparent Prot. G2, CAV3, CAV1, CAVY (ext.), Prot. G3, and CAV2/2R clades received extremely low bootstrap support and are therefore represented by a polytomy in the final tree. *, caveolin-related protein from *Salpingoeca rosetta*.

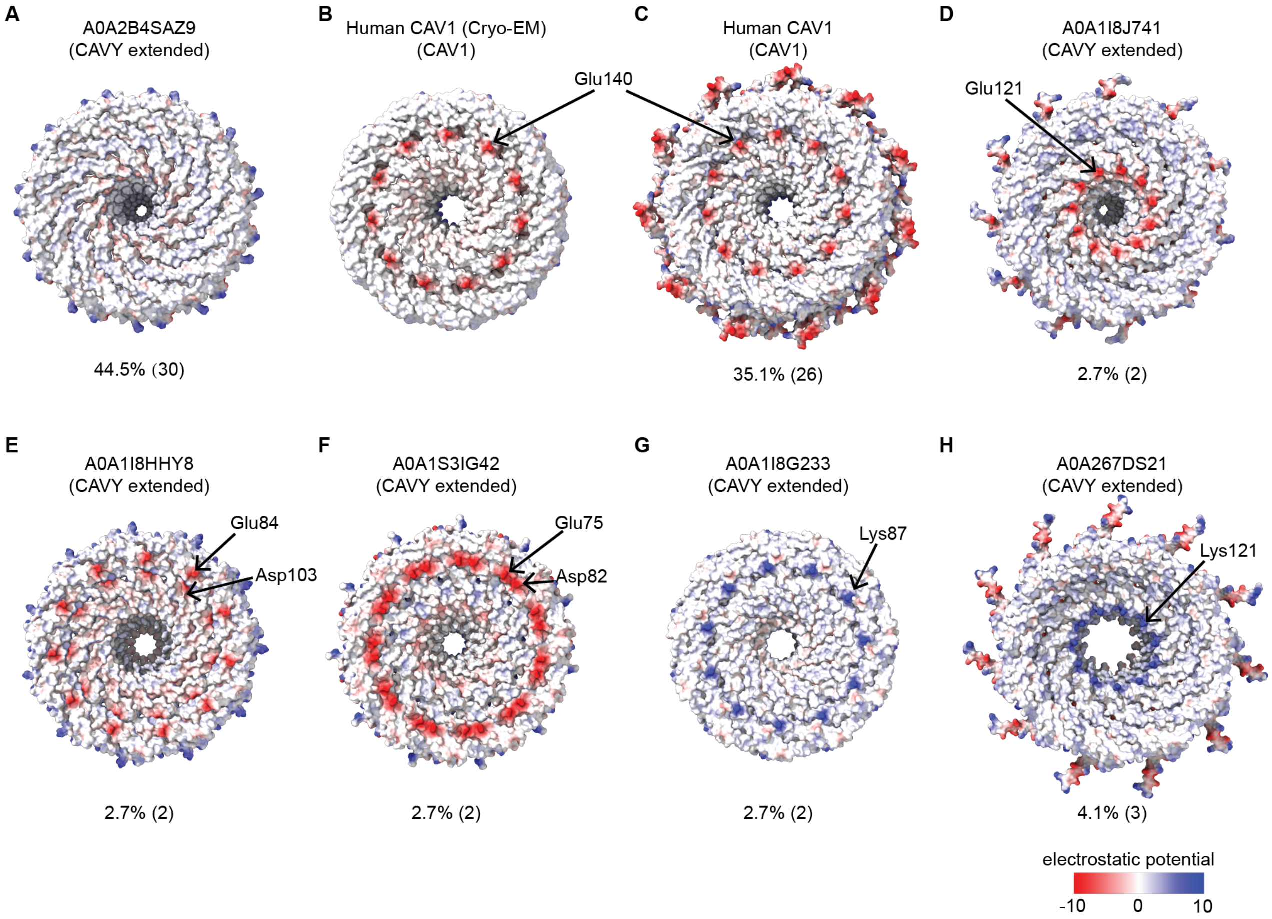

**Fig. S2. Electrostatic potential distribution patterns on the proposed lipid bilayer facing surface of the computationally modeled caveolin oligomers**. Examples of different patterns of charged residues on the proposed membrane facing surface are shown for representative caveolin oligomers predicted by AF2.2. They include **(A)** completely neutral surface; **(B-C)** a negatively charged ring contributed by a single glutamic acid (Glu) located in the middle of the spoke region; **(D)** a negatively charged ring contributed by a single Glu located near the C-terminal region of the spoke region; (**E)** two negatively charged rings contributed by a Glu or an aspartic acid (Asp) in the middle of the spoke region; (**F**) a single negatively charged ring contributed by a Glu and an Asp in the middle of the spoke region, **(G)** a positively charged ring contributed by a lysine (Lys) in the middle of the spoke region, **(H)** a positively charged ring contributed by a Lys near the C-terminal region of the spoke region. The percentage of caveolin complexes exhibiting each pattern is listed below each model (from a total of 74 caveolins investigated).

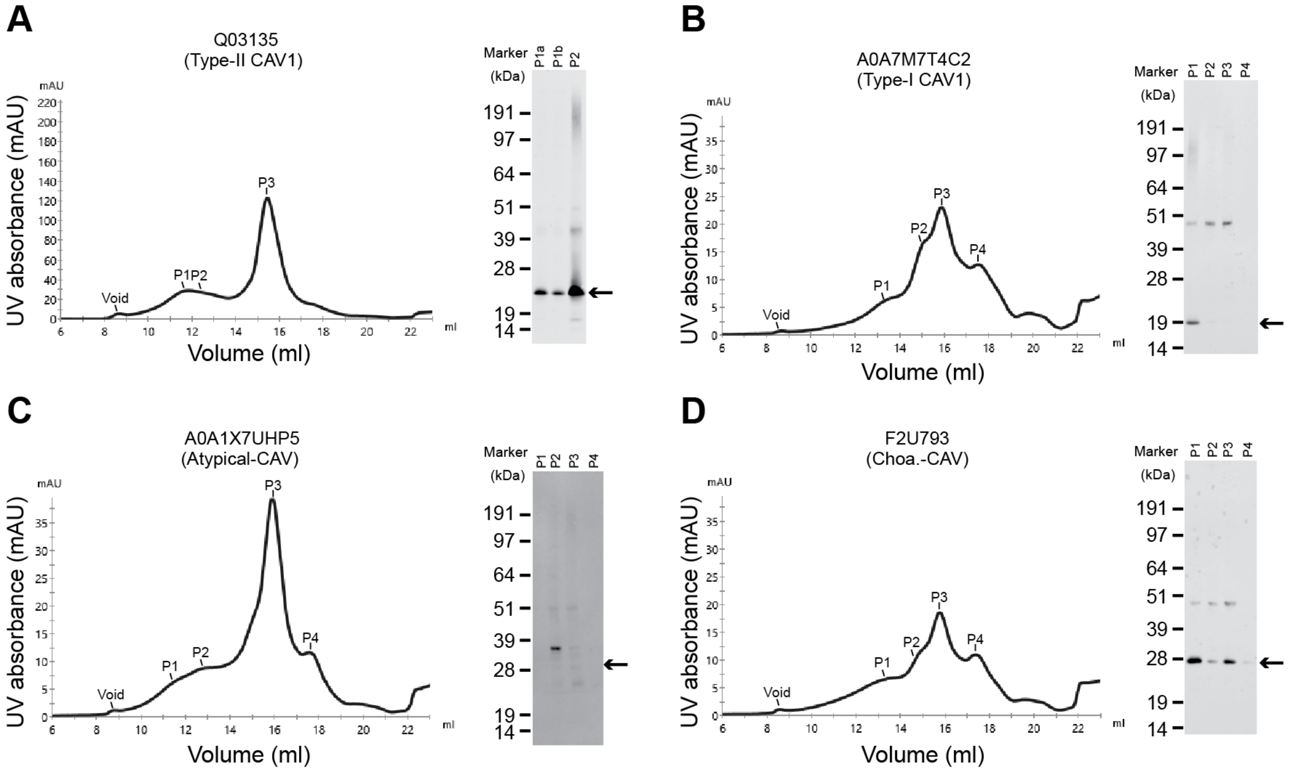

**Fig. S3. Fast protein liquid chromotagraphy (FPLC) traces and Westerns blots of caveolin purifications.** The indicated caveolin proteins were purified from *E. coli* membranes and applied to a Superose®6 10/300 GL column. Elution profiles and Western blotting results are shown for **(A)** Human Cav1 (Type II-CAV, Q03135), **(B)** *S. purpuratus* caveoli (Type I-CAV, A0A7M7T4C2), **(C)** *A. queenslandica* caveolin (Atypical-CAV, A0A1X7UHP5), and **(D)** *S. rosetta* caveolin (Choa-CAV, F2U793). The position of the void and shoulders corresponding to various peaks (P1-P4) are indicated on each FPLC trace. Arrows on the Western blots point to the expected position for monomers for each of the caveolins based on their predicted molecular weight.

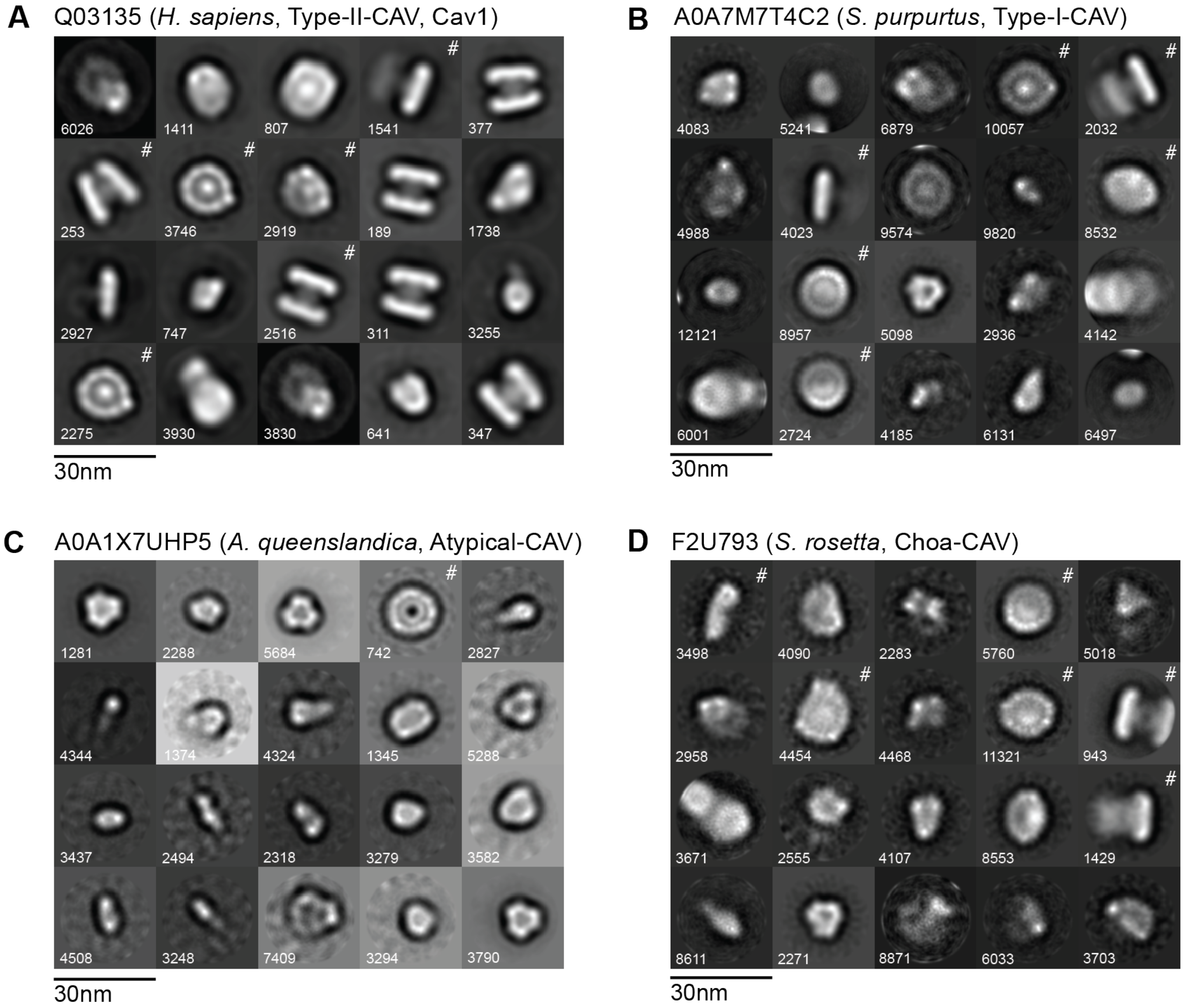

**Fig. S4. Negative stain EM averages of caveolin complexes.** Negative stain 2D class averages of human Cav1 (**A**), *S. purpuratus* caveolin (**B**), *A. queenslandica* caveolin (**C**), and *S. rosetta* caveolin (**D**). Classes denoted with # are shown in **Figure 5**. Scale bar, 30 nm. The majority of 2D classes of *A. queenslandica* caveolin are contaminant proteins. Consequently, only one class is marked as the caveolin complex.

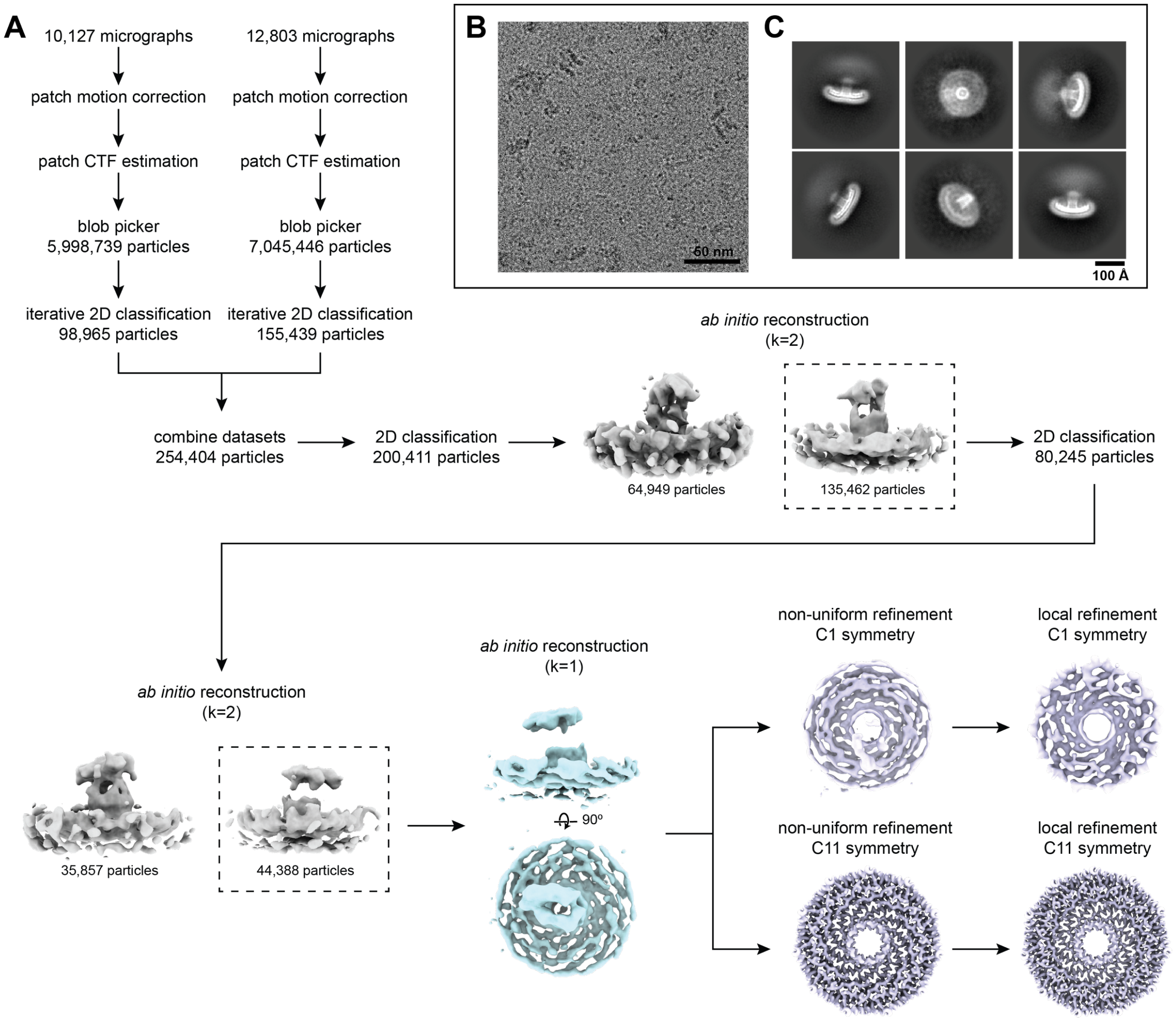

**Fig. S5. Flowchart of cryo-EM processing steps for the *S. purpuratus* caveolin complex.** (**A**) Flowchart representing the classification and analysis of *S. purpuratus* caveolin complex micrographs. Two independently collected datasets were combined following pre-processing, particle picking, and initial 2D classification. *Ab initio* reconstructions that were used for further processing are noted with dashed boxes. *Ab initio* reconstruction used as an input for non-uniform refinement is shown in light blue in an *en face* view and rotated 90˚ around the x-axis. Non-uniform and local refinements with no symmetry applied (C1) and elevenfold symmetry applied (C11) of the *S. purpuratus* caveolin complex are shown in lavender in an *en face* view. (**B**) Representative micrograph of *S. purpuratus* caveolin complex. Scale bar, 50nm. (**C**) Representative *S. purpuratus* caveolin complex 2D classes. Box size, 352 pix$^2$ (390.7 Å x 390.7 Å). Scale bar, 100Å.

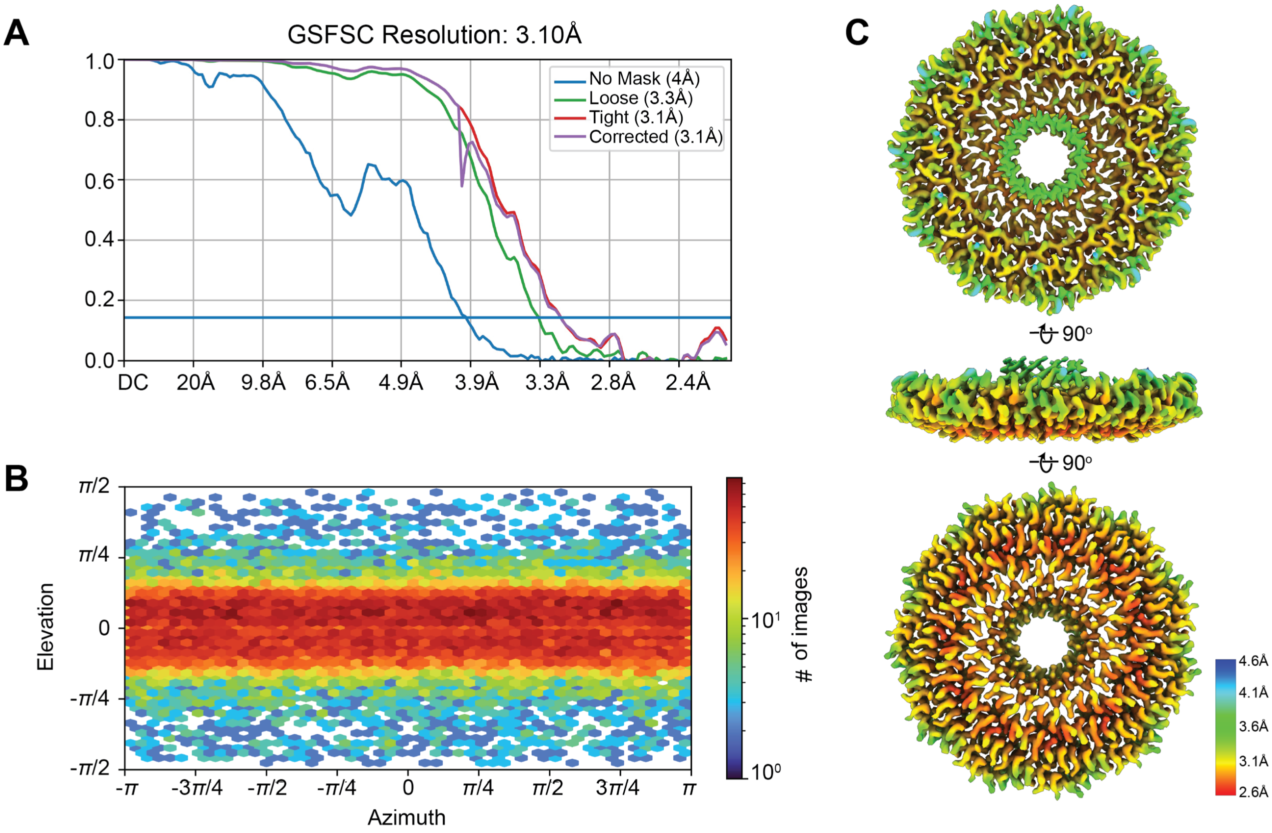

**Fig. S6. Cryo-EM processing of *S. purpuratus* caveolin complex.** (**A**) Gold Standard Fourier Shell Correlation (GS-FSC) of C11 refinement with no mask (blue line), loose mask (green line), tight mask (red line), and corrected (purple). Blue horizontal line, Fourier Shell Correlation (FSC) = 0.143. (**B**) Euler angle plot of angles of particle distribution for the C11 reconstruction of the *S. purpuratus* caveolin complex. (**C**) Heat map of local resolution of C11 3D reconstruction, rotated around the x-axis.

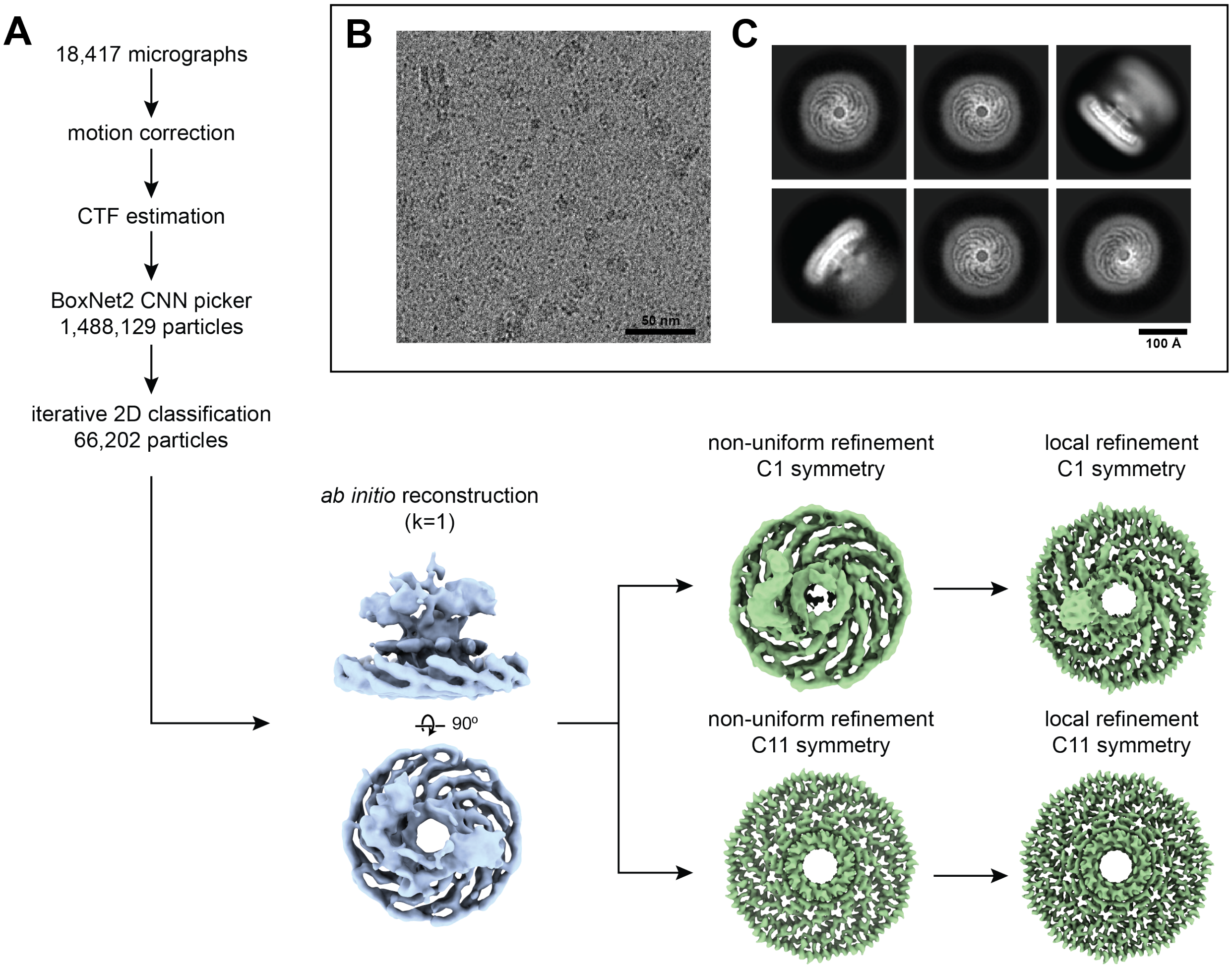

**Fig. S7. Flowchart of cryo-EM processing steps for *S. rosetta* caveolin complex.** (**A**) Flowchart depicting the classification and analysis of *S. rosetta* caveolin complex micrographs. *Ab initio* reconstruction used as an input for non-uniform refinement is shown in light blue in an *en face* view and rotated 90˚ around the x-axis. Non-uniform and local refinements with no symmetry applied (C1) and 11-fold symmetry applied (C11) of the *S. rosetta* caveolin complex are shown in green in an *en face* view. (**B**) Representative micrograph of *S. rosetta* caveolin complex. Scale bar, 50nm. (**C**) Representative *S. rosetta* caveolin complex 2D classes. Box size, 300 pix$^2$ (261Å x 261Å). Scale bar, 100Å.

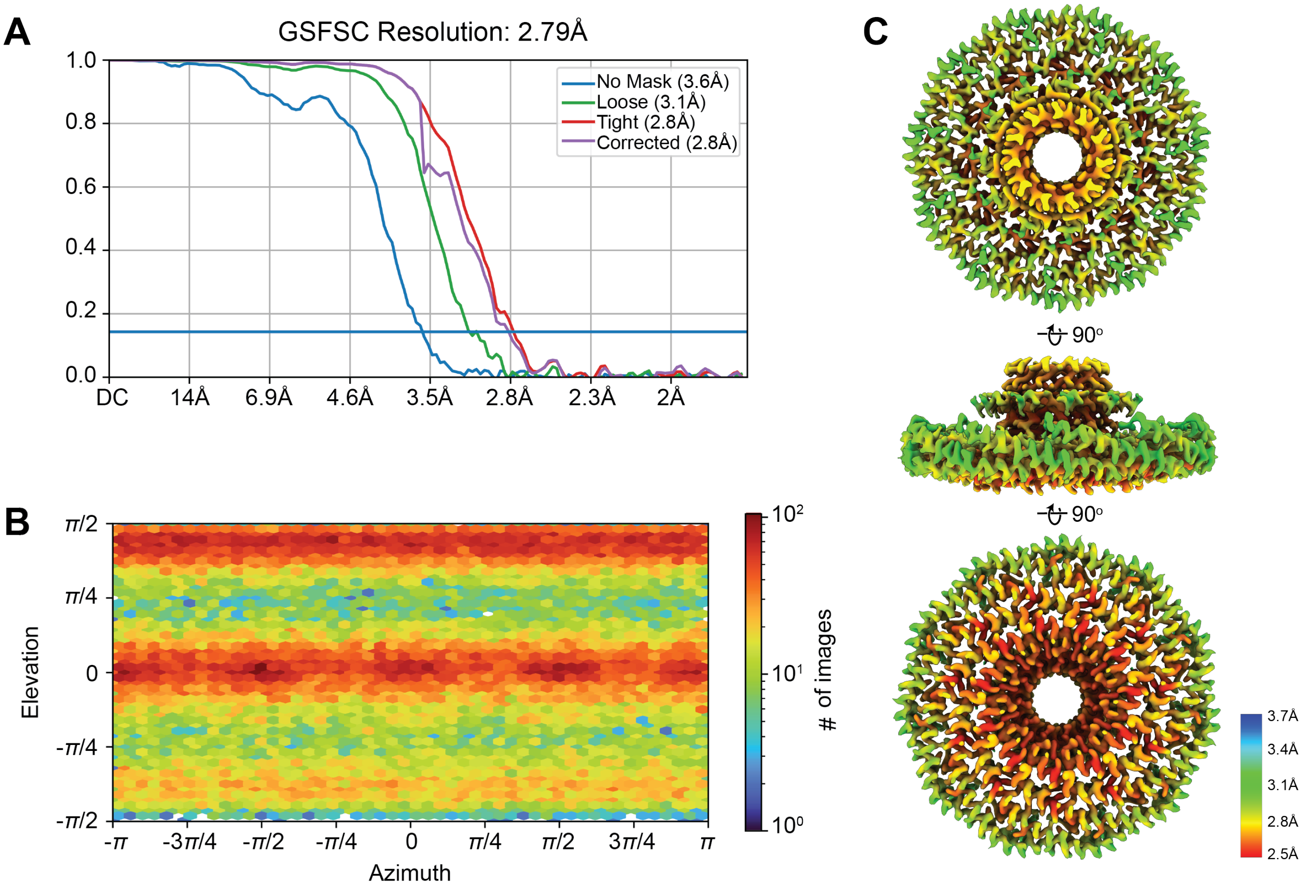

**Fig. S8. Cryo-EM processing of *S. rosetta* caveolin complex.** (**A**) Gold Standard Fourier Shell Correlation (GS-FSC) of C11 refinement with no mask (blue line), loose mask (green line), tight mask (red line), and corrected (purple). Blue horizontal line, Fourier Shell Correlation (FSC) = 0.143. (**B**) Euler angle plot of angles of particle distribution for the C11 reconstruction of the *S. rosetta* caveolin complex. (**C**) Heat map of local resolution of C11 3D reconstruction, rotated around the x-axis.

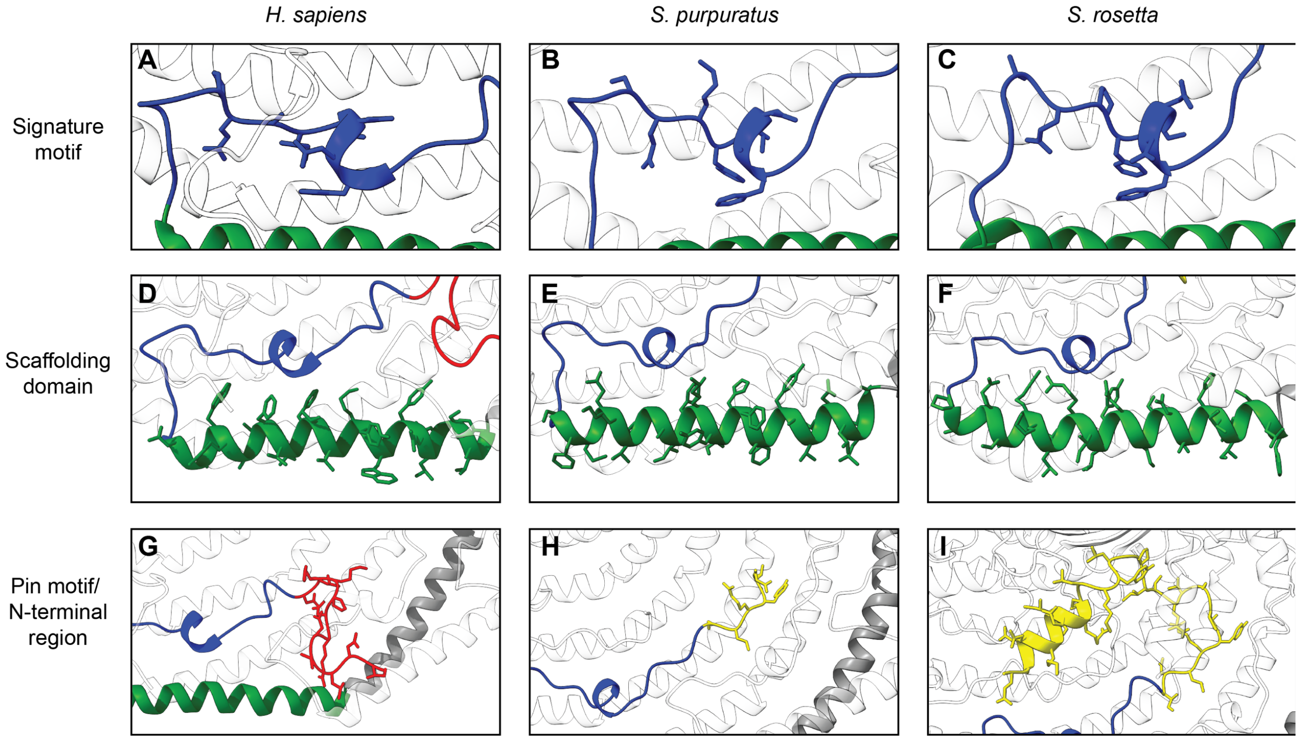

**Fig. S9. Comparison of the signature motif, scaffolding domain, and pin motif/N-terminal region for *H. sapiens*, *S. rosetta,* and *S. purpuratus* caveolins.** Detailed view of the signature motifs (**A-C**) (blue), scaffolding domains (**D-F**) (green), and pin motif (**G**) (red) or N-terminal variable regions (**H & I**) (yellow) of the human Cav1, *S. rosetta* caveolin and *S. purpuratus* caveolin complexes. One protomer is colored according to the structural elements as described in Figure 2 while other protomers of the complex are depicted in transparent gray. The first and last residues of the motifs are labeled, and any residues that are absolutely conserved between the three caveolins are labeled and marked with an asterisk.

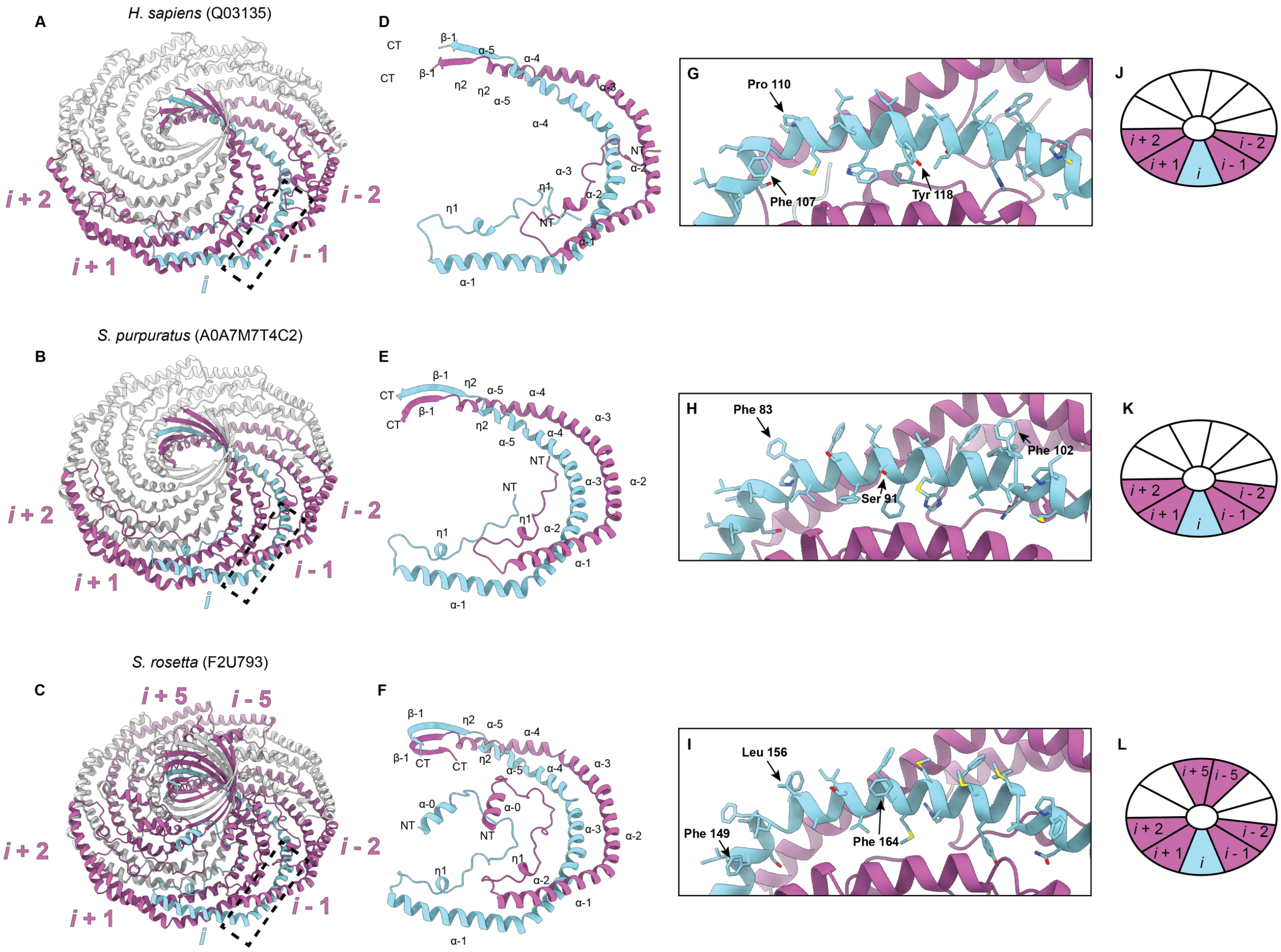

**Fig. S10. Evolutionarily diverse caveolin complexes exhibit extensive contacts between protomers.** (**A-C**) Overall structure of caveolin complexes highlighting one protomer, *i*, in light blue and its interacting protomers in magenta. The interacting protomers are labeled *i* - 2 to *i* + 2 for the caveolin complexes with the *S. rosetta* caveolin complex also exhibiting interactions at protomers *i* - 5 and *i* + 5. (**D-F**) Packing of two protomers with secondary structure elements labeled. (**G-I**) Zoomed-in view of membrane-facing residues indicated by dashed box on overall caveolin structures in (A-C). Complexes are rotated -160˚ around the x-axis from (A-C) to show the membrane-facing surface. Membrane-interacting residues of interest are noted. (**J-L**) Cartoon depiction of caveolin complexes to illustrate the organization of interacting protomers. Color and labeling scheme remain the same as (A-C).

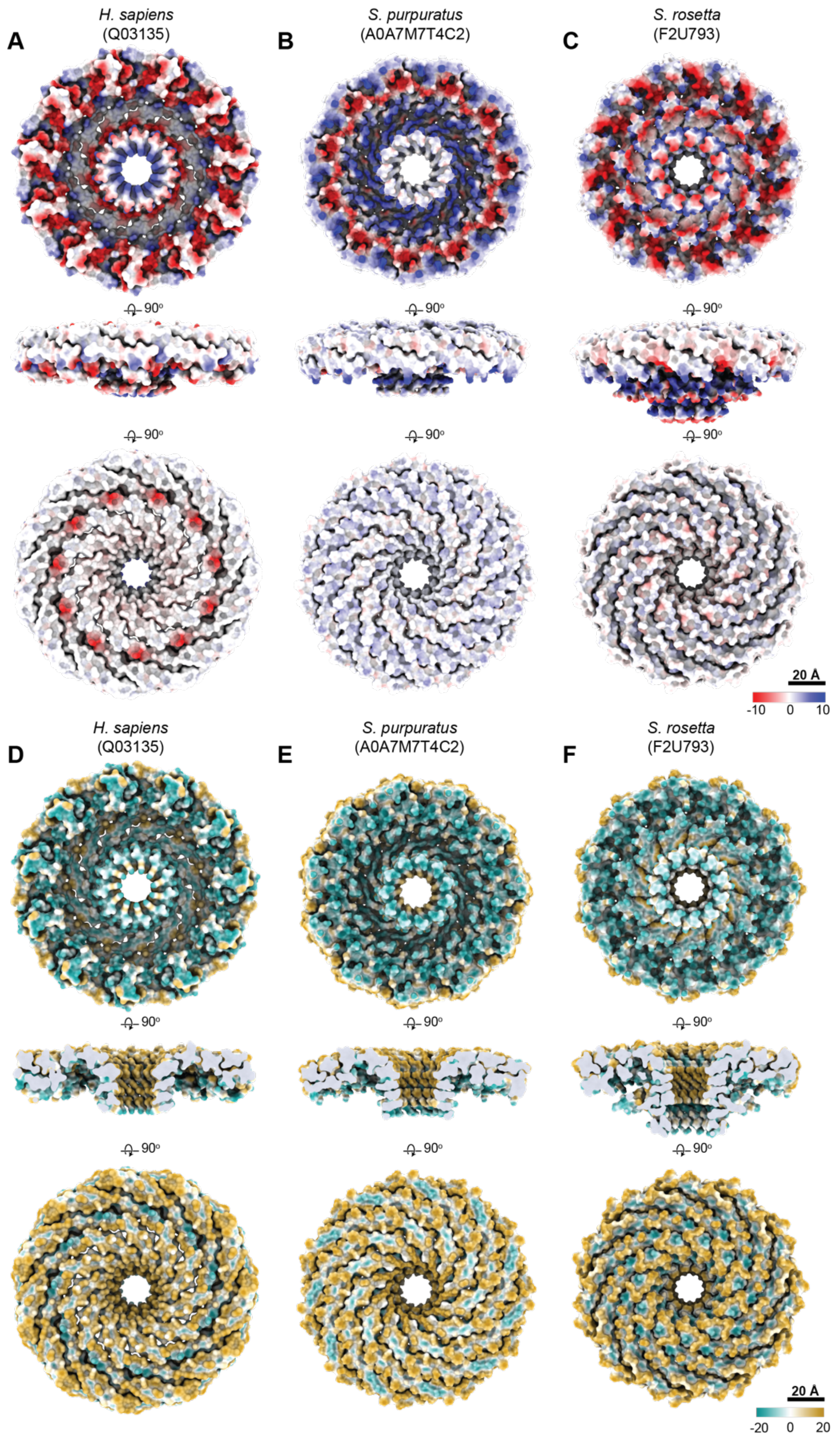

**Fig. S11. Distribution of charged residues and hydrophobicity of the predicted membrane and cytoplasmic facing surfaces of *H. sapiens*, *S. rosetta,* and *S. purpuratus* caveolin complexes. (A-C)** Space filling models of the caveolin complexes rotated 90˚, showing (**A-C**) the charge of the amino acids or (**D-F**) hydrophobicity values. Note that side views in A-C are shown with the surface of the complex whereas a cut through the center of the complex is shown in D-F**.**

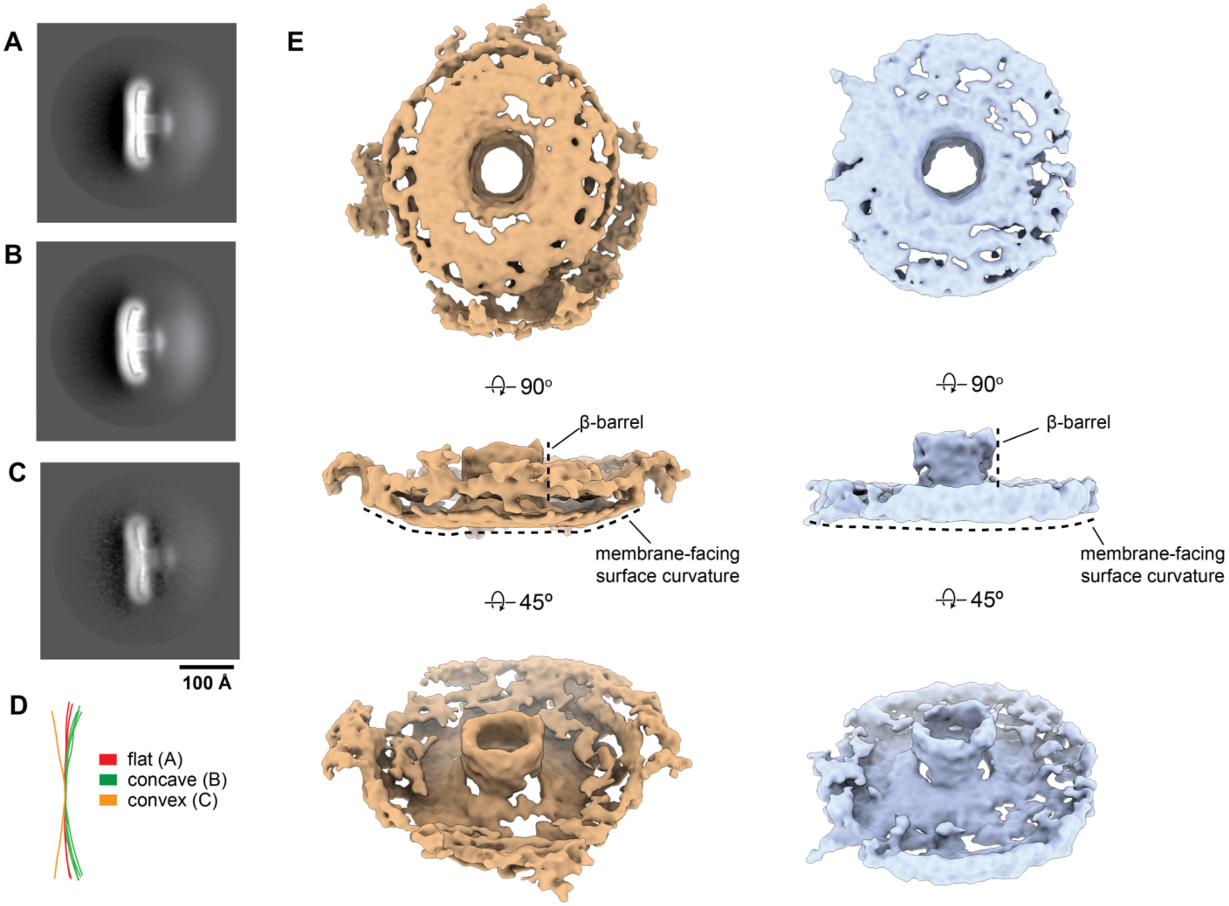

**Fig. S12. *S. purpuratus* caveolin displays various curvatures in 2D classes and 3D variability analysis**. (**A-C**) *S. purpuratus* caveolin 2D class averages that show a flat (**A**), concave (**B**), and convex (**C**) curvature of the complex. Scale bar, 100 Å. (**D**) The difference in curvature is highlighted with overlaid traces of the membrane-facing surface from 2D class averages. (**E**) Structures representing the negative and positive values along the reaction coordinate of a 3D variability analysis (3DVA) component calculated with a resolution limit of 8 Å. The complex on the left (orange) shows a concave membrane surface curvature while the complex on the right (light blue) shows a flat membrane surface curvature and "lifting" of the β-barrel above the rim of the complex. The proposed membrane facing surface is shown (top), rotated 90˚ around the x-axis to show a side view (middle), and rotated an additional 45˚ to show a view of the predicted cytoplasmic facing surface (bottom).

**Table S1. Alignment of the caveolin sequences clustered in Figure S1.** Sequences were truncated to a region corresponding to residues 54–158 in human Cav1 prior to alignment. Residues were colored by percentage identity. Four standard annotations for the alignment (conservation, quality, occupancy and consensus) are shown below the sequences. The number of "+" symbols in each cell represents the frequency of occurrence of a specific structural unit in the caveolins of the corresponding clade. The number in parentheses indicates the average number of amino acids constituting that structural unit within the caveolin clades (rounded to the nearest integer). For Type I-CAV caveolins, where the C-term β-strand is predicted to be discontinuous in two segments, the average is calculated separately for caveolins with a single C-term β-strand and for those with two segments. The results are separated by a comma for the two types of the C-term β-strand, and the averages for the two segments are separated by a forward slash.

**Table S2. Cryo-EM data collection, refinement, and validation statistics**

| | *S. purpuratus* Caveolin complex (EMDB-XXXXX) (PDB-XXXX) | *S. rosetta* Caveolin complex (EMDB-XXXXX) (PDB-XXXX) |
|---|---|---|
| **Data collection and processing** | | |
| Magnification | 81,000x | 105,000x |
| Voltage (kV) | 300 | 300 |
| Electron exposure ($e^-/Å^2$) | 60.0 | 59.2 |
| Defocus range (μm) | -0.5 to -3 | -0.5 to -3 |
| Pixel size (Å) | 1.11 | 0.87 |
| Symmetry imposed | C11 | C11 |
| Initial particle images (no.) | 13,044,185 | 1,488,129 |
| Final particle images (no.) | 135,462 | 66,202 |
| Map resolution (Å) | 3.1 | 2.9 |
| FSC threshold | 0.143 | 0.143 |
| Map resolution range (Å) | 2.6-4.0 | 2.5-3.3 |
| **Refinement** | | |
| Initial model used (PDB code) | Initial model generated by ModelAngelo | Initial model generated by ModelAngelo |
| Model resolution (Å) | 3.3 | 3.0 |
| FSC threshold | 0.5 | 0.5 |
| Map sharpening *B* factor ($Å^2$) | 147.90 | 148.57 |
| Model composition | | |
| Non-hydrogen atoms | 11,088 | 13,409 |
| Protein residues | 1364 | 1,683 |
| Ligands | None | None |
| *B* factors ($Å^2$) | | |
| Protein | -100 | -80 |
| Ligand | None | None |
| R.m.s. deviations | | |
| Bond lengths (Å) | 0.002 | 0.003 |
| Bond angles (º) | 0.580 | 0.659 |
| Validation | | |
| MolProbity score | 0.94 | 0.69 |
| Clash score | 1.58 | 0.56 |
| Poor rotamers (%) | 0.00 | 0.64 |
| Ramachandran plot | | |
| Favored (%) | 97.84 | 98.01 |
| Allowed (%) | 2.16 | 1.99 |
| Disallowed (%) | 0.00 | 0.00 |

**Other supplementary flies**

**File S1. Source data used in the ALG analysis.**

**File S2. Prediction results of the structures of caveolin monomers or oligomers by AlphaFold2.1 with different protomer numbers.** The 2D example color sketches were generated from the 3D model by AlphaFold2_advanced notebook. All right panels of the 2D schematics are colored by pLDDT confidence values. For the left panels, the monomers were colored by N-term to C-term, the multimers were colored by chain. Five models were generated for each prediction. The 2D sketches were based on rank 1 models (R1) in this supplemental file if there is no special note was left under the pLDDT value.

**File S3. Prediction results of the structures of caveolin monomers or oligomers by AlphaFold2.2 with different protomer numbers.** Five models were generated for each prediction. For each prediction, the rank 1 model was displayed in ribbons/slabs mode in the left panel. Models were all colored by pLDDT values. The predicted aligned error (PAE) plots of all five models were displayed in the right panels.

**Movie 1. 3D variability analysis of *S. purpuratus* Cav.** 3DVA of *S. purpuratus* caveolin shows 3D density maps along the variability component that exhibit the complex with a concave and flattened membrane surface conformation. Movement of the *S. purpuratus* caveolin complex from the concave to flattened membrane surface conformations results in the α-helices in the central spoke region "lifting" the beta barrel ~5Å. The complex is shown at a side view and then rotated ~45˚ along the x-axis.
