## Supplementary material for "Evolutionarily diverse caveolins share a common structural framework built around amphipathic discs": File S2

### File S1. Prediction results of the structures of caveolins monomer or oligomers by AlphaFold2.1 with different protomer numbers.

#### Note:

The 2D example color sketches were generated from the 3D model by AlphaFold2_advanced notebook.

The left panel of  the monomers were colored by N → C; the left panel of the oligomers were colored by chain; the right panels were colored by pLDDT values as below color key indicated:

plDDT: Very low (<50) Low (60) OK (70) Confident (80) Very high (>90)

Five models were generated for each prediction. The 2D sketches were based on rank 1 models (R1) in this supplemental file if there is no special note was left under the pLDDT value.

### *Salpingoeca rosetta*

## F2U793

| | *pLDDT* | | |
|---|---|---|---|
| 1-mer | 73.91 | 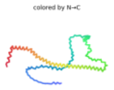  | 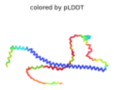  |
| 2-mer | 59.81 | 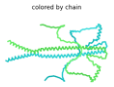  | 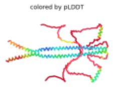  |
| 3-mer | 47.29 | 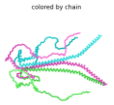  | 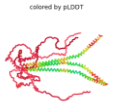  |
| 4-mer | 57.45 | 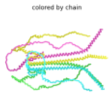  | 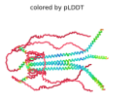  |
| 5-mer | 42.52 | 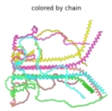  | 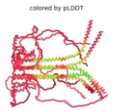  |
| 6-mer | 42.05 | 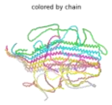  | 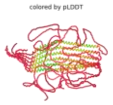  |
| 7-mer | 45.02 | 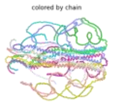  | 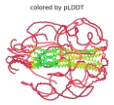  |

### *Amphimedon queenslandica*

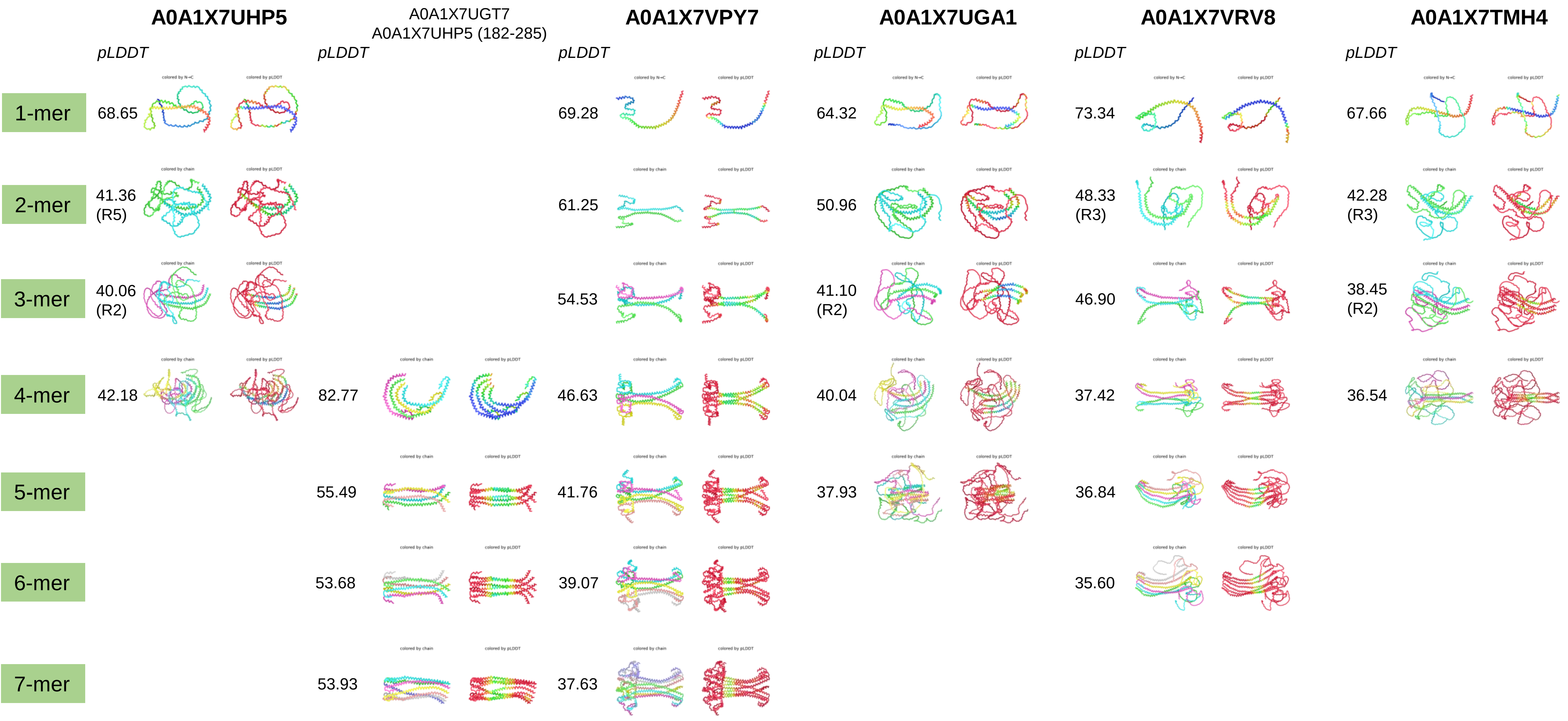

### *Oscarella carmela*

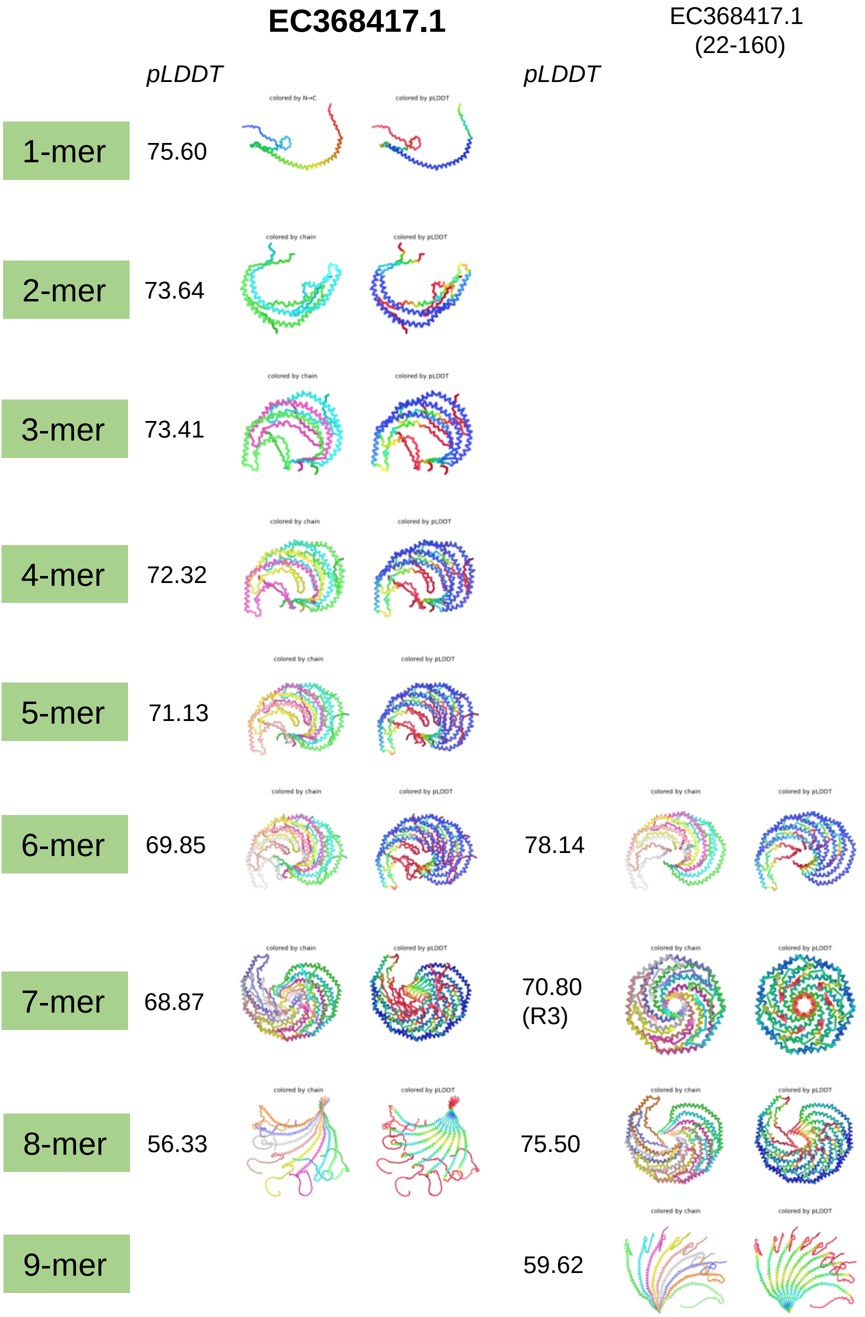

### *Trichoplax adhaerens*

| | **B3RWV2** pLDDT | | B3RWV2 (1-151) pLDDT | |
|---|---|---|---|---|
| 1-mer | 84.57 | 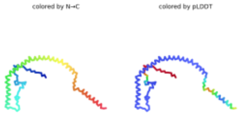 | | |
| 2-mer | 77.88 | 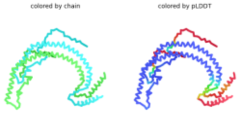 | | |
| 3-mer | 75.07 | 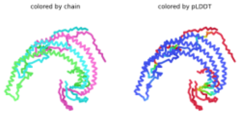 | | |
| 4-mer | 74.30 | 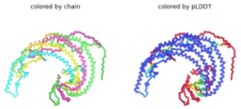 | | |
| 5-mer | 73.37 | 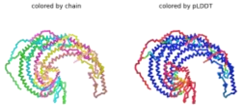 | | |
| 6-mer | 70.10 | 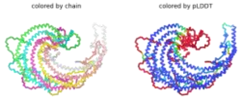 | | |
| 7-mer | 69.58 | 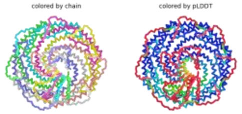 | 75.76 (R2) | 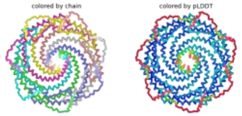 |
| 8-mer | | | 77.55 | 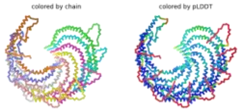 |

### *Stylophora pistillata*

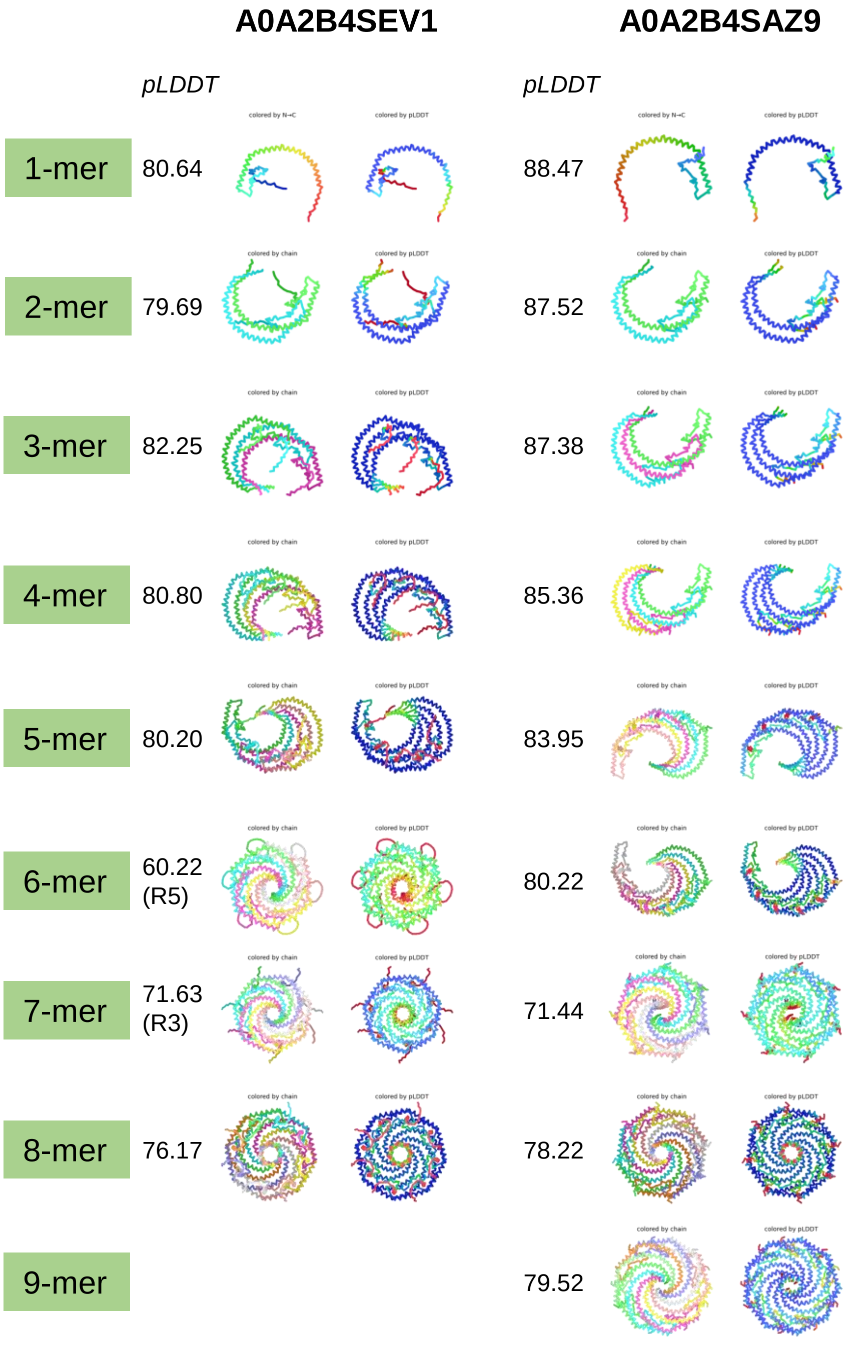

### *Homo sapiens*

| | Q03135 (CAV1) pLDDT | Q03135 32-178 (Beta-CAV1) pLDDT | P51636 (CAV2) pLDDT | P51636 (CAV2) date | P51636 (CAV2) pLDDT | P51636 (CAV2) date | | P56539 (CAV3) pLDDT |
|---|---|---|---|---|---|---|---|---|
| 1-mer | 78.74 | 85.84 | 78.75 | 12/9/2021 | 78.42 | 9/7/2021 | | 89.43 |
| 2-mer | 71.65 | 80.75 | 77.96 | 11/14/2021 | 78.05 | 9/8/2021 | | 86.34 |
| 3-mer | 72.78 | 83.56 | 78.13 | 12/9/2021 | 78.22 | 9/7/2021 | | 86.93 |
| 4-mer | 72.63 | 83.92 | 77.83 | 12/9/2021 | 78.28 | 9/7/2021 | | 86.56 |
| 5-mer | 73.18 | 83.13 | 77.62 | 12/9/2021 | 76.36 | 9/7//2021 | | 86.01 |
| 6-mer | 69.58 | 67.52 (R4) | 73.79 | 12/9/2021 | 74.37 | 9/7/2021 | | 85.48 |
| 7-mer | 68.40 (R2) | 77.40 (R2) | 67.51 | 11/14/2021 | 73.70 | 9/7/2021 | 65.78(R3) | 80.75 (R2) |
| 8-mer | | 80.60 (R2) | 58.30 | 11/14/2021 | 58.30 | 9/7/2021 | | 80.57 |
| 8-mer | | 64.71 | | | | | | |

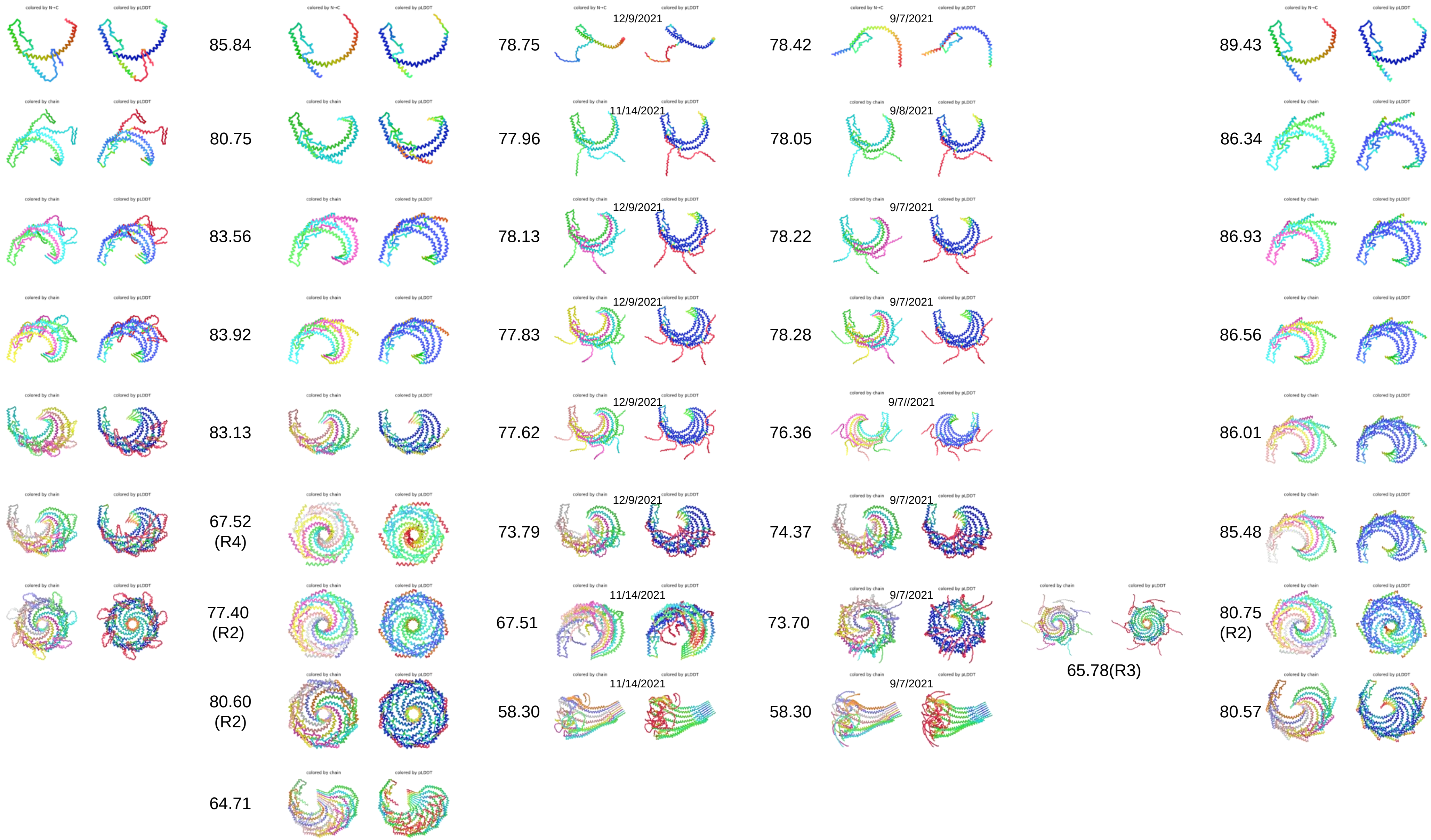

*Strongylocentrotus purpuratus*

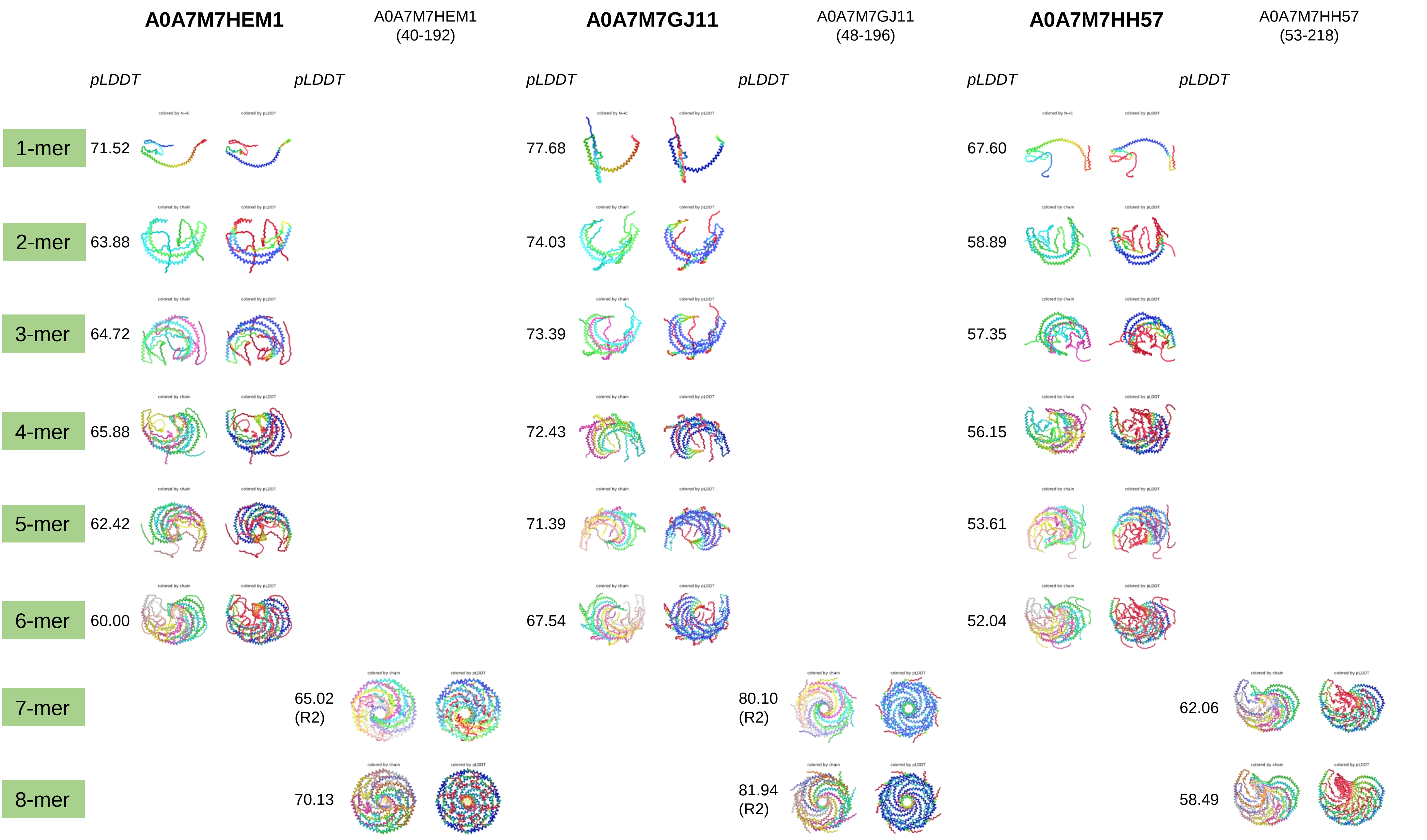

*Strongylocentrotus purpuratus*

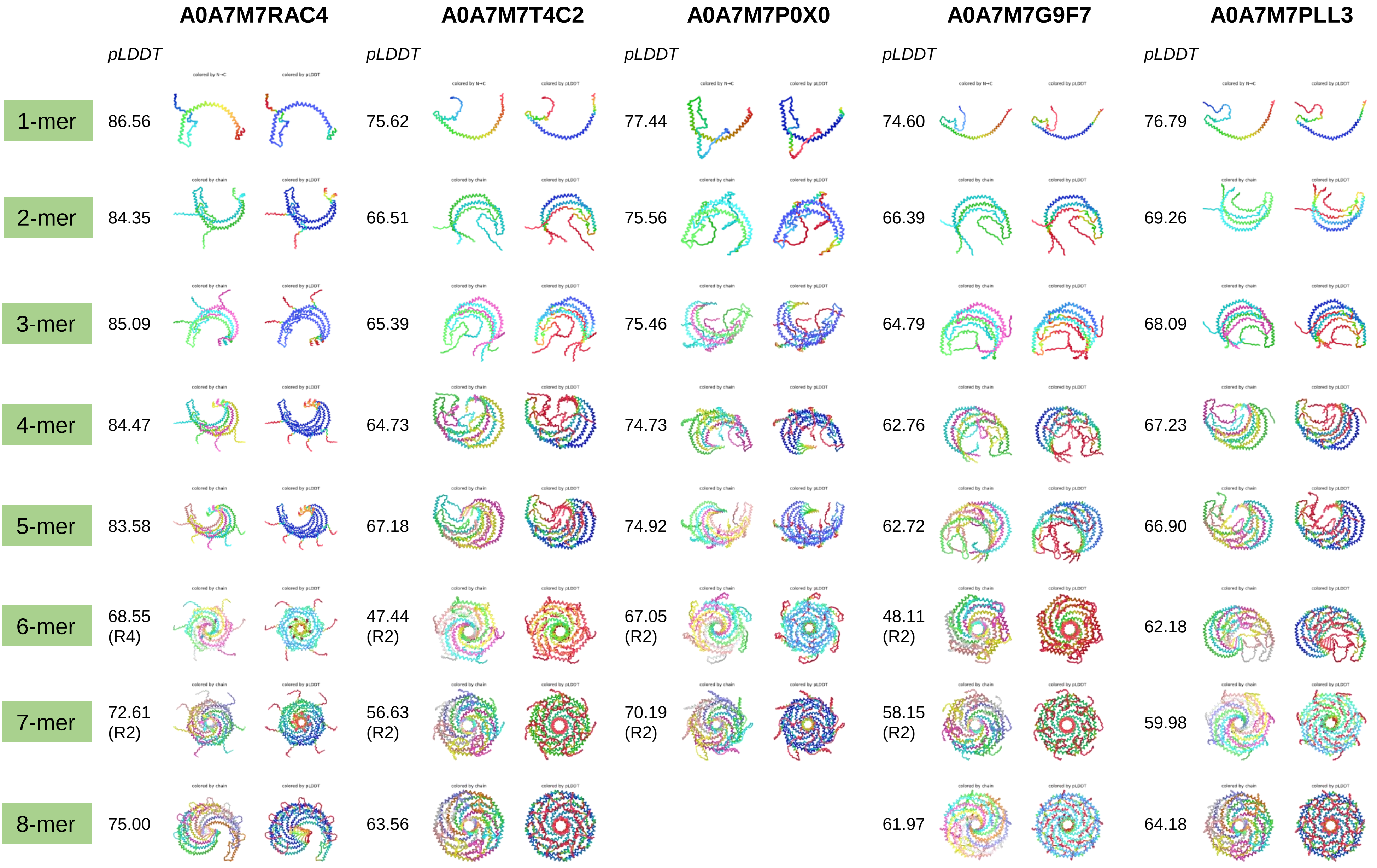

### *Priapulus caudatus*

| | XP_014678552.1 | XP_014664738.1 |
|---|---|---|
| | *pLDDT* | *pLDDT* |
| 1-mer | 92.04 | 63.94 |
| 2-mer | 87.97 | 51.44 |
| 3-mer | 88.48 | 52.25 |
| 4-mer | 87.26 | 48.50 |
| 5-mer | 86.68 | 47.72 |
| 6-mer | 72.19 (R3) | 47.78 |
| 7-mer | 77.05 (R2) | 45.26 |
| 8-mer | 83.06 | 41.55 |
| 9-mer | 48.14 | |

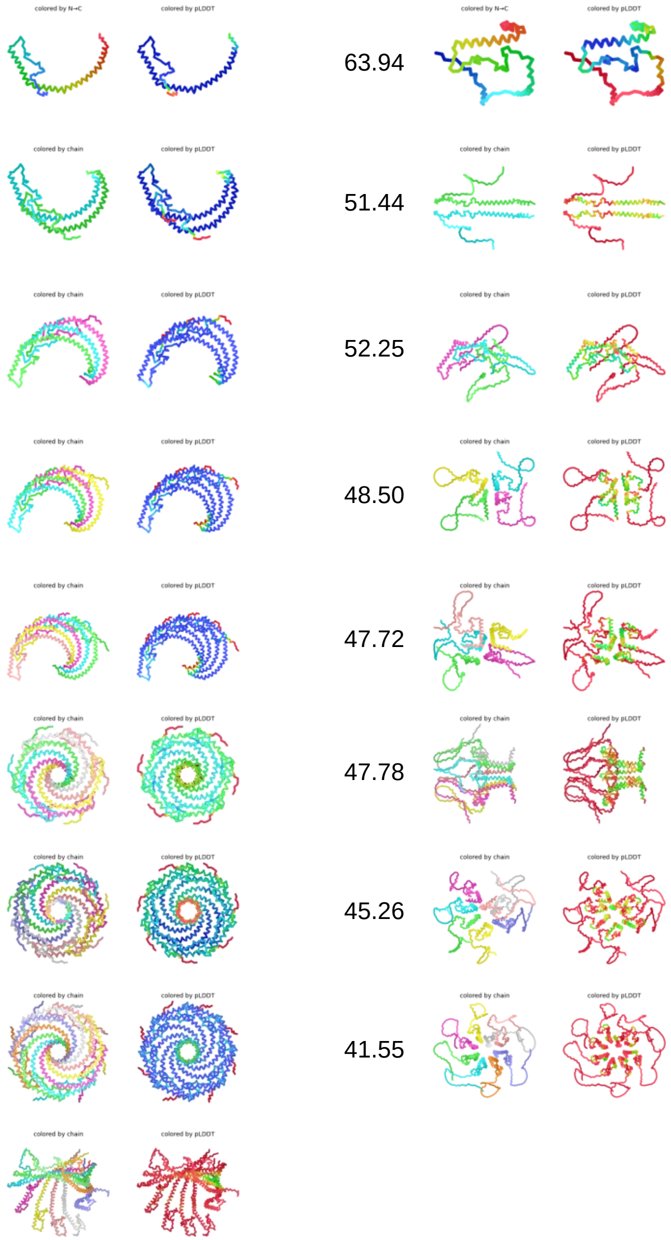

### *Apis mellifera*

## A0A7M7R2L2

*pLDDT*

**1-mer** 80.39

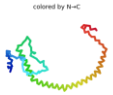

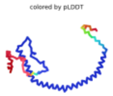


**2-mer** 72.87

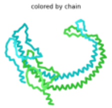

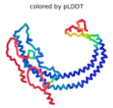


**3-mer** 74.73

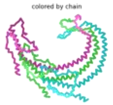

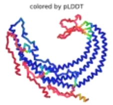


**4-mer** 74.72

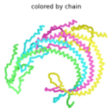

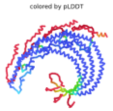


**5-mer** 74.02

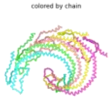

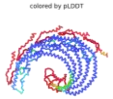


**6-mer** 73.72

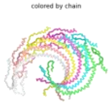

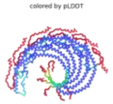


**7-mer** 65.85

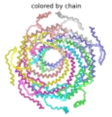

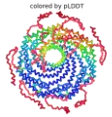


**8-mer**

#### A0A7M7GWE0<br>A0A7M7R2L2 (24-177)

*pLDDT*

**7-mer** 70.61 (R2)

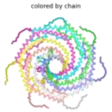

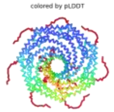


**8-mer** 76.08

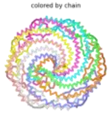

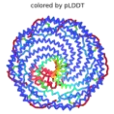

### *Caenorhabditis elegans*

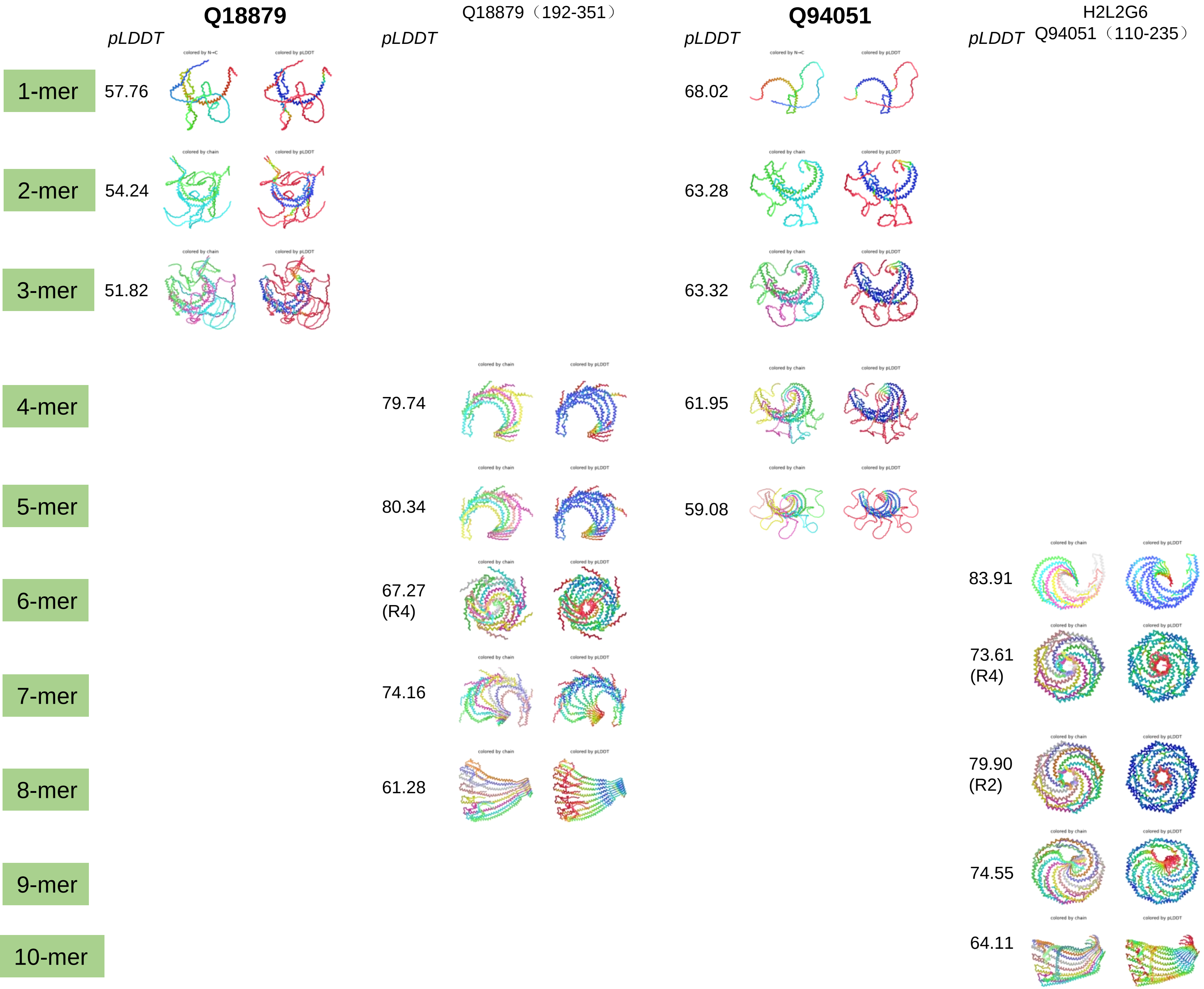

### *Brachionus plicatilis*

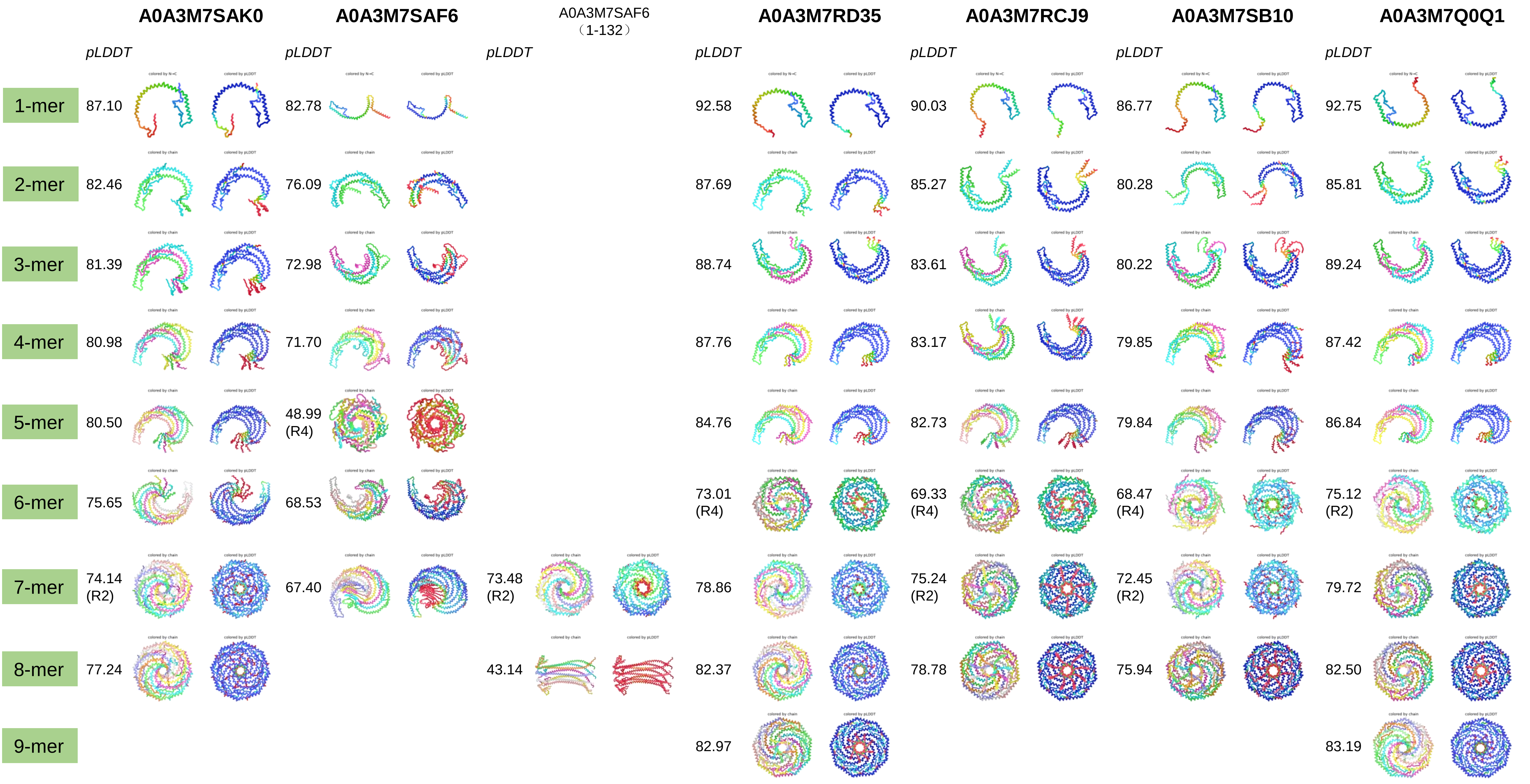

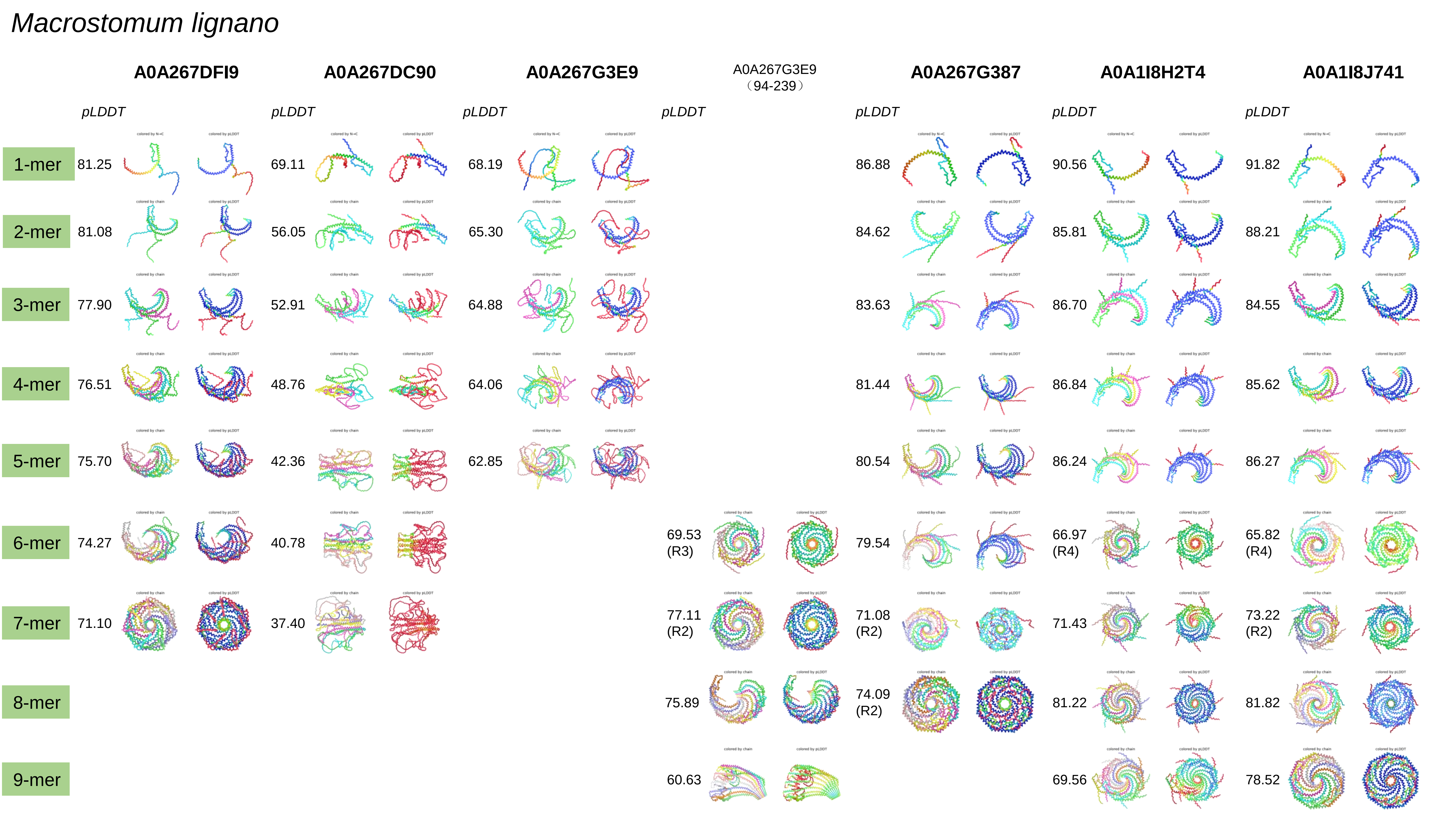
Macrostomum lignano
A0A267DFI9
A0A267DC90
A0A267G3E9
A0A267G3E9 (94-239)
A0A267G387
A0A1I8H2T4
A0A1I8J741
pLDDT
1-mer
81.25
69.11
68.19
86.88
90.56
91.82
2-mer
81.08
56.05
65.30
84.62
85.81
88.21
3-mer
77.90
52.91
64.88
83.63
86.70
84.55
4-mer
76.51
48.76
64.06
81.44
86.84
85.62
5-mer
75.70
42.36
62.85
80.54
86.24
86.27
6-mer
74.27
40.78
69.53 (R3)
79.54
66.97 (R4)
65.82 (R4)
7-mer
71.10
37.40
77.11 (R2)
71.08 (R2)
71.43
73.22 (R2)
8-mer
75.89
74.09 (R2)
81.22
81.82
9-mer
60.63
69.56
78.52

### *Macrostomum lignano*

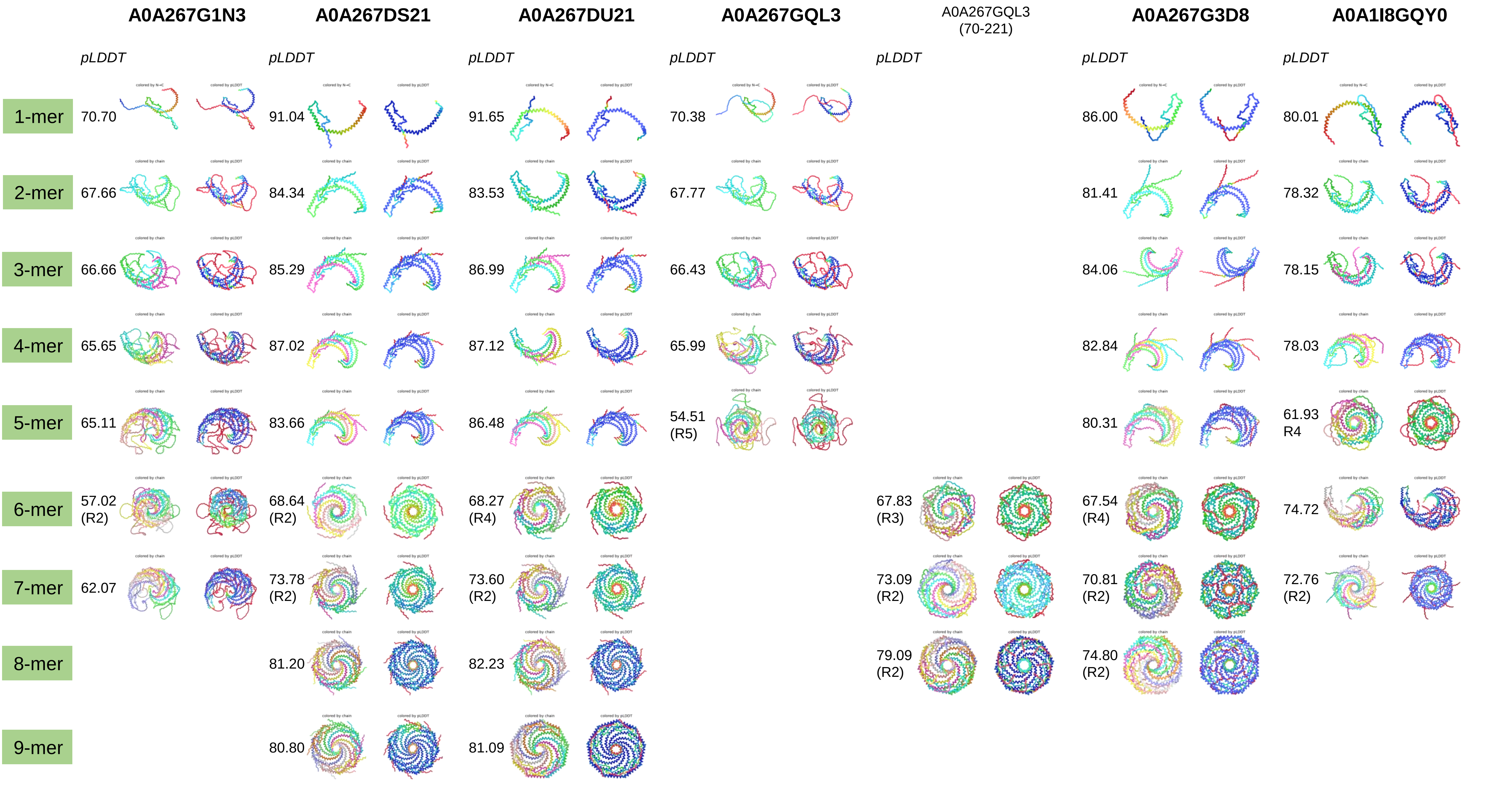

*Macrostomum lignano*

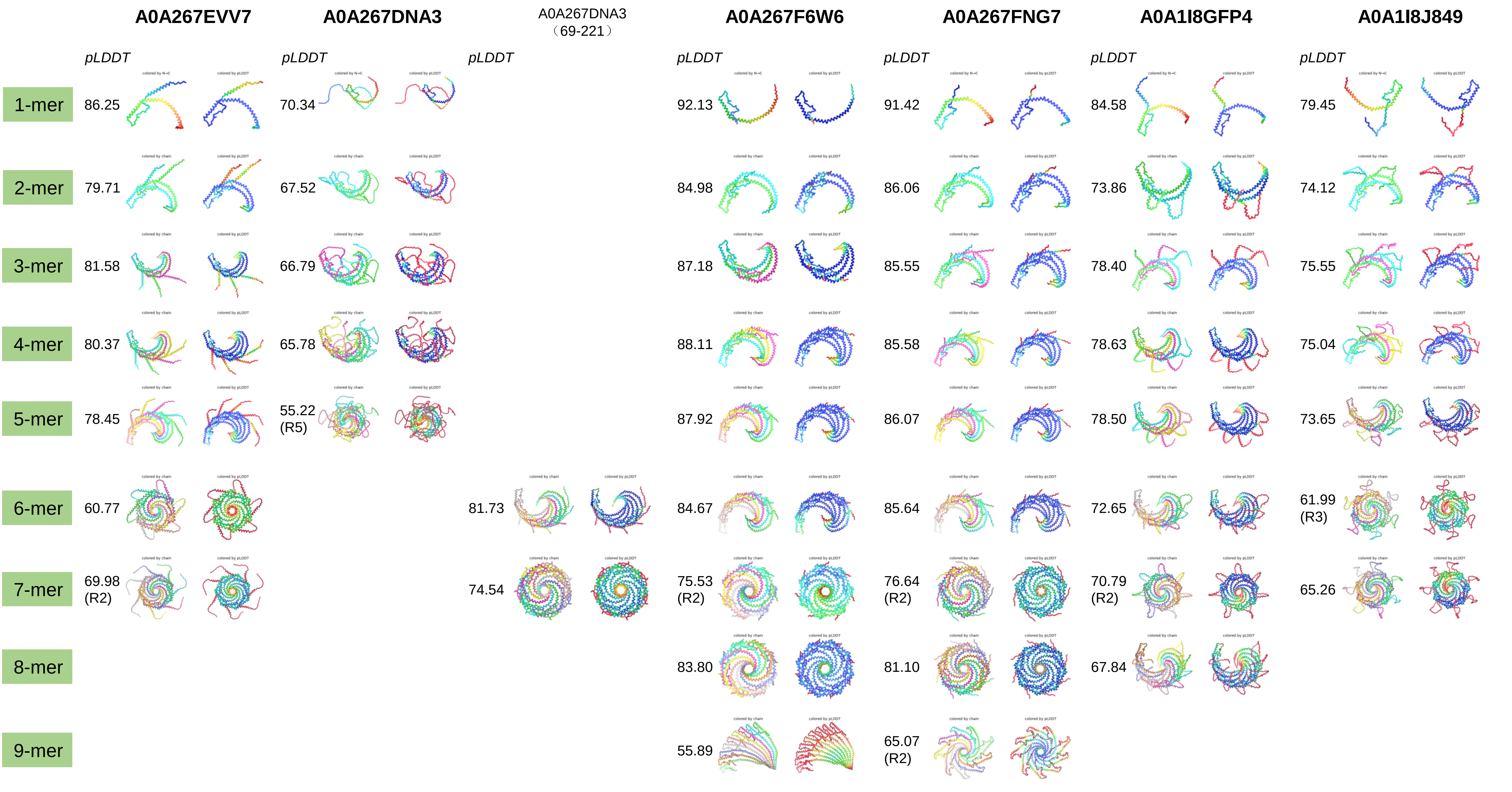

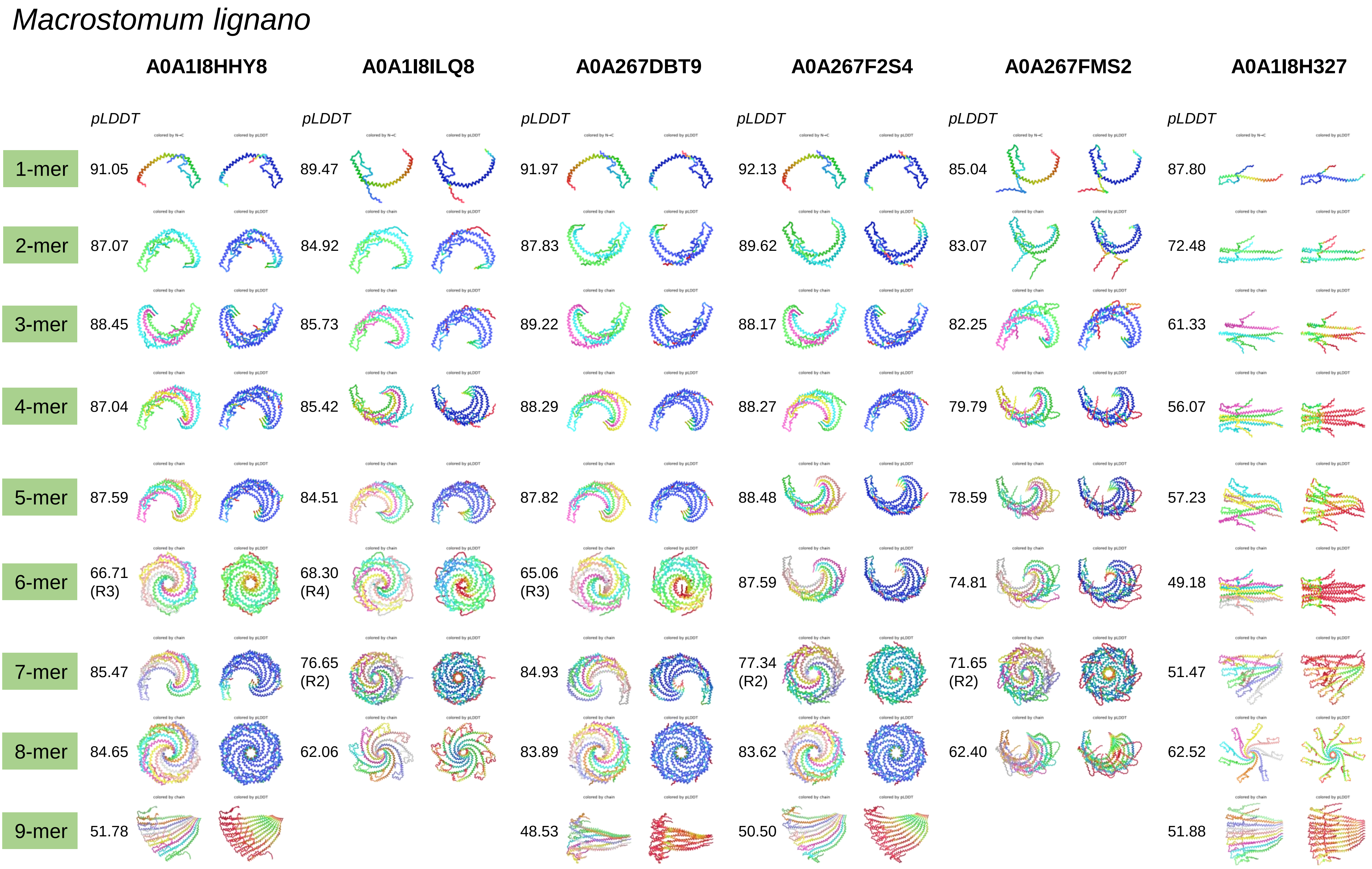
Macrostomum lignano
A0A1I8HHY8
A0A1I8ILQ8
A0A267DBT9
A0A267F2S4
A0A267FMS2
A0A1I8H327
pLDDT
1-mer
91.05
89.47
91.97
92.13
85.04
87.80
2-mer
87.07
84.92
87.83
89.62
83.07
72.48
3-mer
88.45
85.73
89.22
88.17
82.25
61.33
4-mer
87.04
85.42
88.29
88.27
79.79
56.07
5-mer
87.59
84.51
87.82
88.48
78.59
57.23
6-mer
66.71 (R3)
68.30 (R4)
65.06 (R3)
87.59
74.81
49.18
7-mer
85.47
76.65 (R2)
84.93
77.34 (R2)
71.65 (R2)
51.47
8-mer
84.65
62.06
83.89
83.62
62.40
62.52
9-mer
51.78
48.53
50.50
51.88

*Macrostomum lignano*

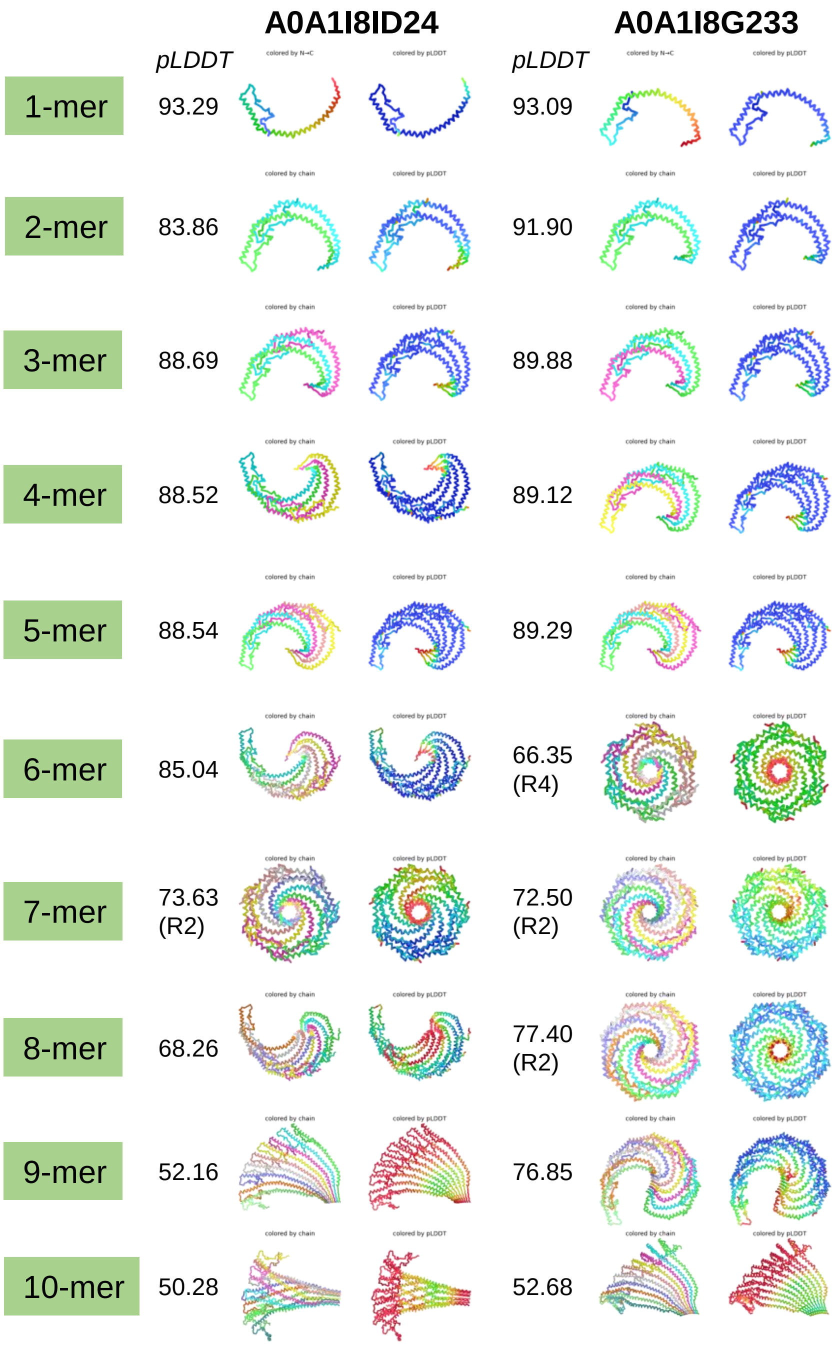

### *Pomacea canaliculata*

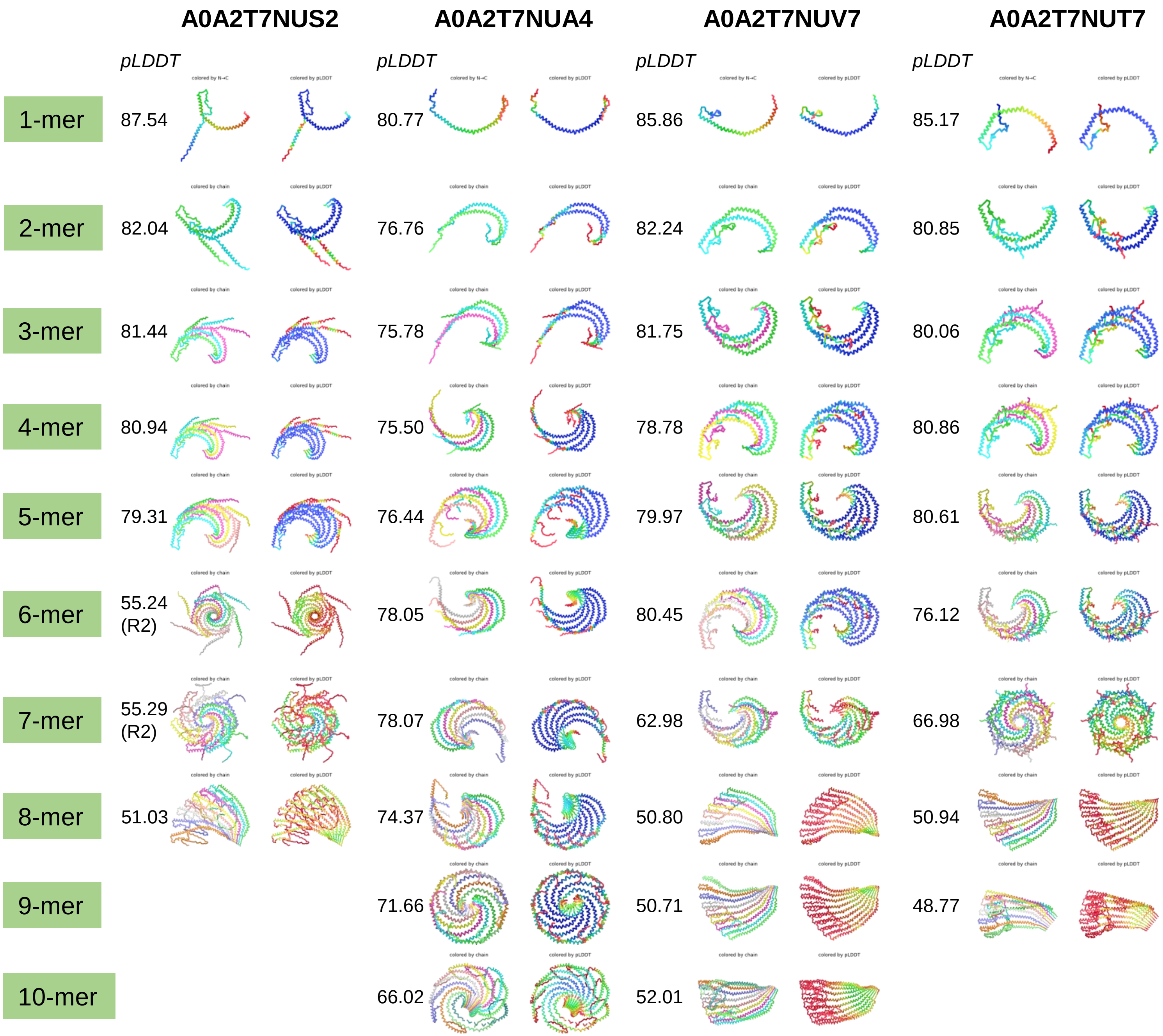

### *Lingula unguis*

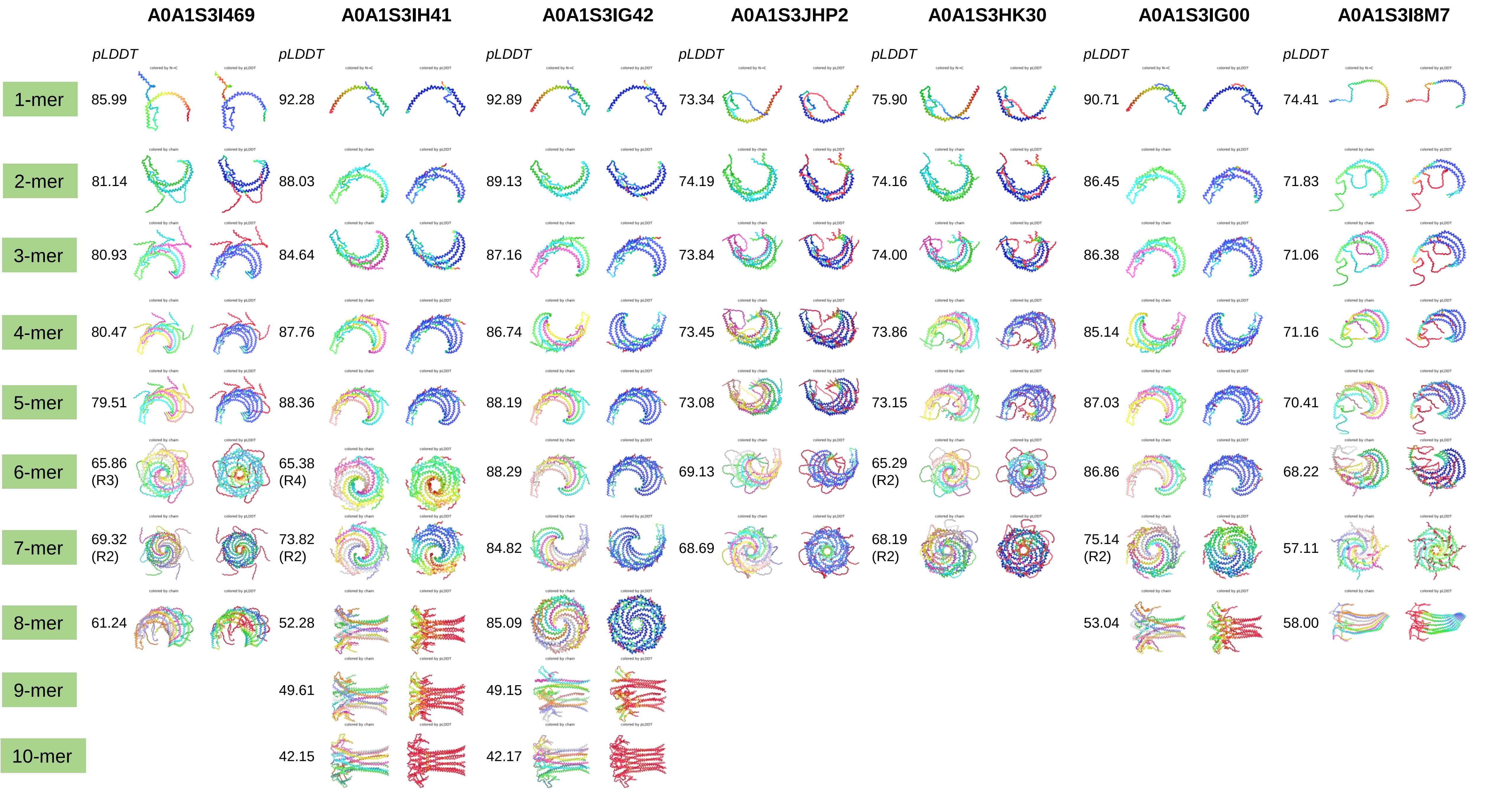

### *Lingula unguis*

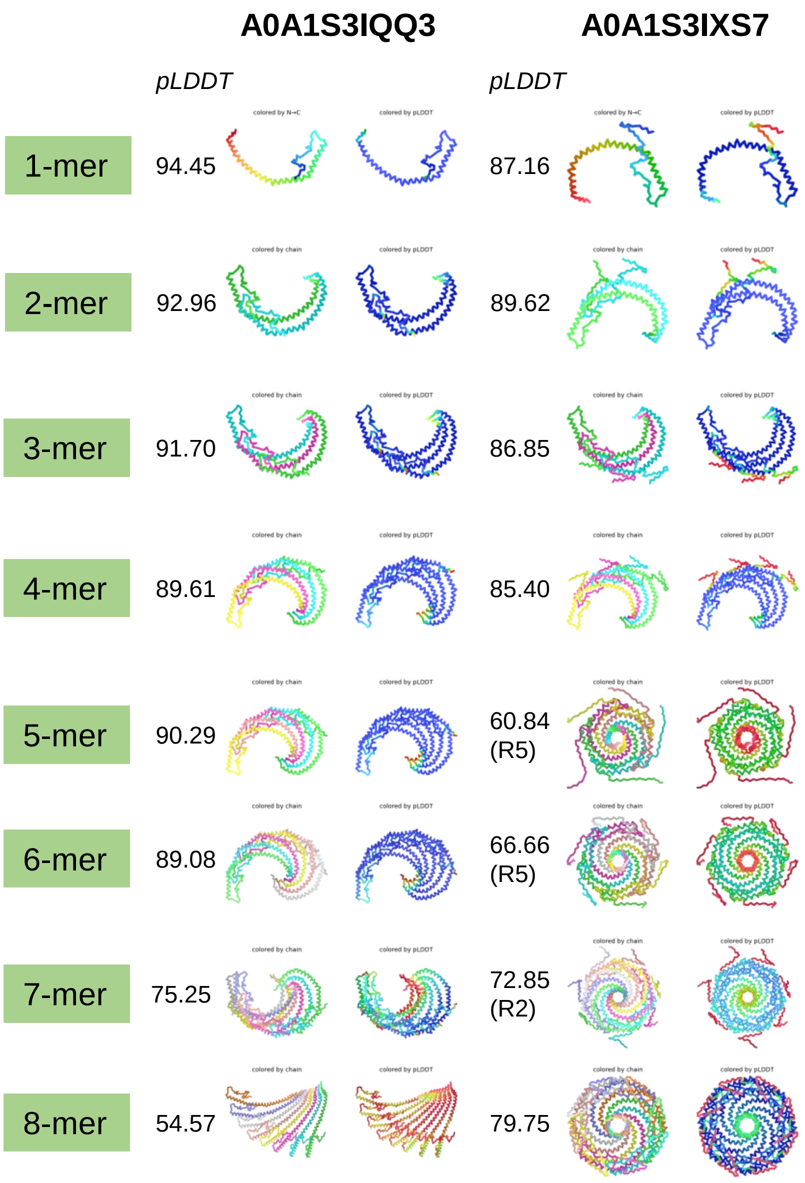

### *Capitella teleta*

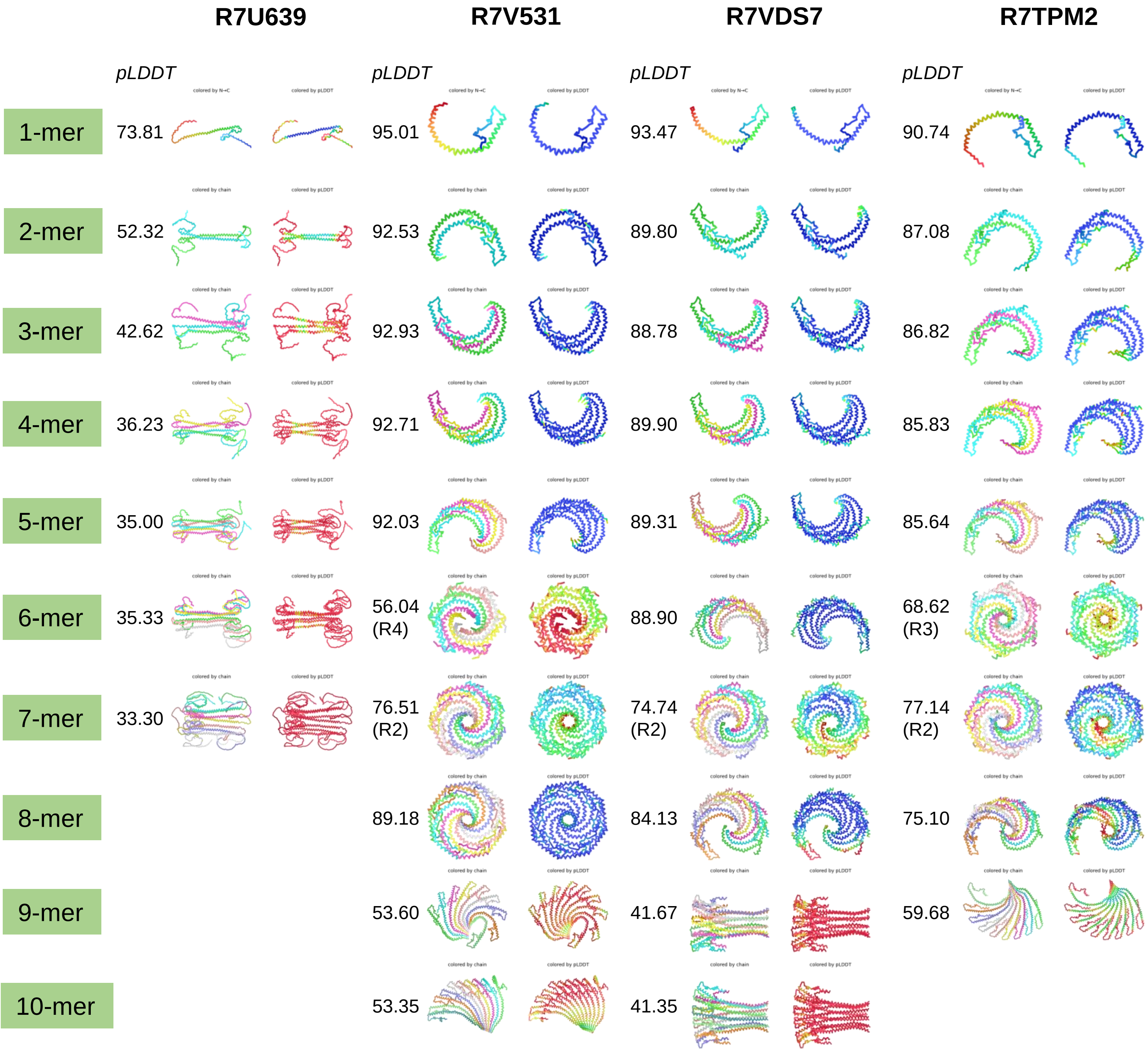
