## Supplementary material for "Evolutionarily diverse caveolins share a common structural framework built around amphipathic discs": File S3

### File S2. Prediction results of the structures of caveolins monomer or oligomers by AlphaFold2.2 with different protomer numbers.

#### Note:

Five models were generated for each prediction. For each prediction, the rank 1 model was displayed in ribbons/slabs mode in the left panel.

Models were all colored by pLDDT values as below color key indicated:

pLDD T: 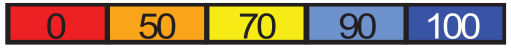


The PAE plots of all five models were displayed in the right panels.

### F2U793 *S. Rosetta*

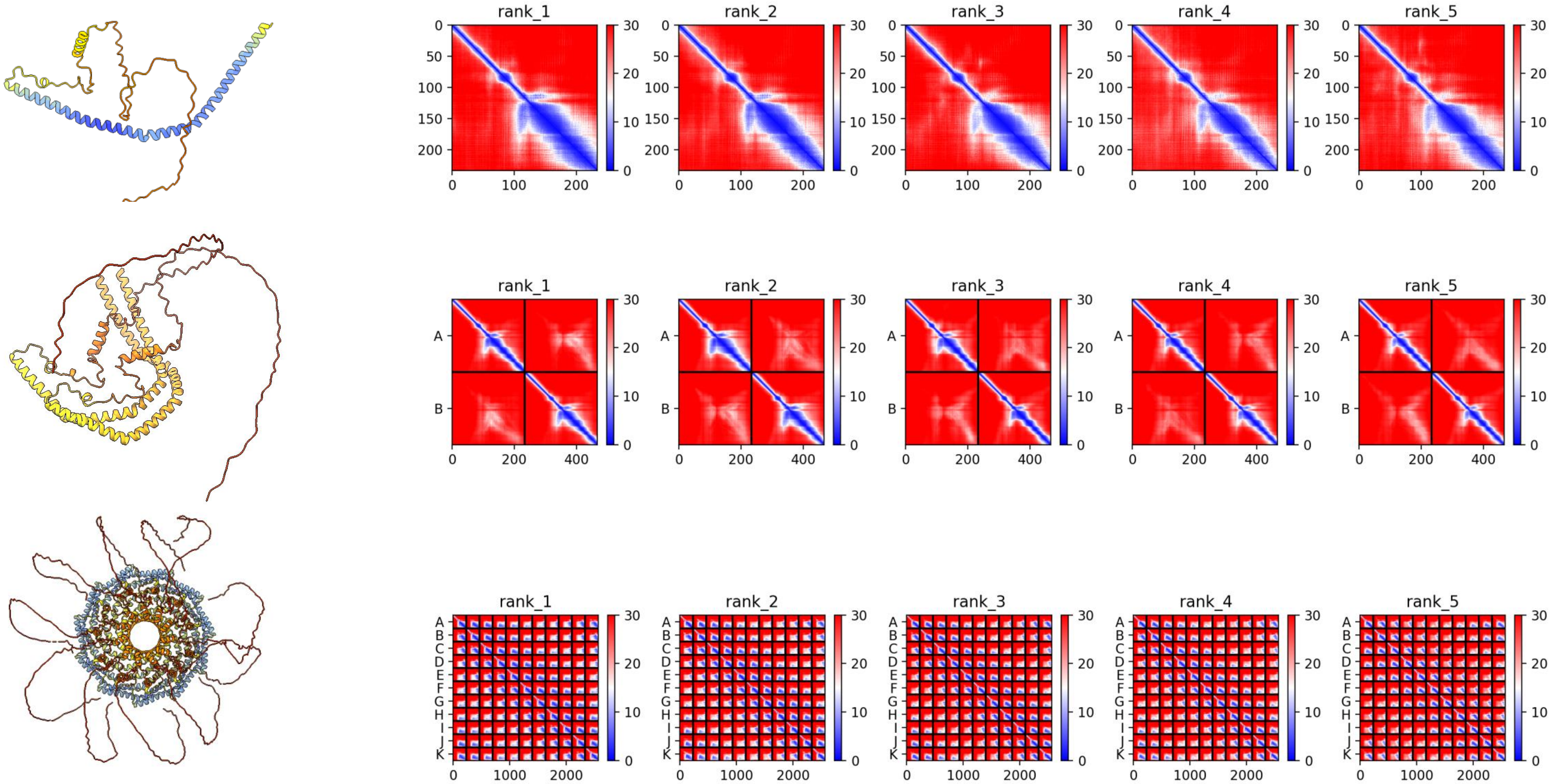

### A0A1X7UHP5 *A. queenslandica*

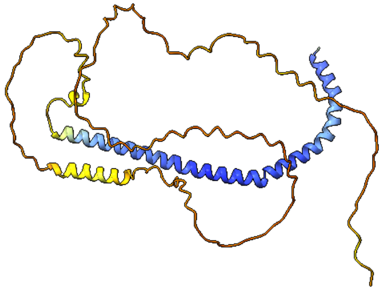

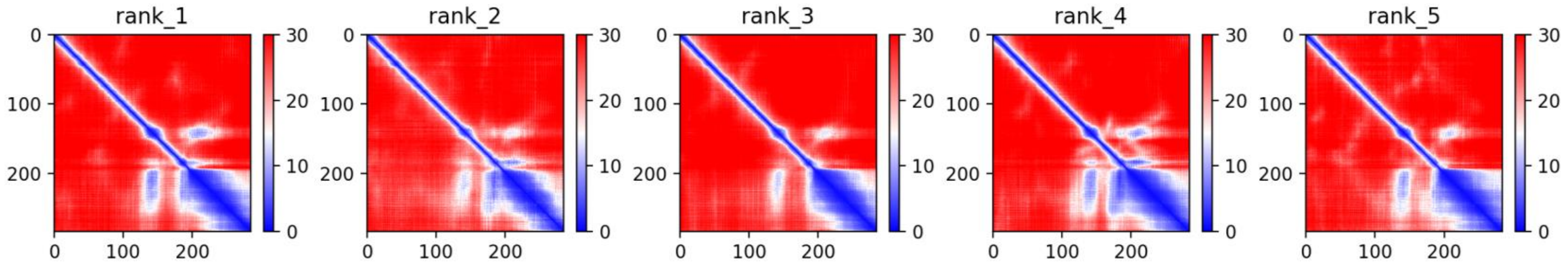


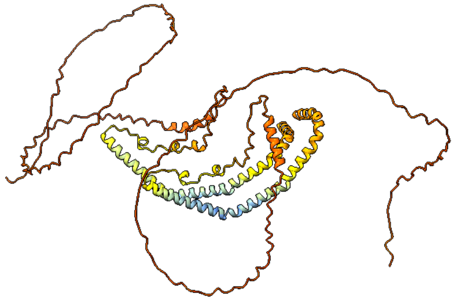

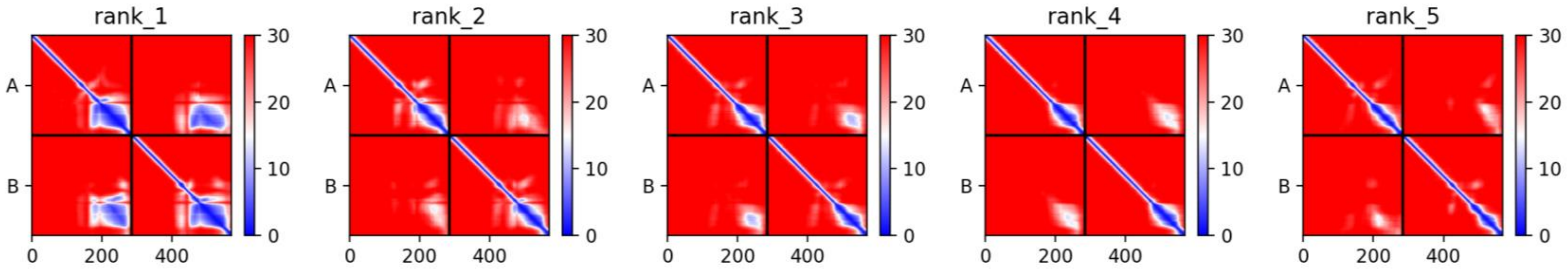


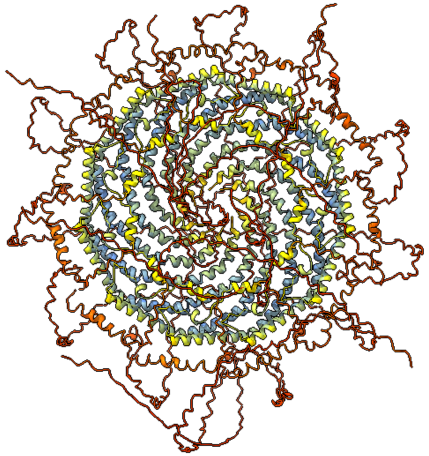

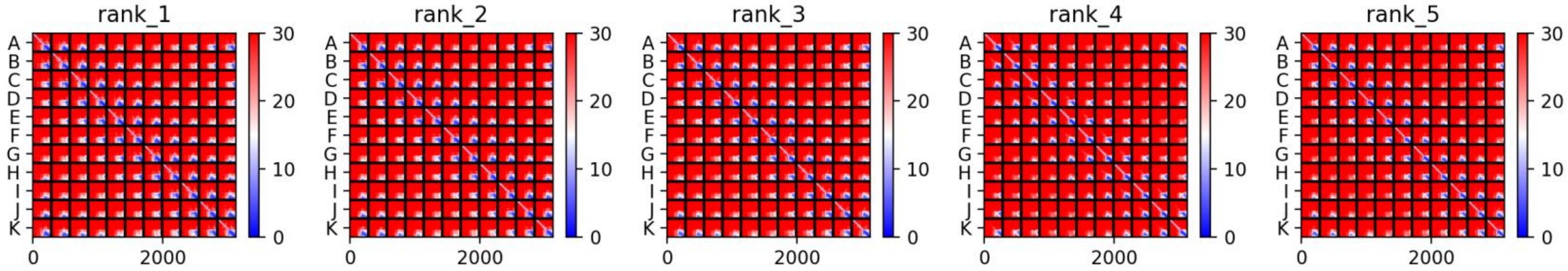

### A0A1X7UGA1 *A. queenslandica*

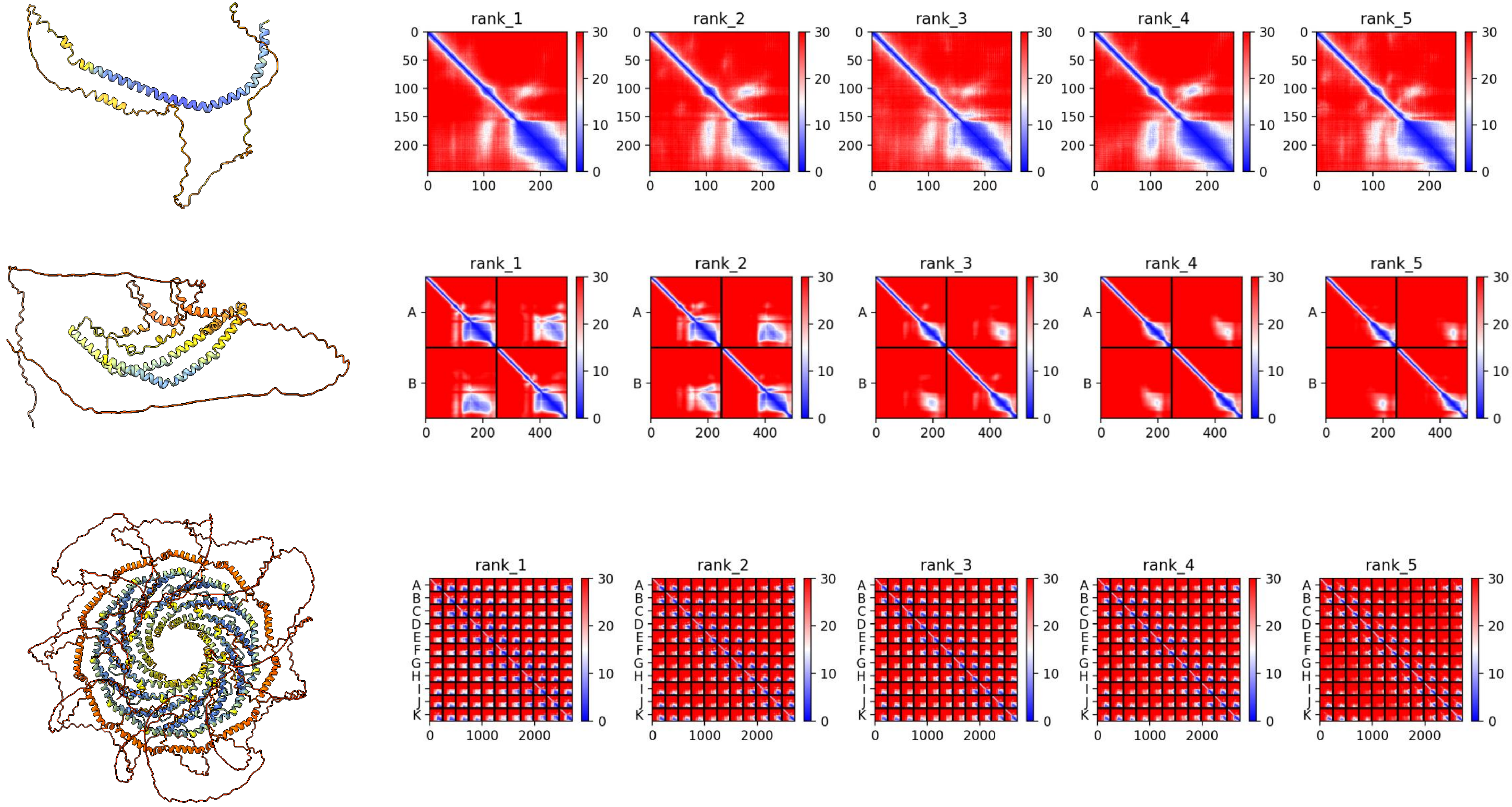

### A0A1X7TMH4 *A. queenslandica*

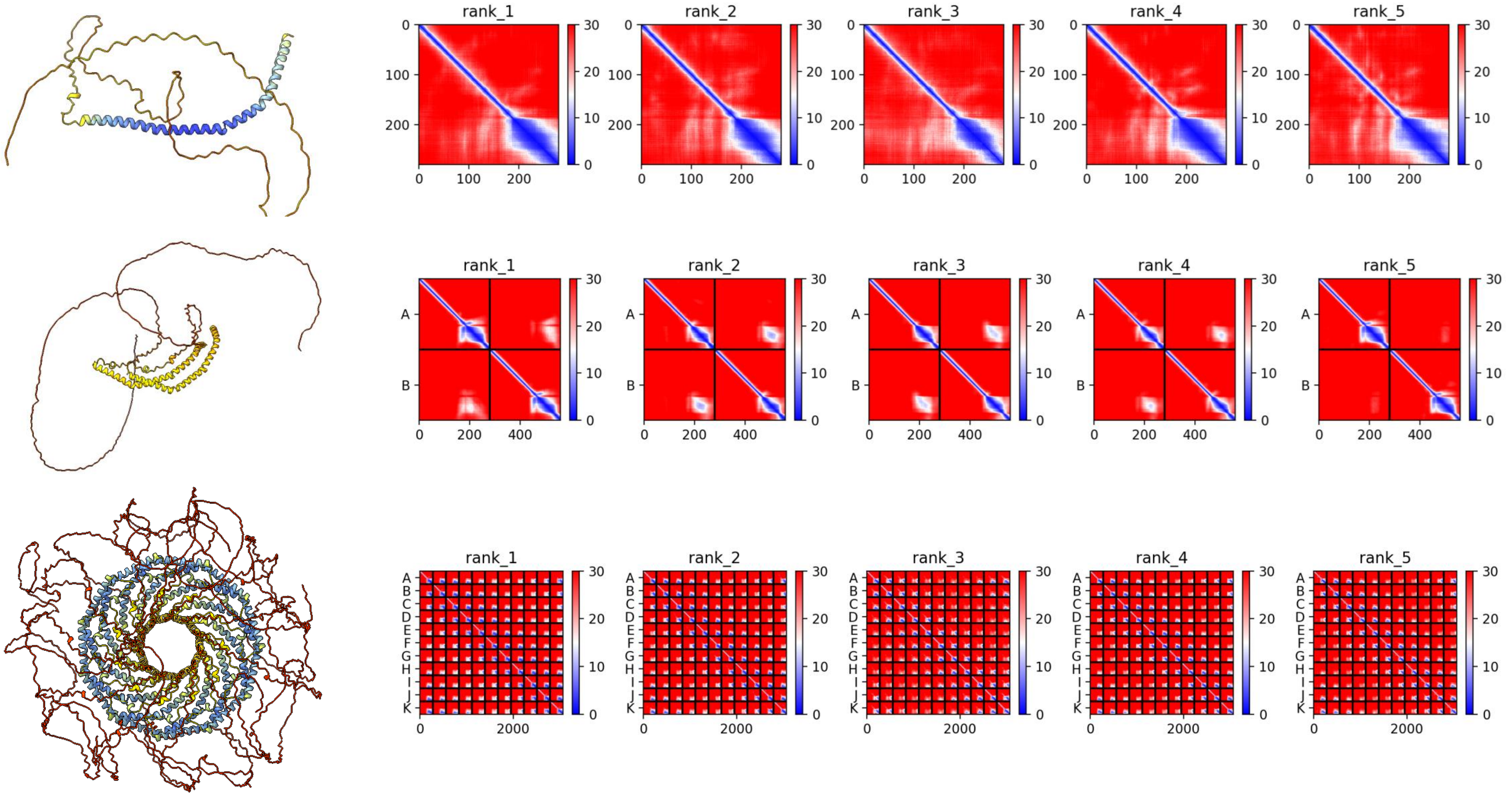

**A0A1X7VPY7** ***A. queenslandica***

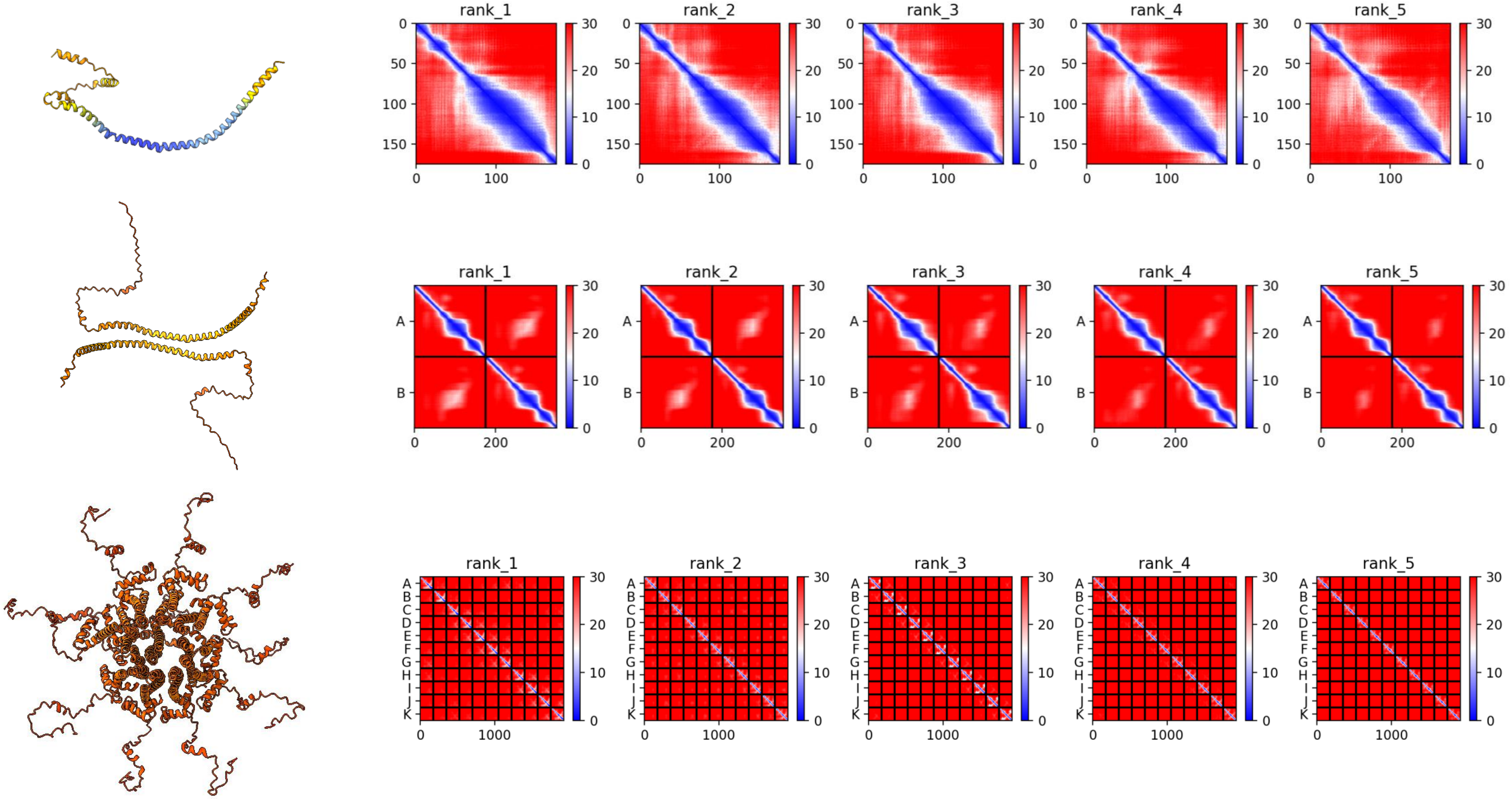

### A0A1X7VRV8 *A. queenslandica*

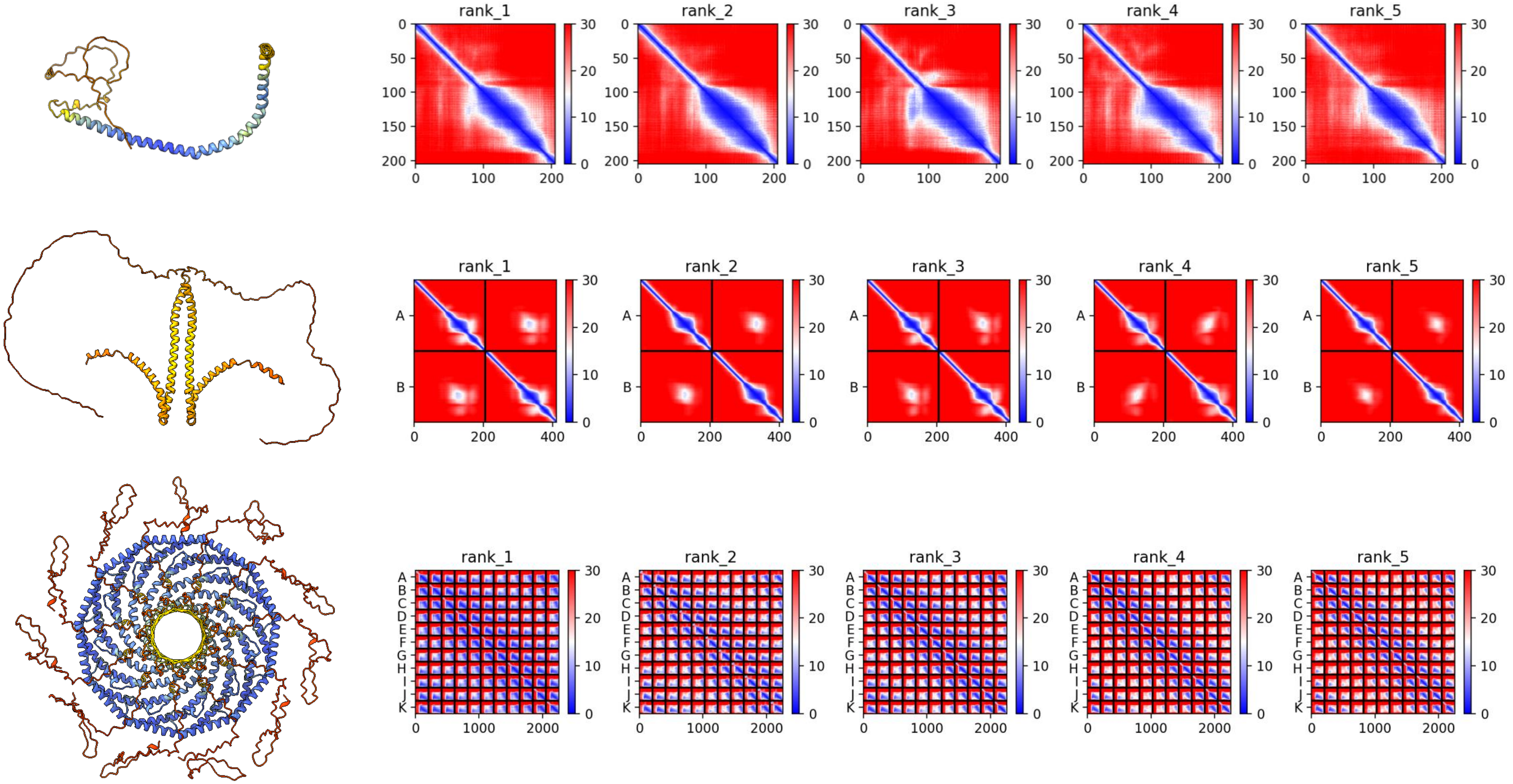

### A0A7M7HH57 *S. purpuratus*

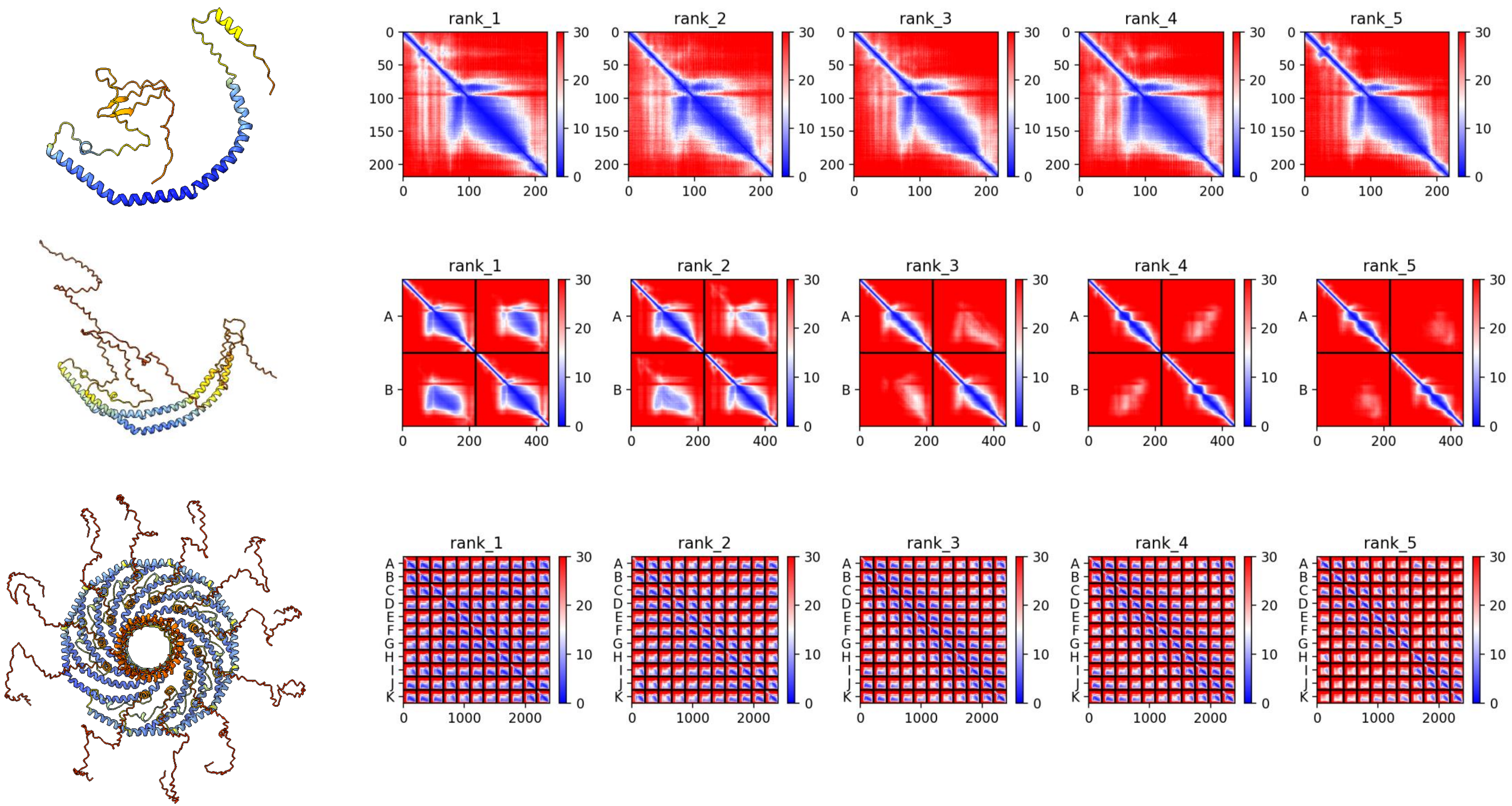

### R7U639 *C. teleta*

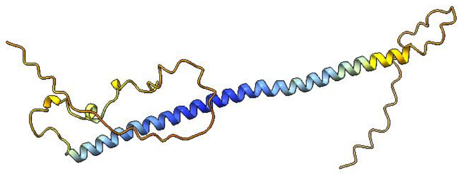

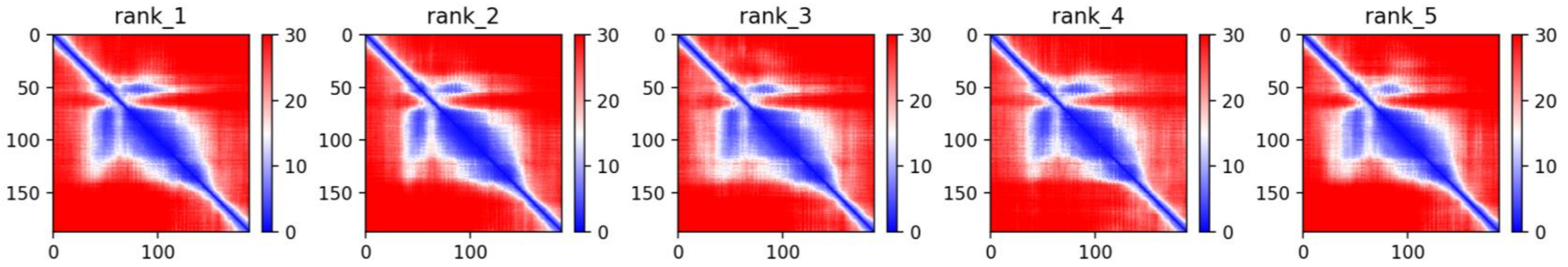


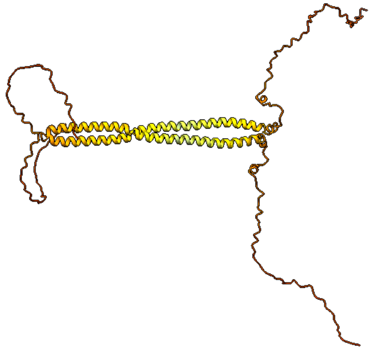

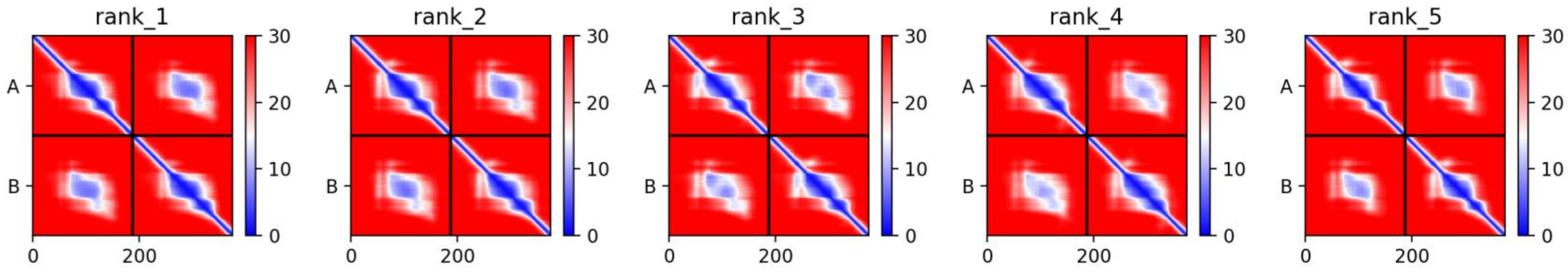


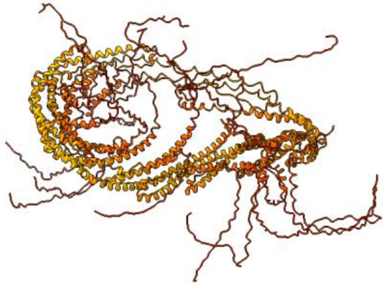

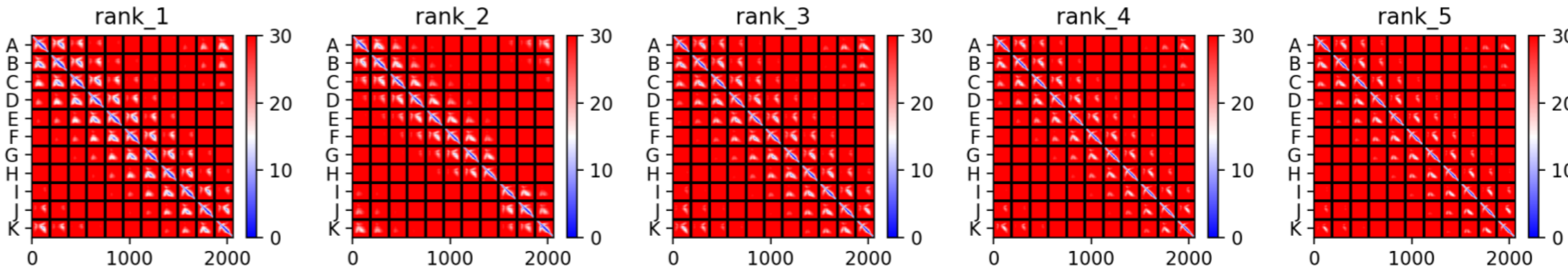

#### A0A7M7HEM1 *S. purpuratus*

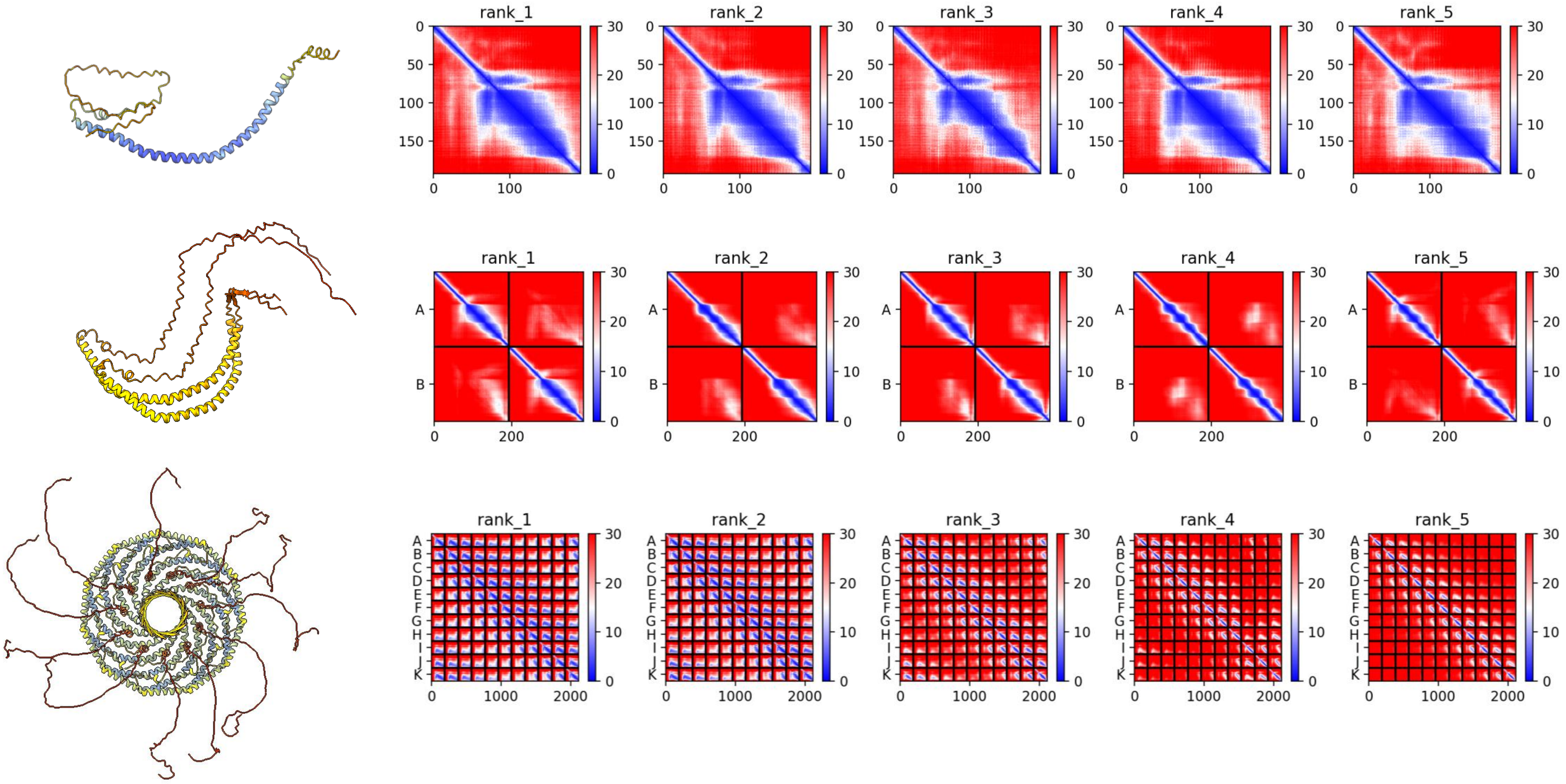

### **A0A7M7T4C2** ***S. purpuratus***

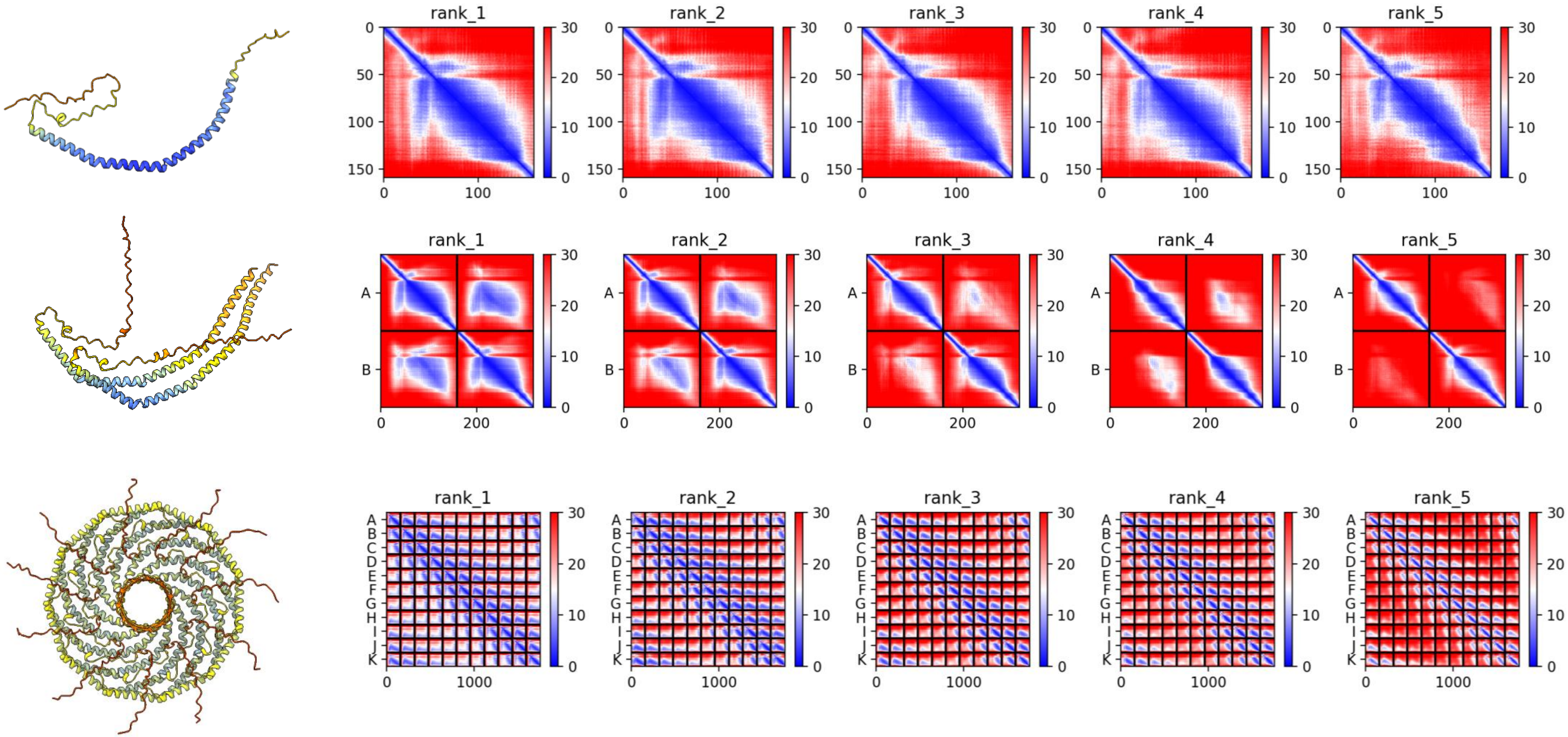

### A0A7M7G9F7 *S. purpuratus*

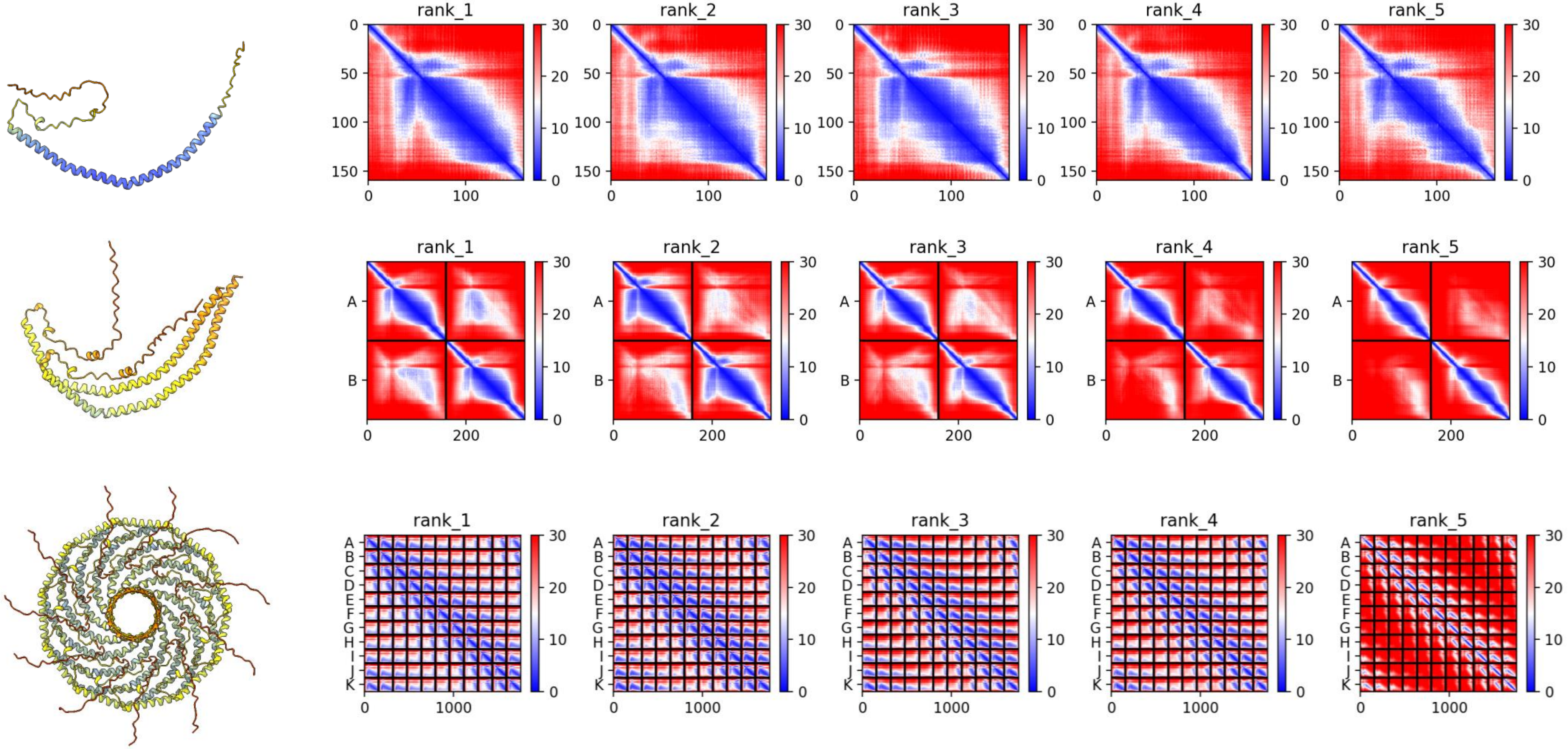

#### A0A7M7PLL3 *S. purpuratus*

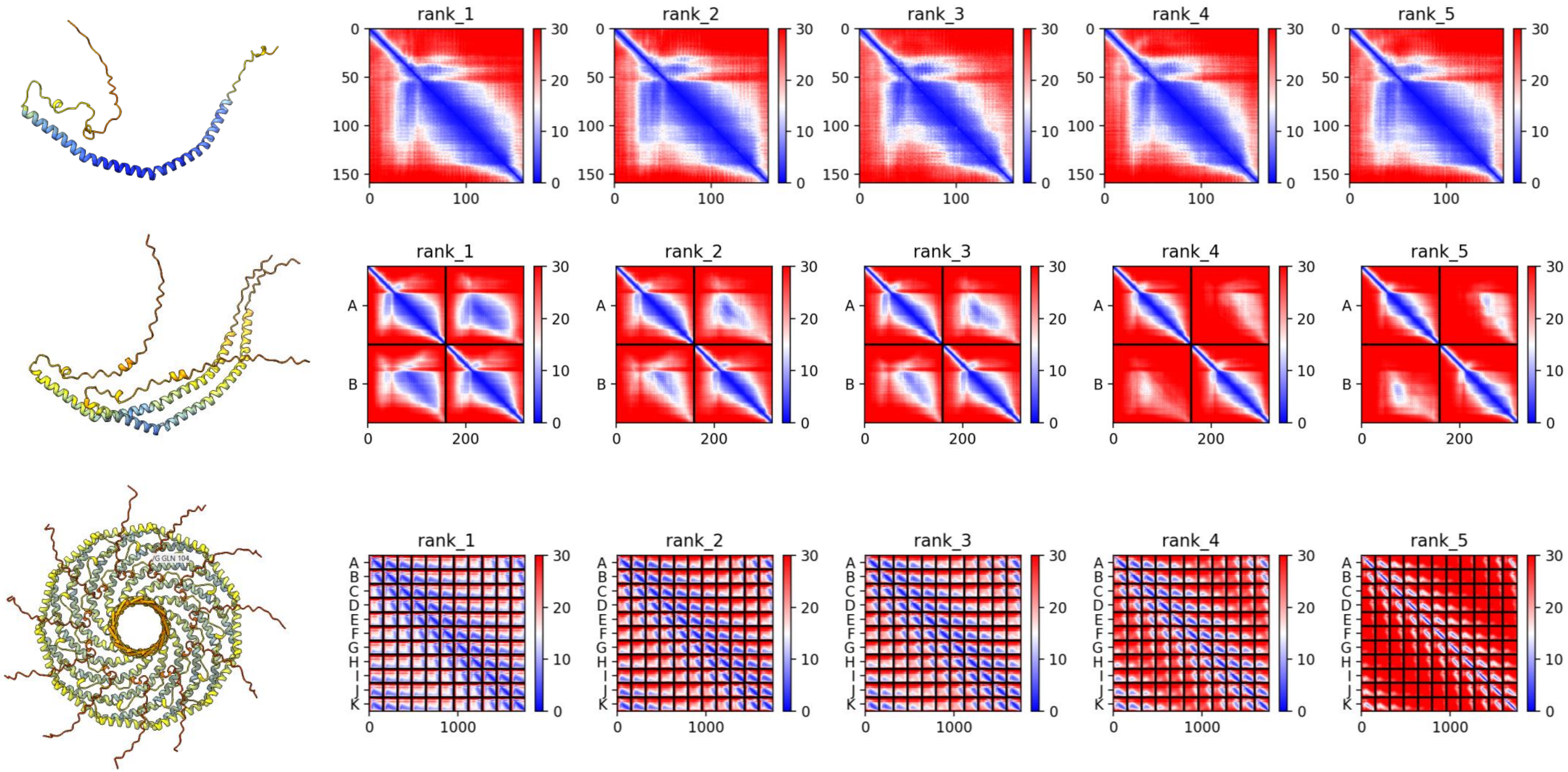

### Q94051 *C. elegans*

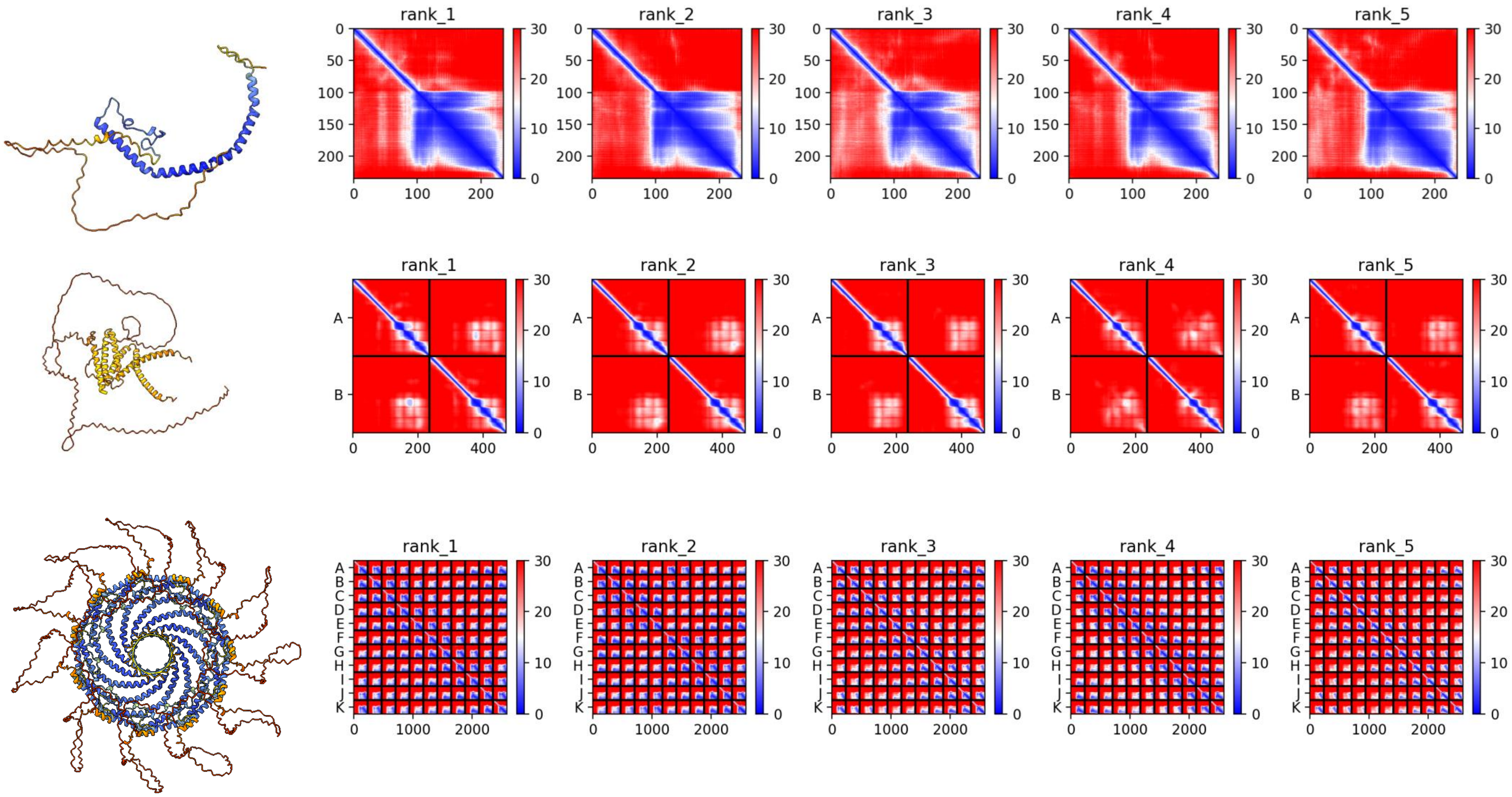

**A0A2T7NUA4** ***P. canaliculata***

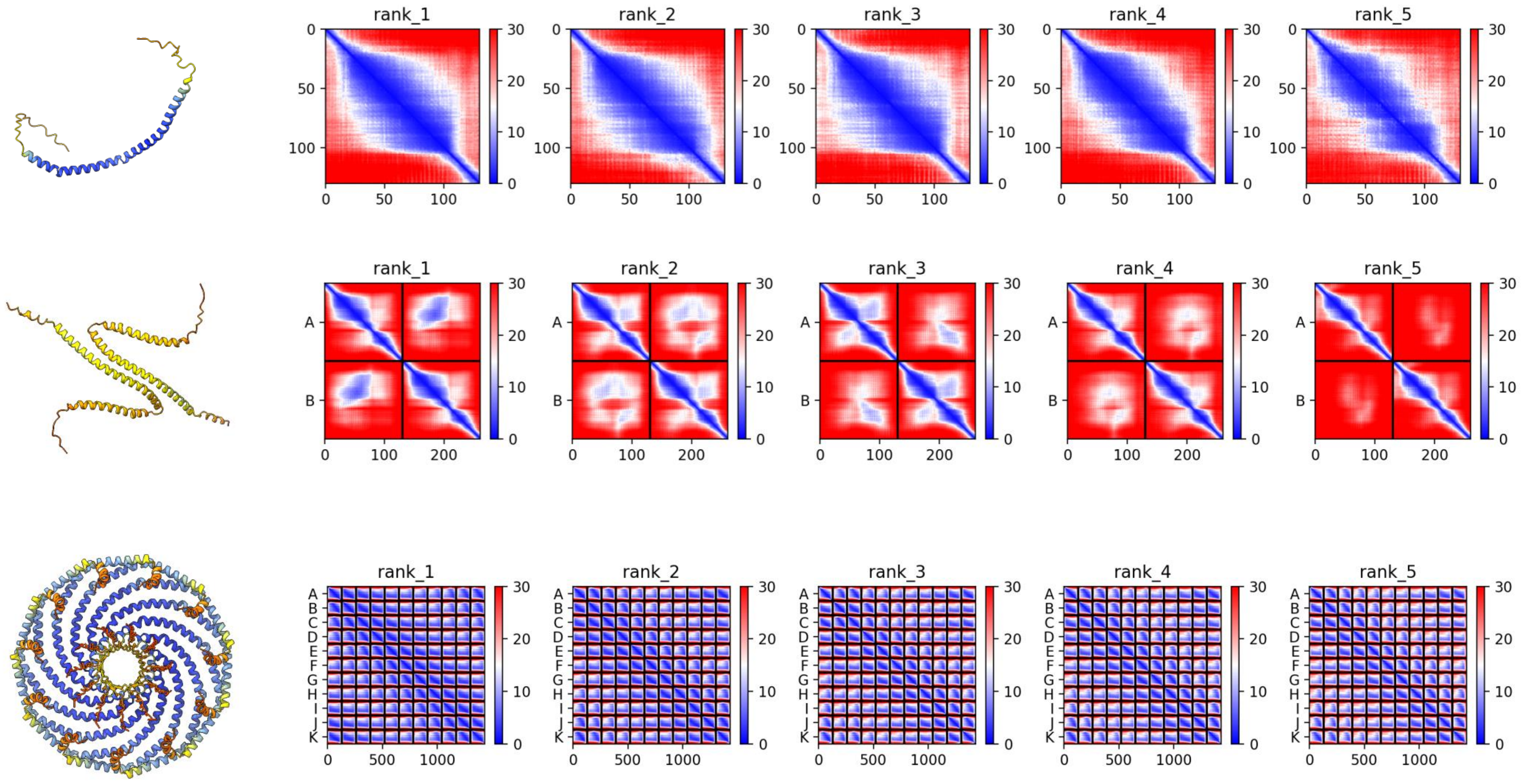

### Q18879 *C. elegans*

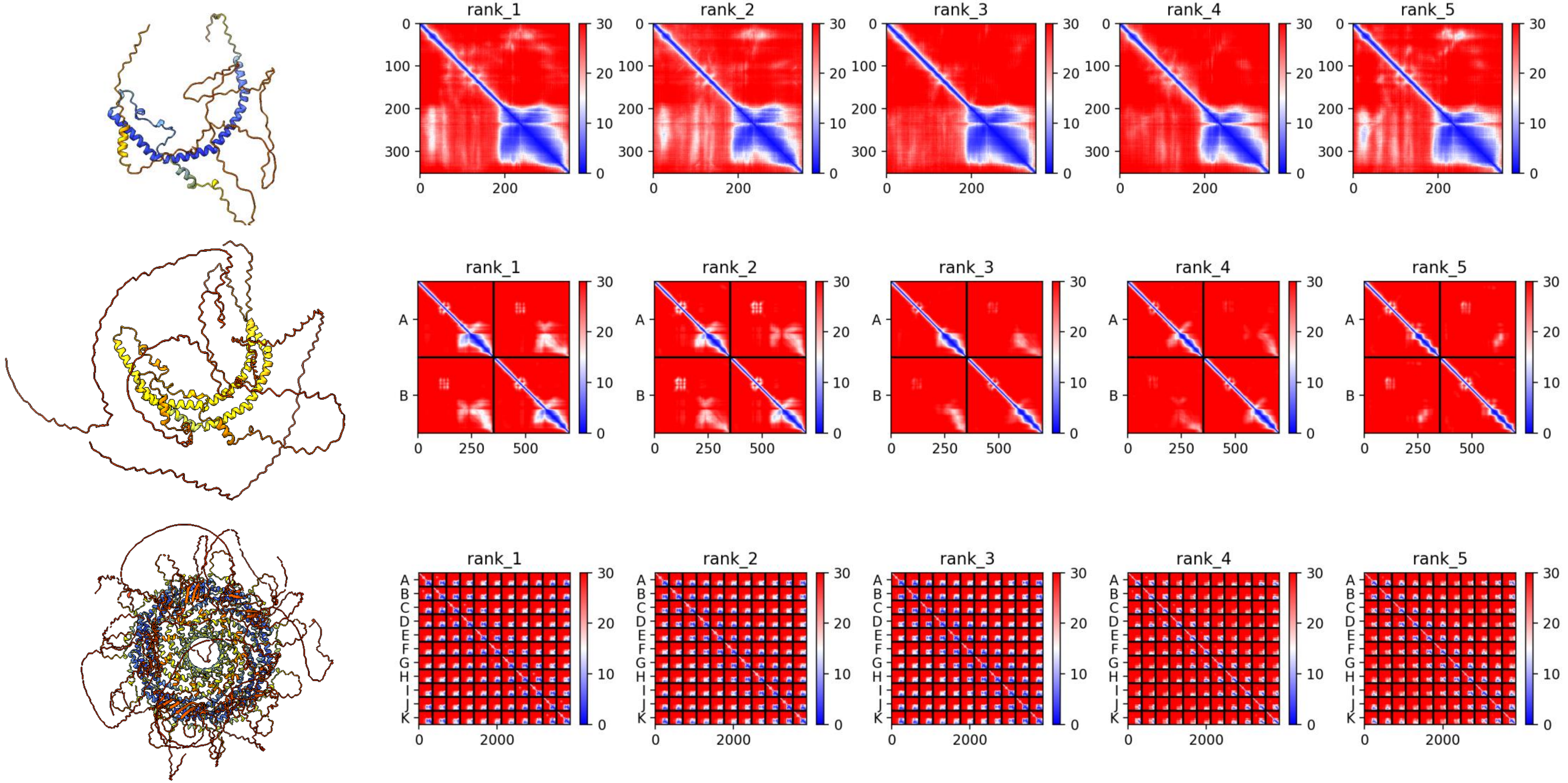

### A0A7M7GJ11 *S. purpuratus*

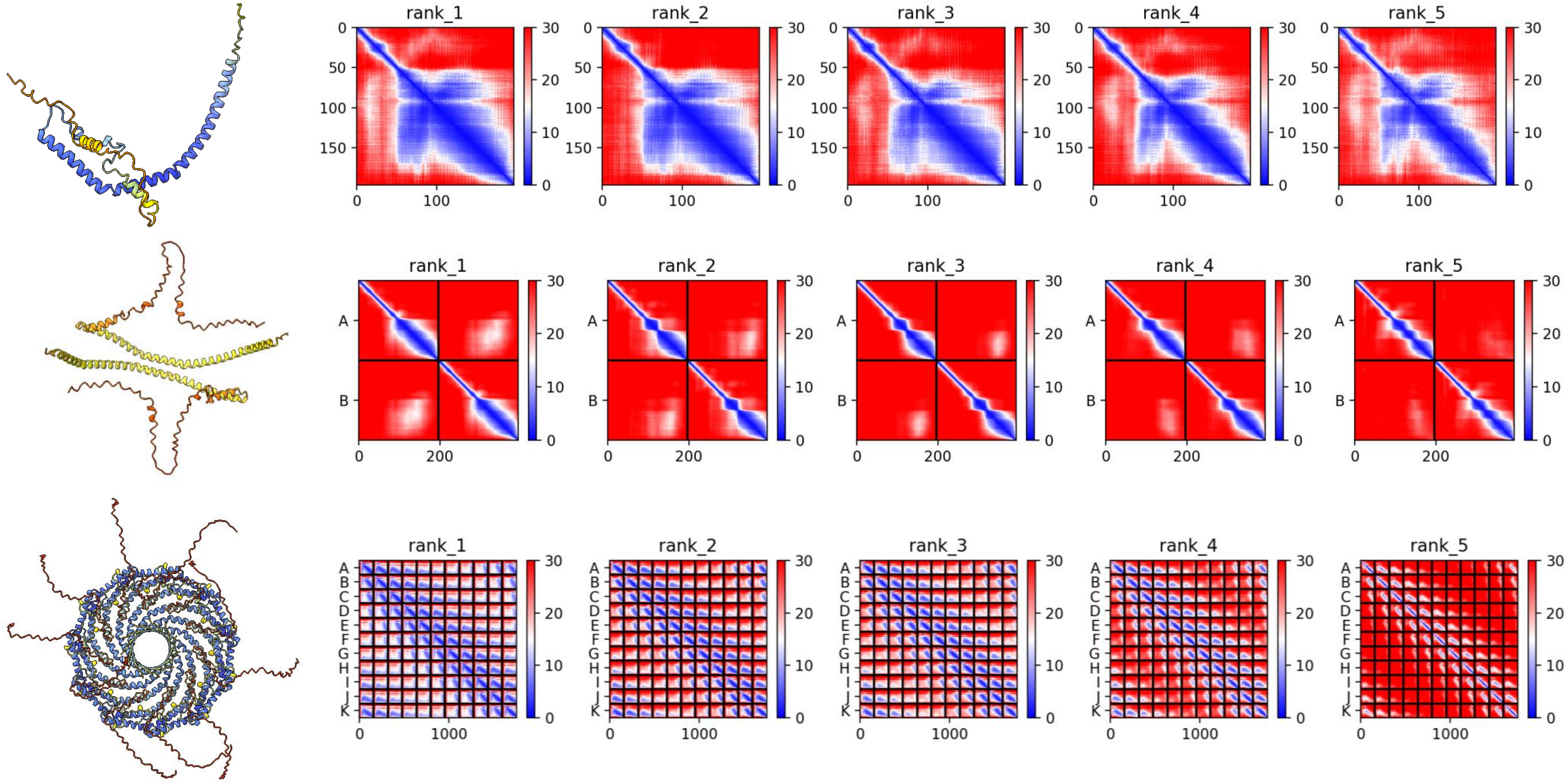

### A0A7M7P0X0 *S. purpuratus*

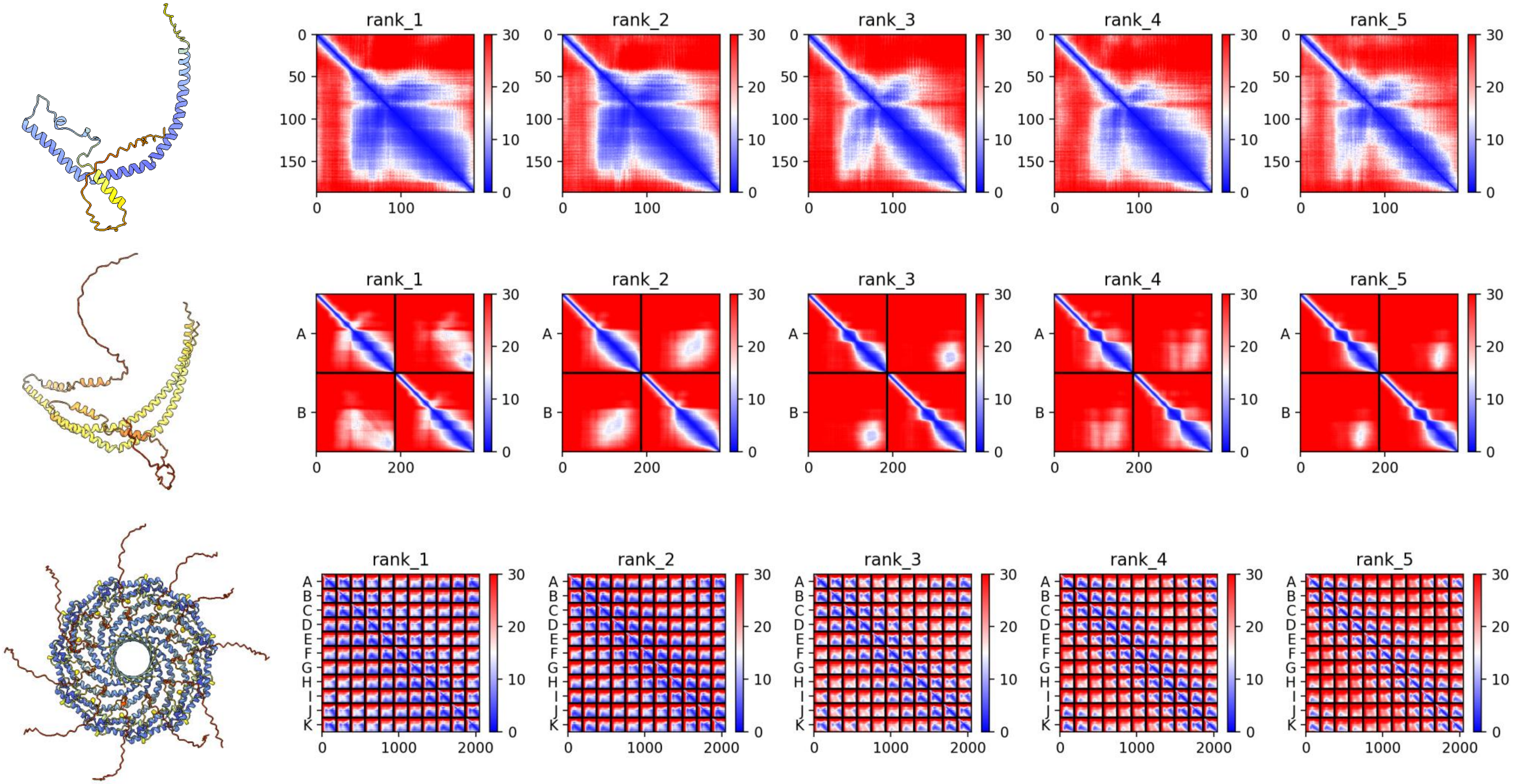

### B3RWV2 *T. adhaerens*

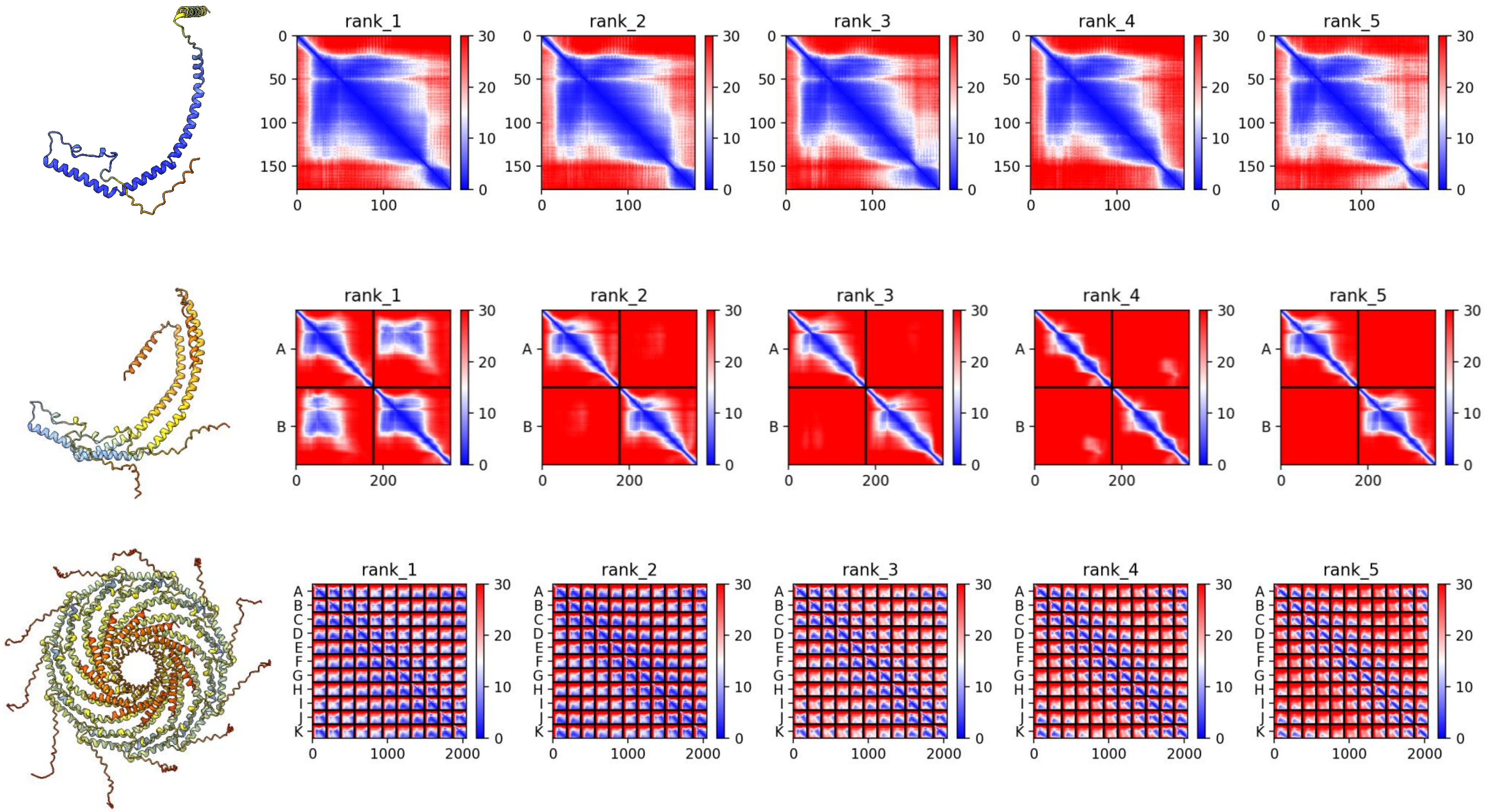

### R7TPM2 *C. teleta*

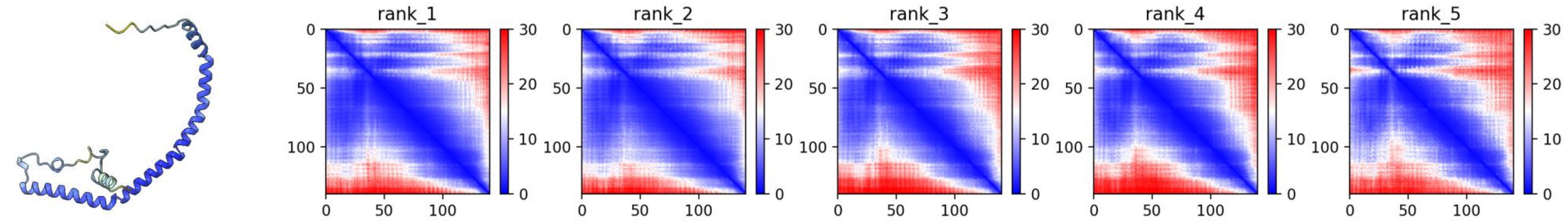


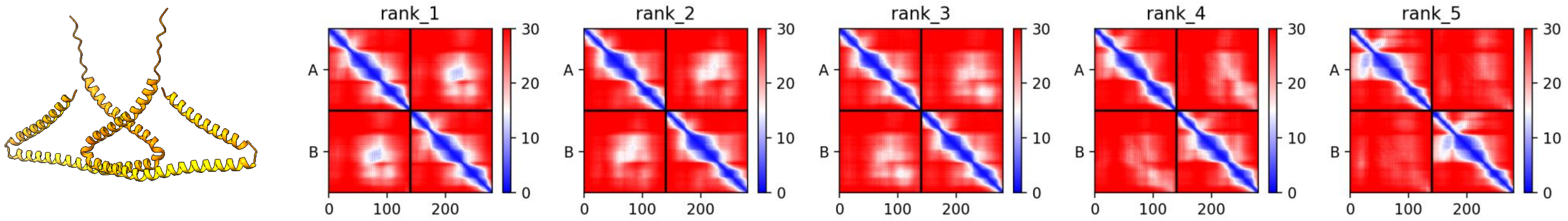


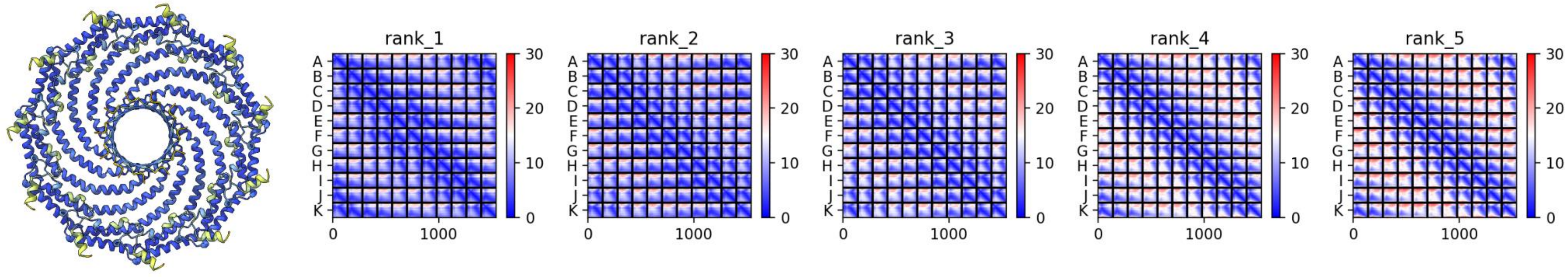

### A0A1I8GQY0 *M. lignano*

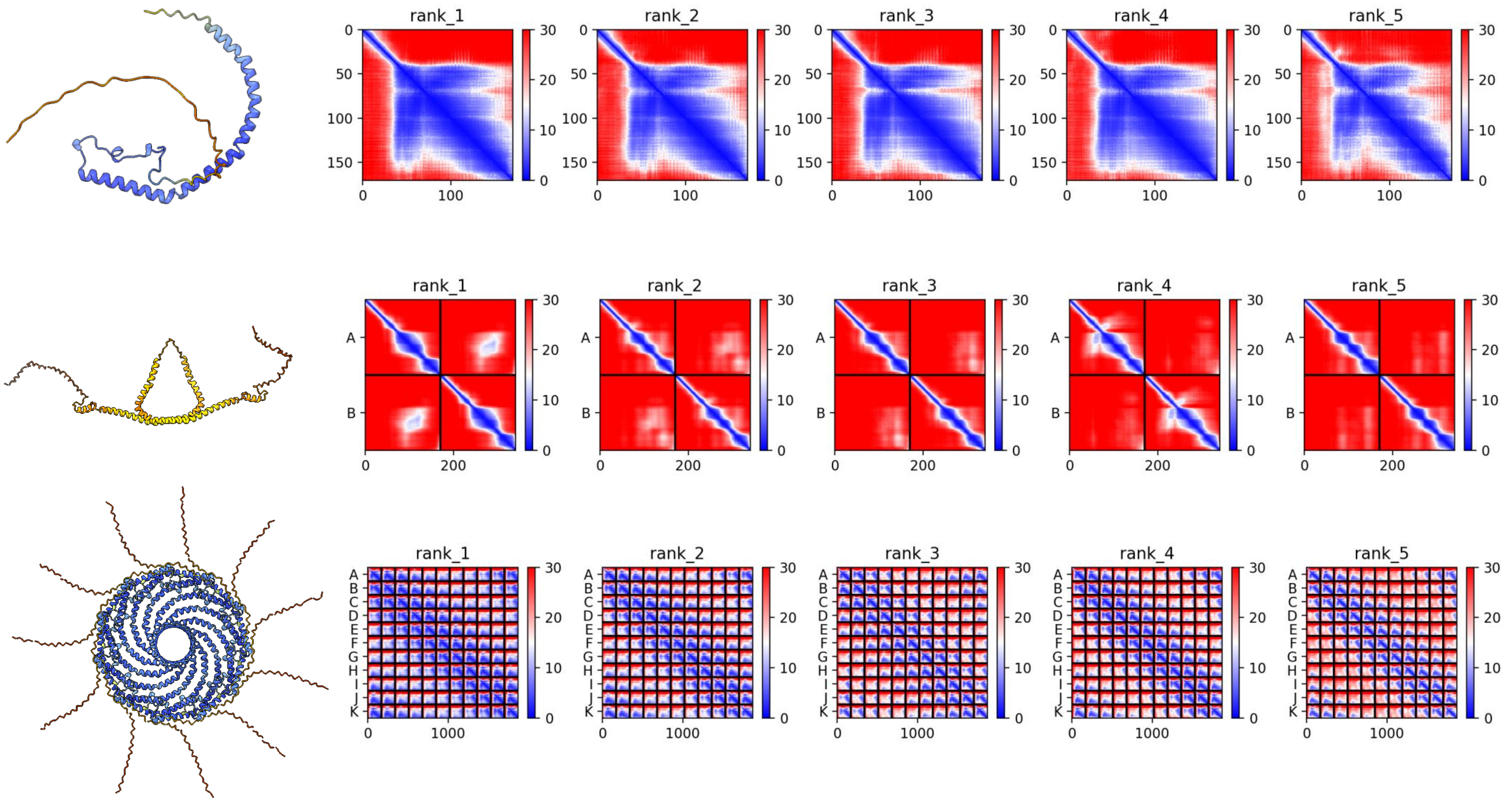

### A0A267DFI9 *M. lignano*

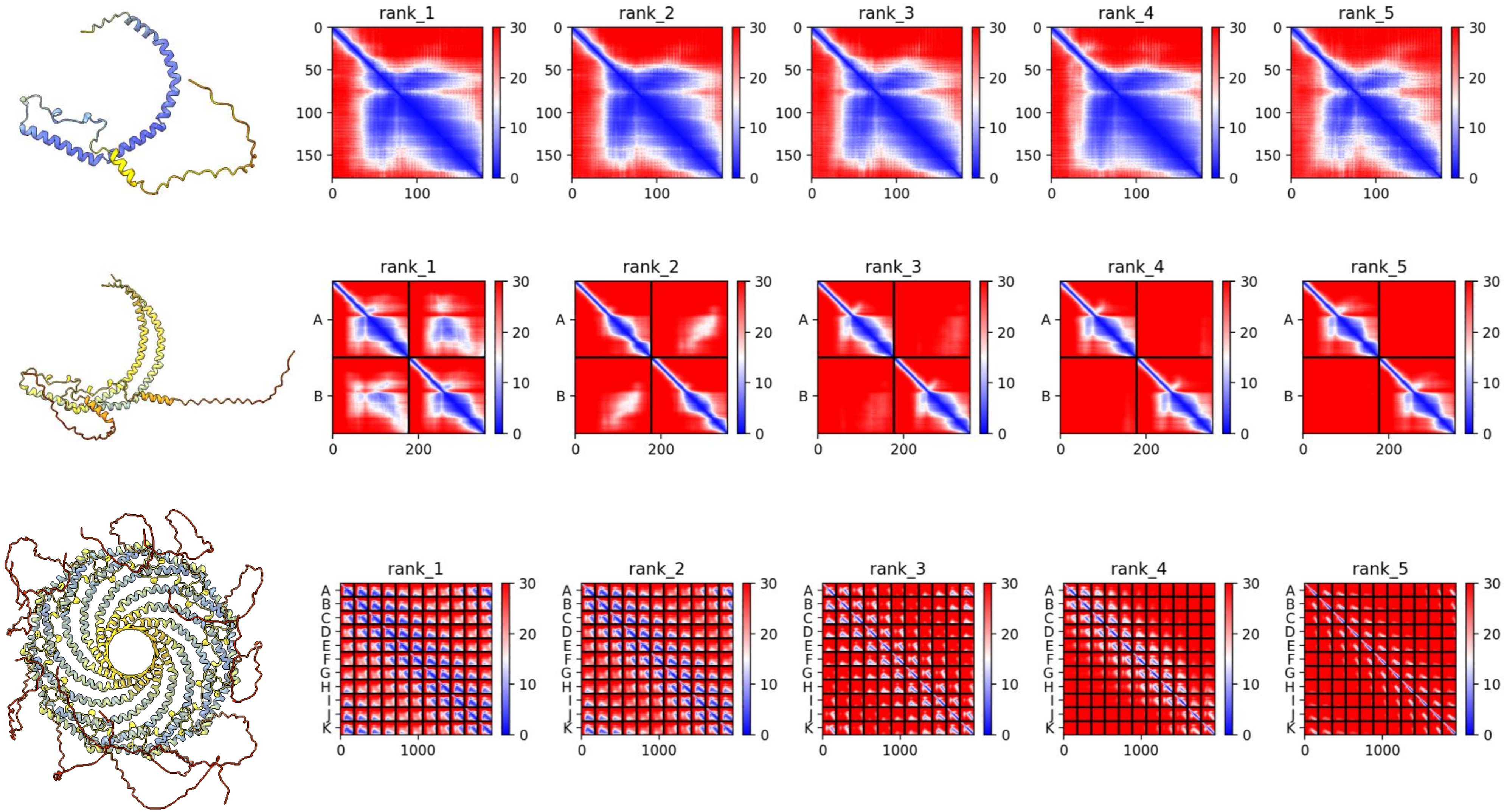

### A0A267G3E9 *M. lignano*

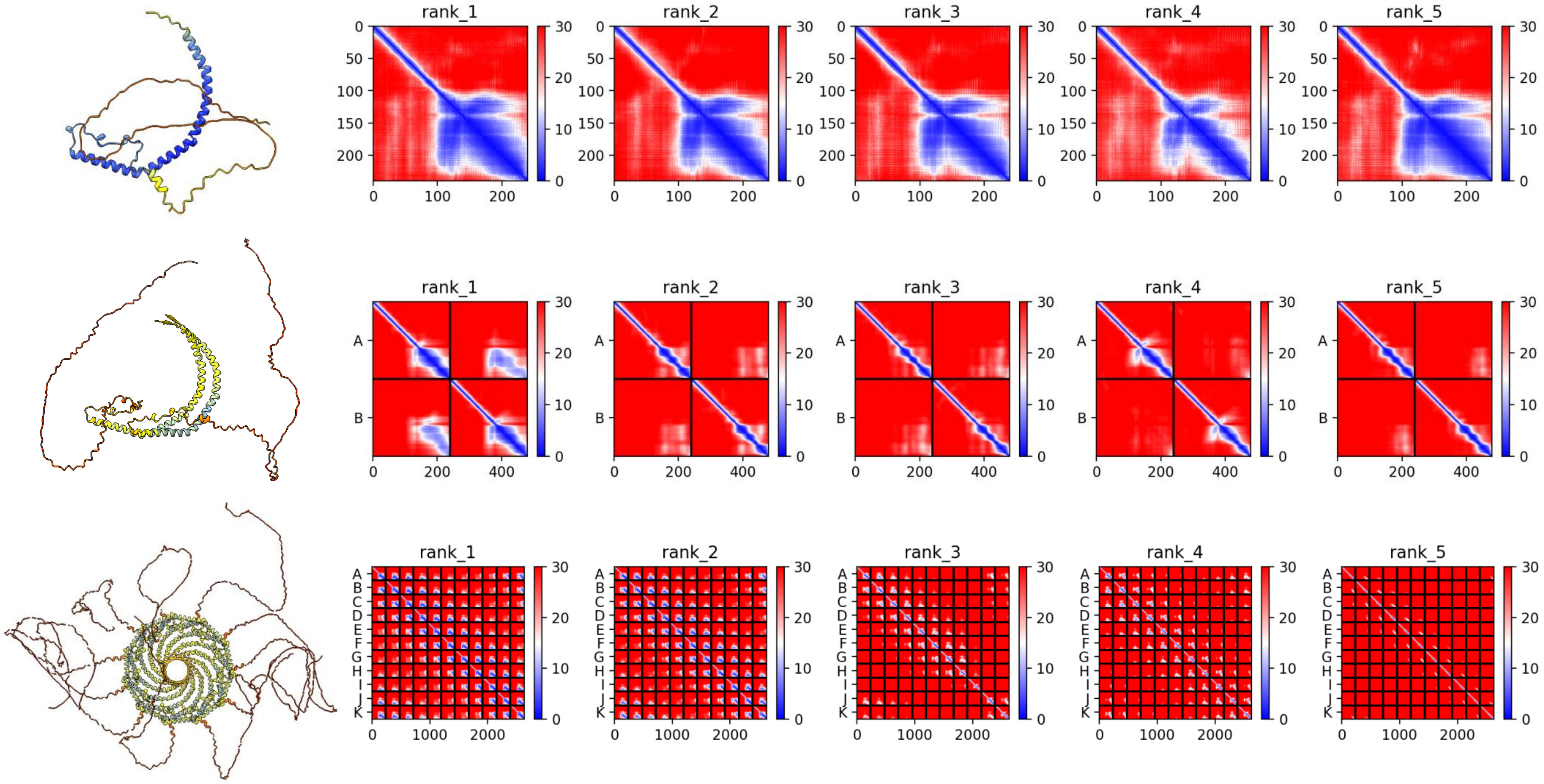

### A0A267GQL3 *M. lignano*

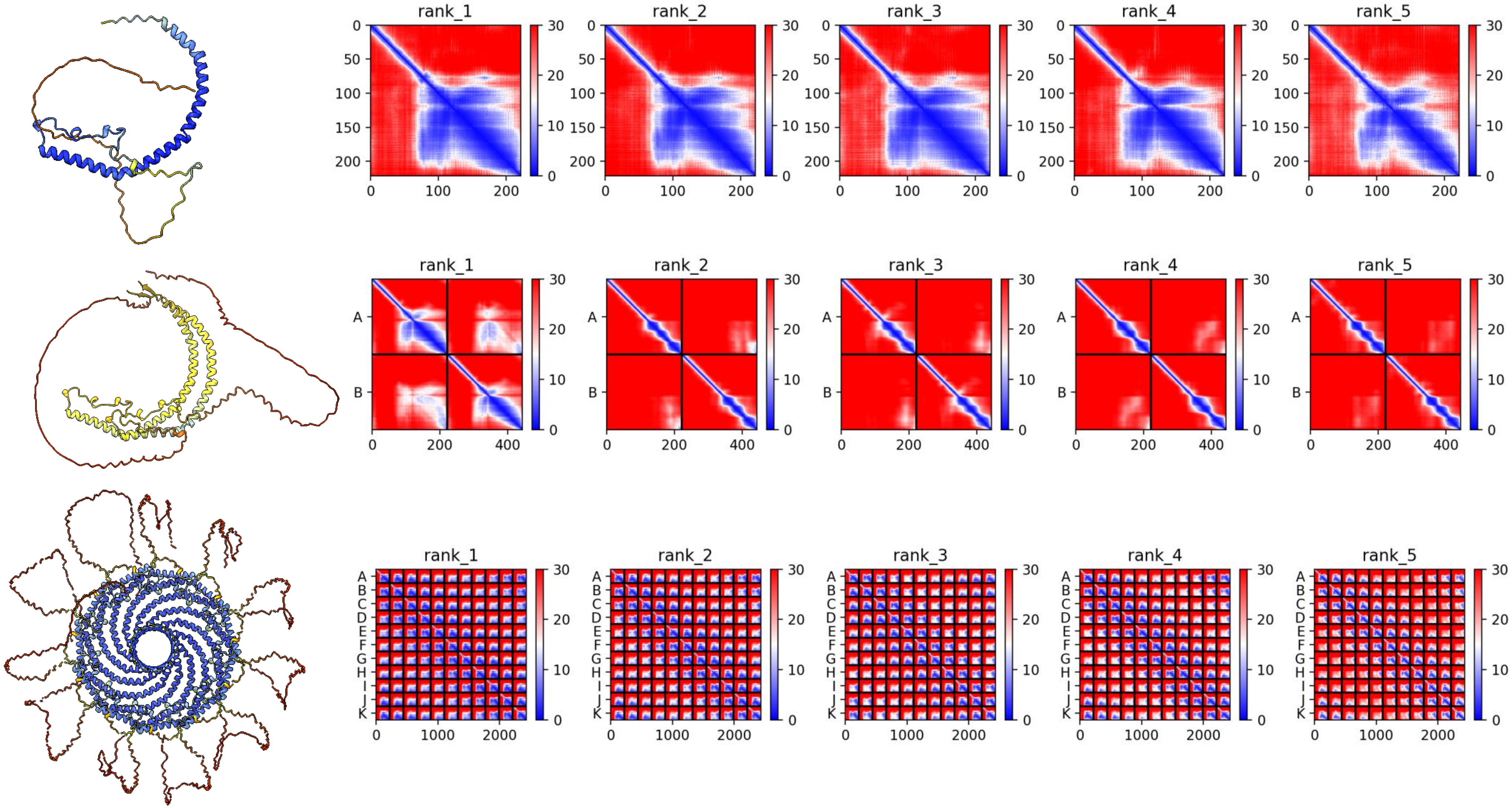

### A0A267G3D8 *M. lignano*

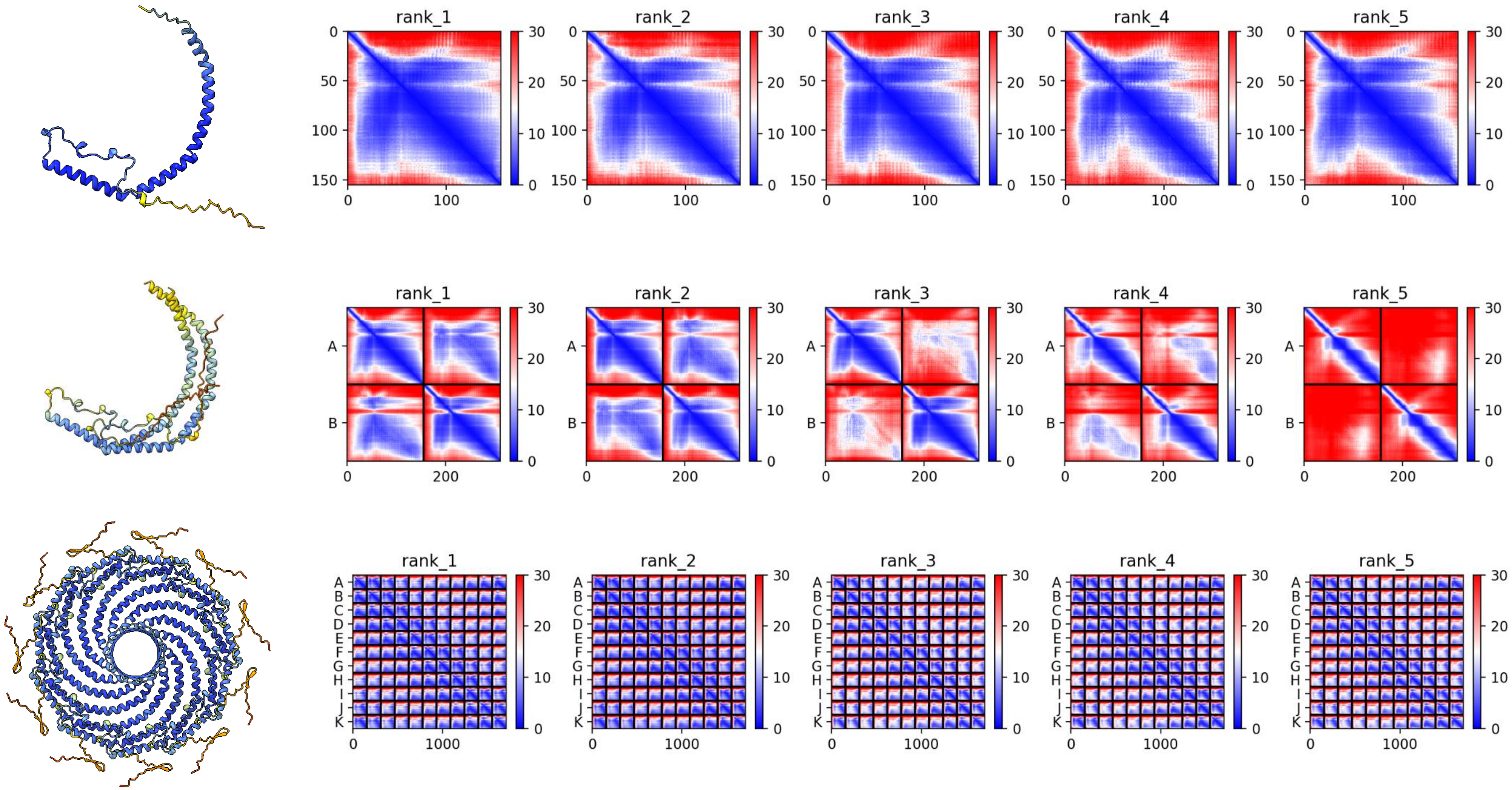

### A0A267G387 *M. lignano*

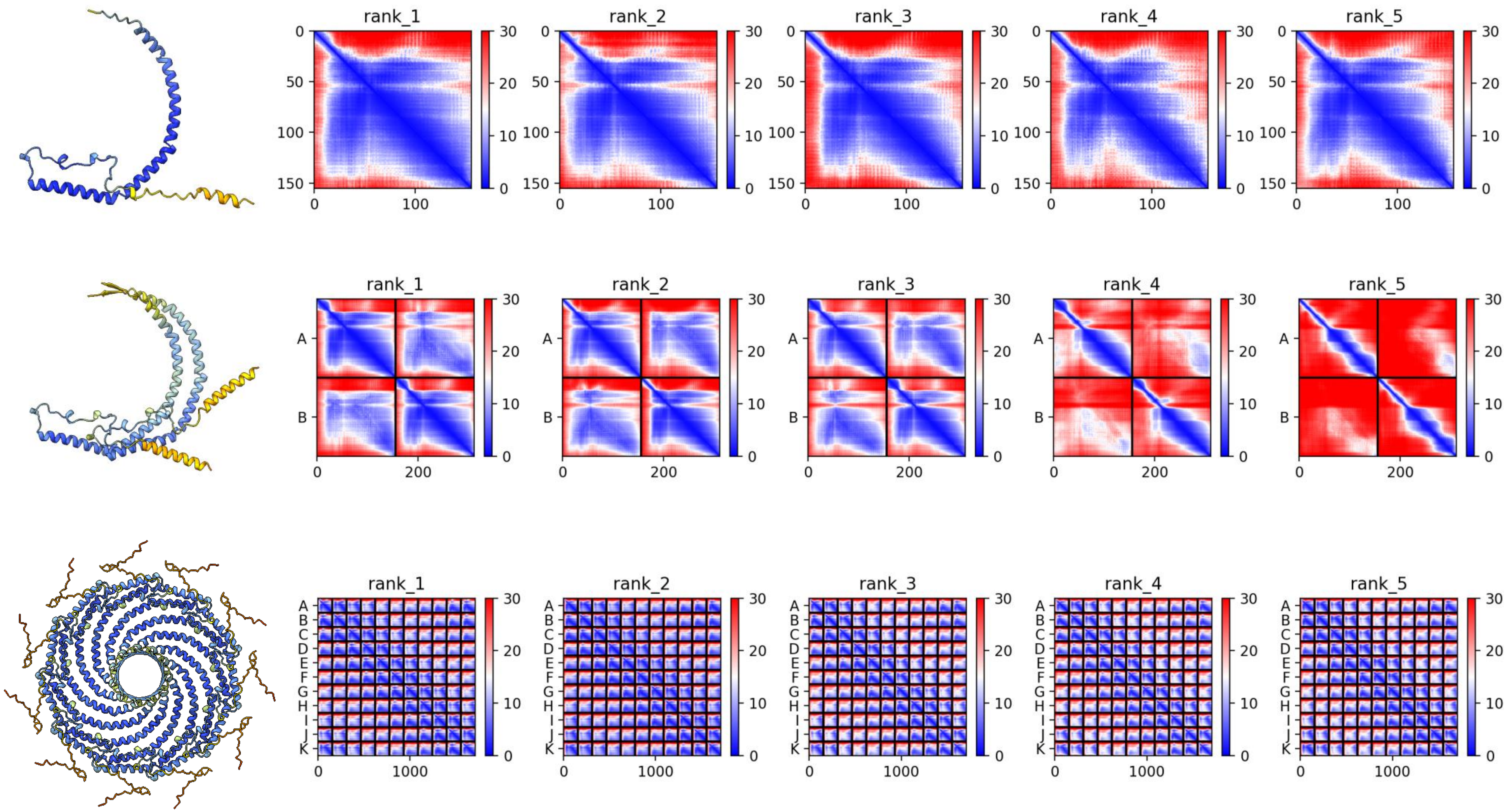

### A0A267G1N3 *M. lignano*

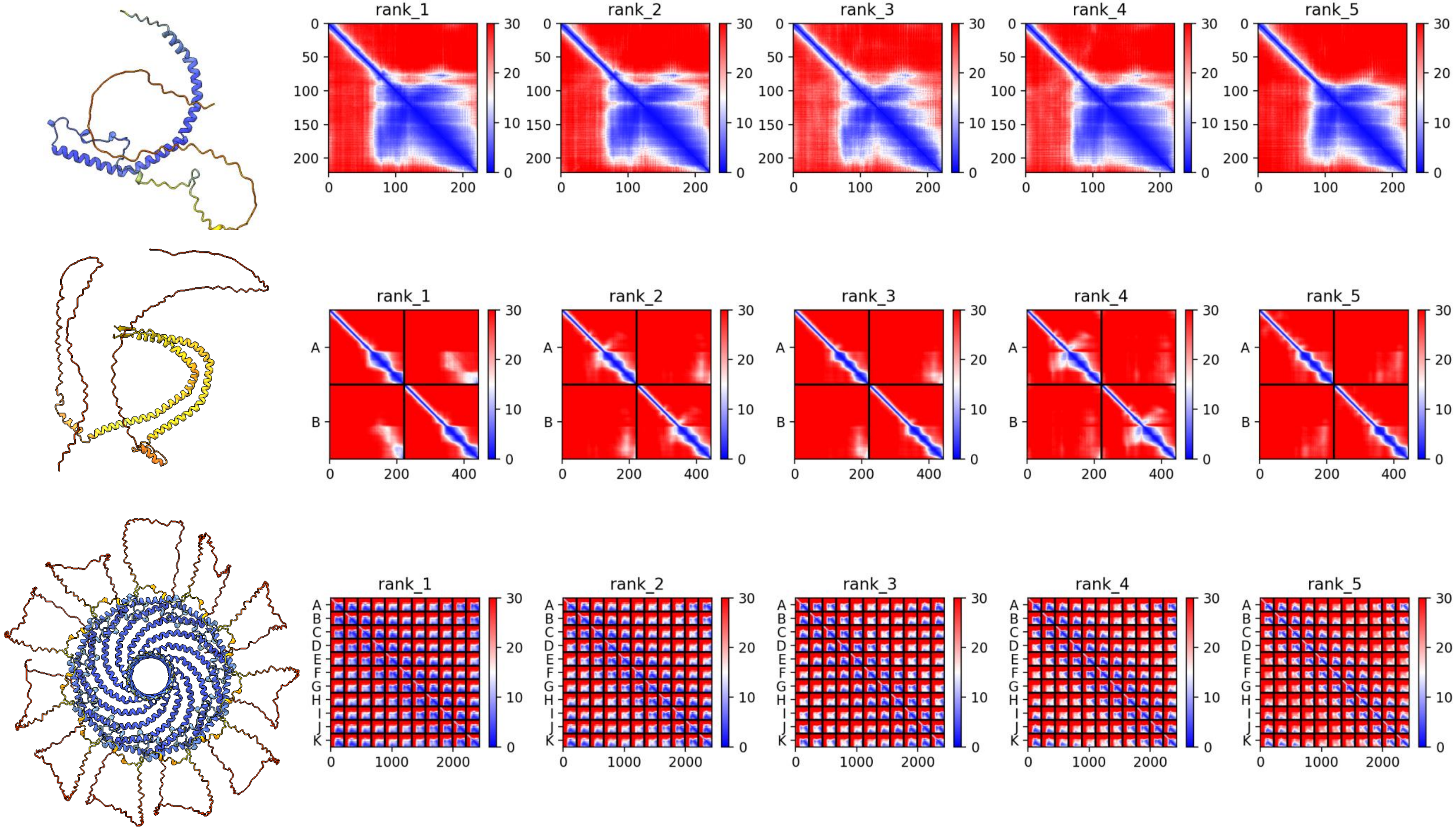

### A0A267DNA3 *M. lignano*

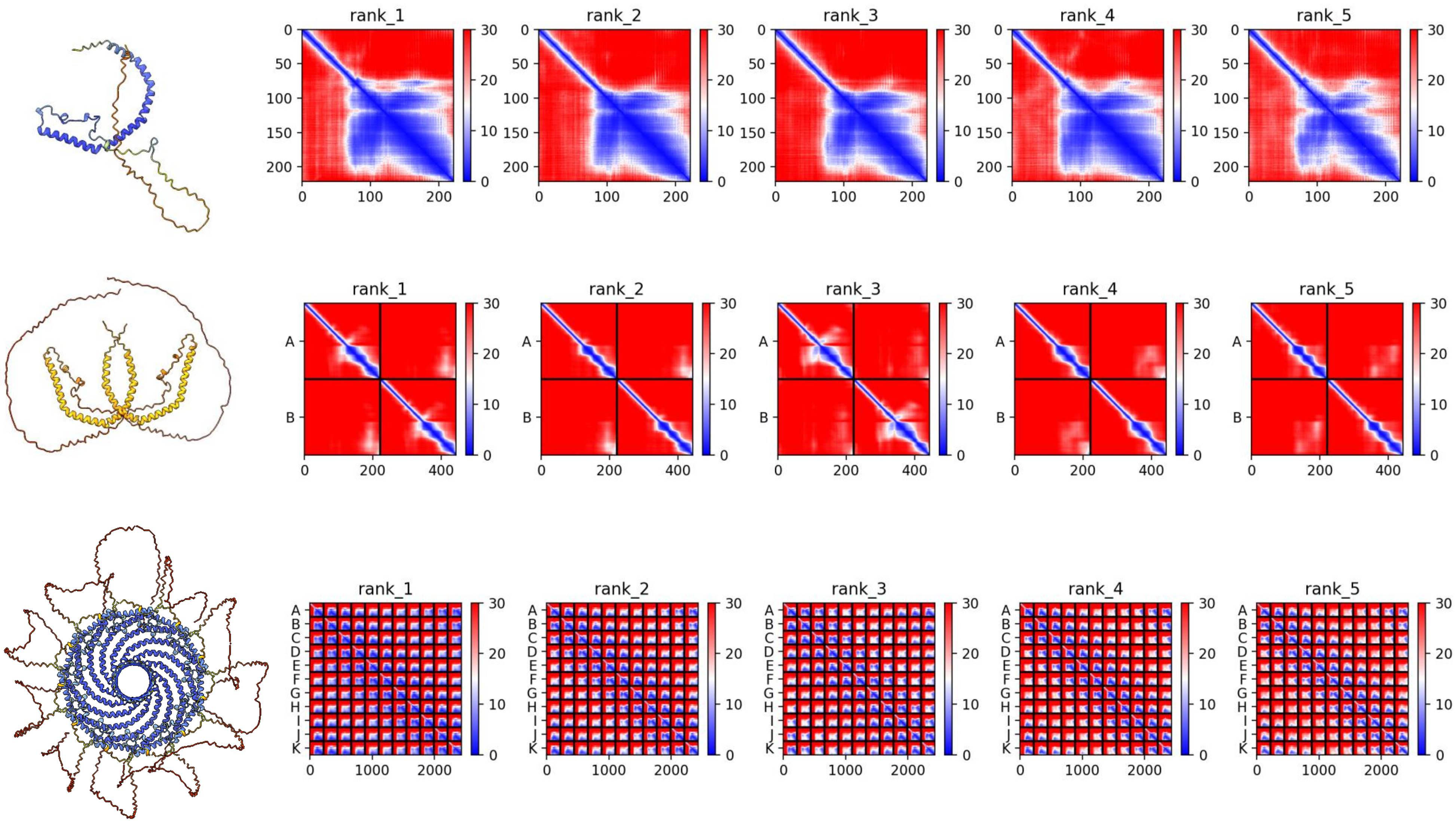

### A0A267FMS2 *M. lignano*

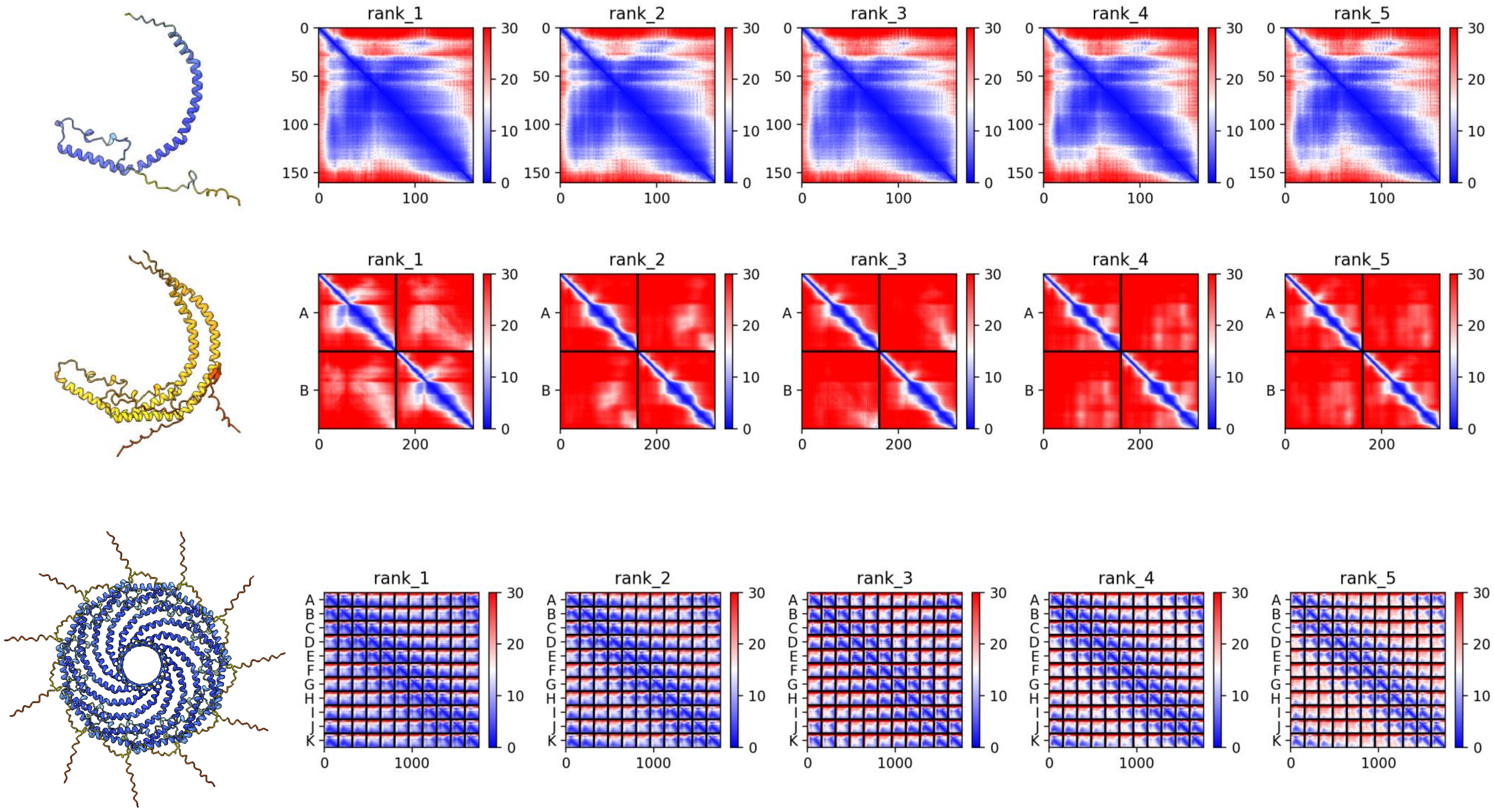

### XP_014664738.1 *P. caudatus*

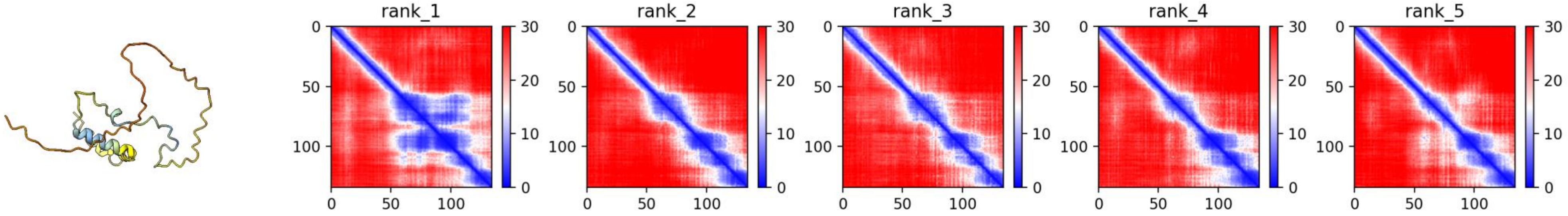


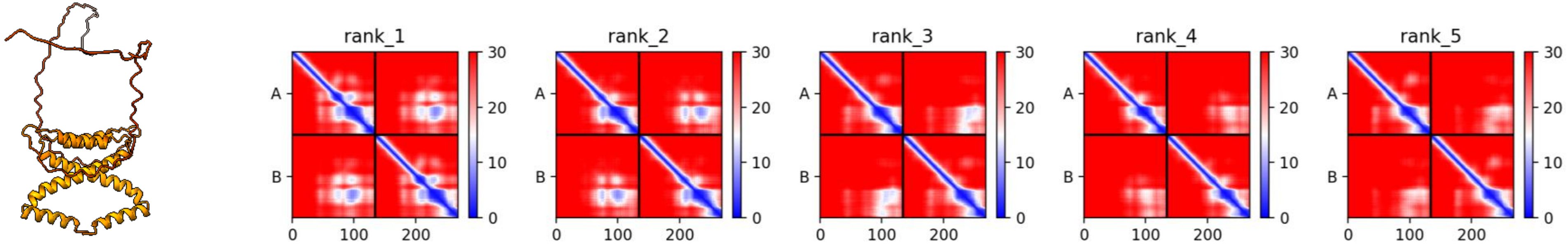


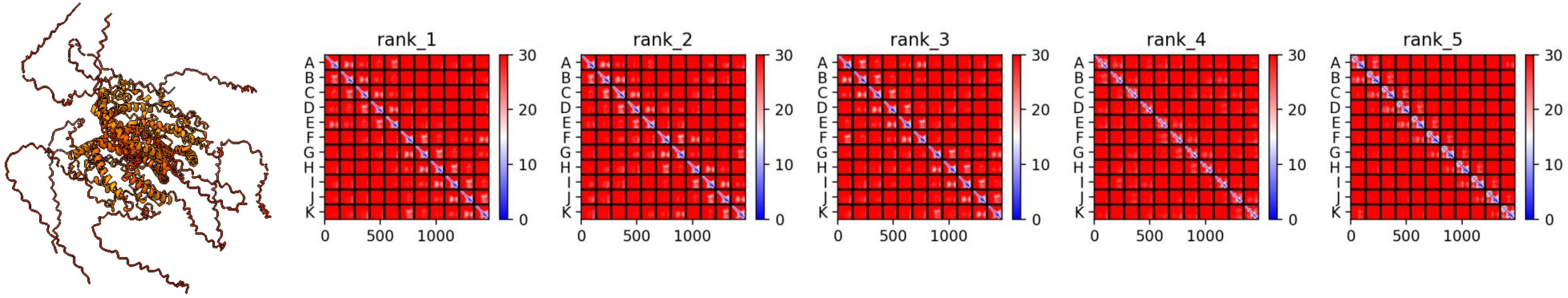

### A0A1S3IXS7 *L. unguis*

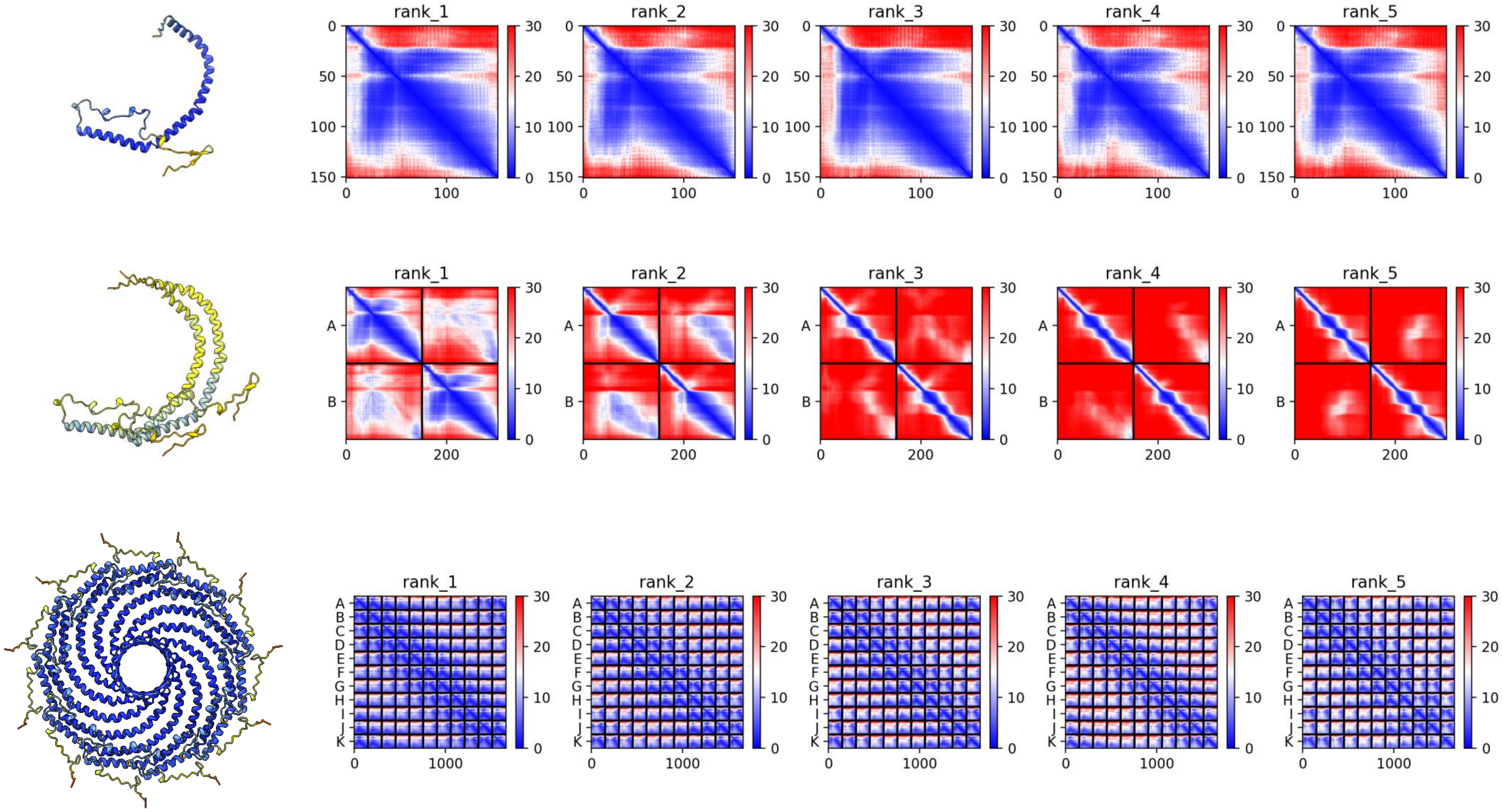

### A0A1S3I8M7 *L. unguis*

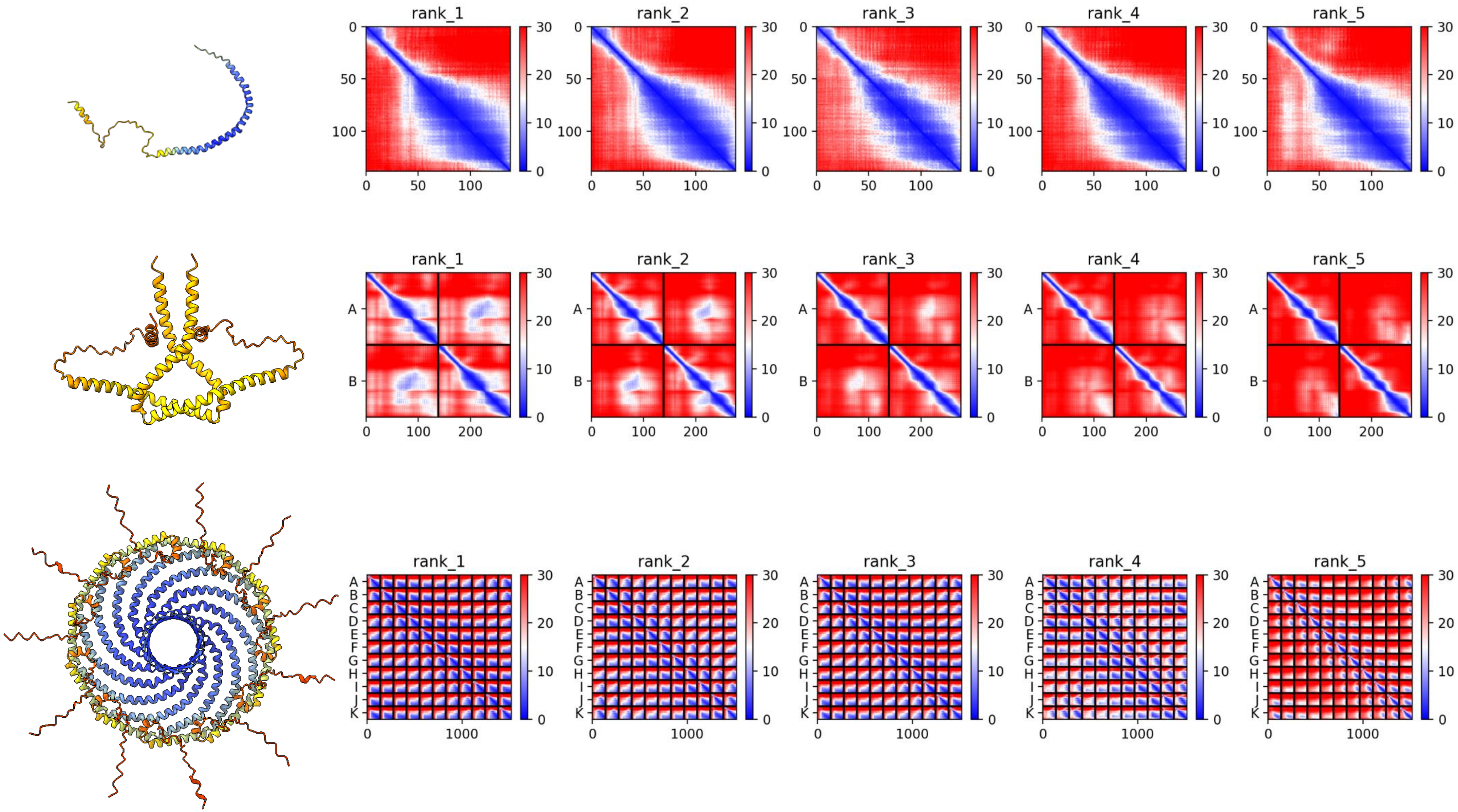

**A0A1S3I469 *L. unguis***

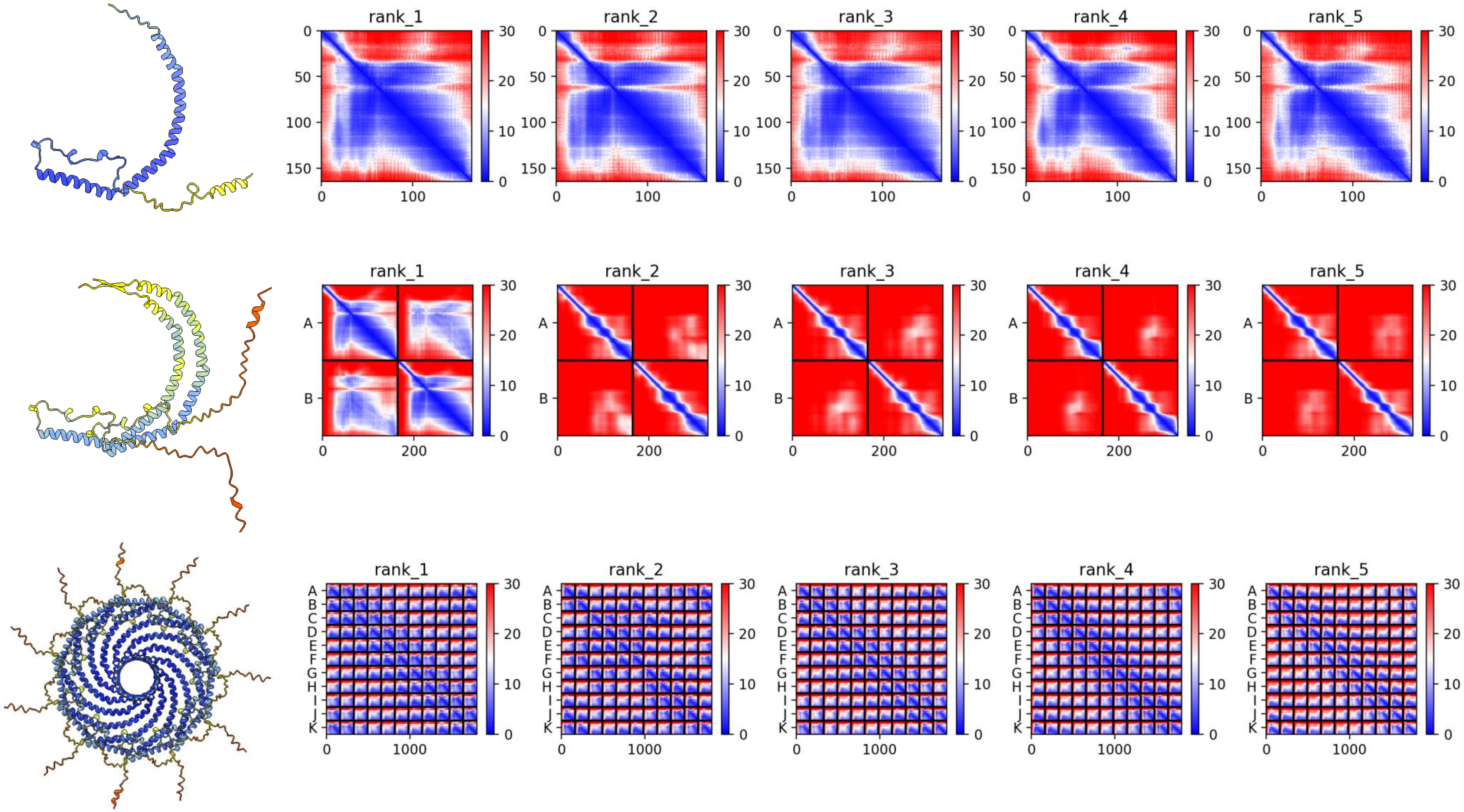

### P51636 (CAV2) *H. sapiens*

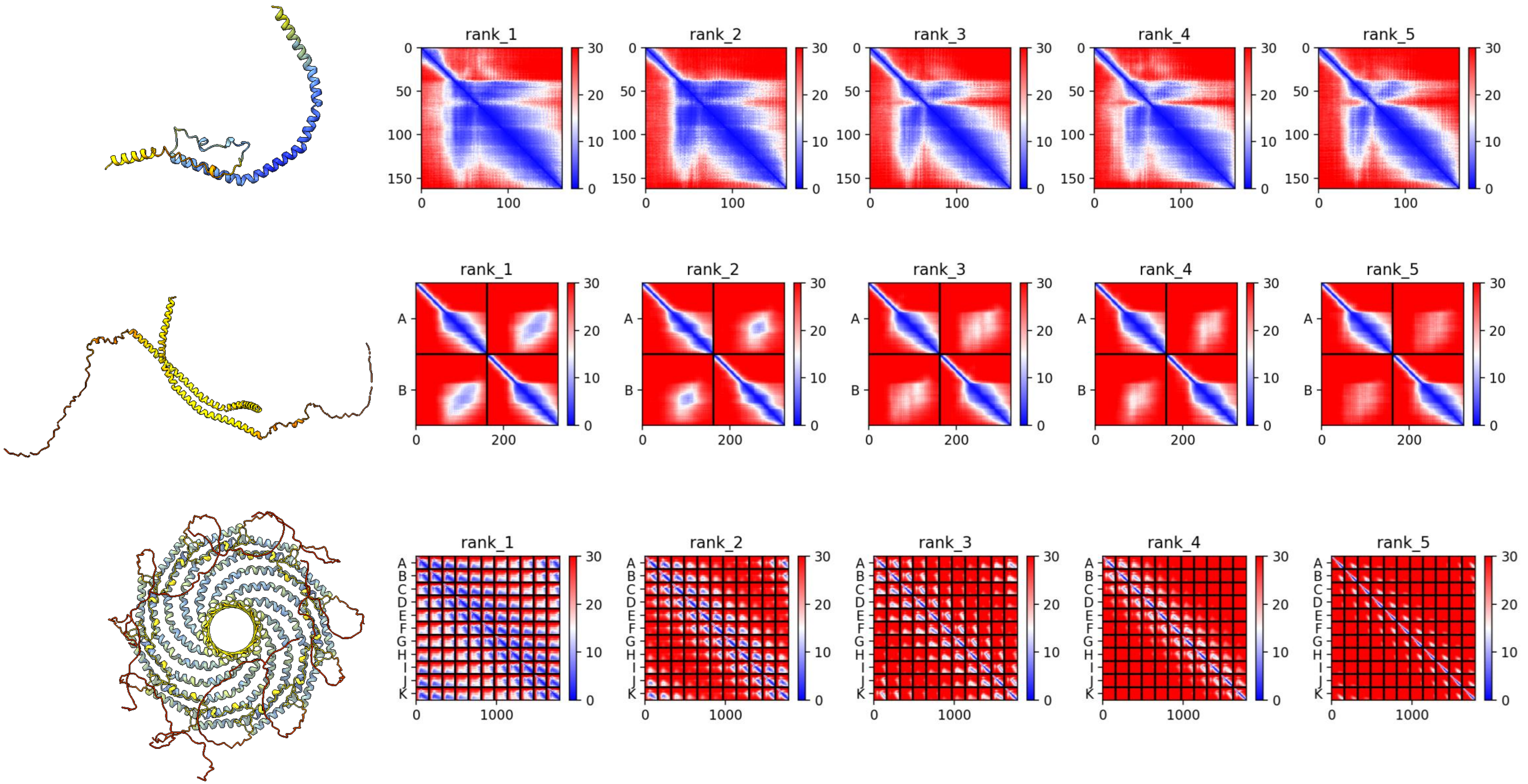

### P56539 (CAV3) *H. sapiens*

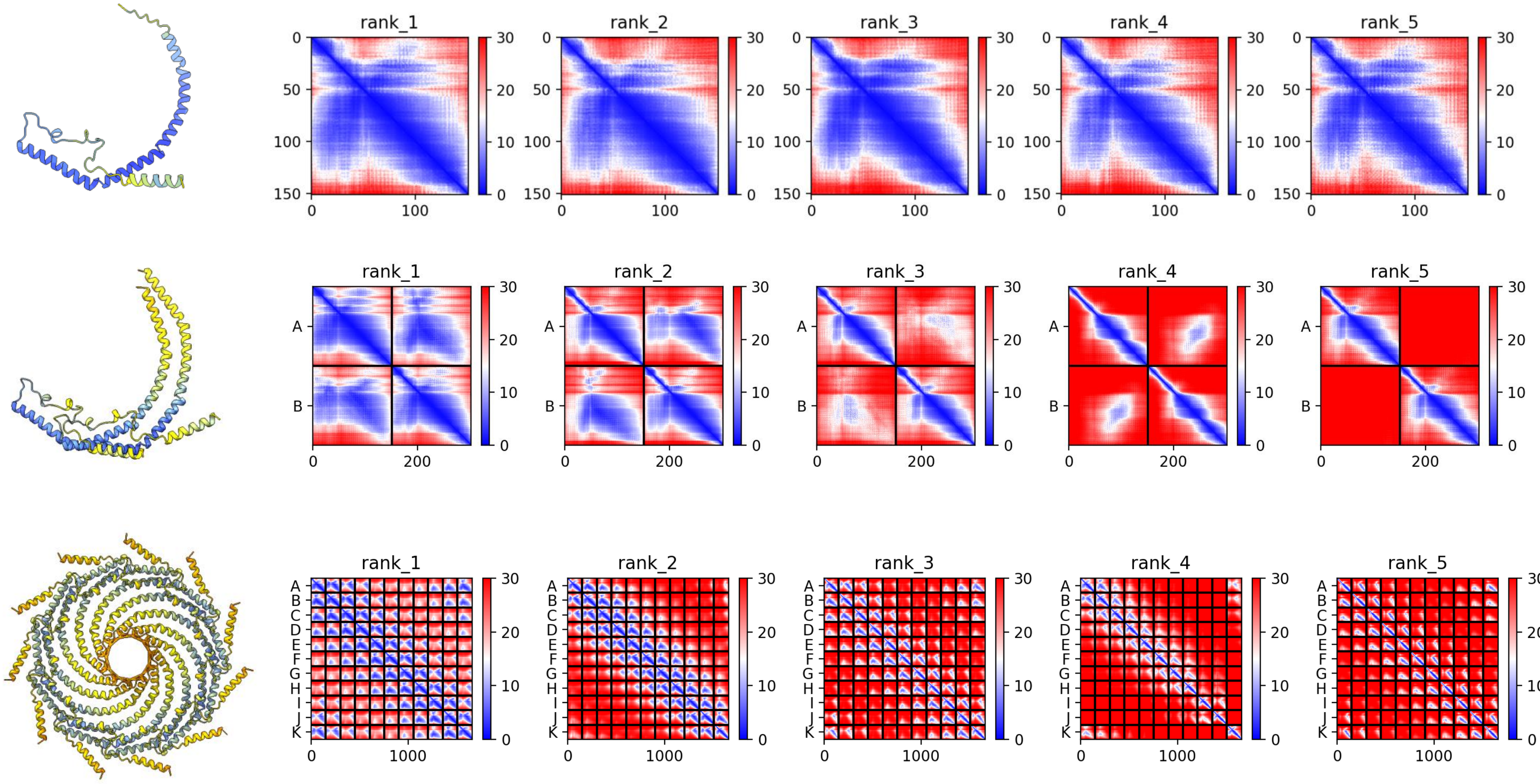

### Q03135 (CAV1) *H. sapiens*

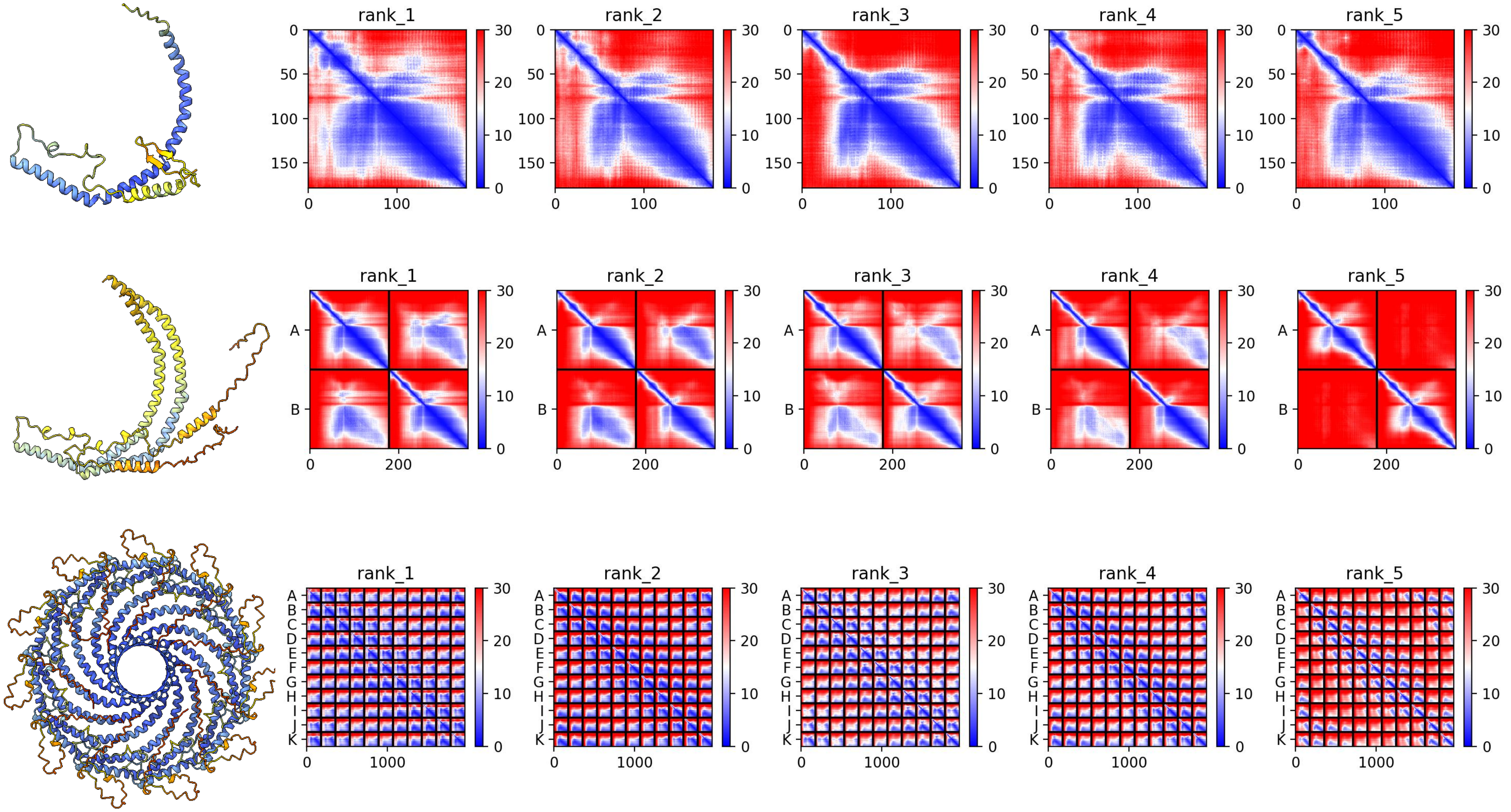

### A0A1S3HK30 *L. unguis*

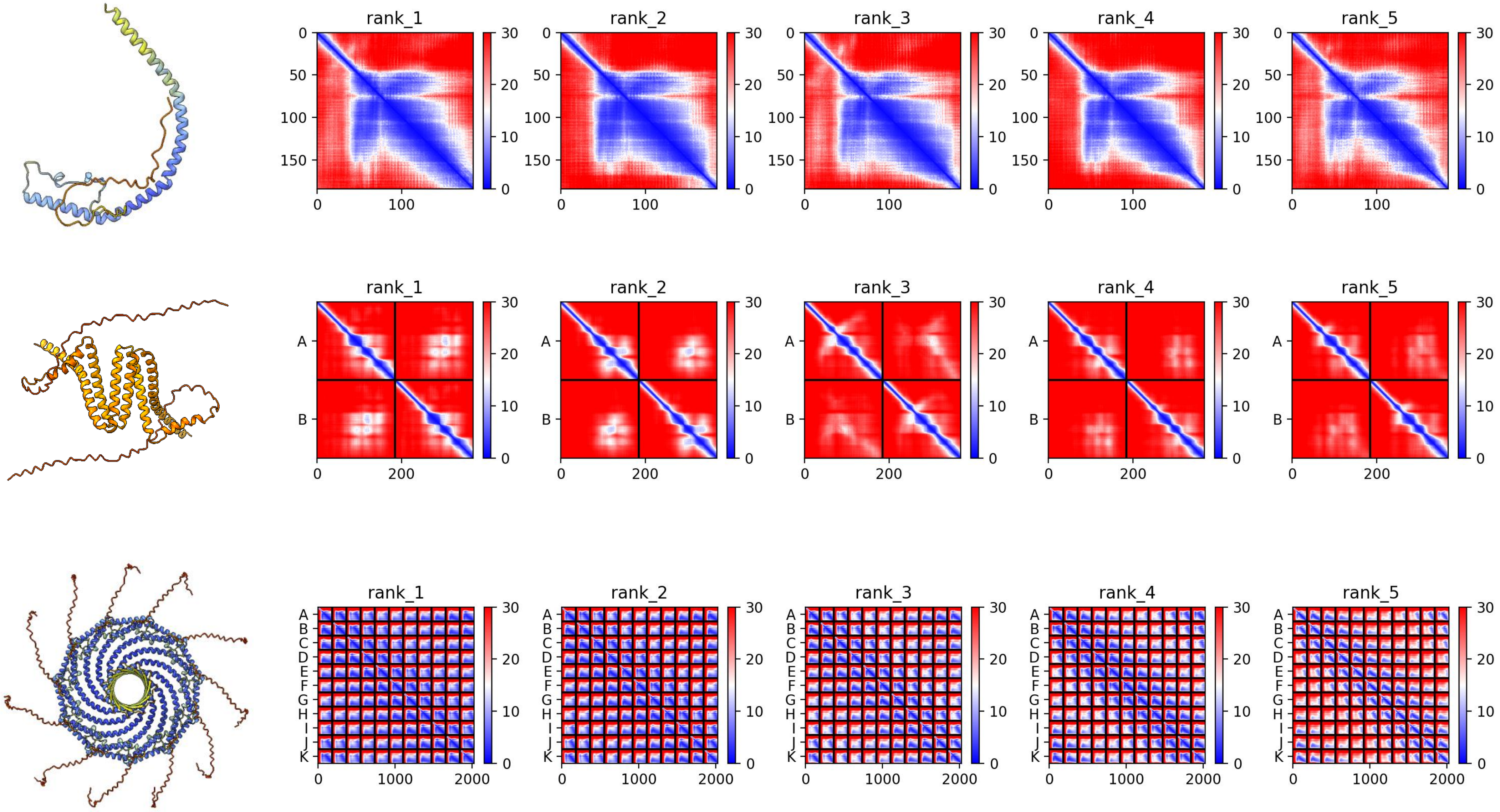

### A0A1S3JHP2 *L. unguis*

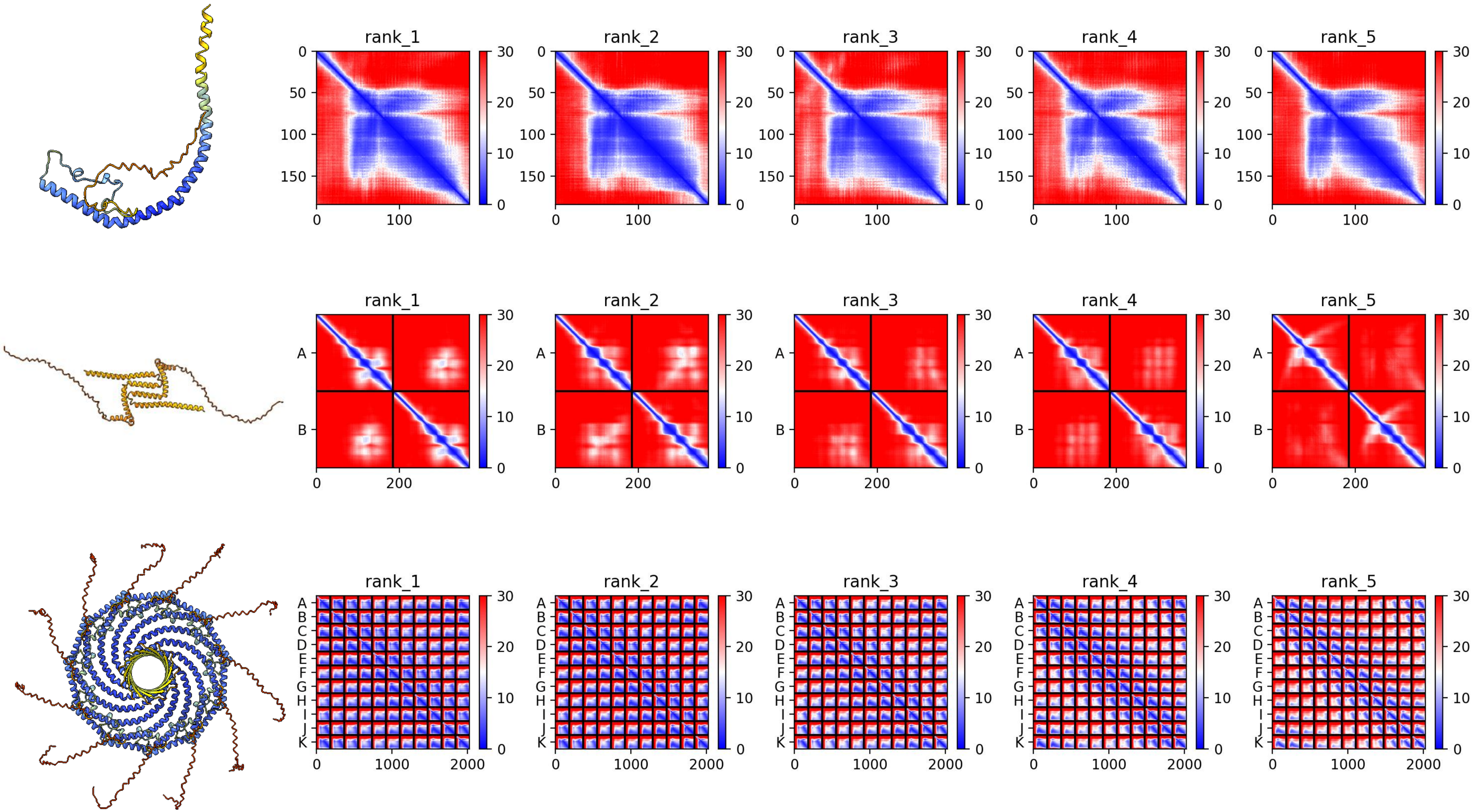

### A0A1S3IQQ3 *L. unguis*

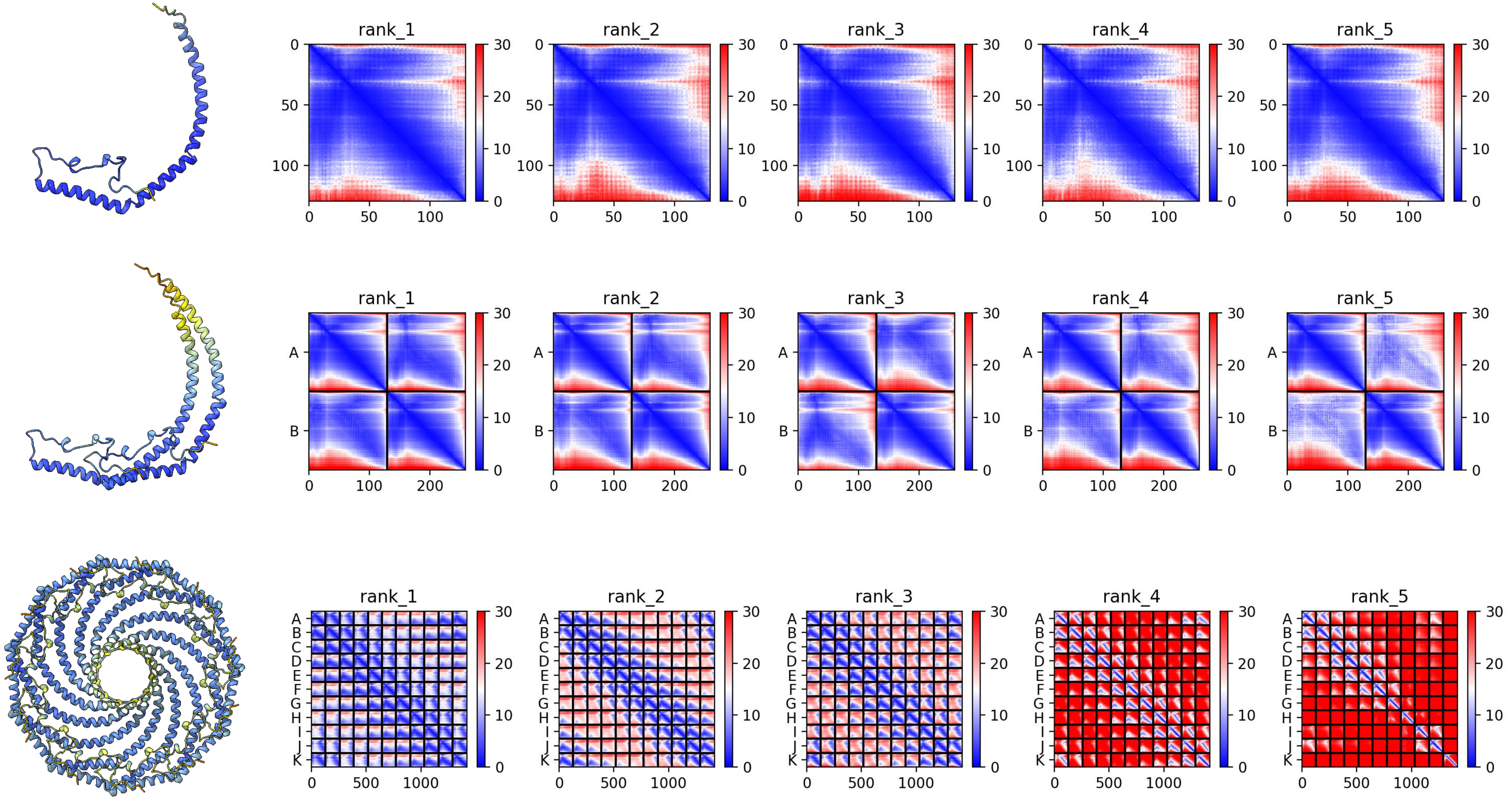

### A0A1S3IG00 *L. unguis*

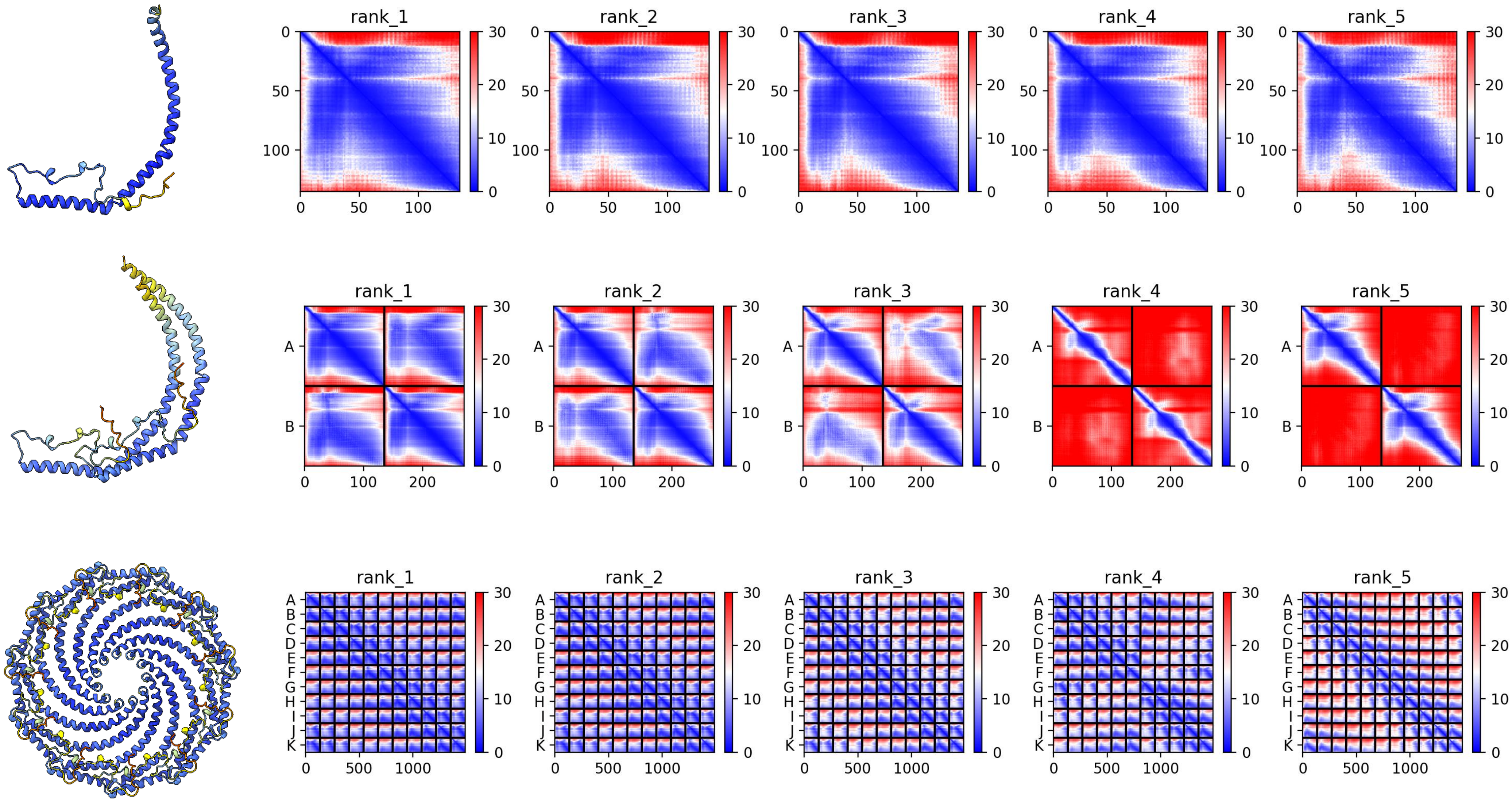

### A0A1S3IG42 *L. unguis*

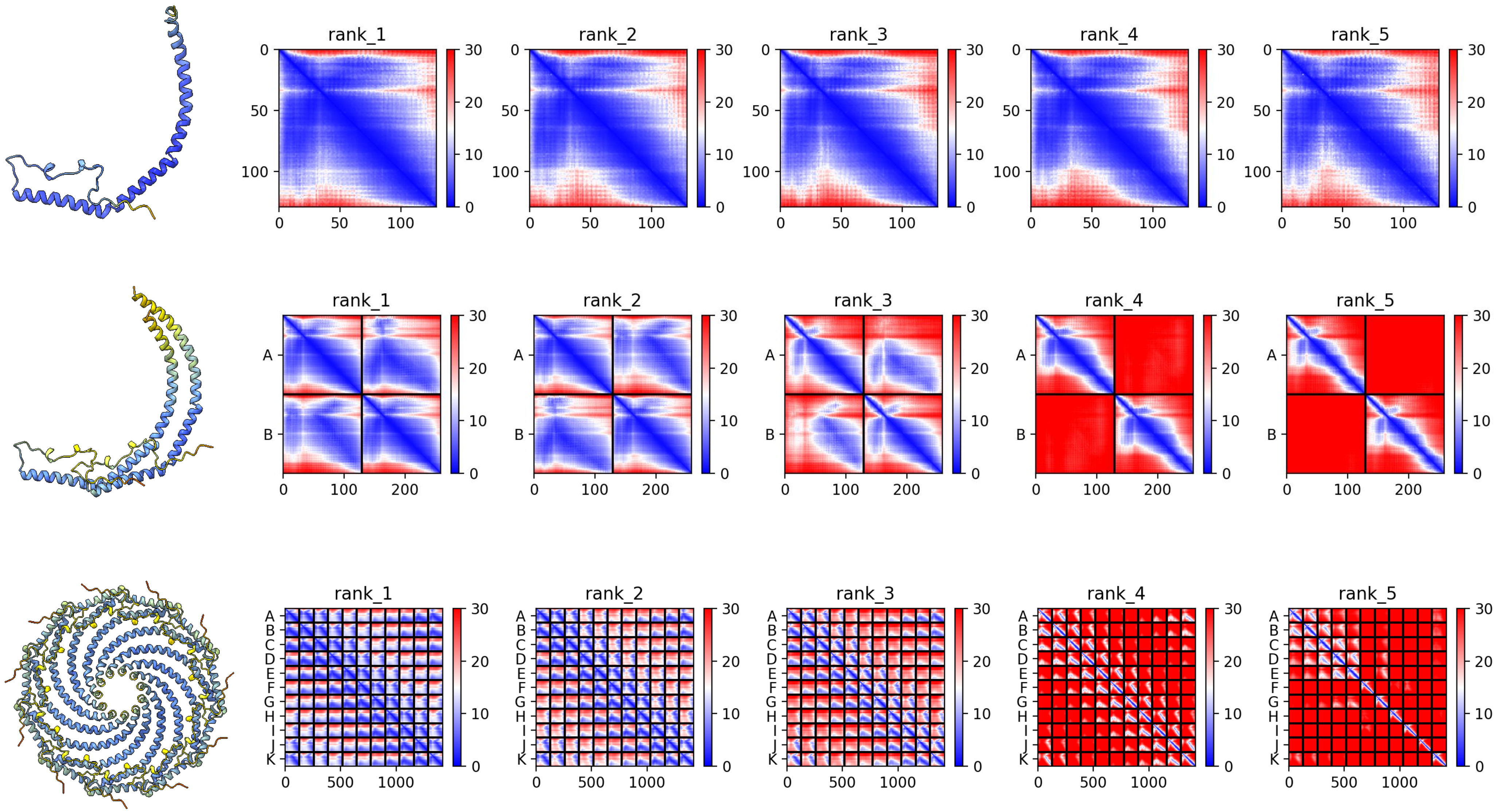

### A0A1S3IH41 *L. unguis*

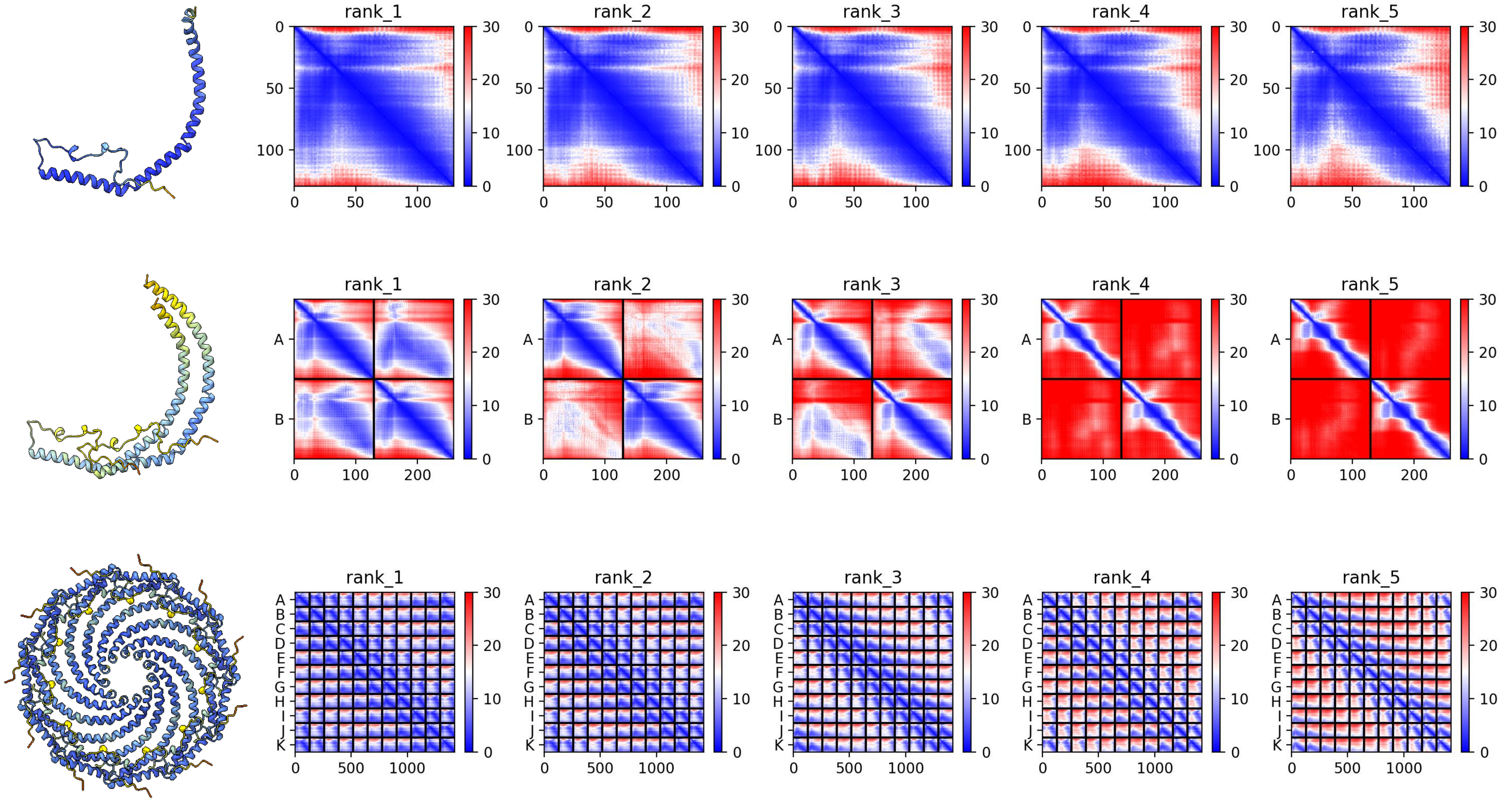

### XP_014678552.1 *P. caudatus*

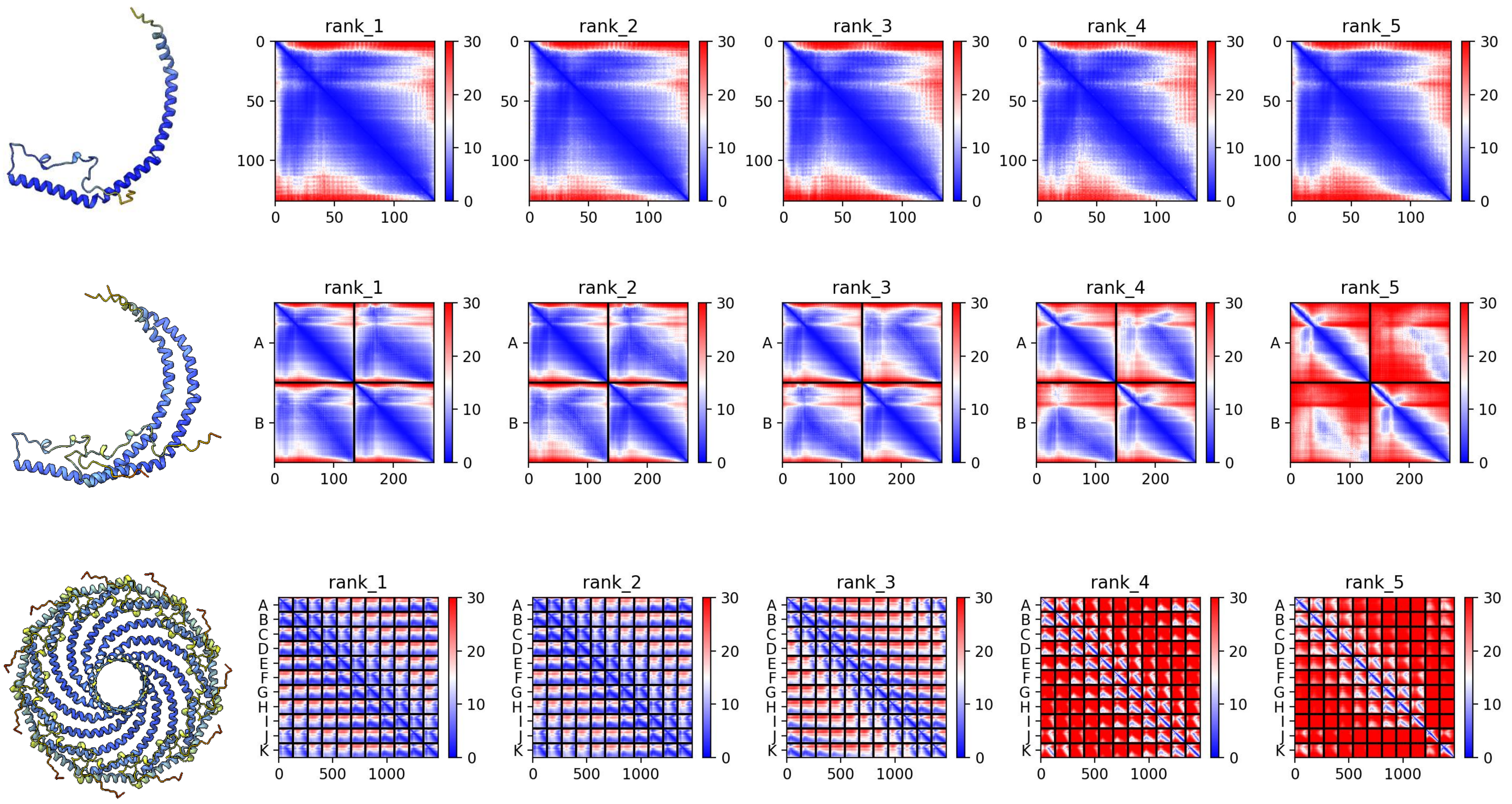

### A0A7M7R2L2 *A. mellifera*

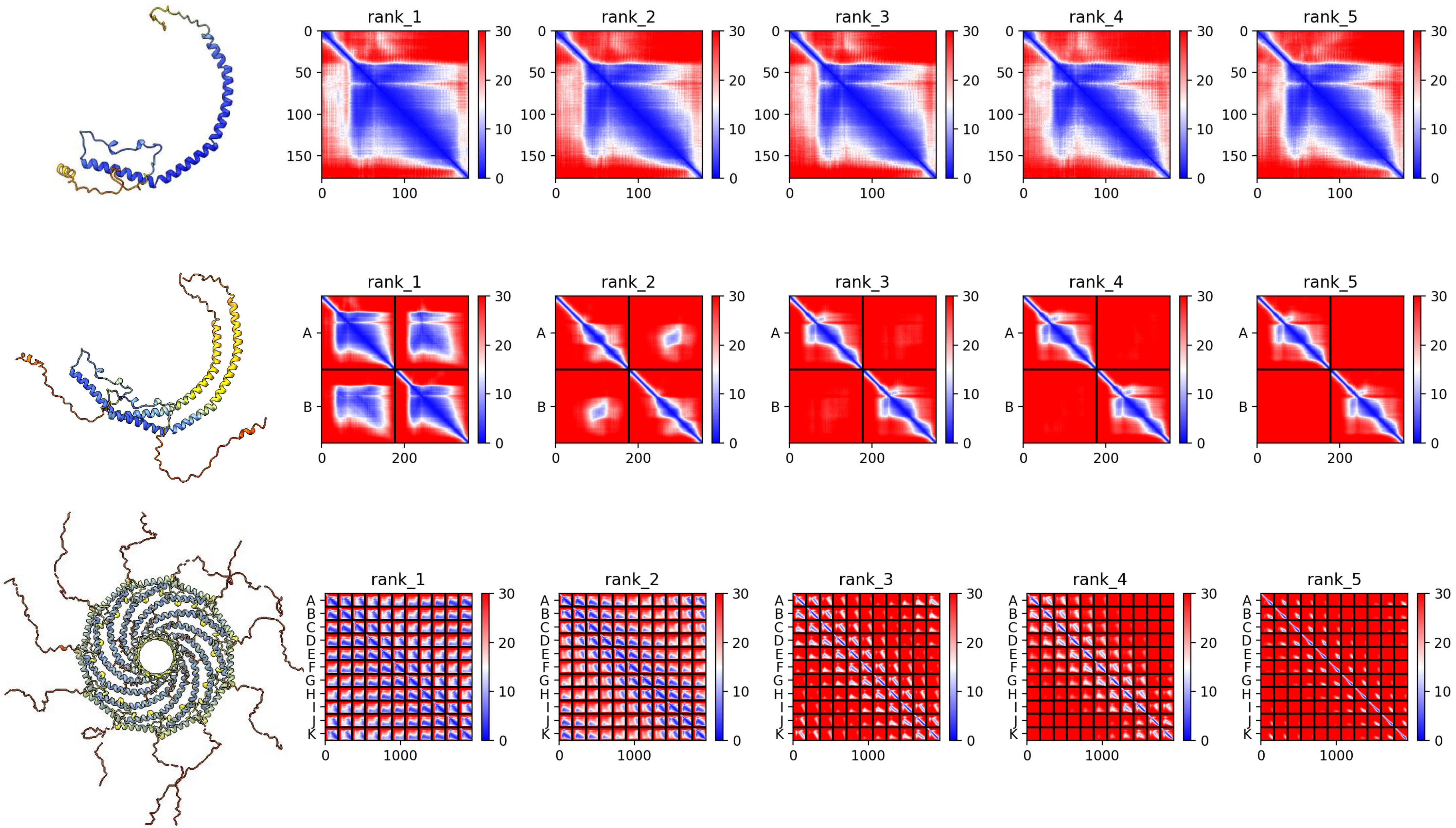

### A0A3M7SAF6 *B. plicatilis*

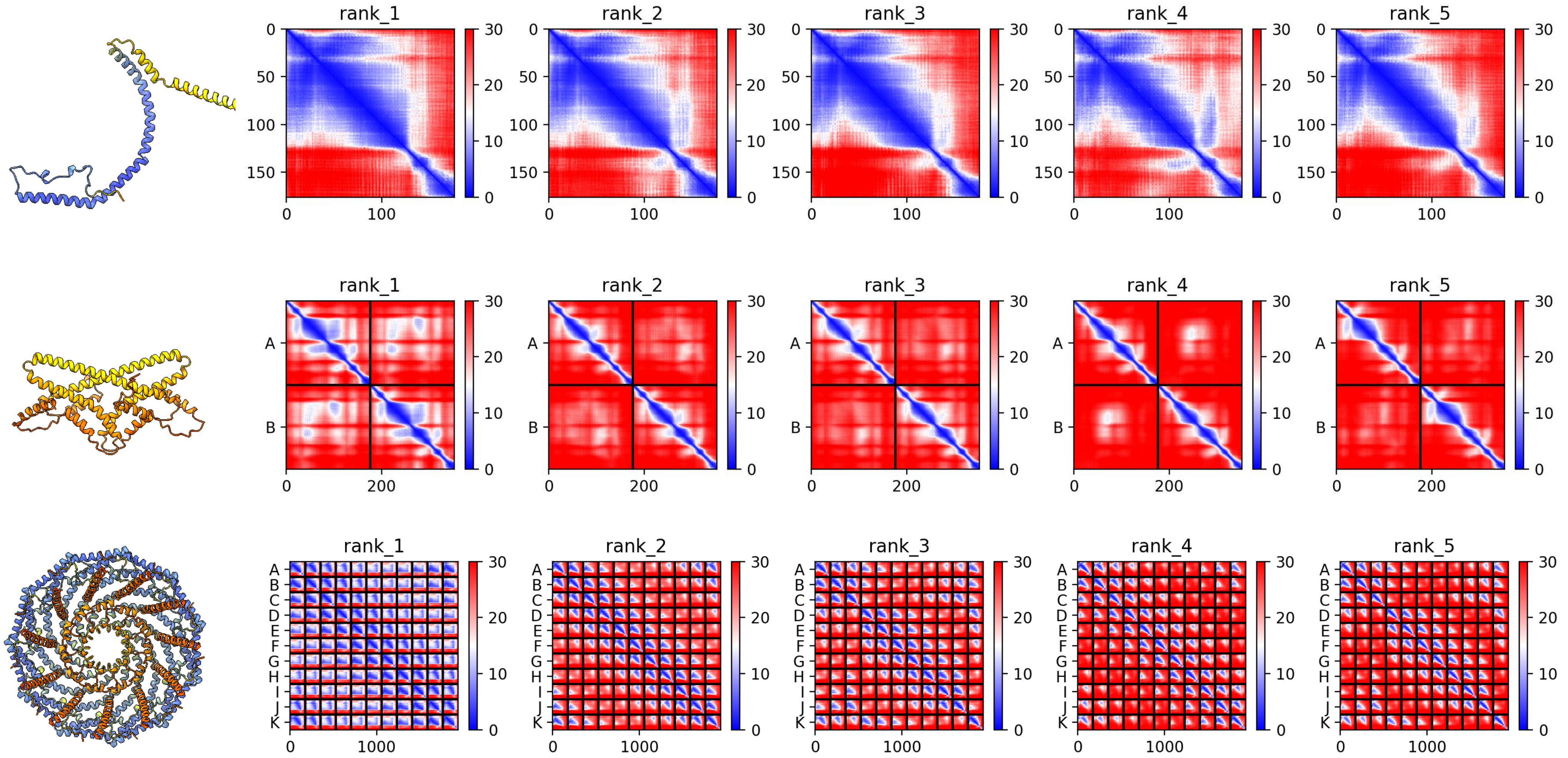

### A0A1I8ILQ8 *M. lignano*

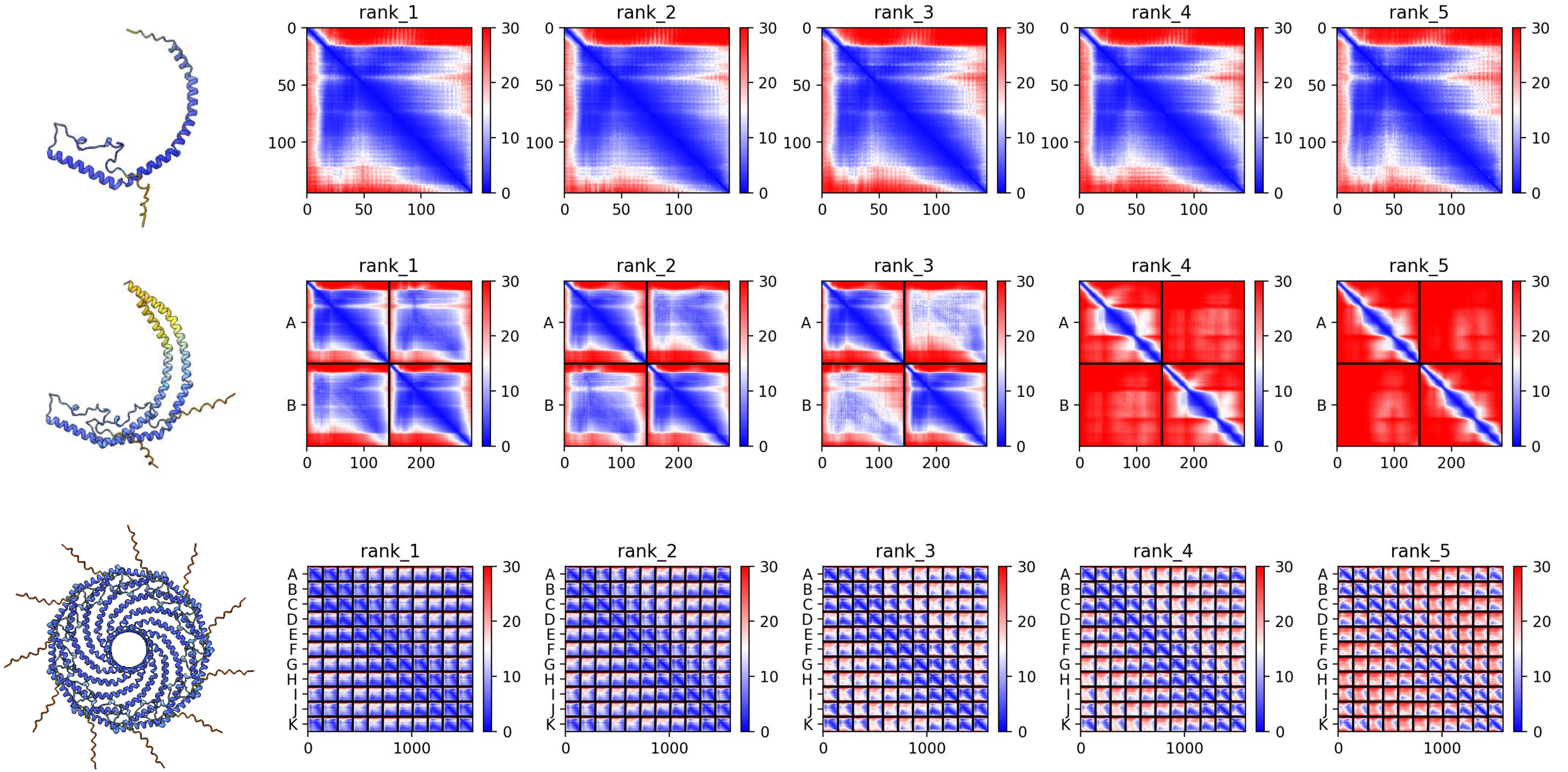

### A0A1I8HHY8 *M. lignano*

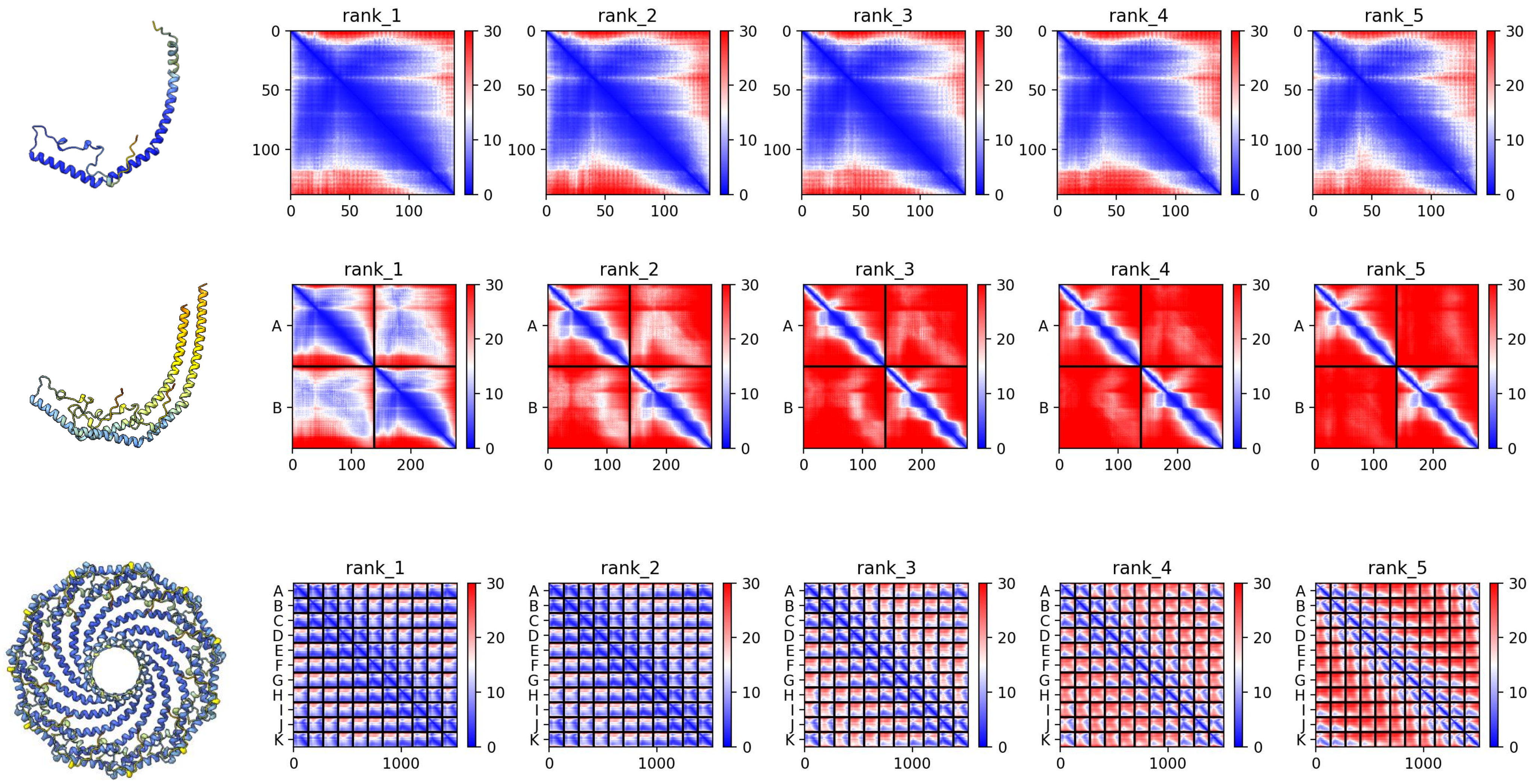

### A0A267DBT9 *M. lignano*

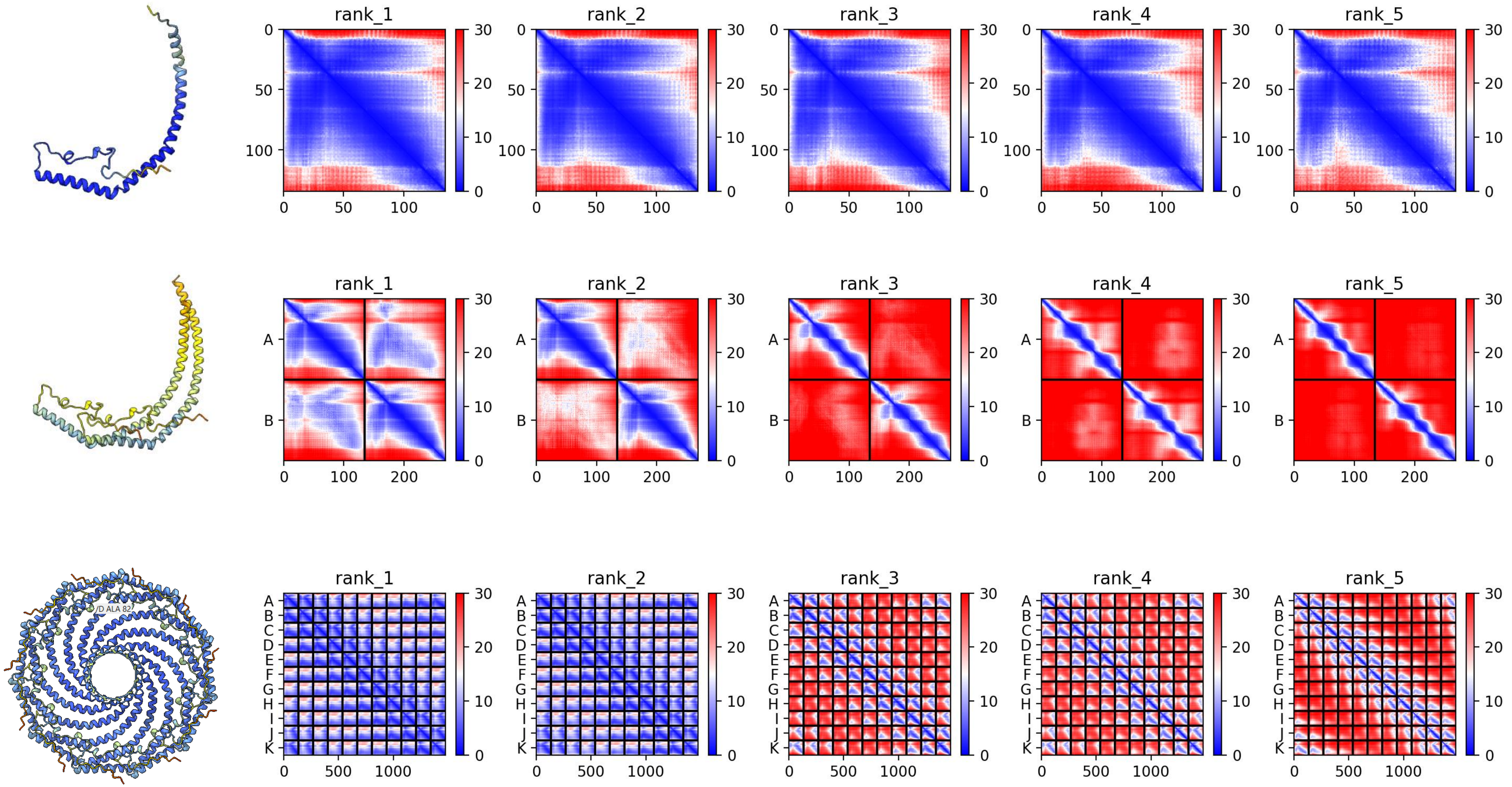

### A0A267F2S4 *M. lignano*

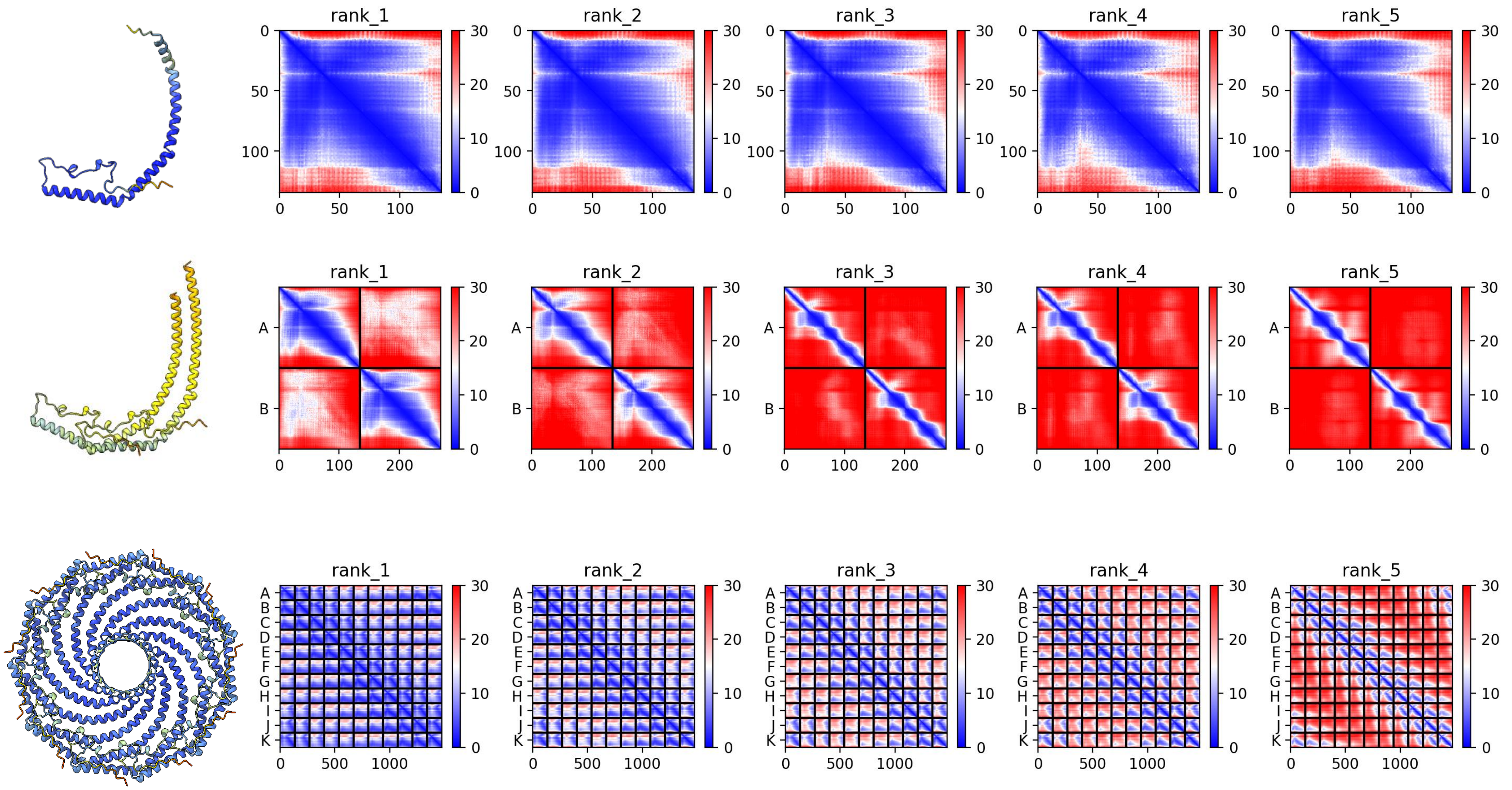

### A0A3M7SB10 *B. plicatilis*

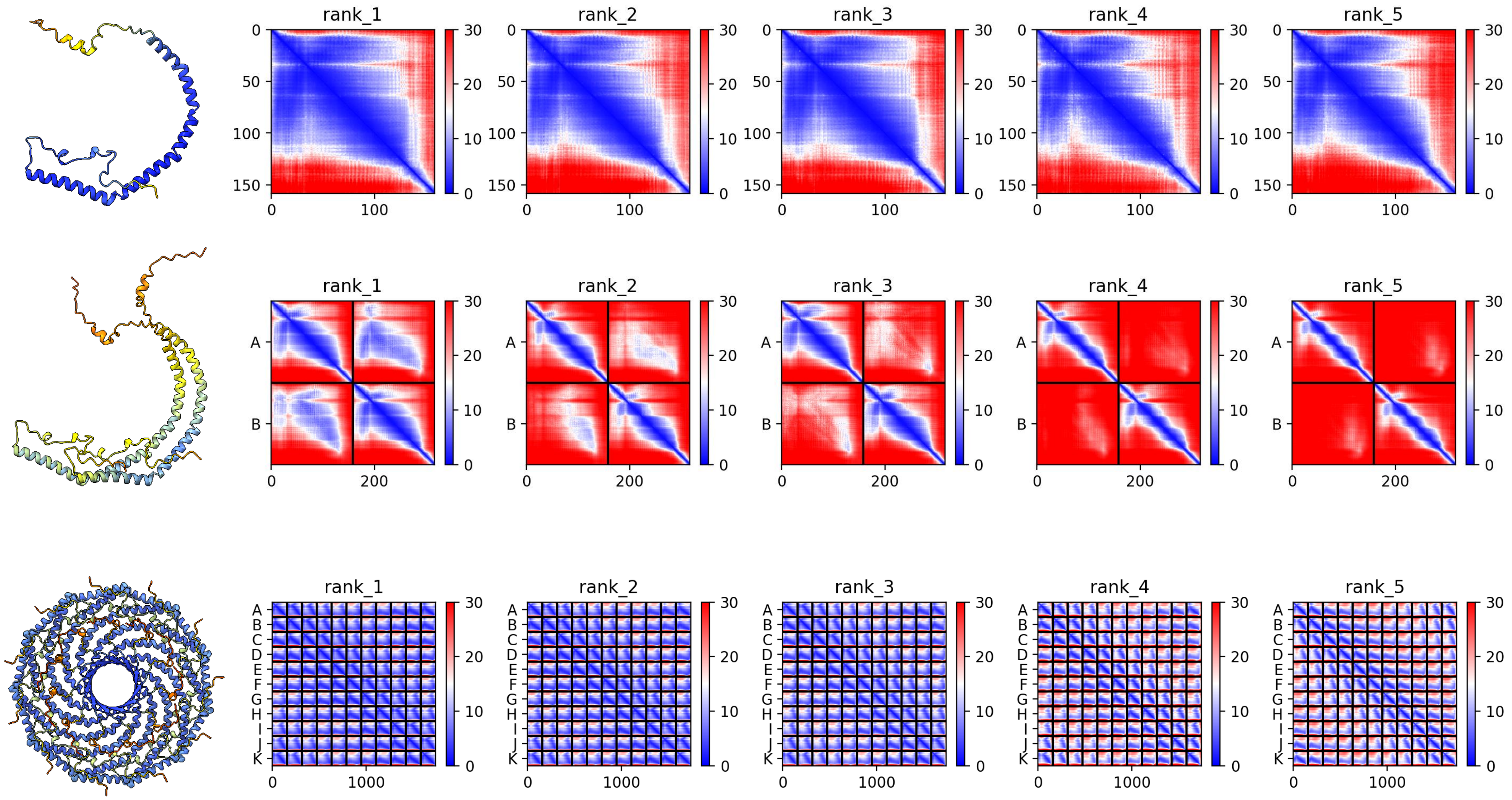

### A0A3M7Q0Q1 *B. plicatilis*

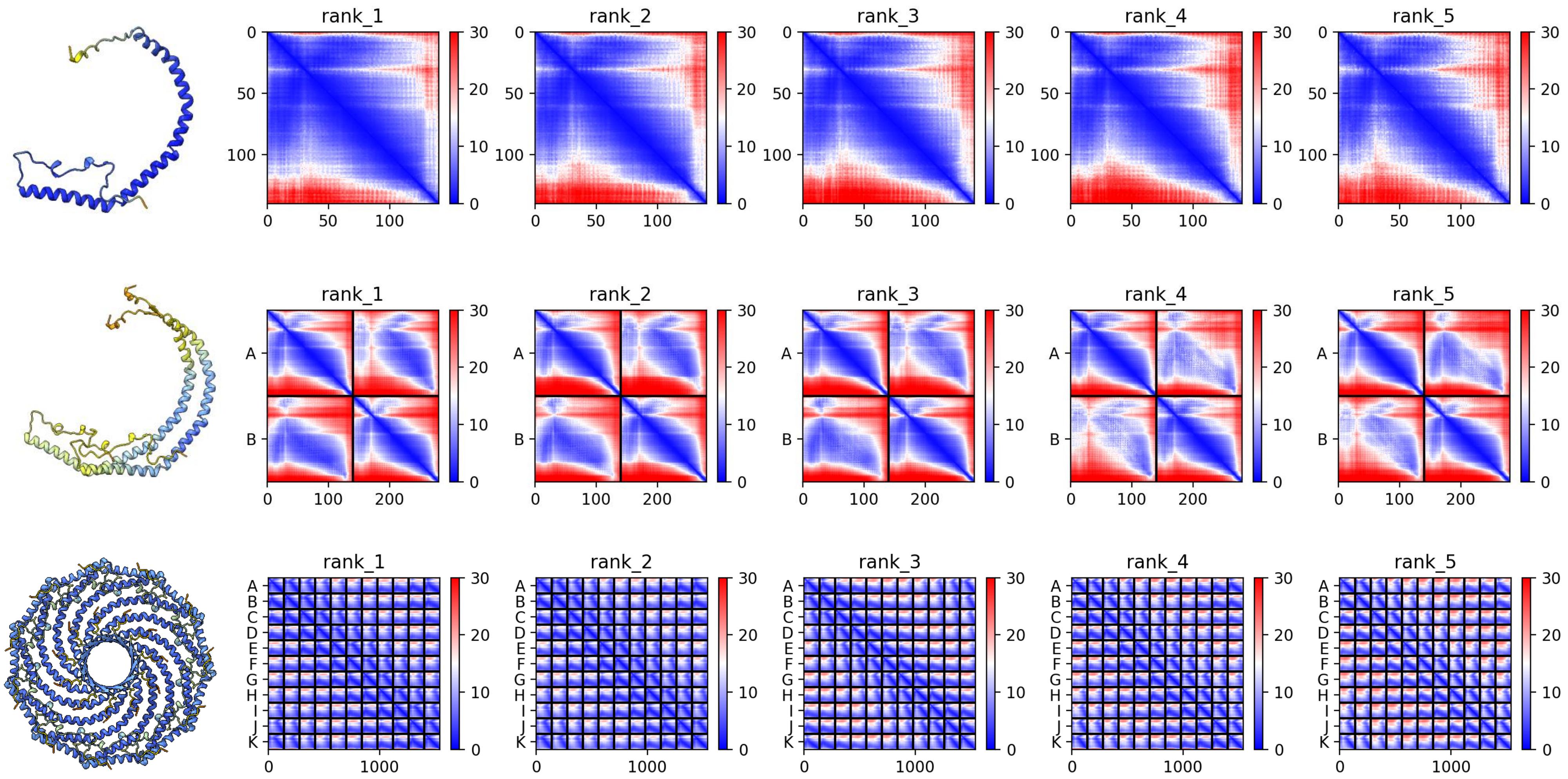

### A0A3M7RD35 *B. plicatilis*

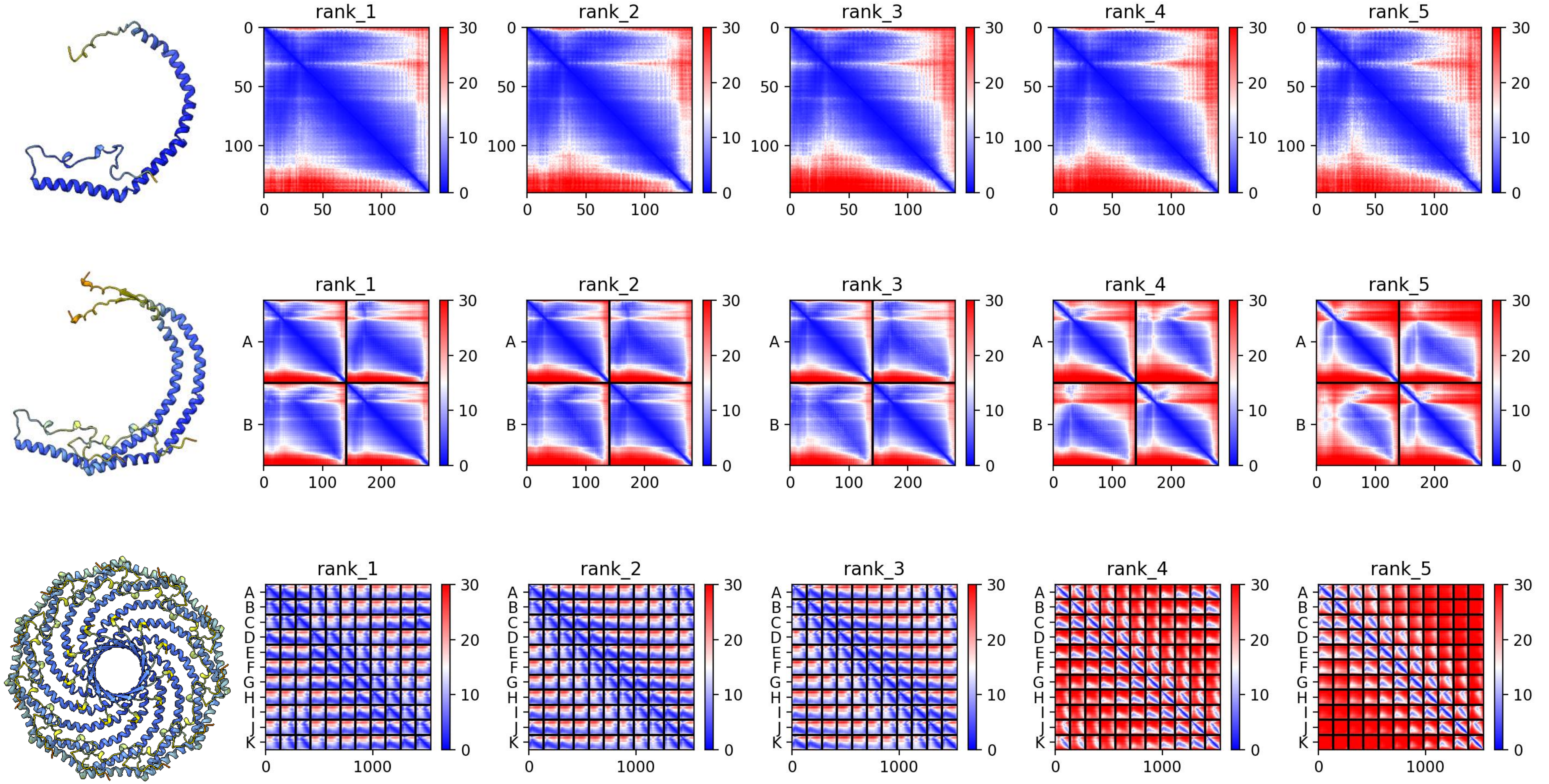

### A0A3M7RCJ9 *B. plicatilis*

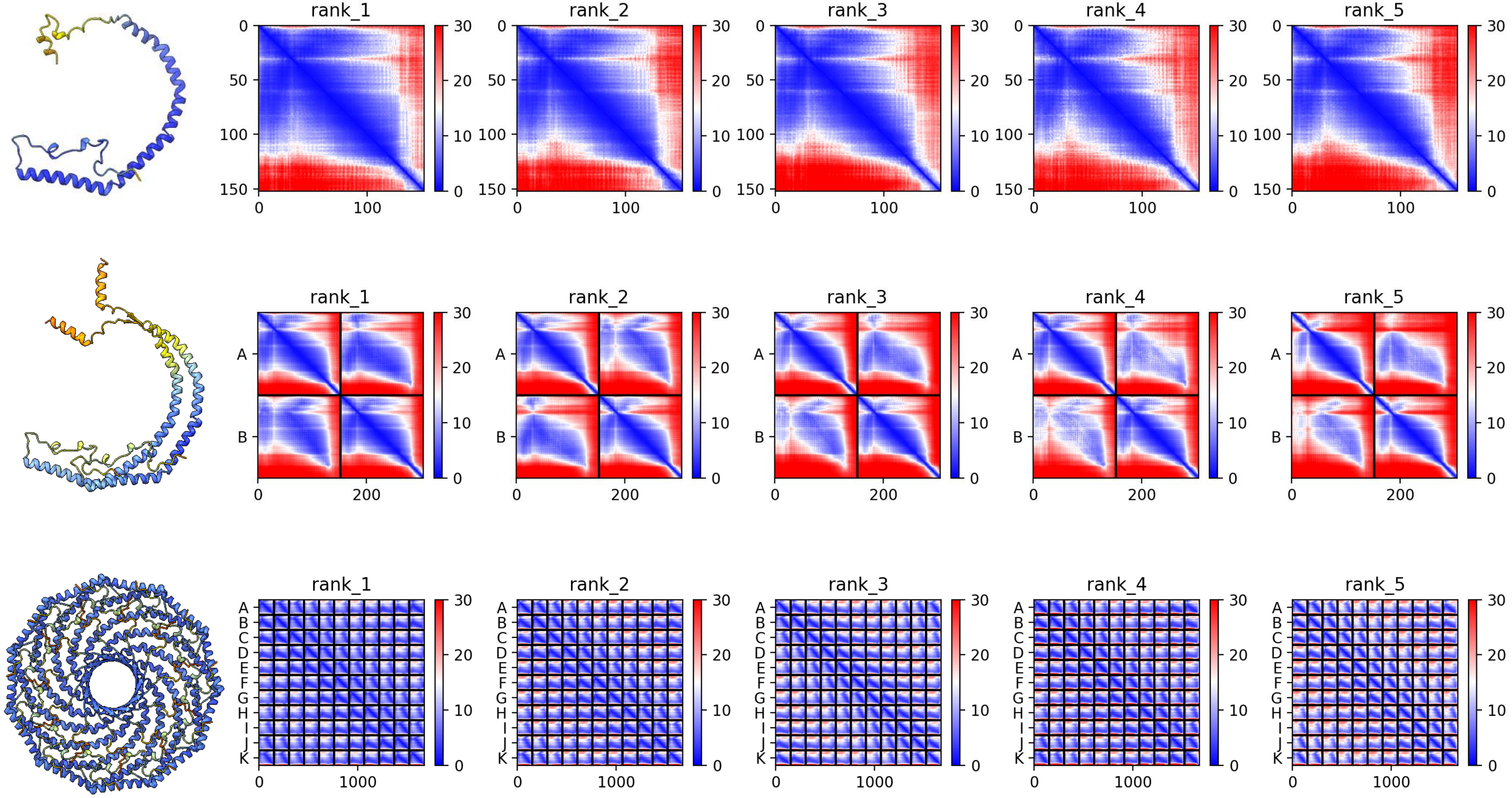

### A0A3M7SAK0 *B. plicatilis*

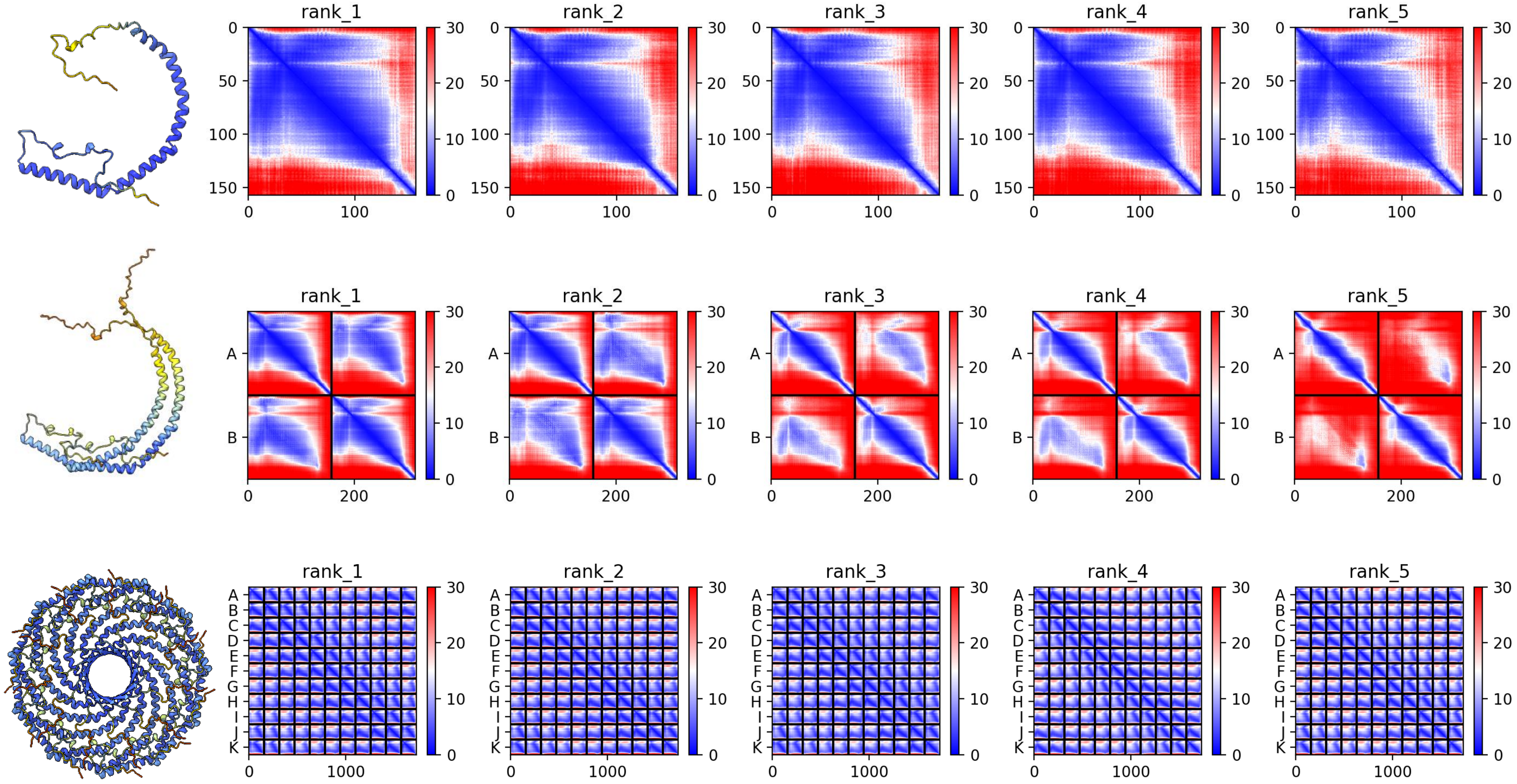

### A0A1I8H2T4 *M. lignano*

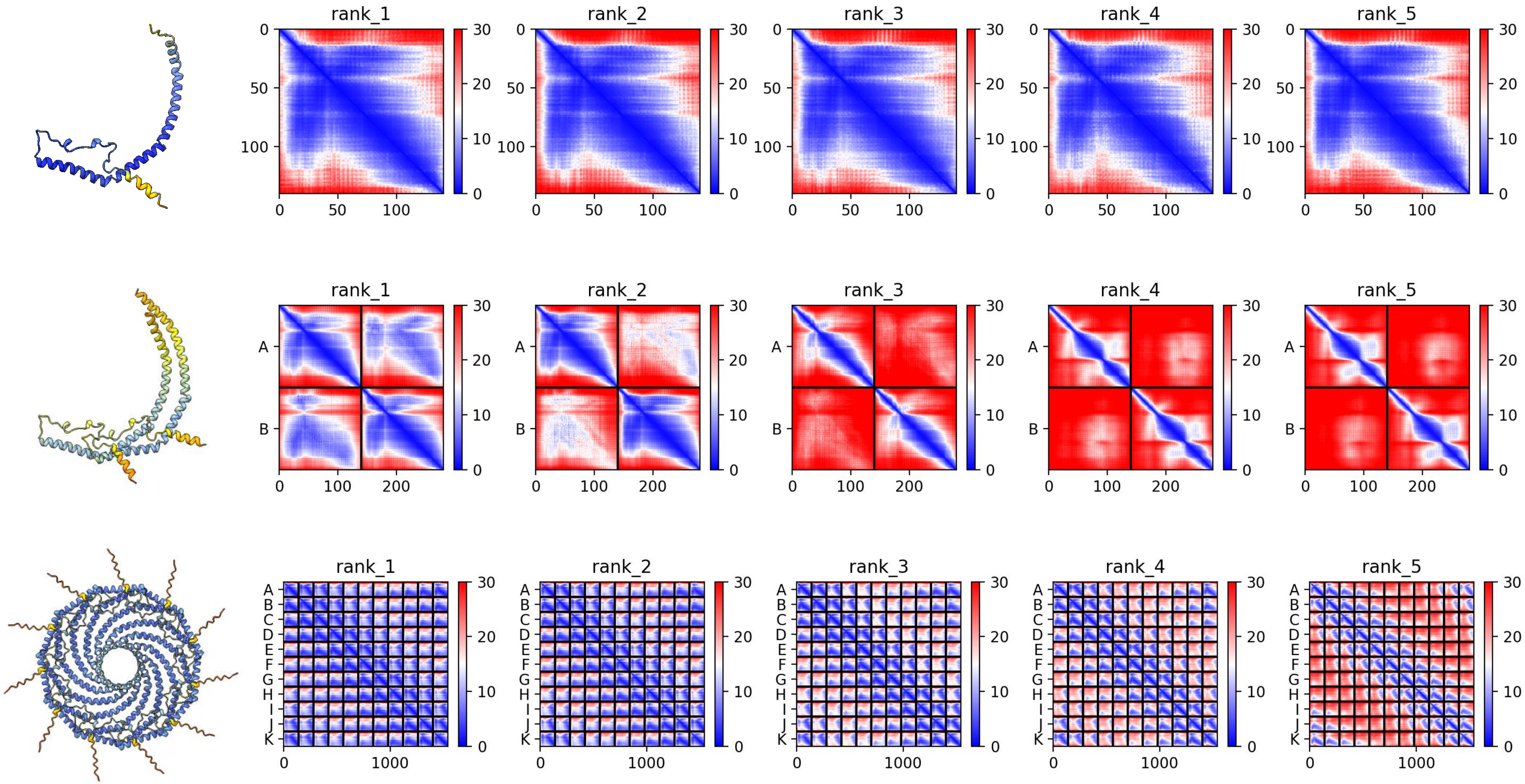

### A0A1I8H327 *M. lignano*

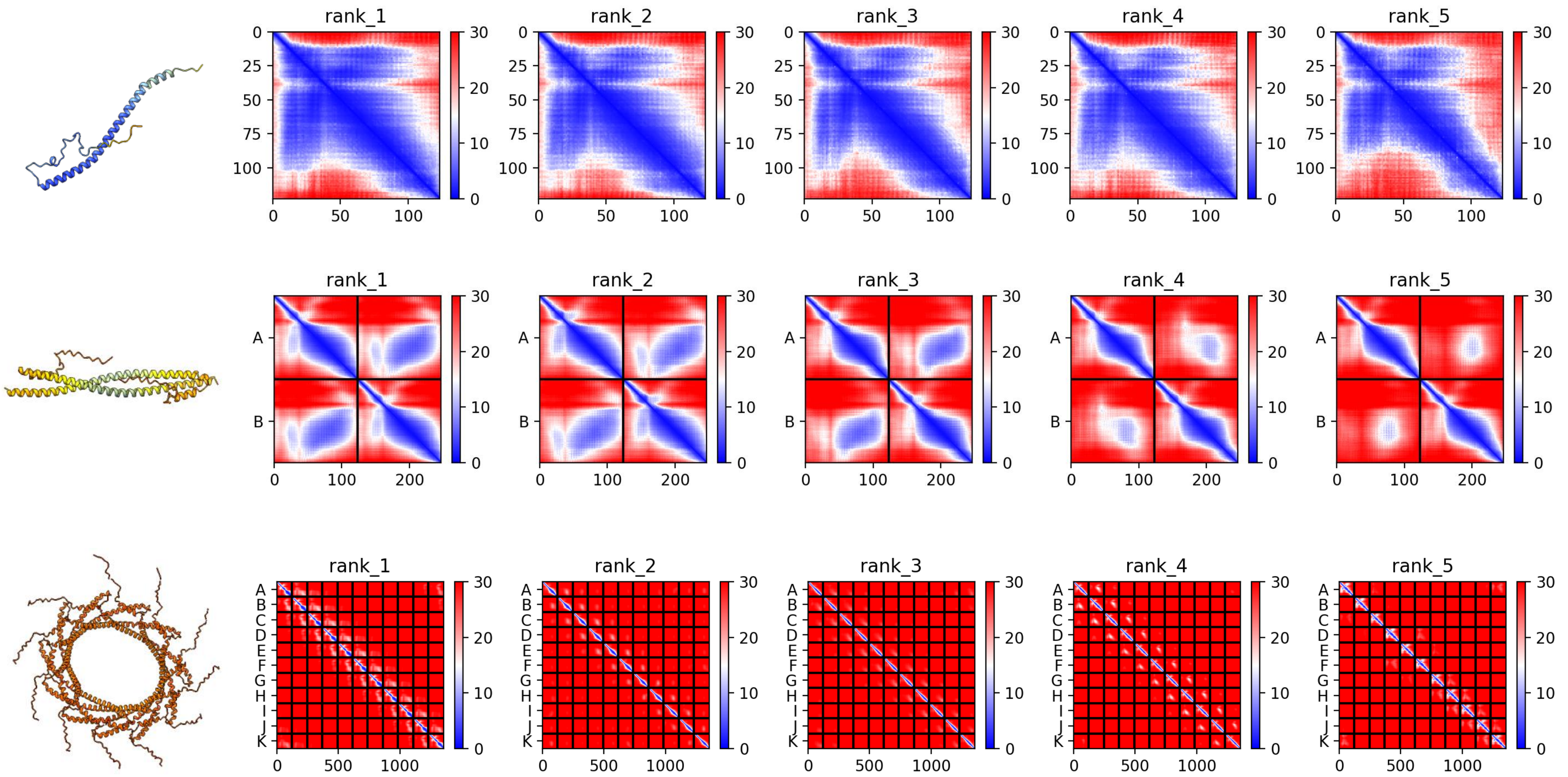

### A0A267DS21 *M. lignano*

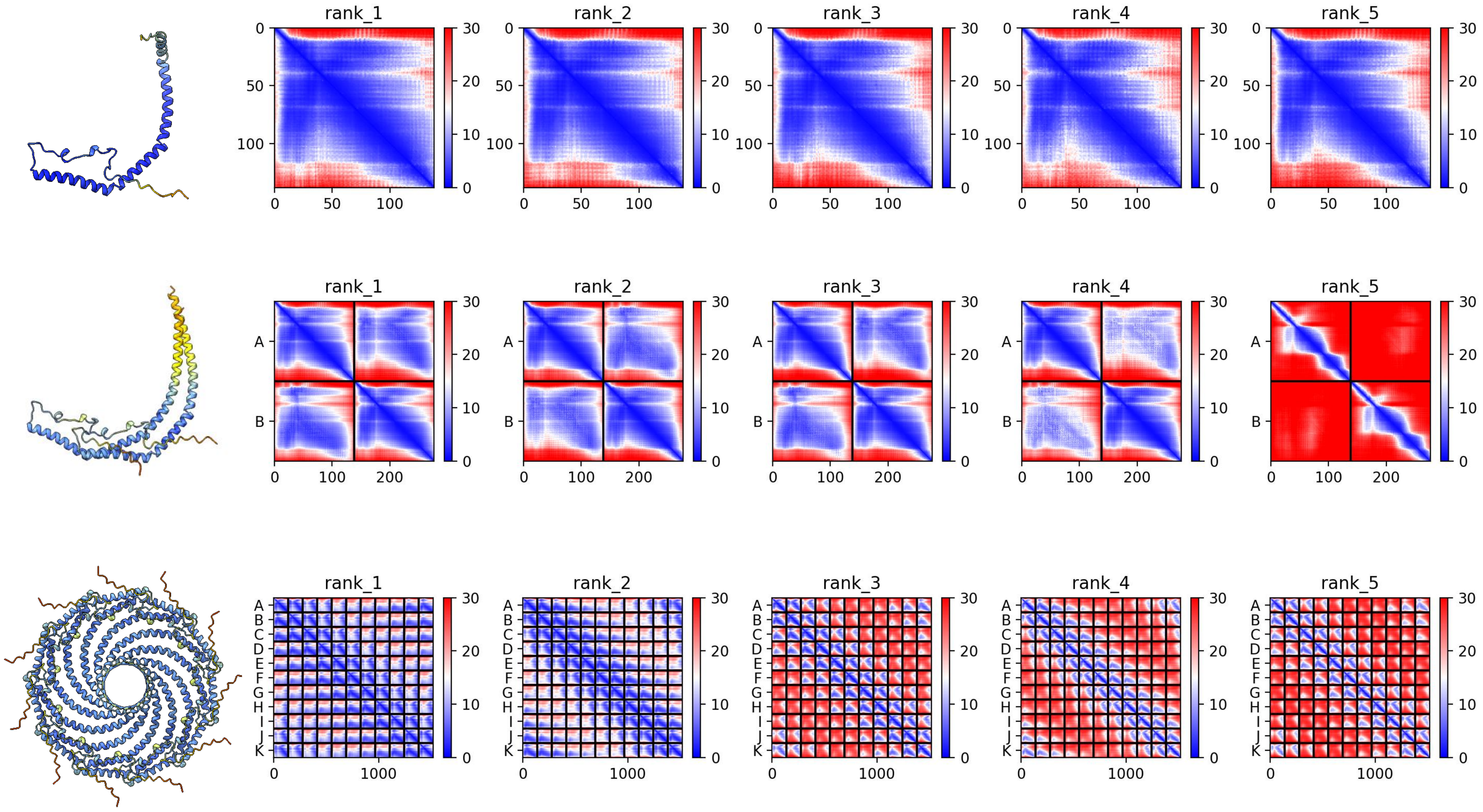

### A0A267DU21 *M. lignano*

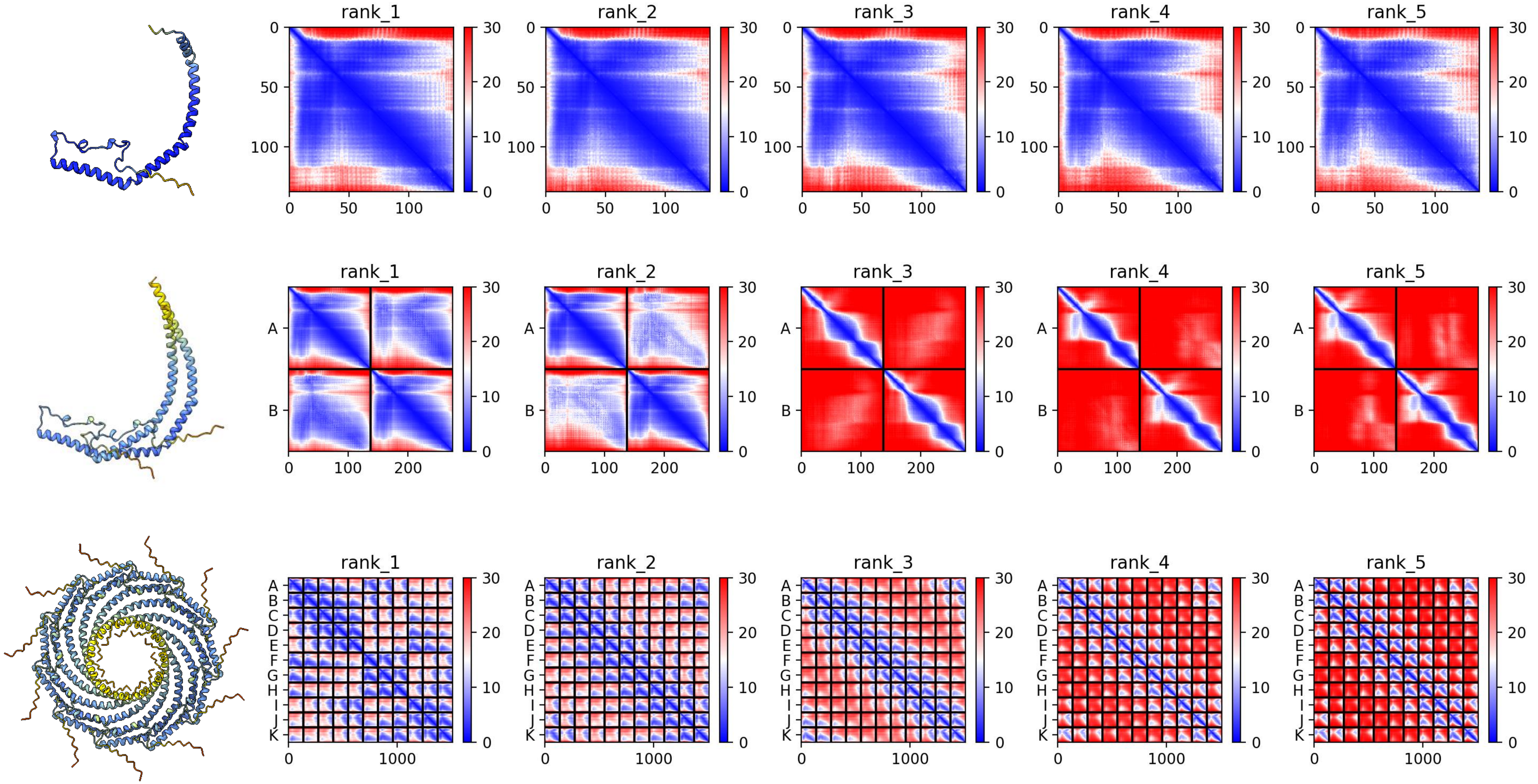

### A0A1I8J849 *M. lignano*

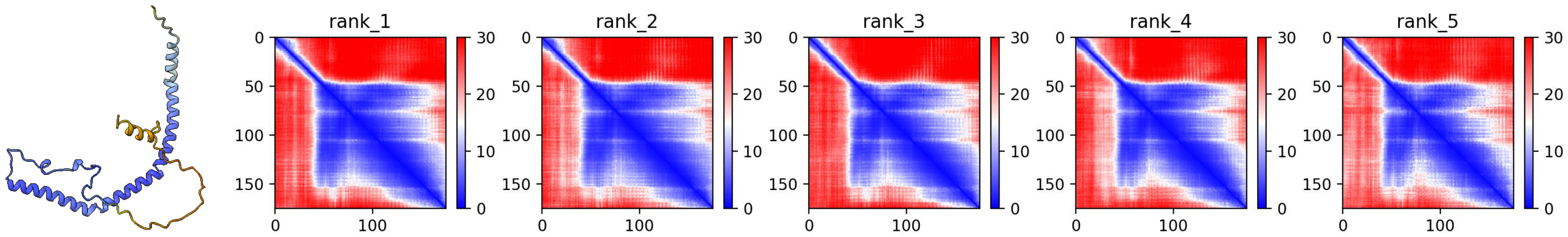


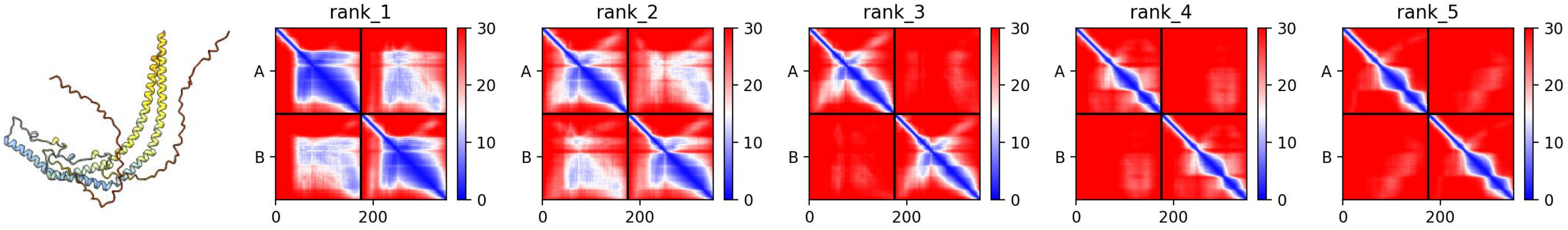


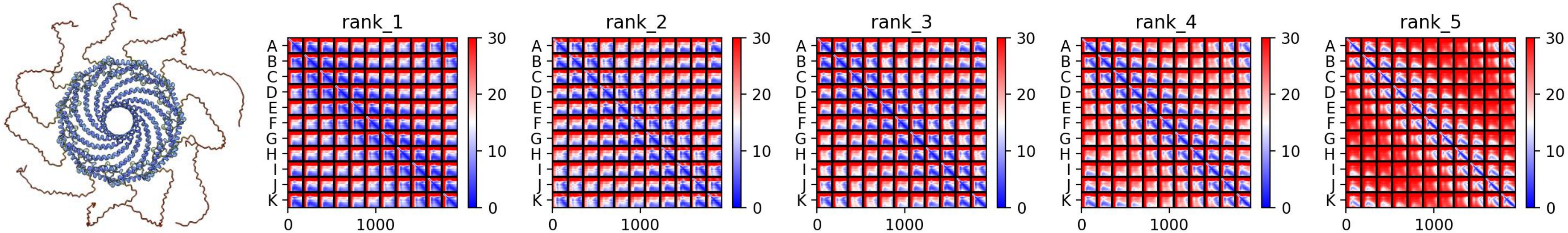

### A0A267FNG7 *M. lignano*

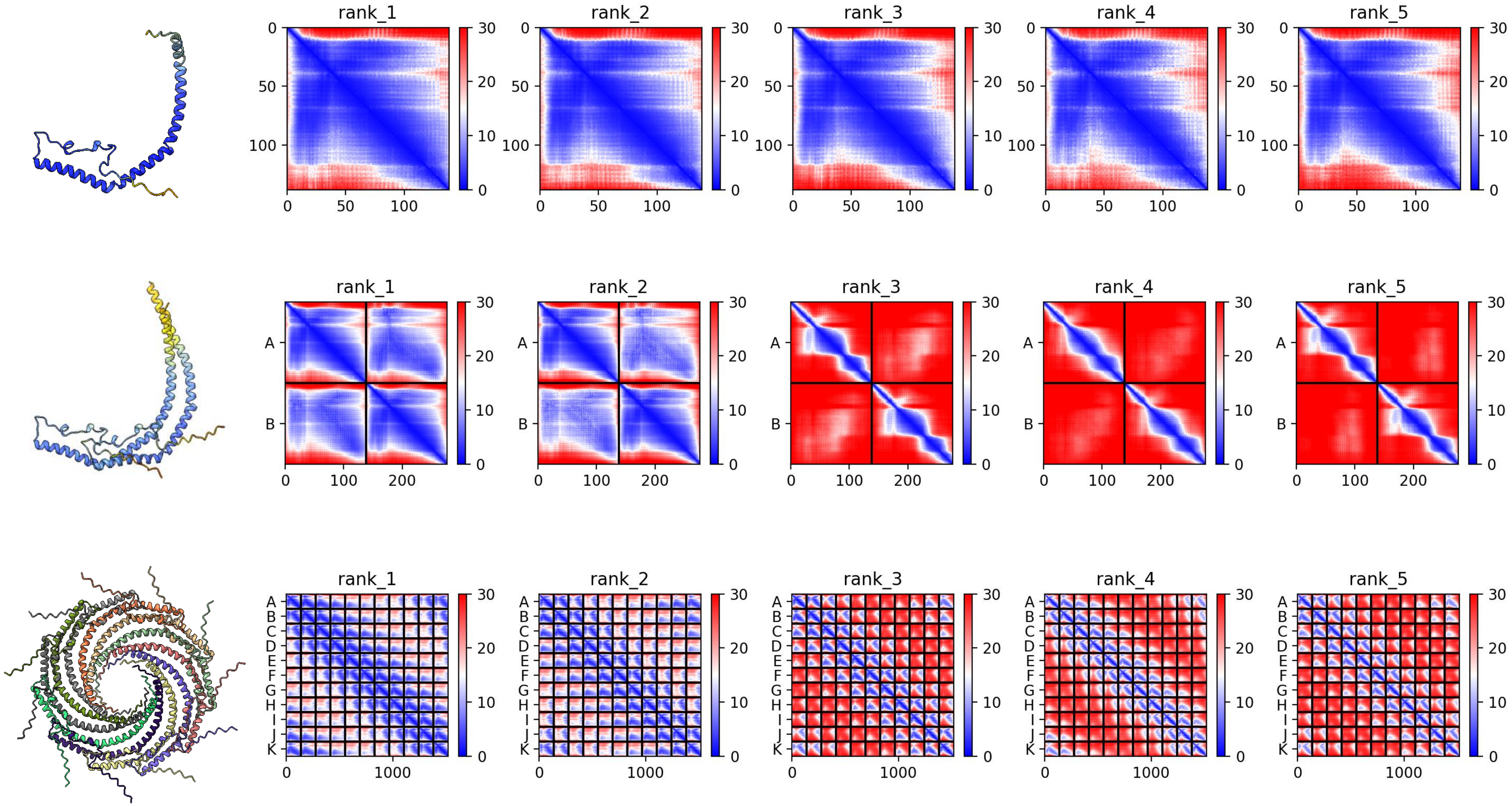

### A0A1I8J741 *M. lignano*

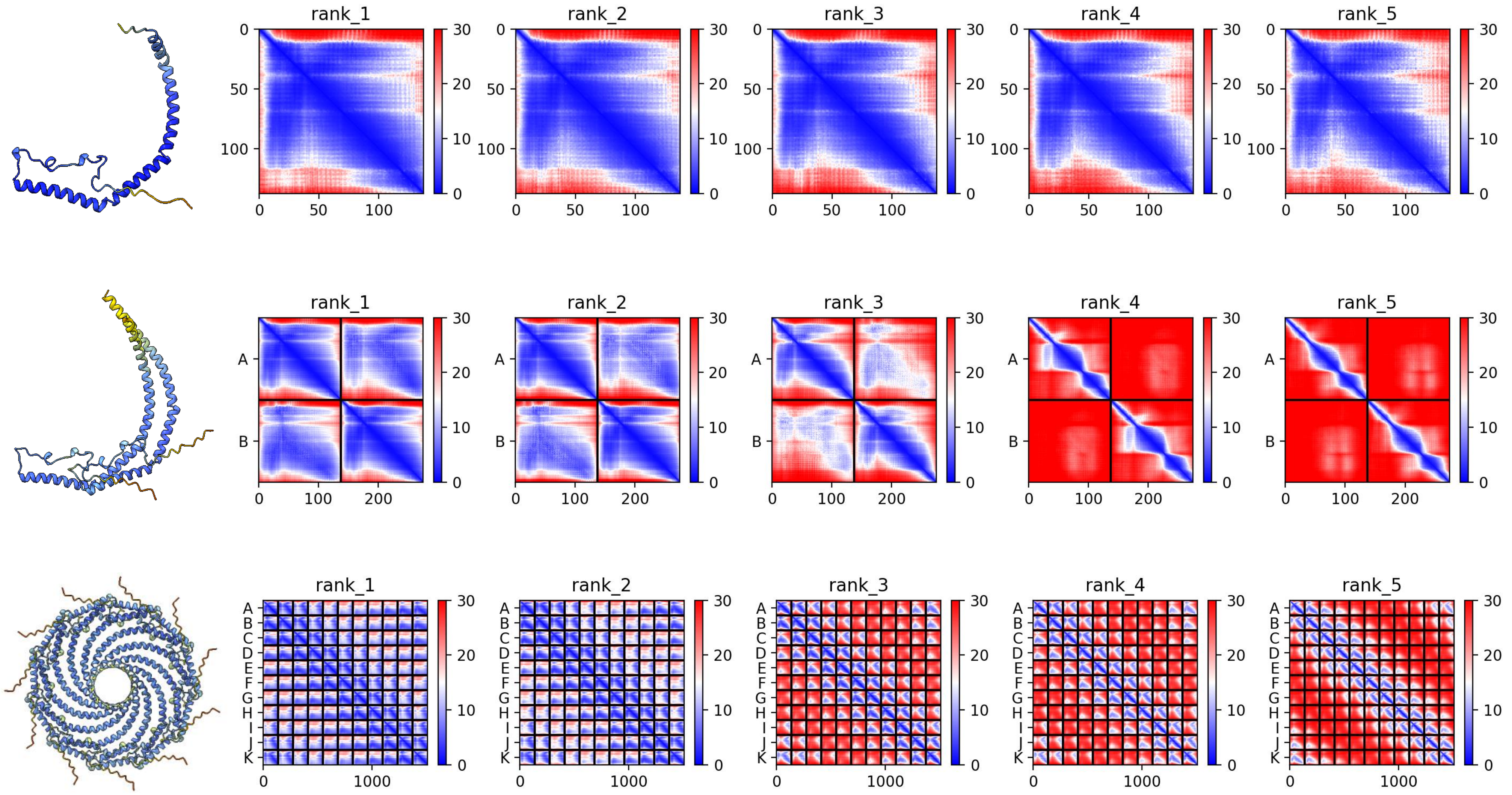

### A0A267DC90 *M. lignano*

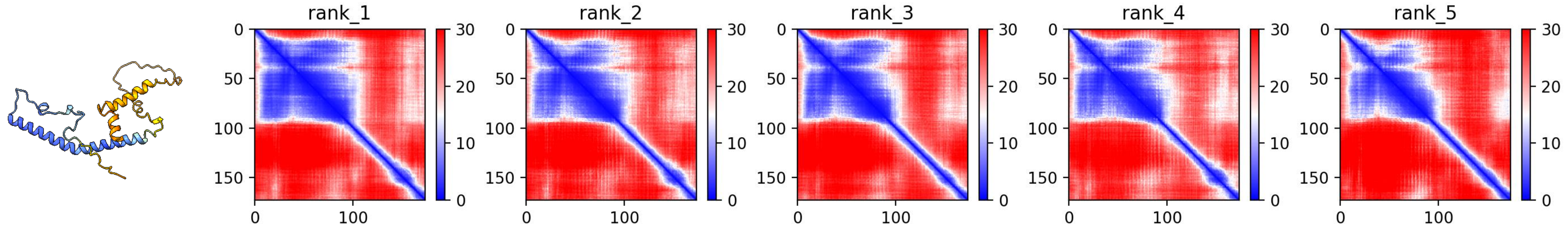


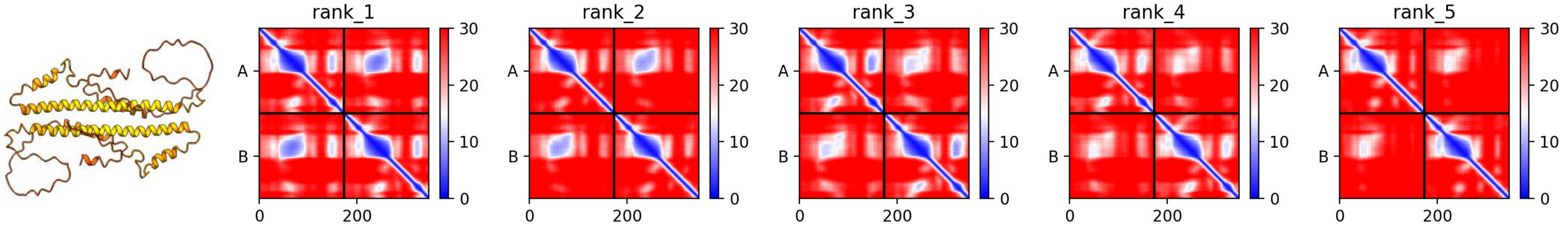


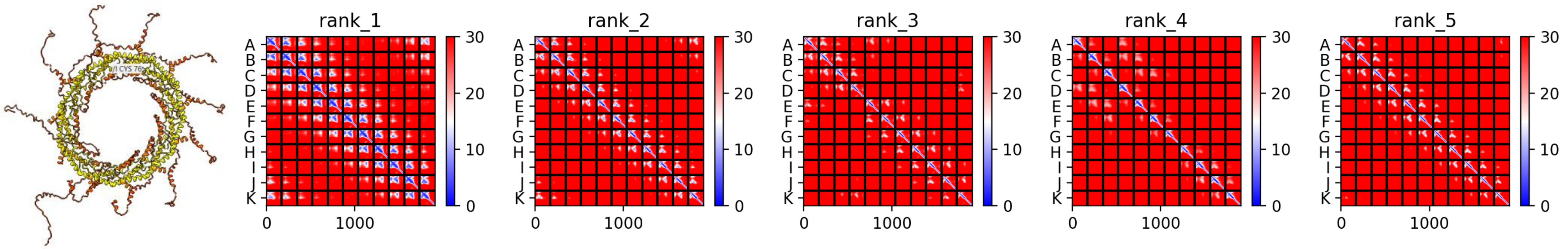

### A0A267F6W6 *M. lignano*

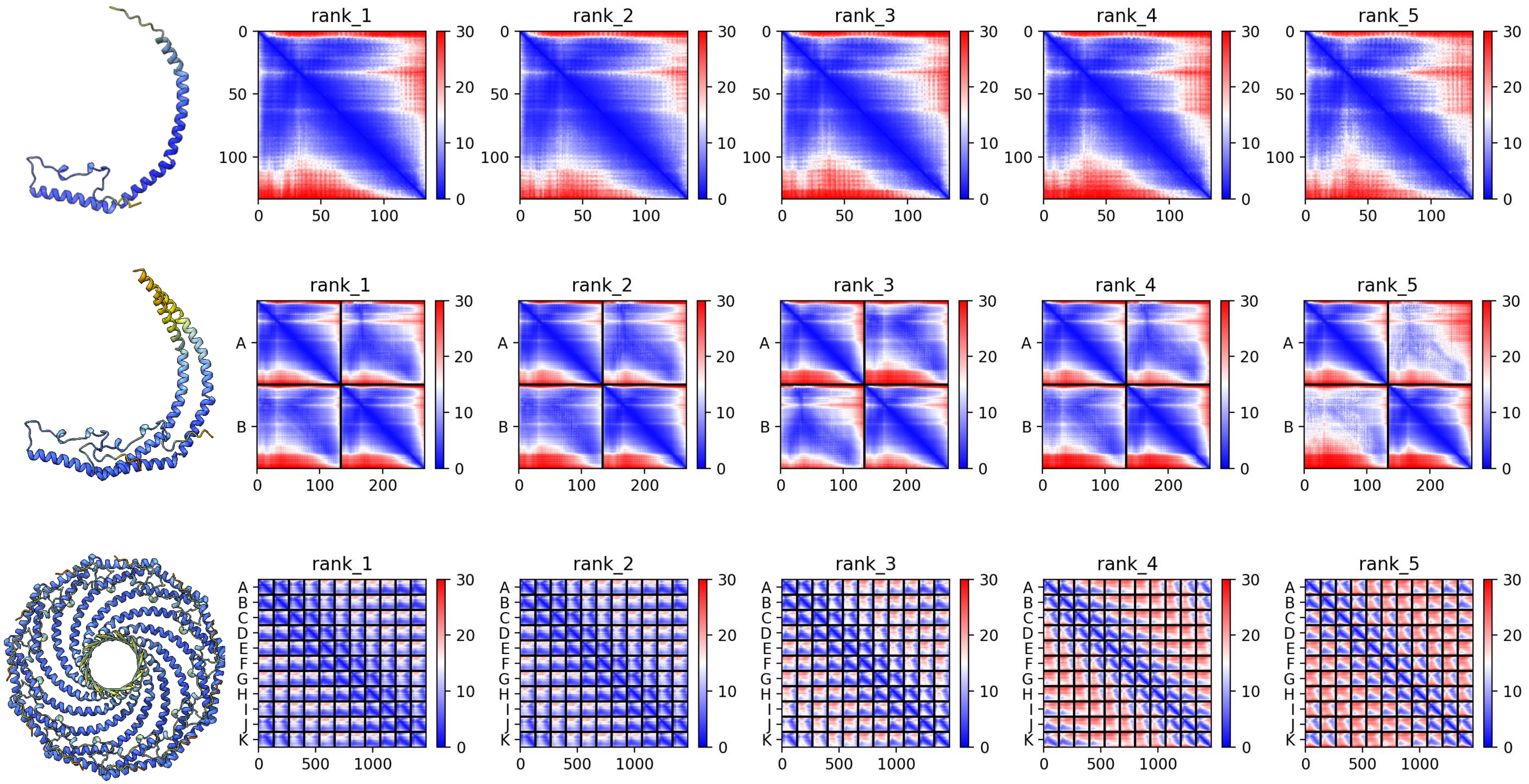

### A0A1I8GFP4 *M. lignano*

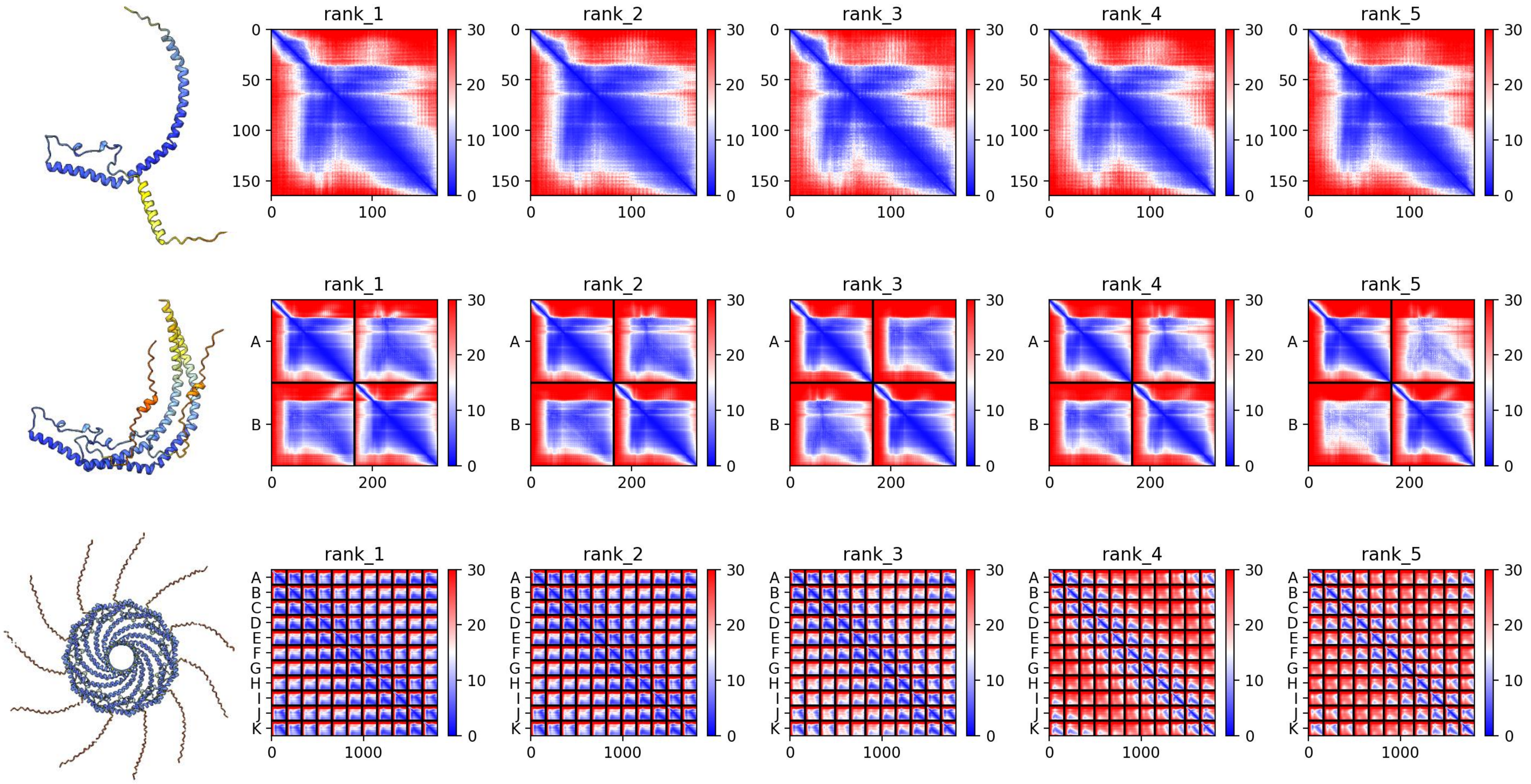

### A0A1I8G233 *M. lignano*

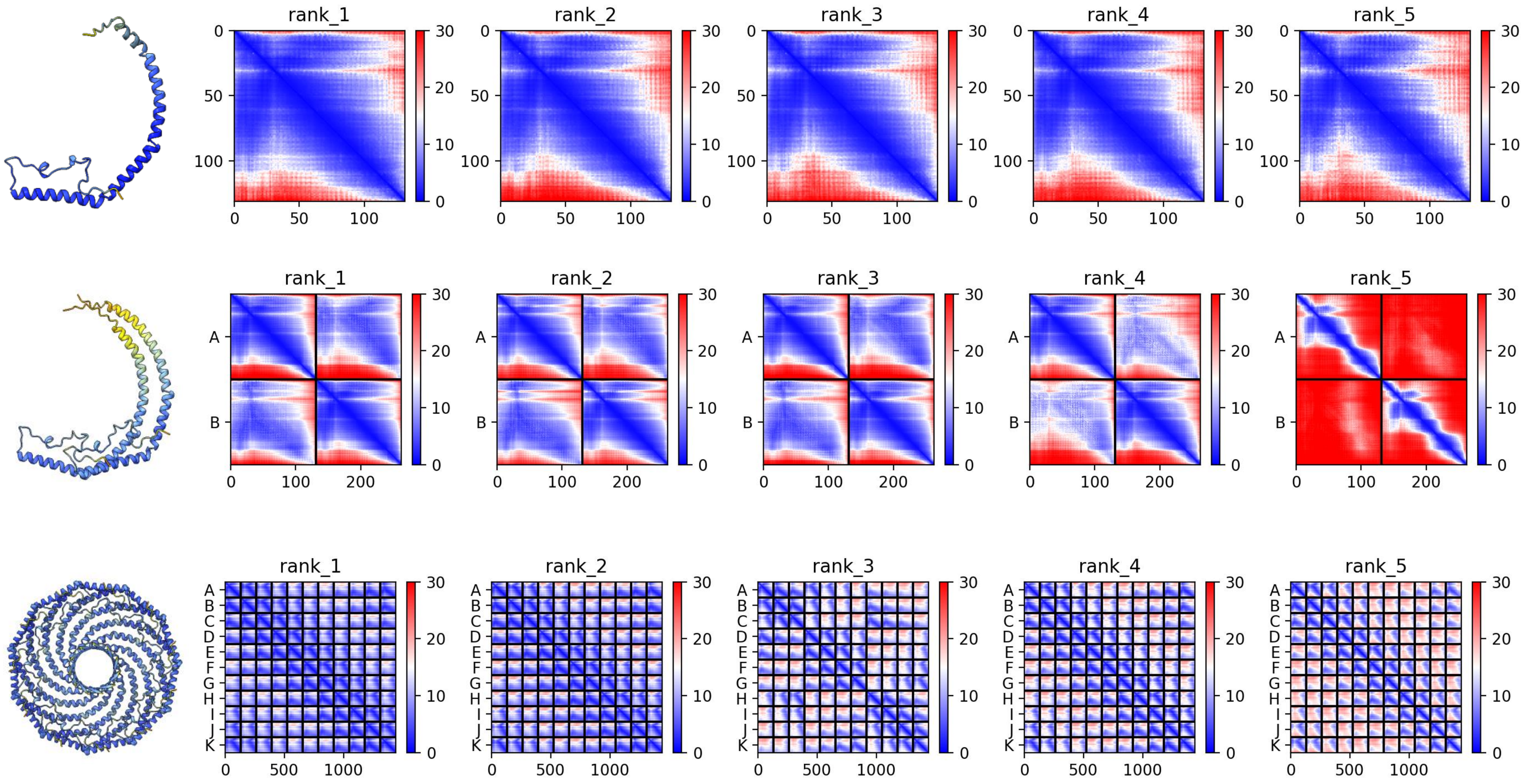

### A0A267EVV7 *M. lignano*

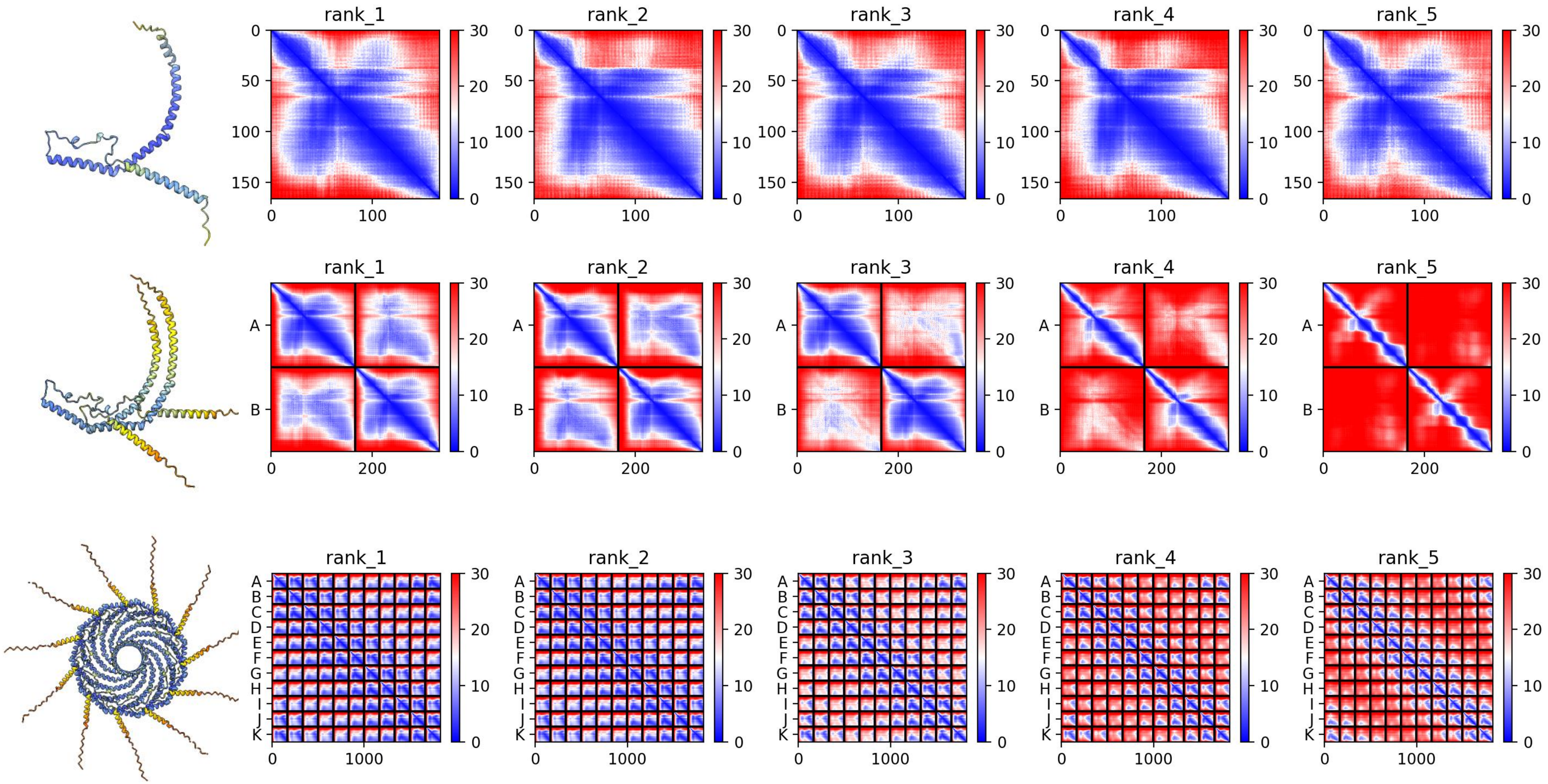

### A0A2T7NUT7 *P. canaliculata*

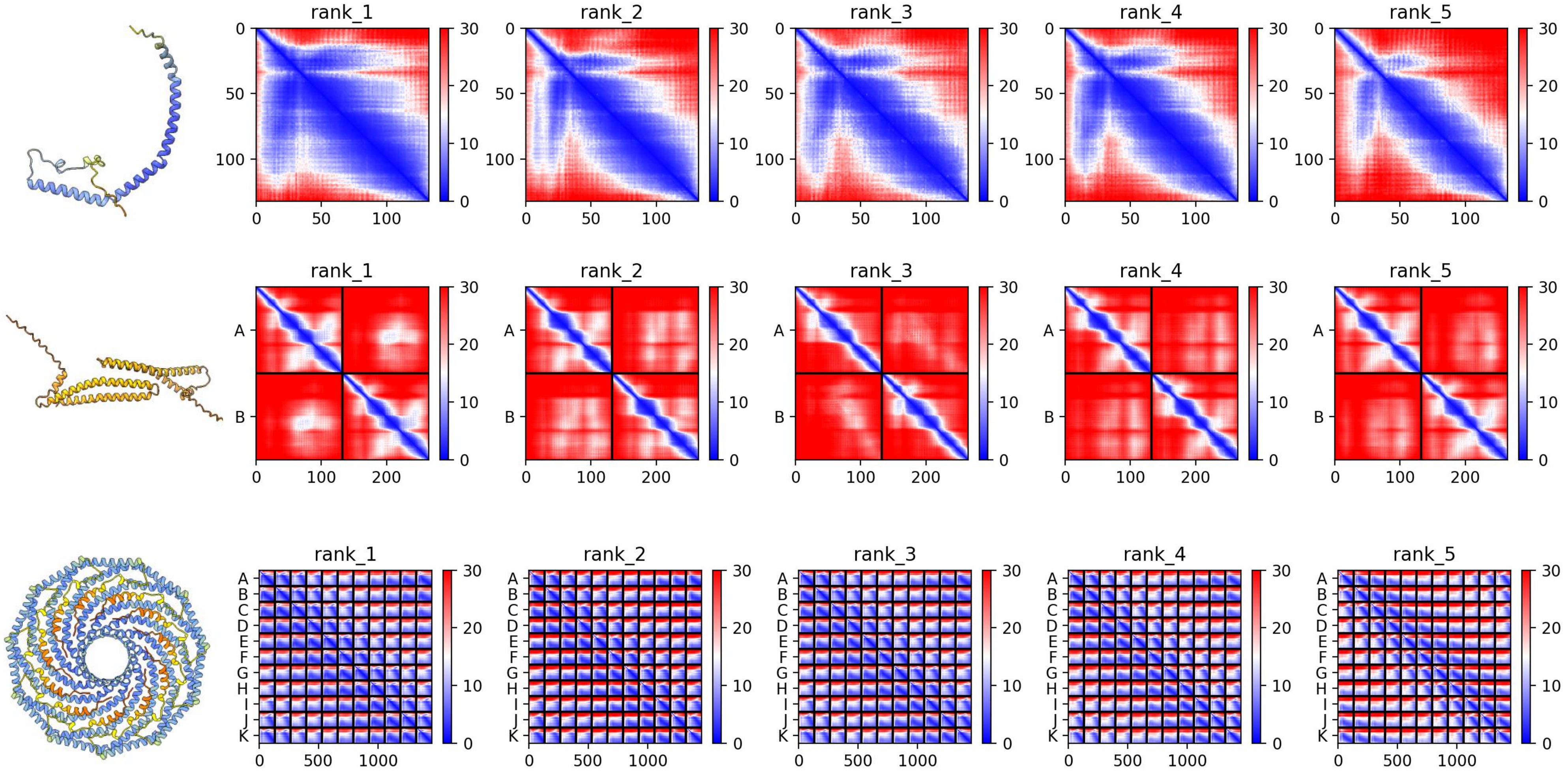

### A0A2T7NUS2 *P. canaliculata*

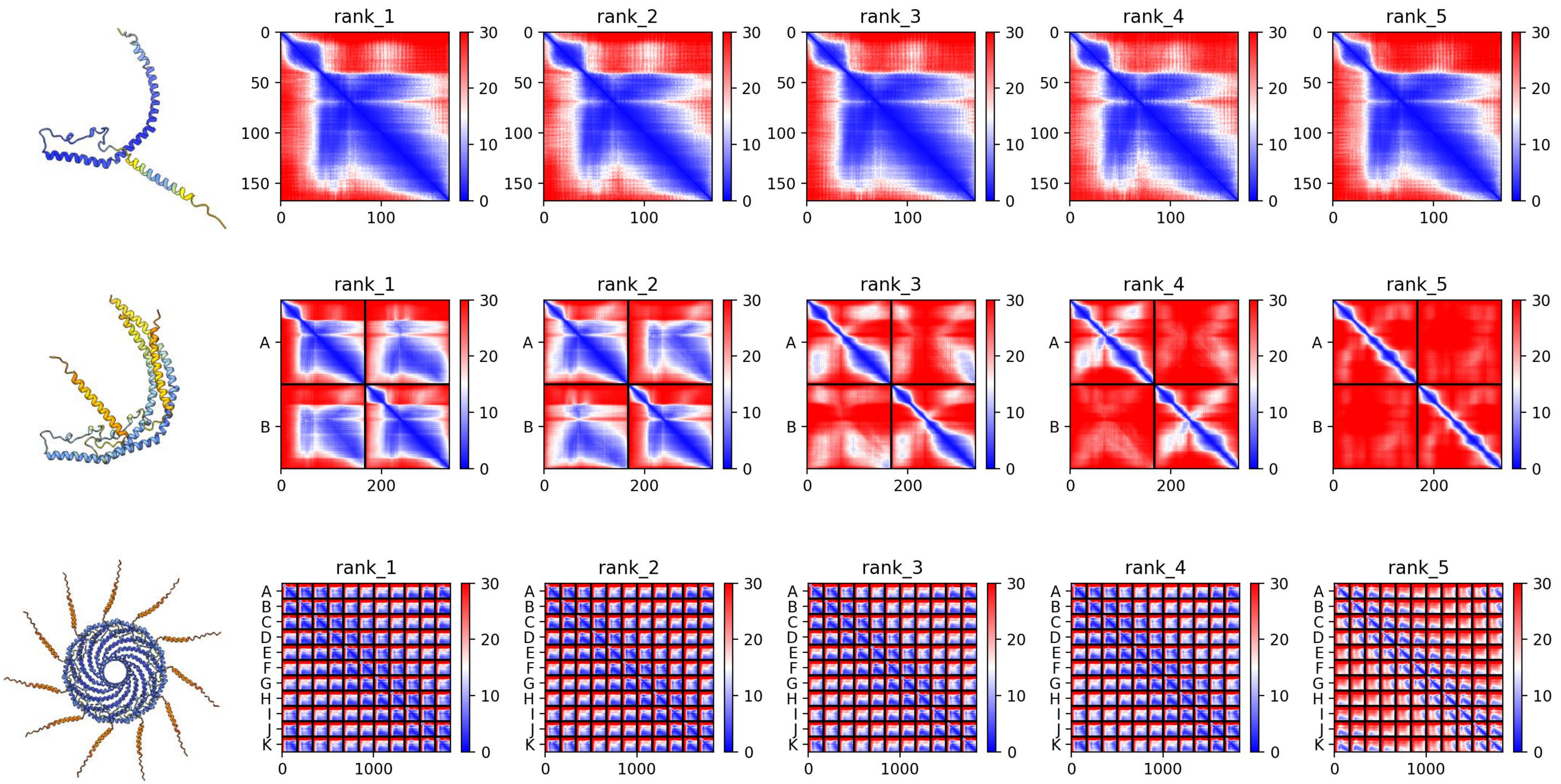

### A0A2T7NUS2 *P. canaliculata*

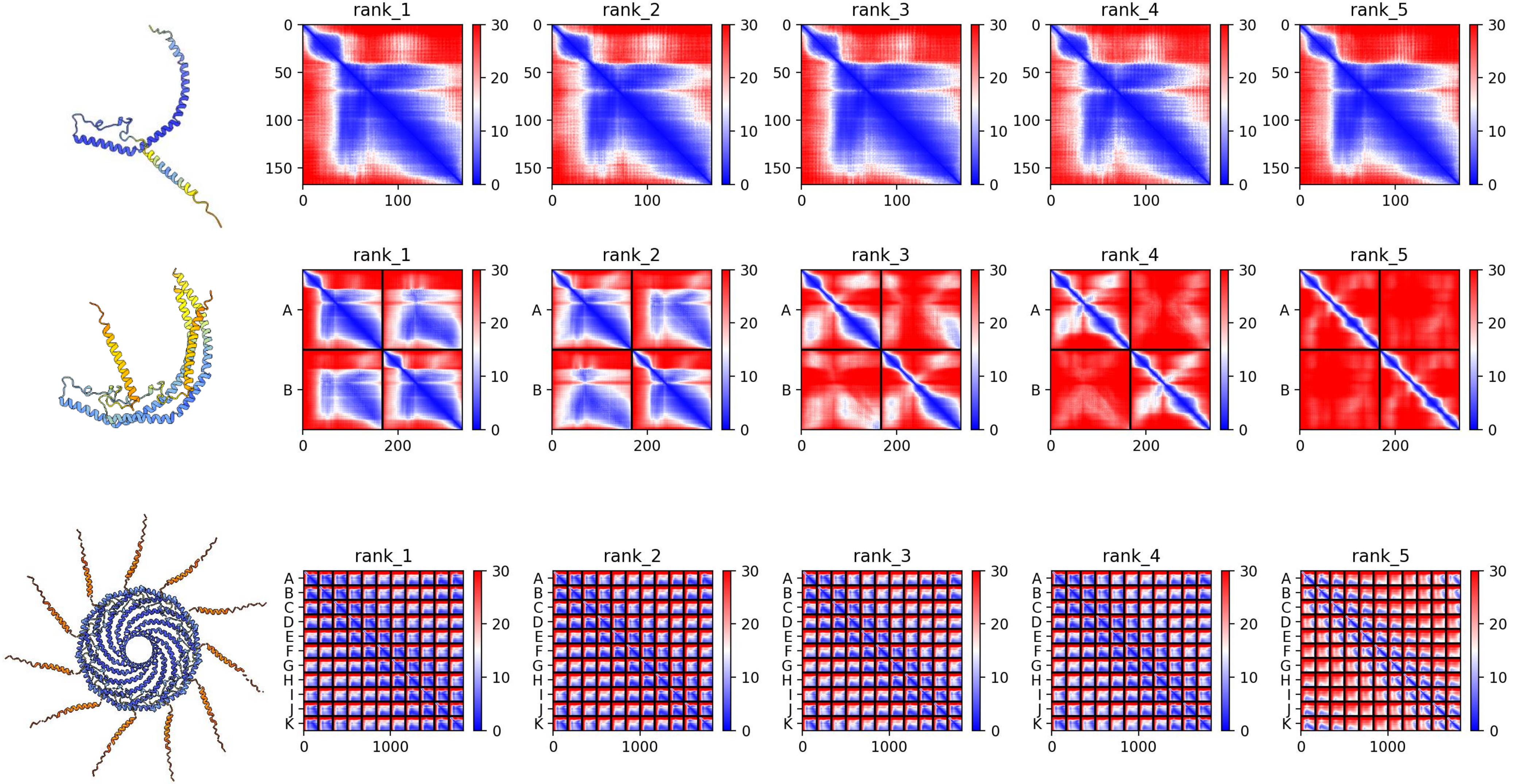

### A0A2T7NUV7 *P. canaliculata*

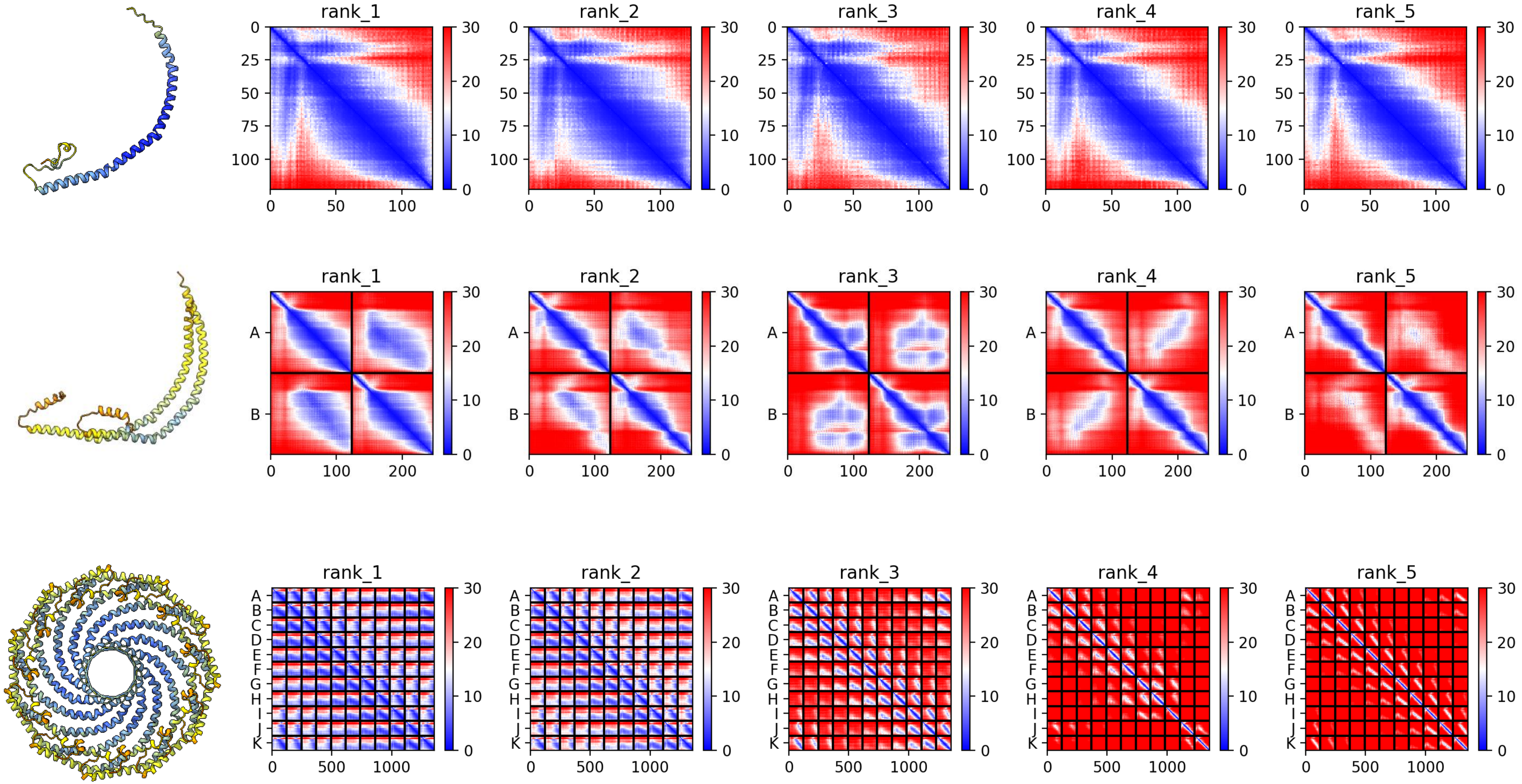

### R7VDS7 *C. teleta*

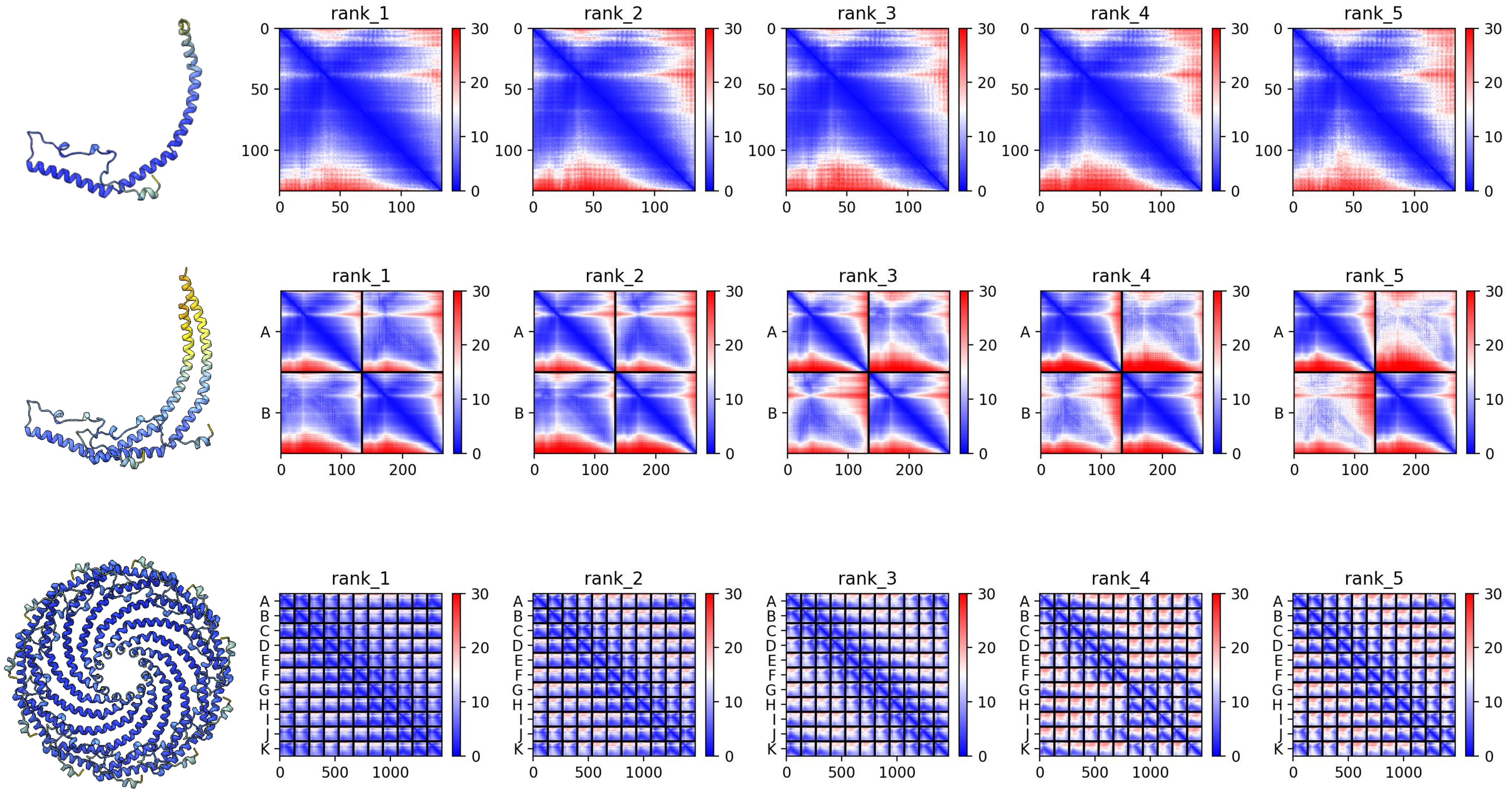

### R7V531 *C. teleta*

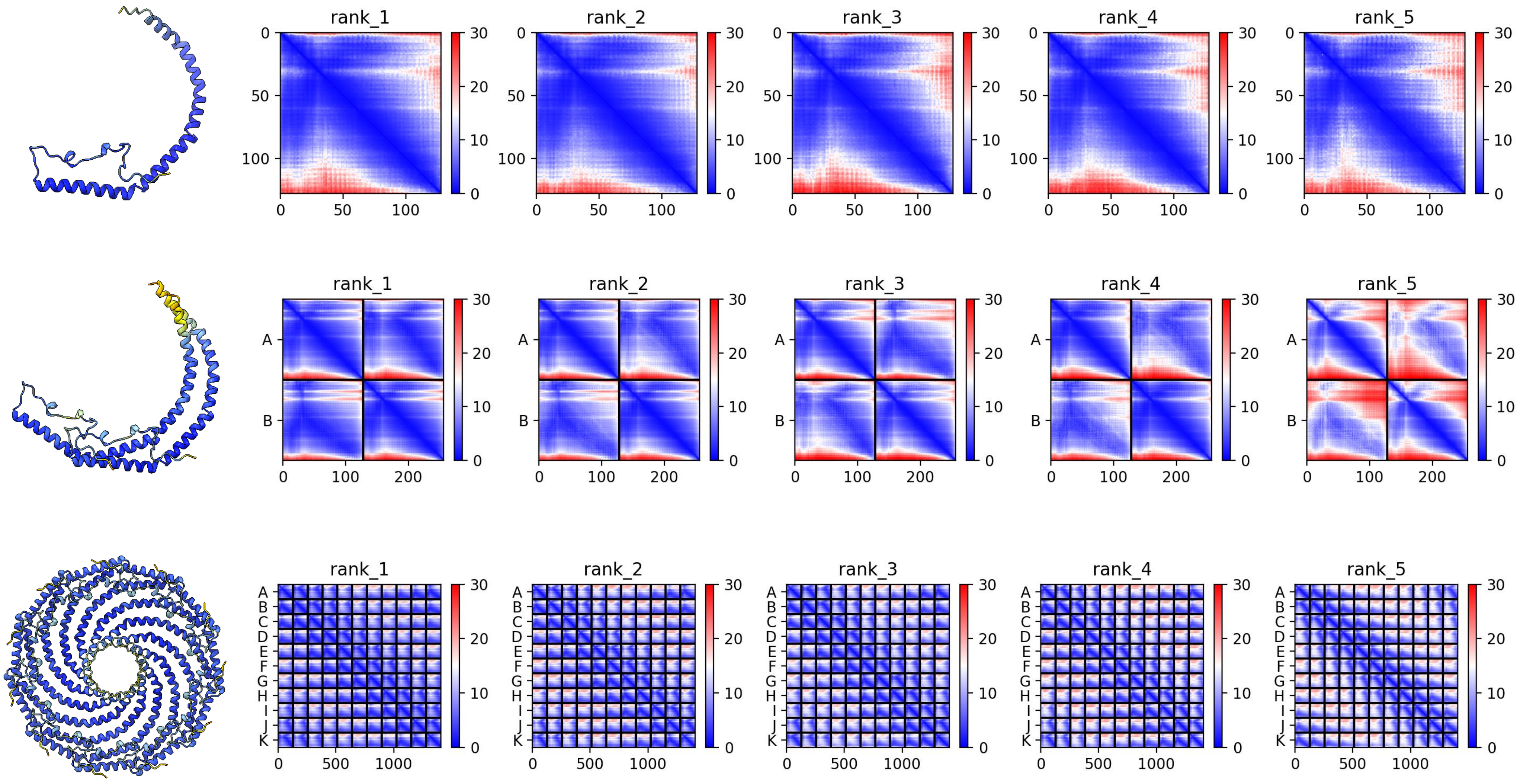

### A0A7M7RAC4 *S. purpuratus*

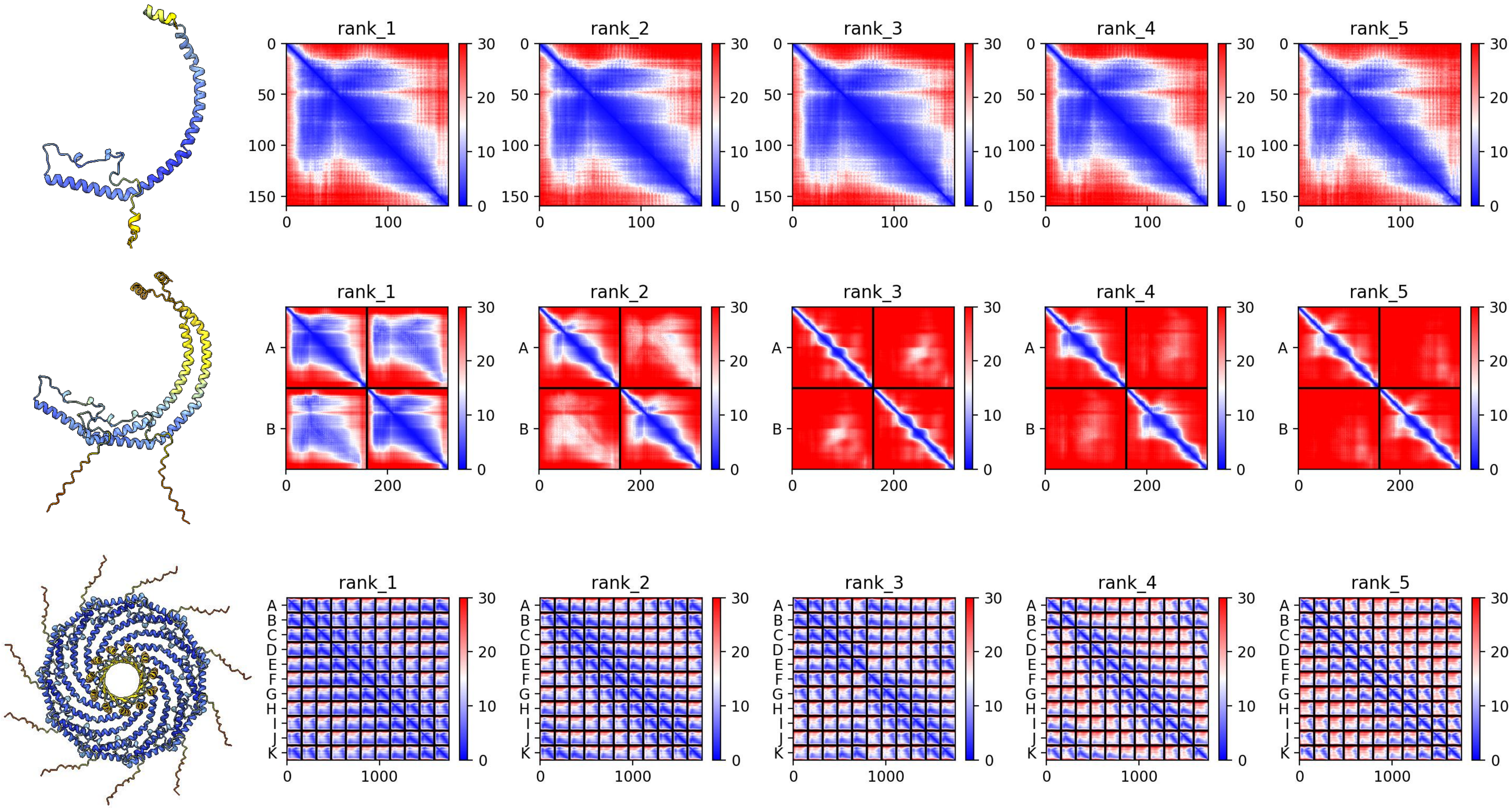

### A0A2B4SEV1 *S. pistillata*

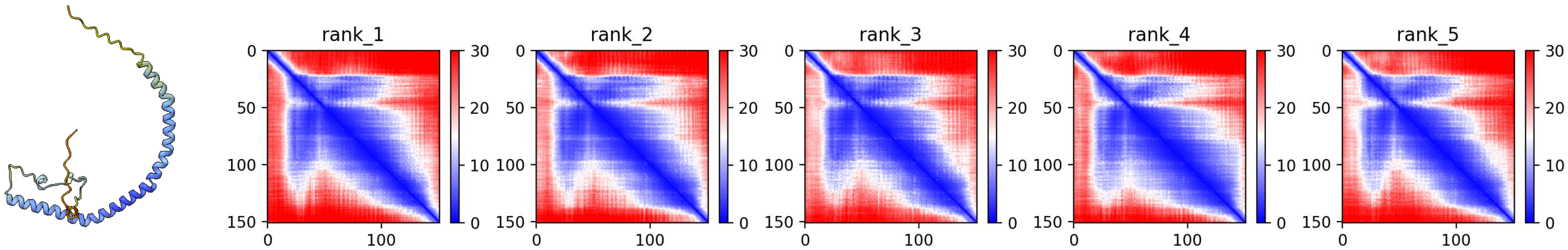


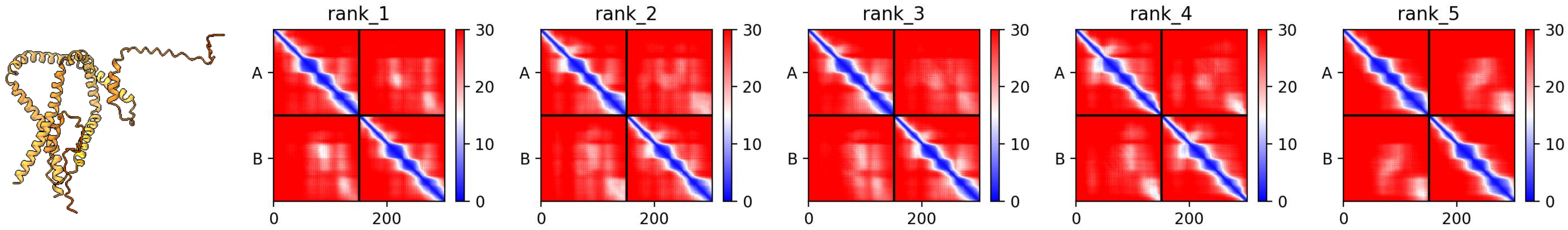


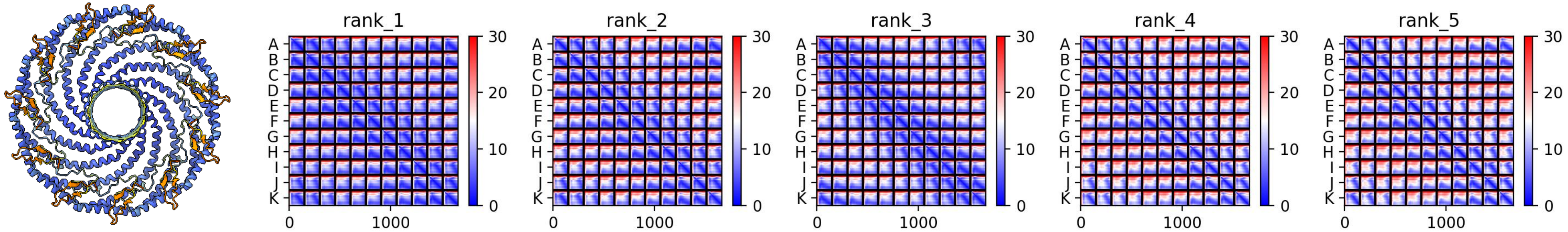

### A0A2B4SAZ9 *S. pistillata*

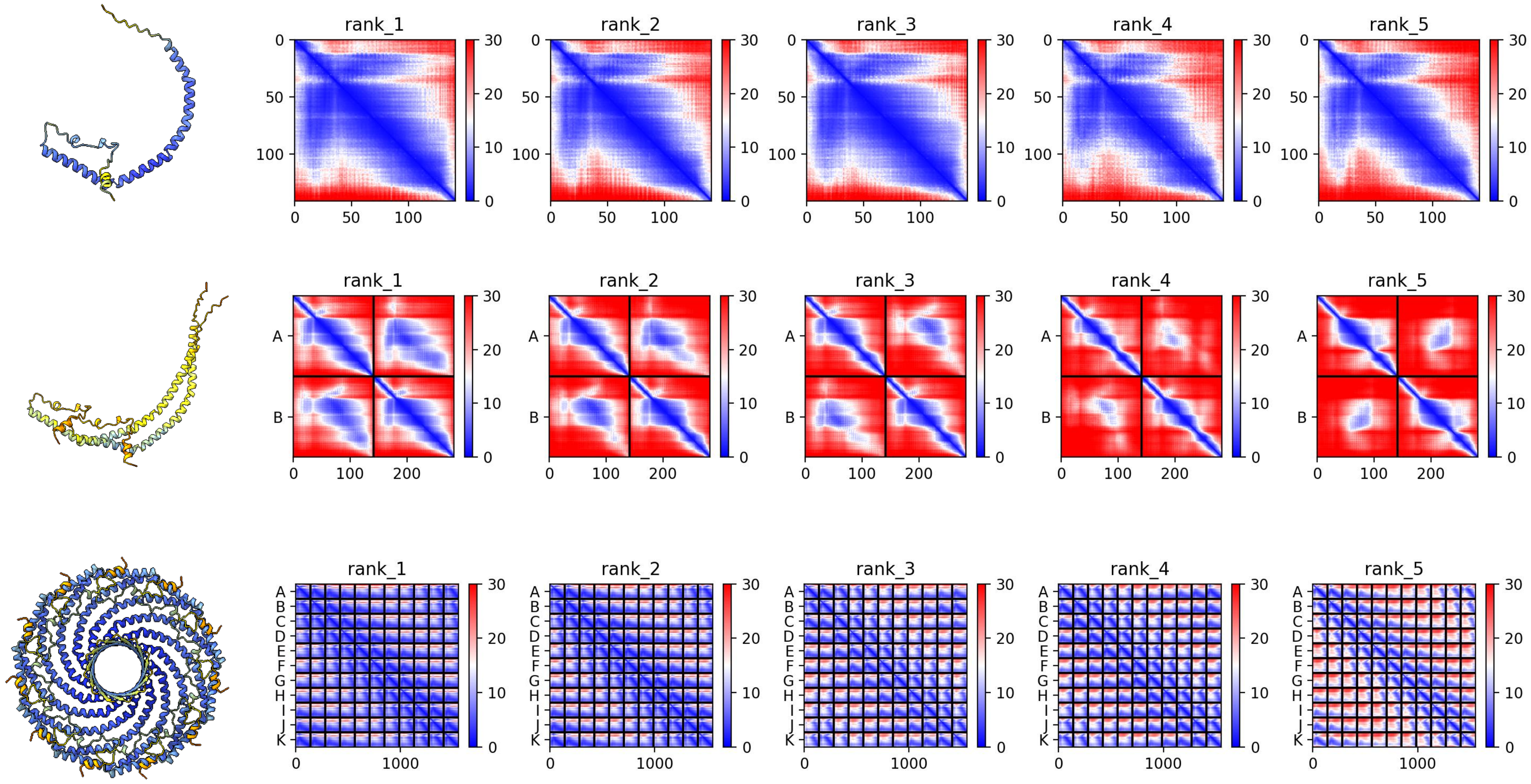

### XP_065842155.1 *O. lobularis*

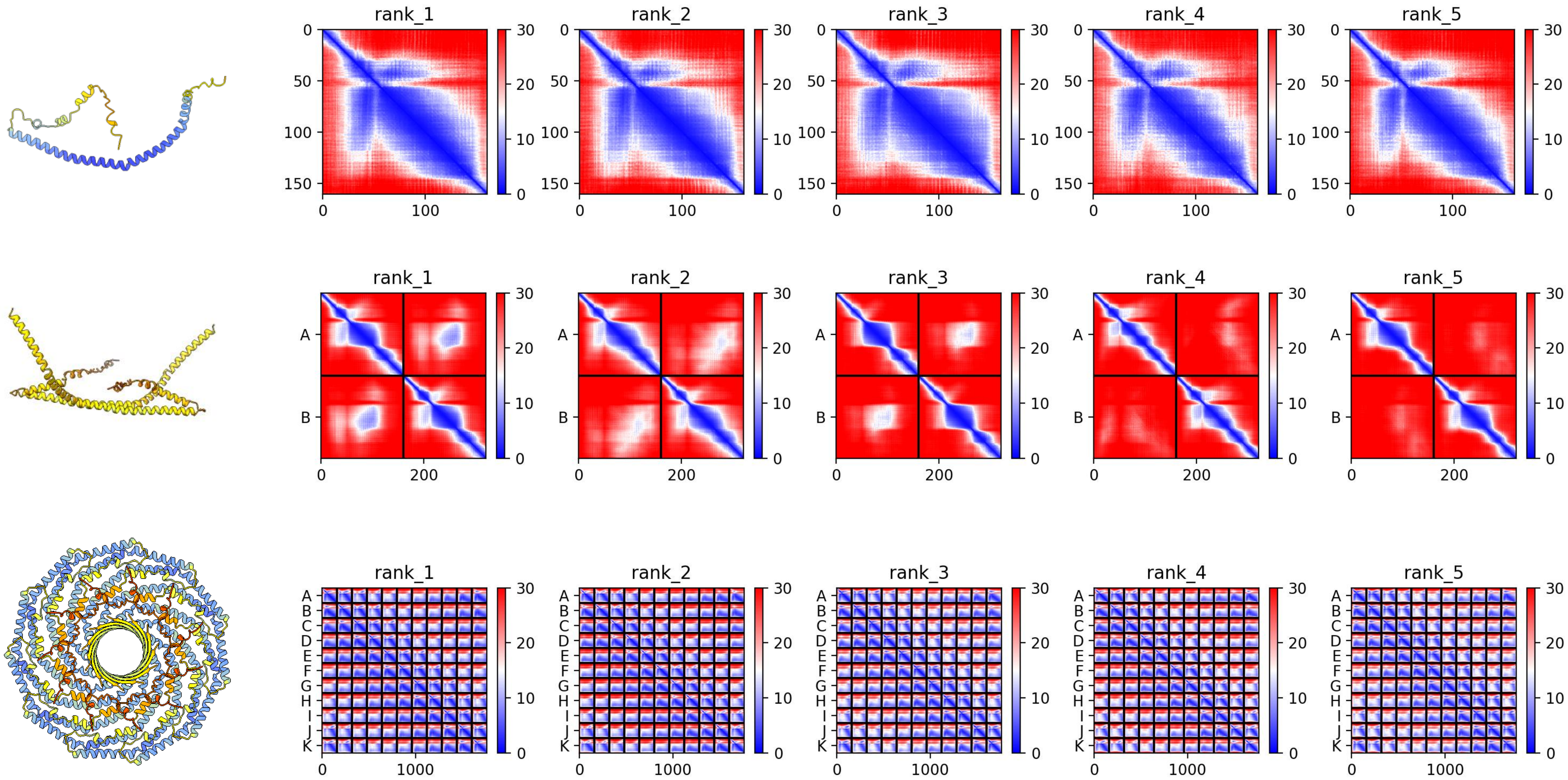

### XP_065840741.1 *O. lobularis*

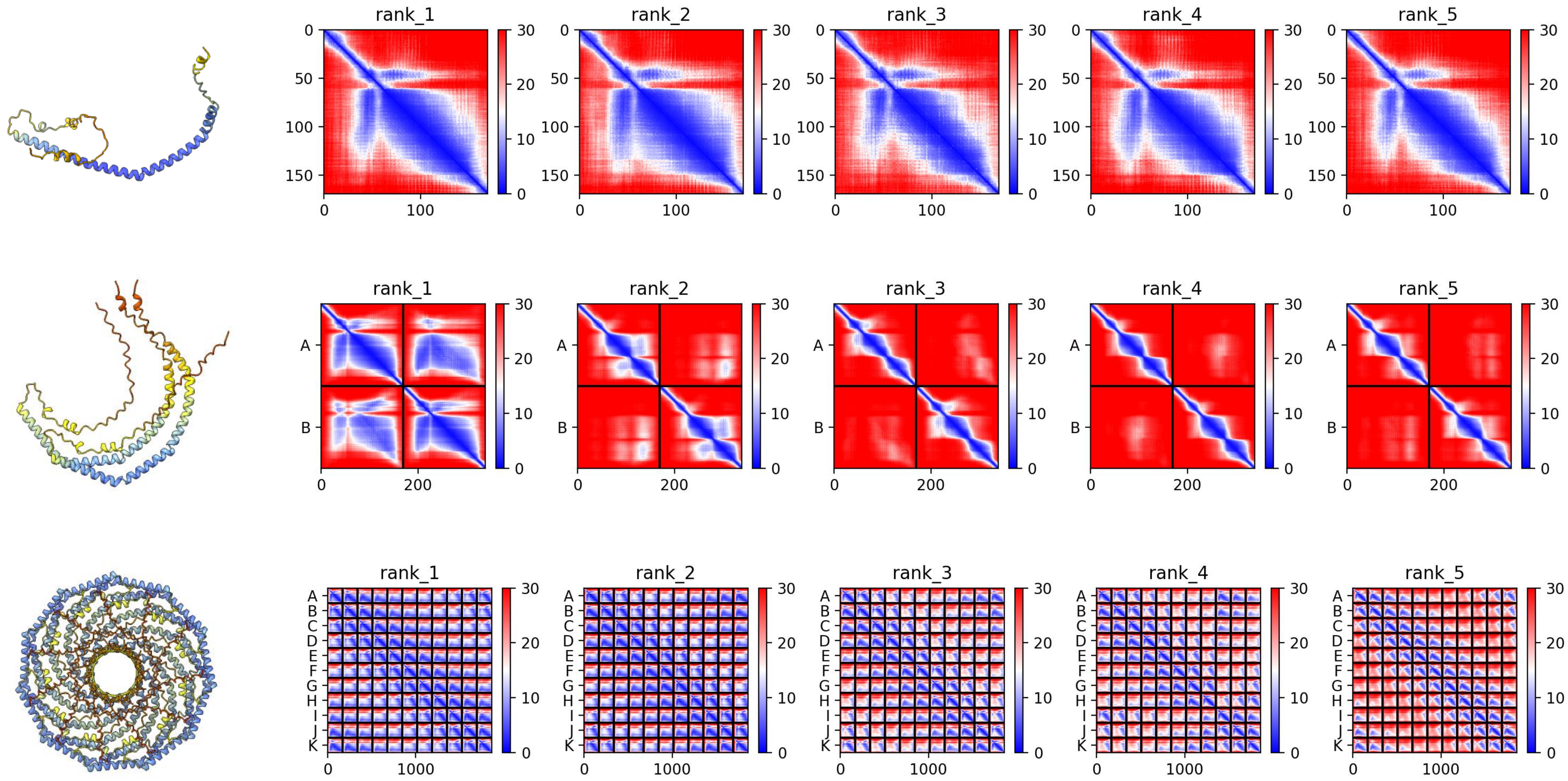

### Em0021g891a *E. muelleri*

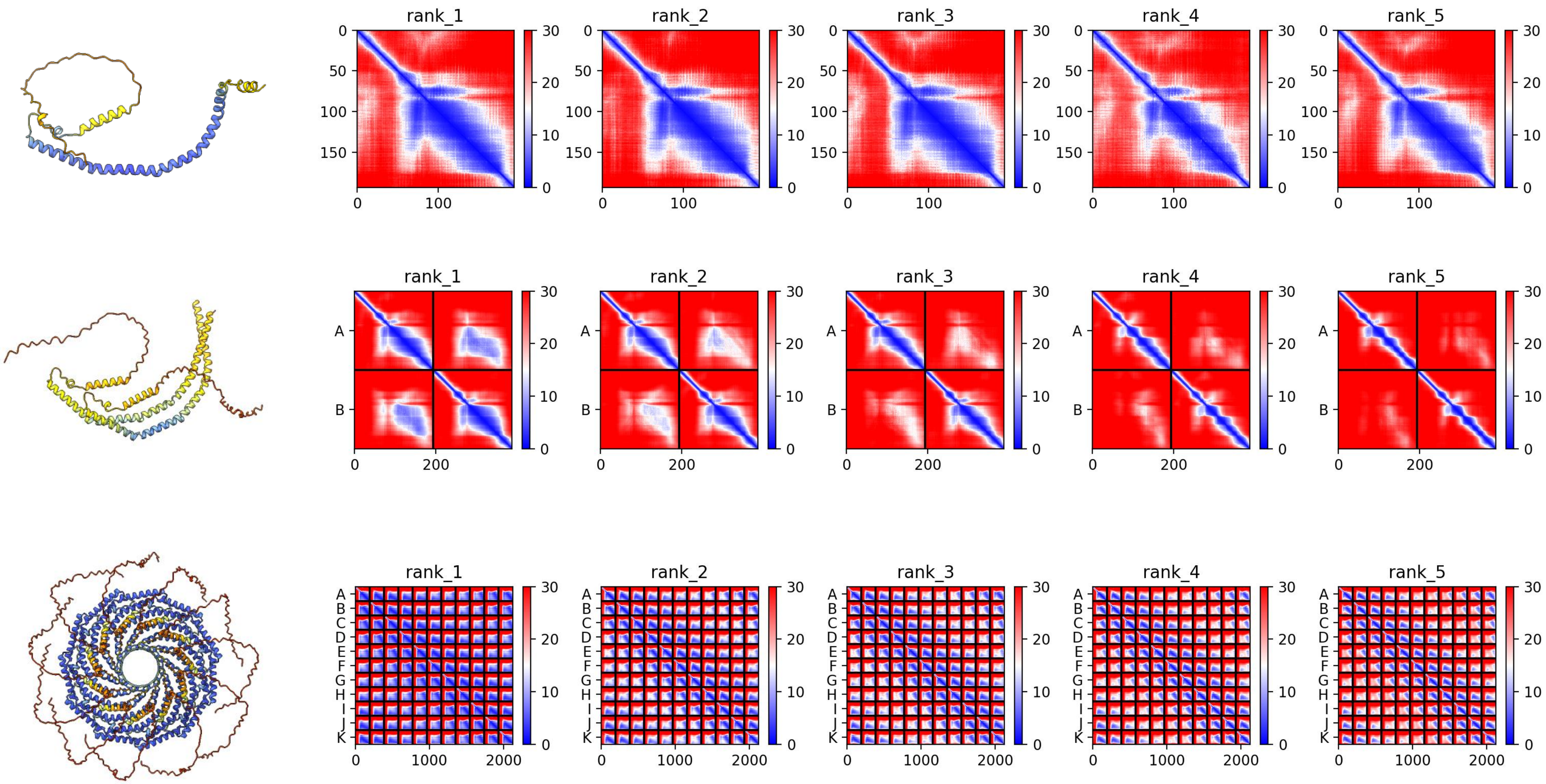

### Em0021g893a *E. muelleri*

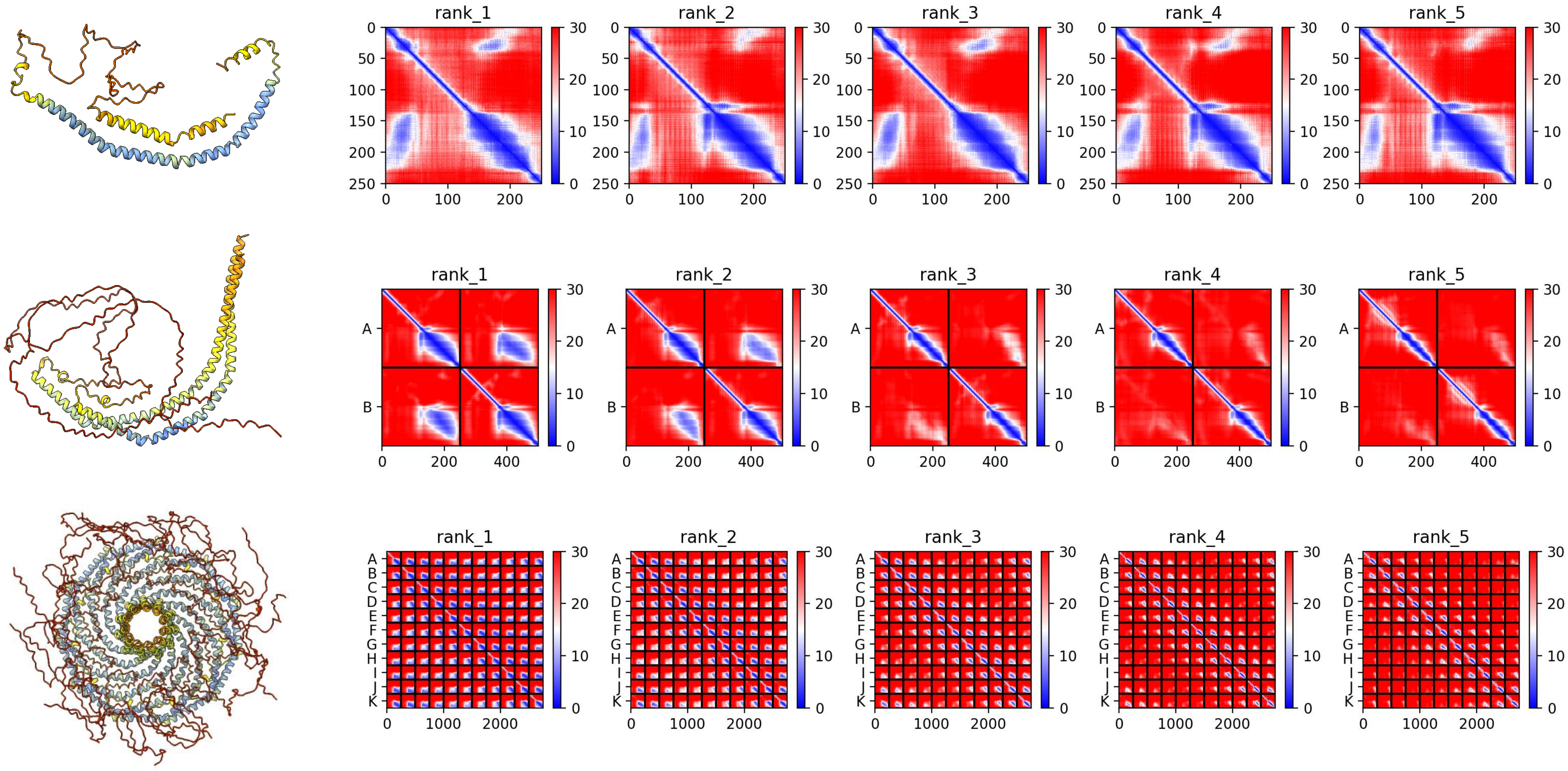

### XP_001633849.3 *N. vectensis*

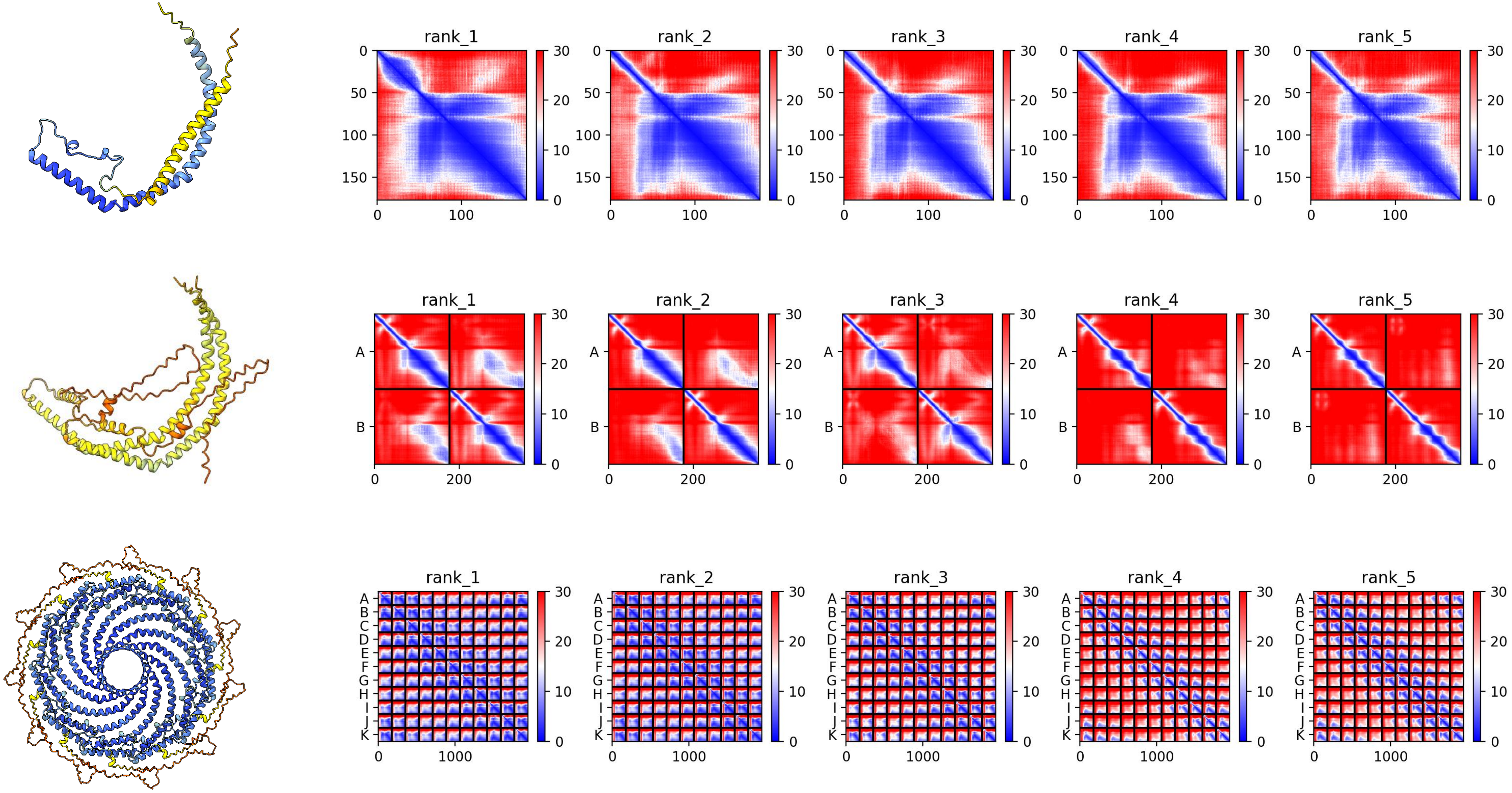

### XP_001625439.2 *N. vectensis*

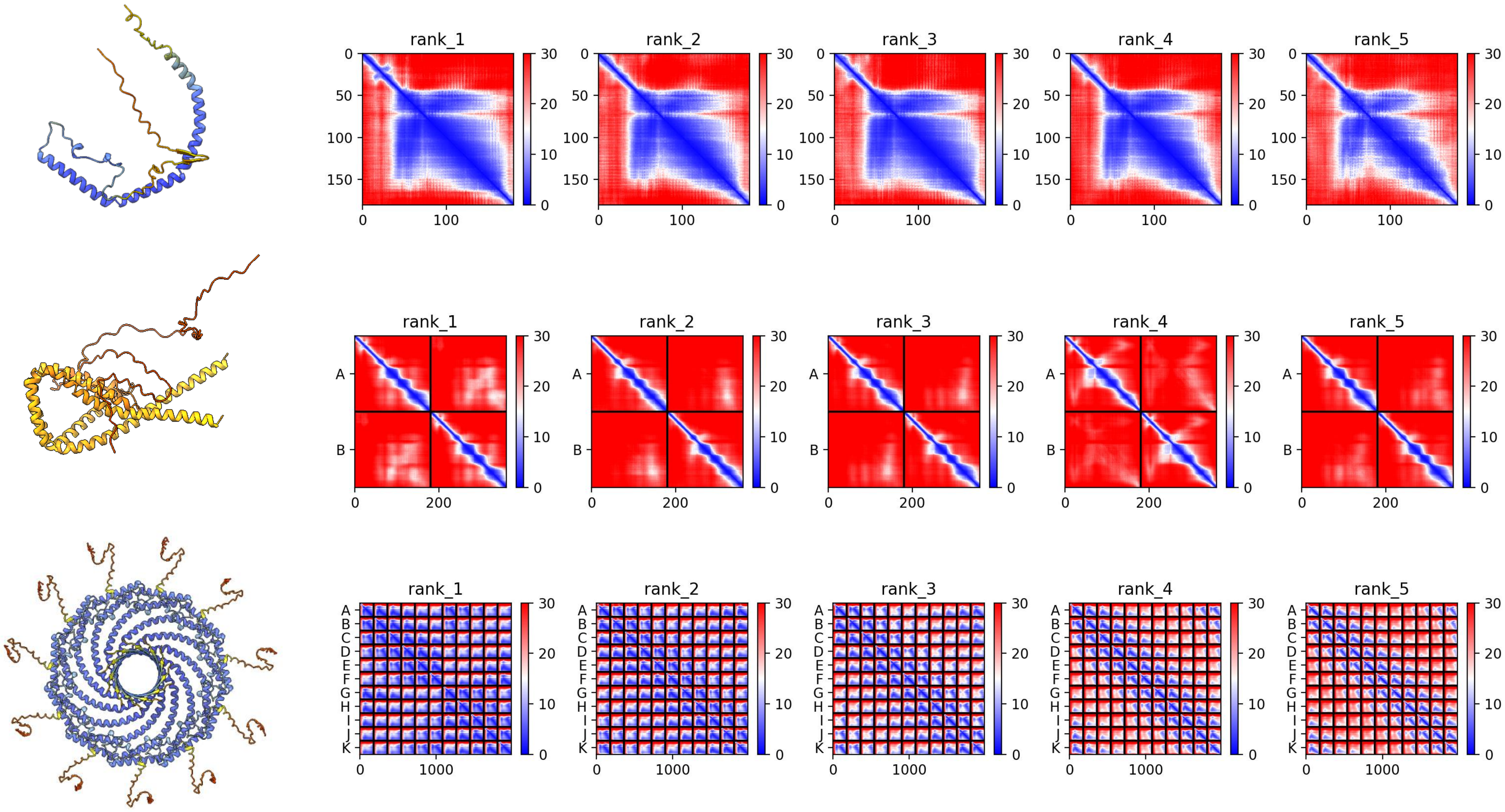

### XP_029191350.1 *A. millepora*

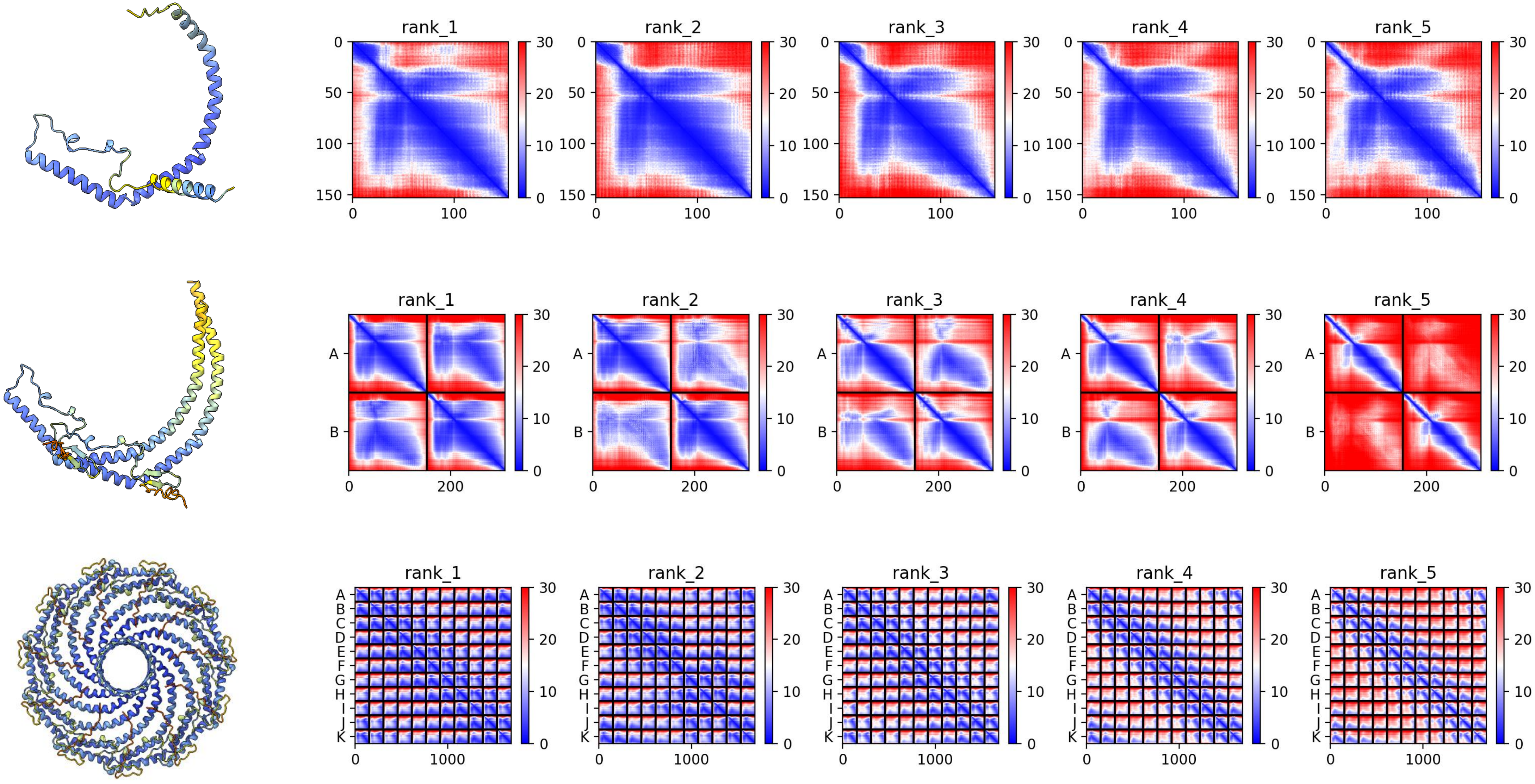

### XP_033745936.1 *P. maximus*

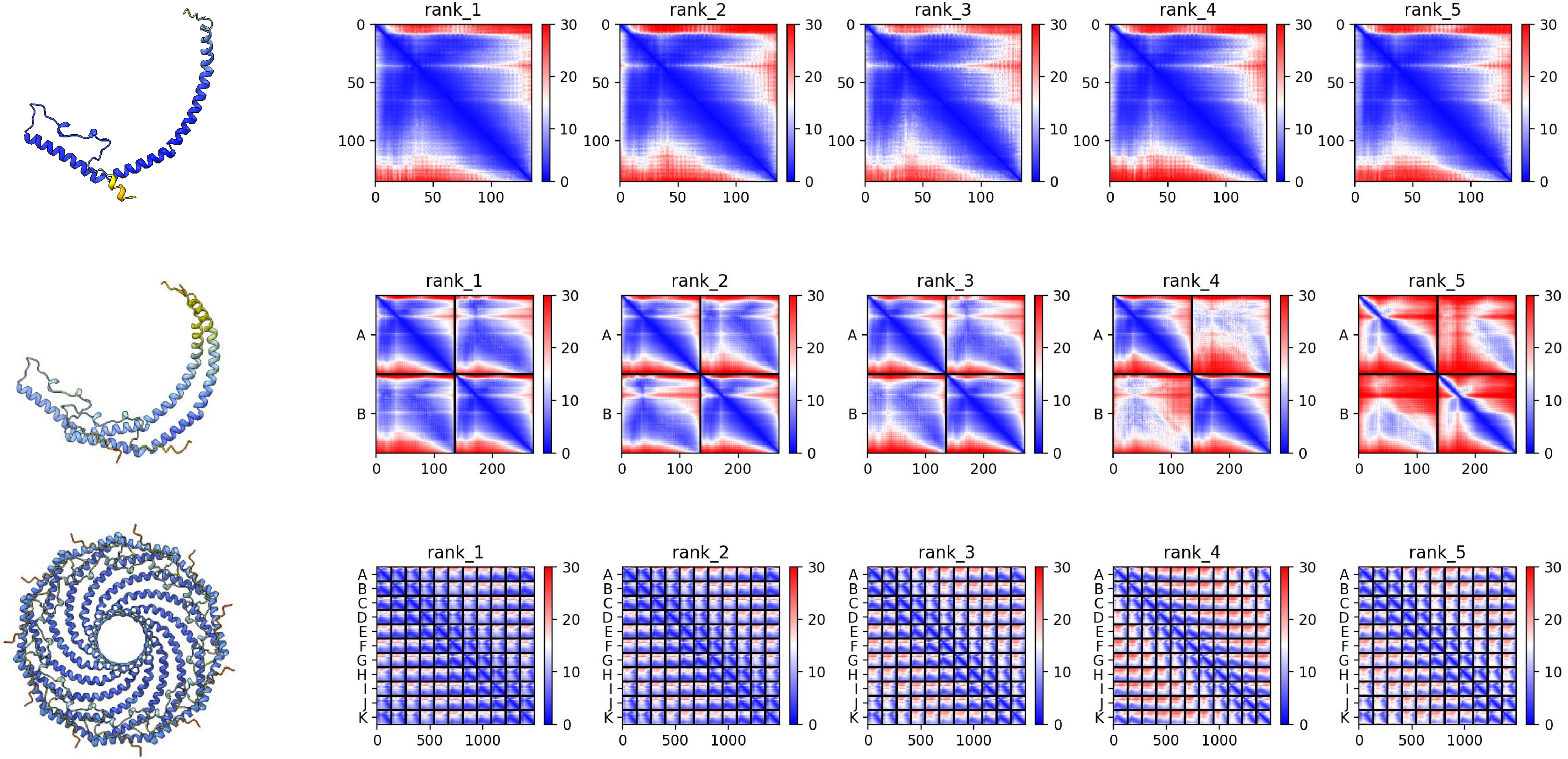

### XP_033740634.1 *P. maximus*

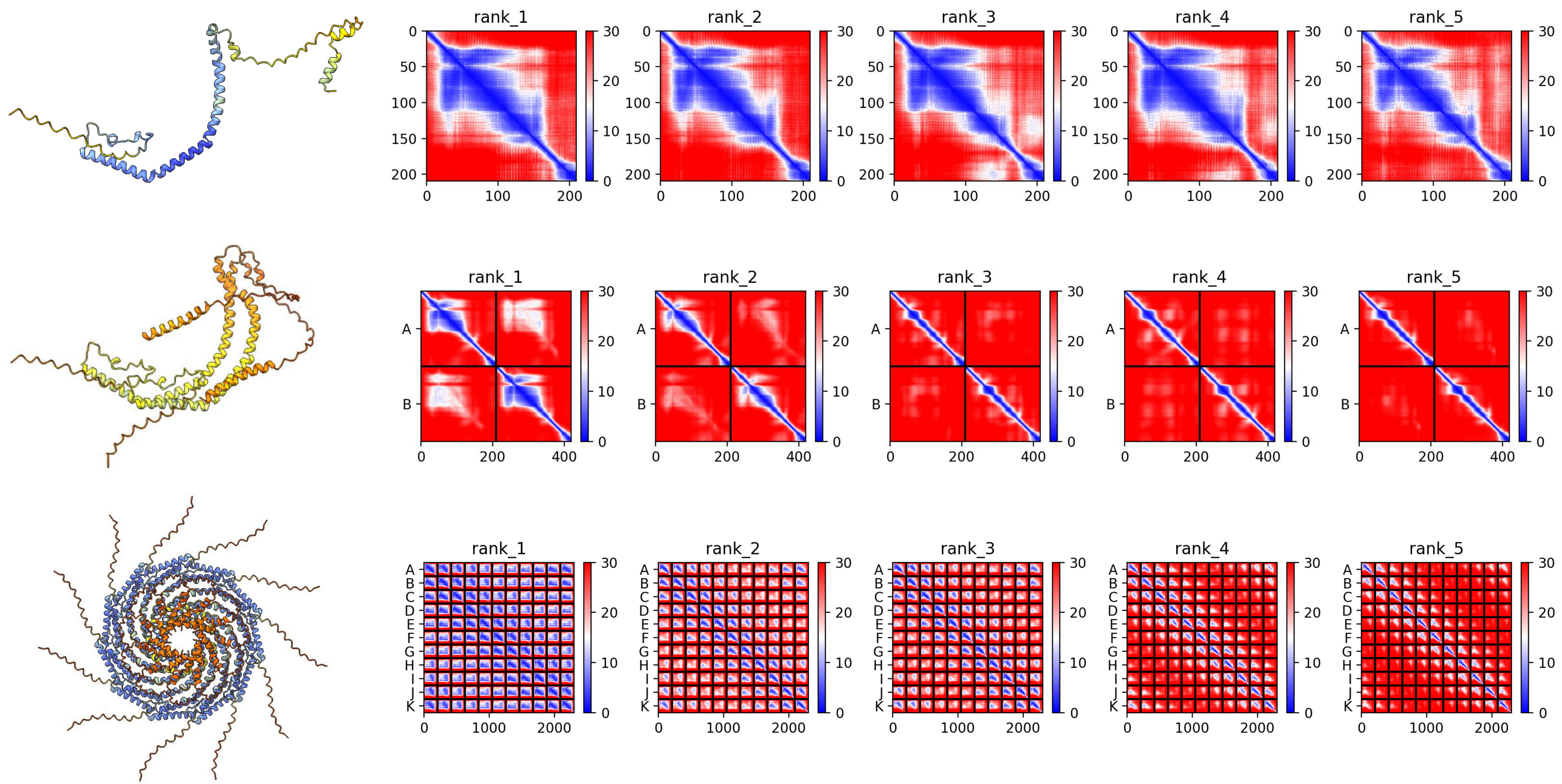

#### XP_052250592.1 *D. polymorpha*

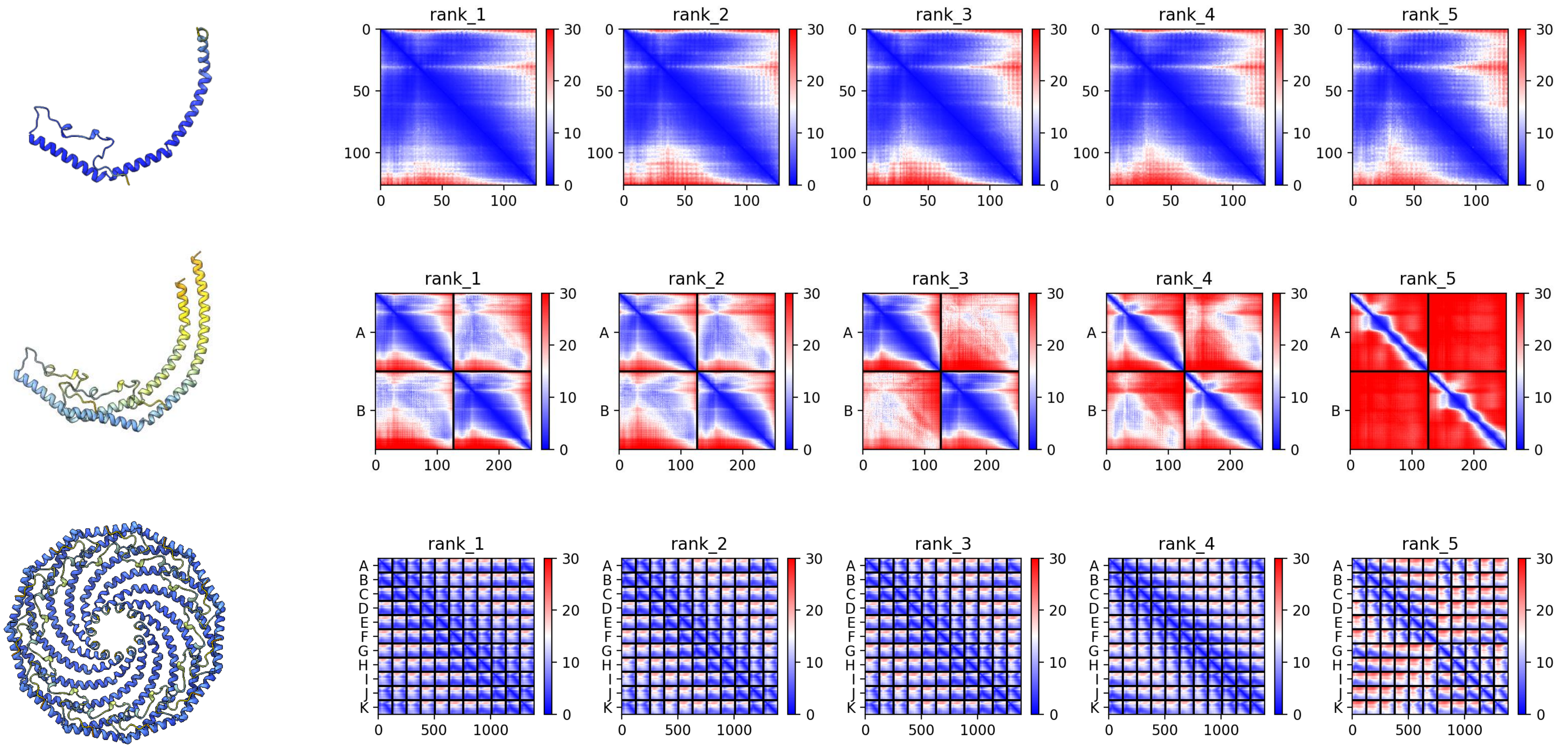

### XP_052249606.1 *D. polymorpha*

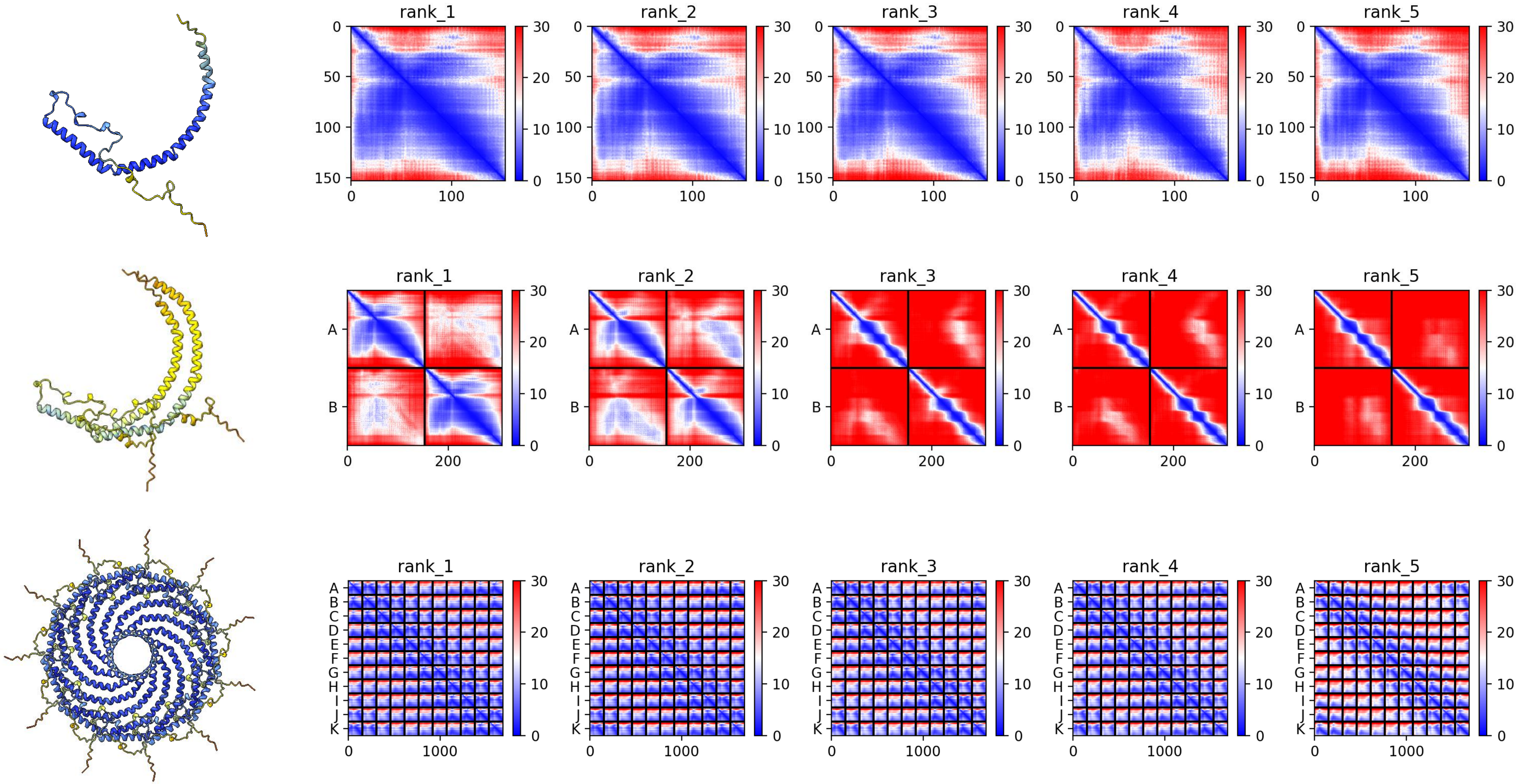

### XP_052234244.1 *D. polymorpha*

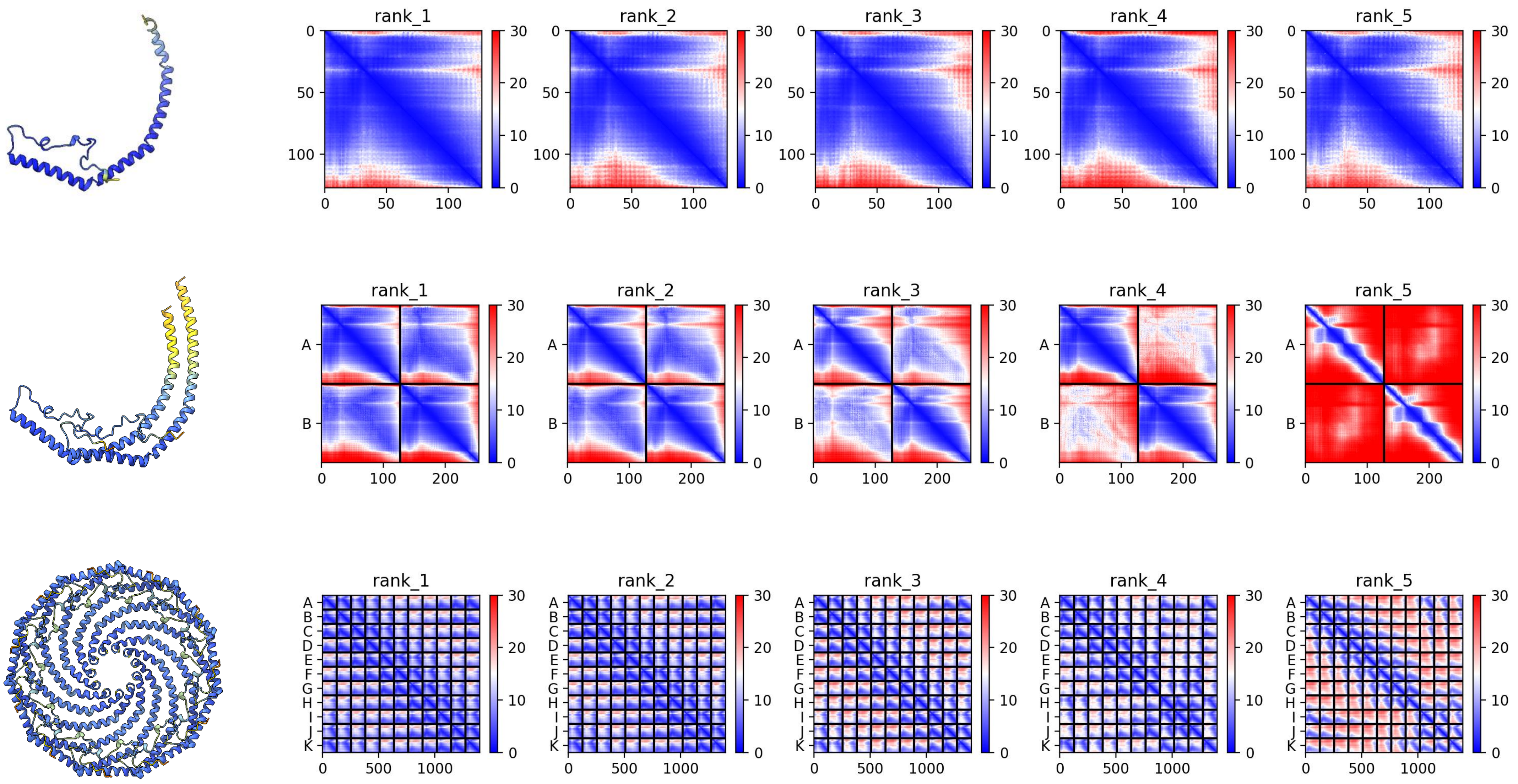

#### XP_013092506.1 *B. glabrata*

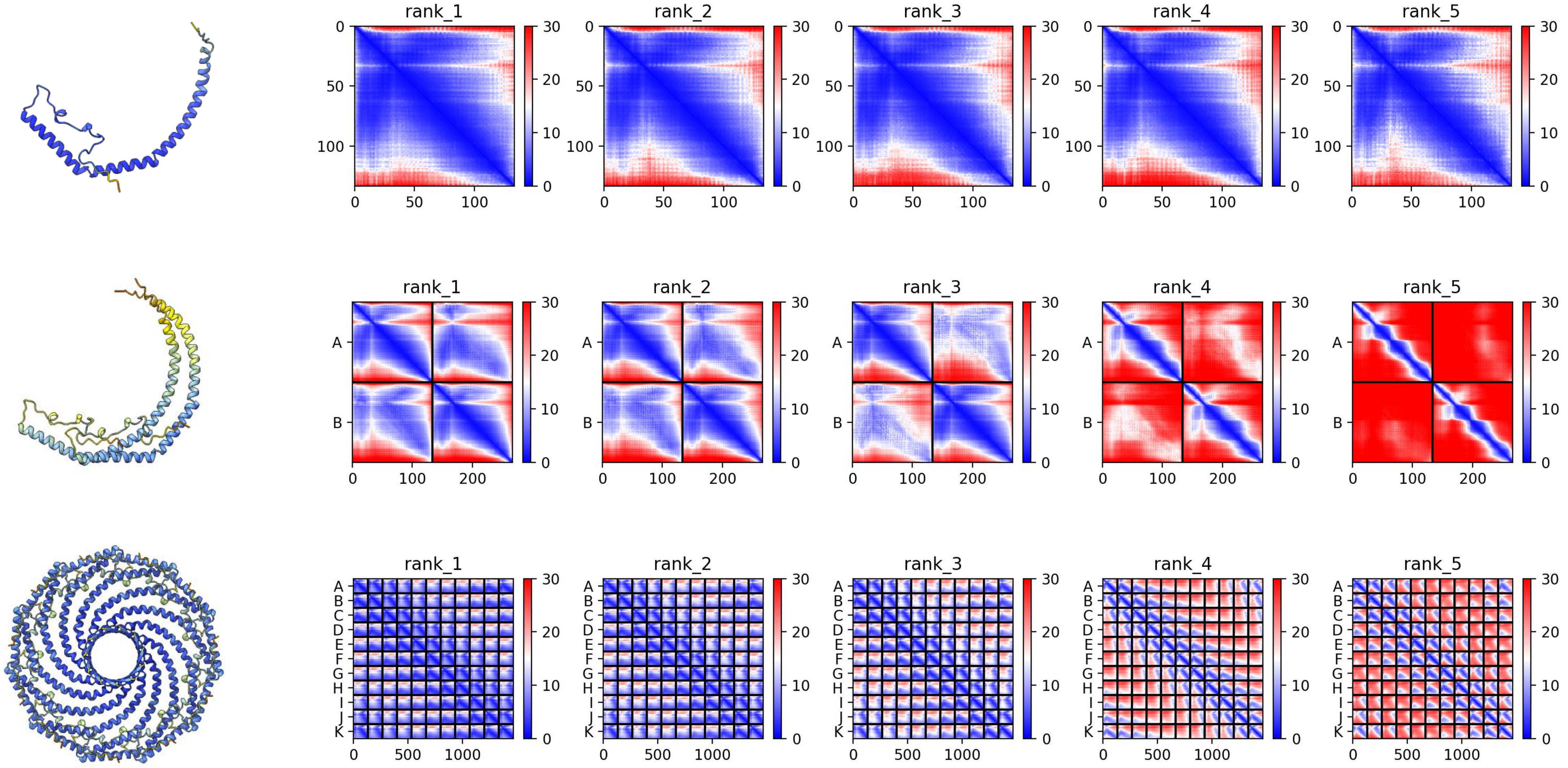

### XP_050413640.1 *P. vulgata*

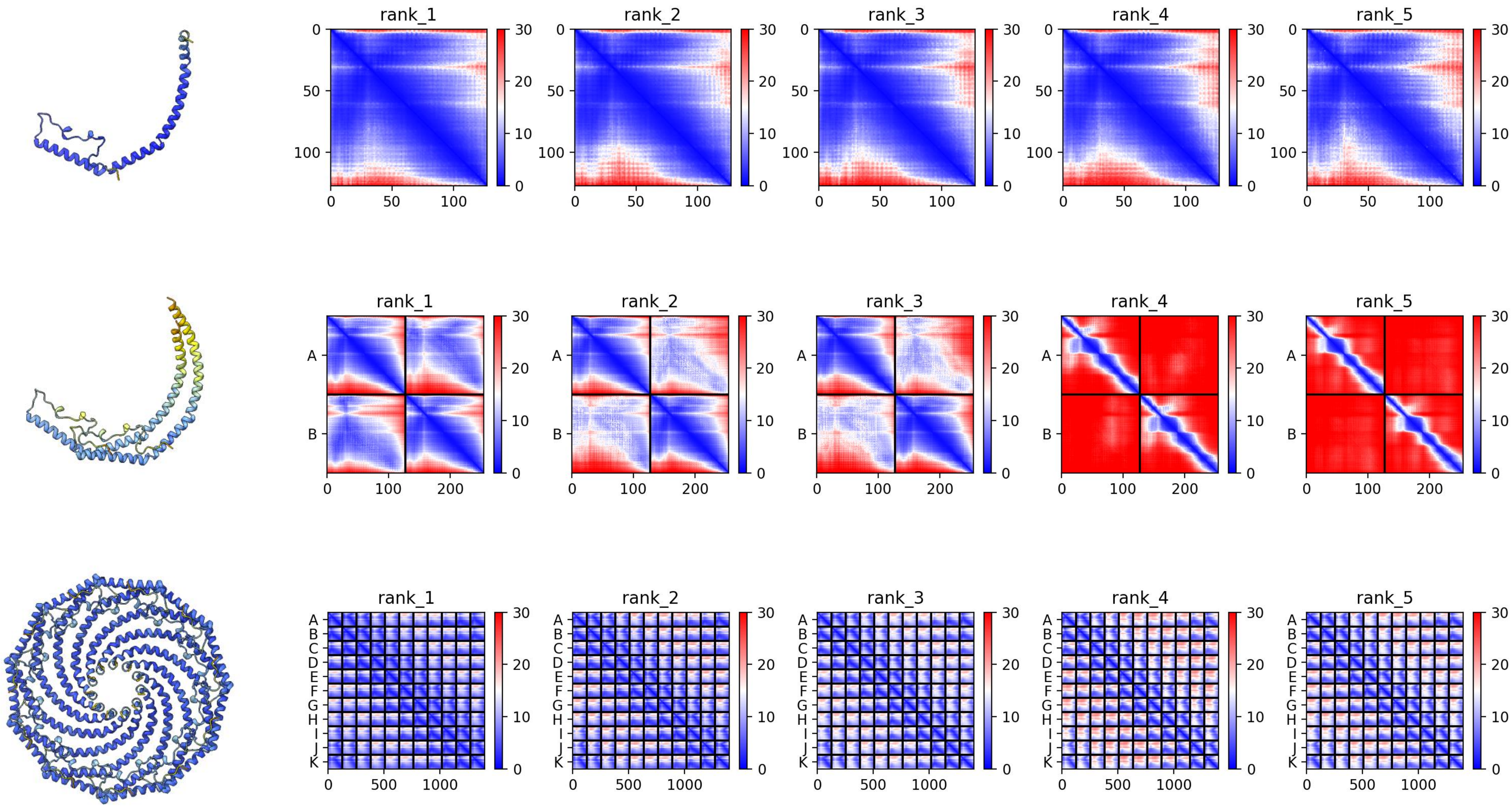

### XP_055958200.1 *P. vulgata*

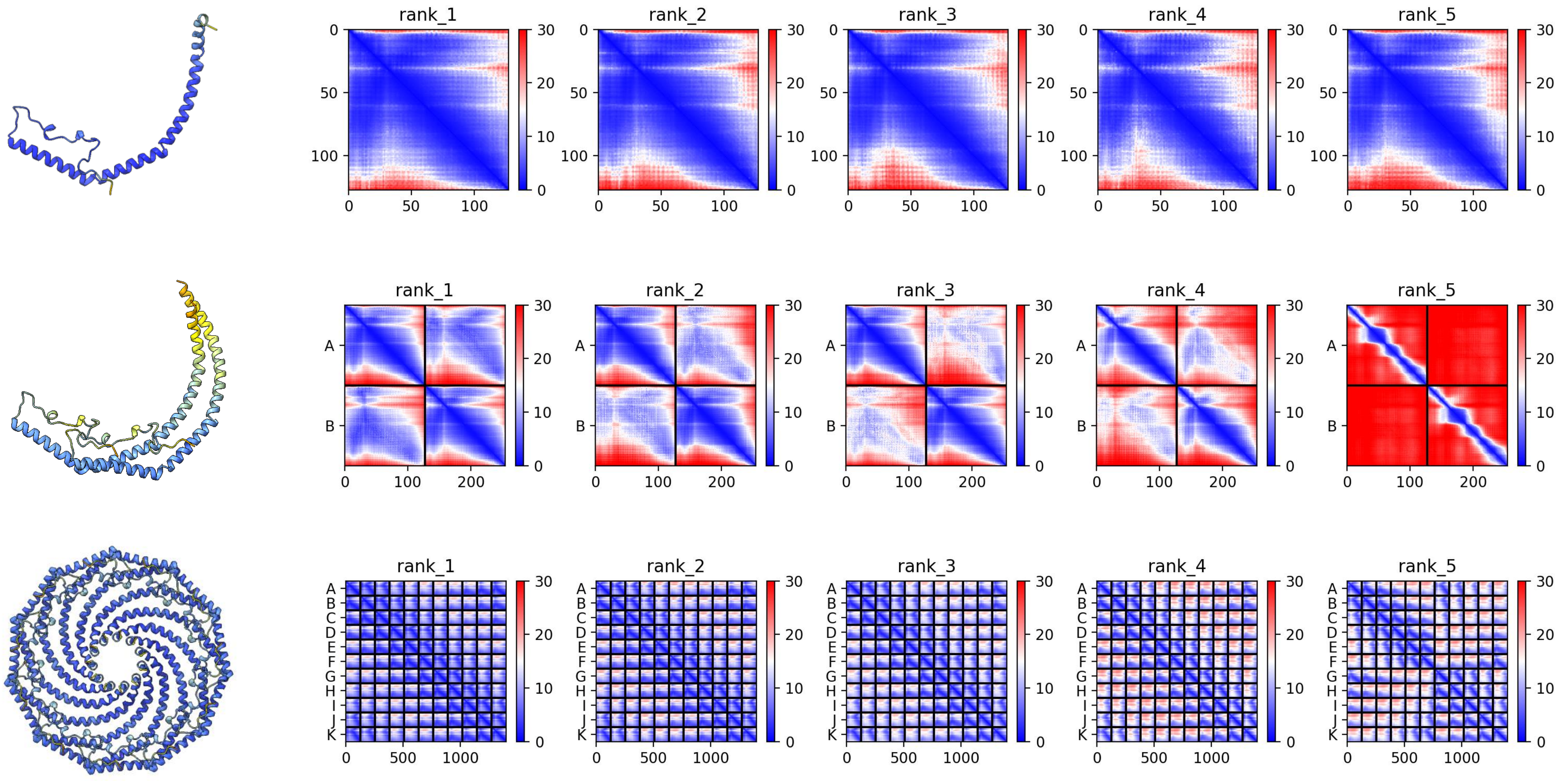

### XP_050414932.1 *P. vulgata*

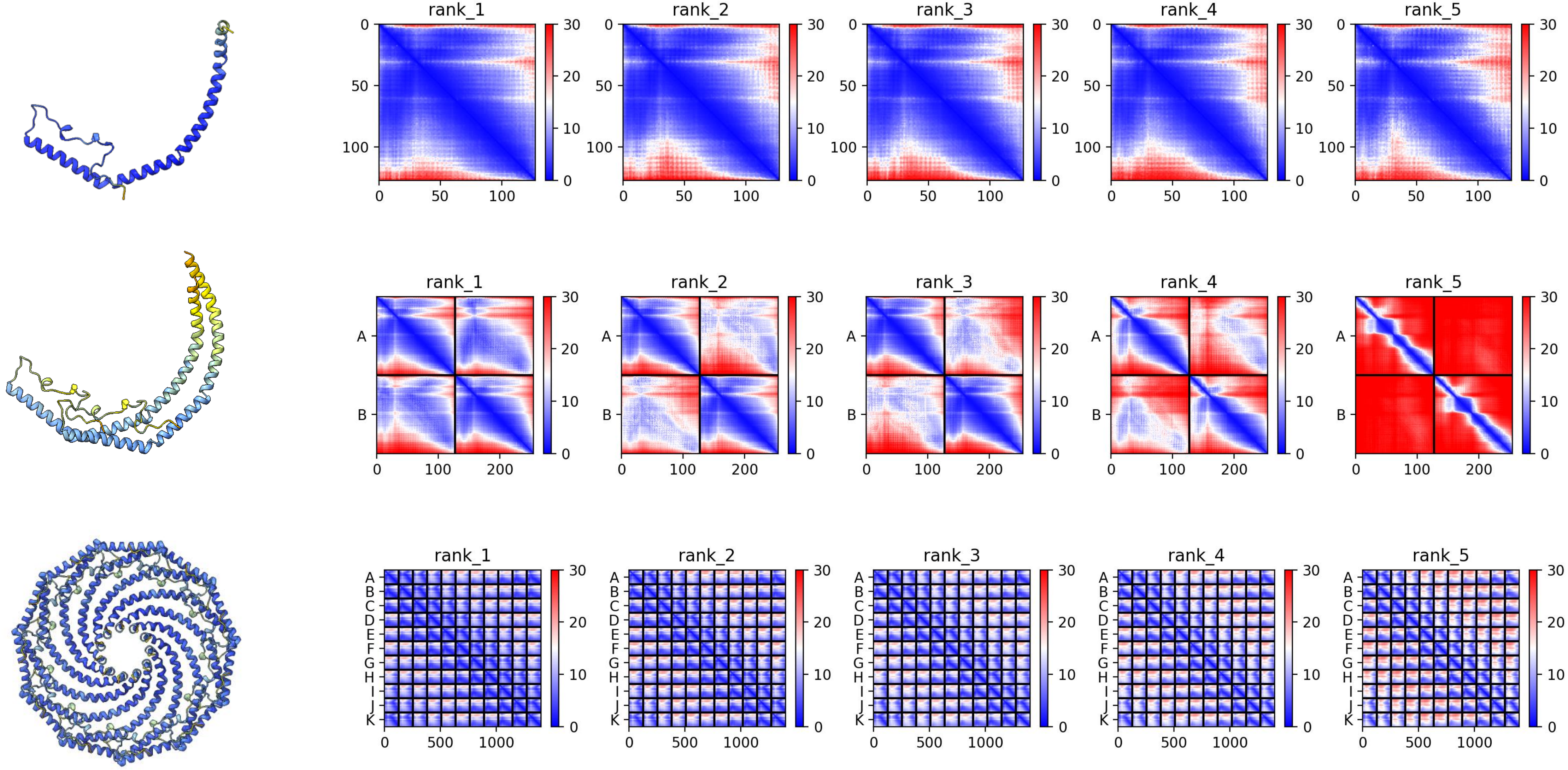

#### XP_050414695.1 *P. vulgata*

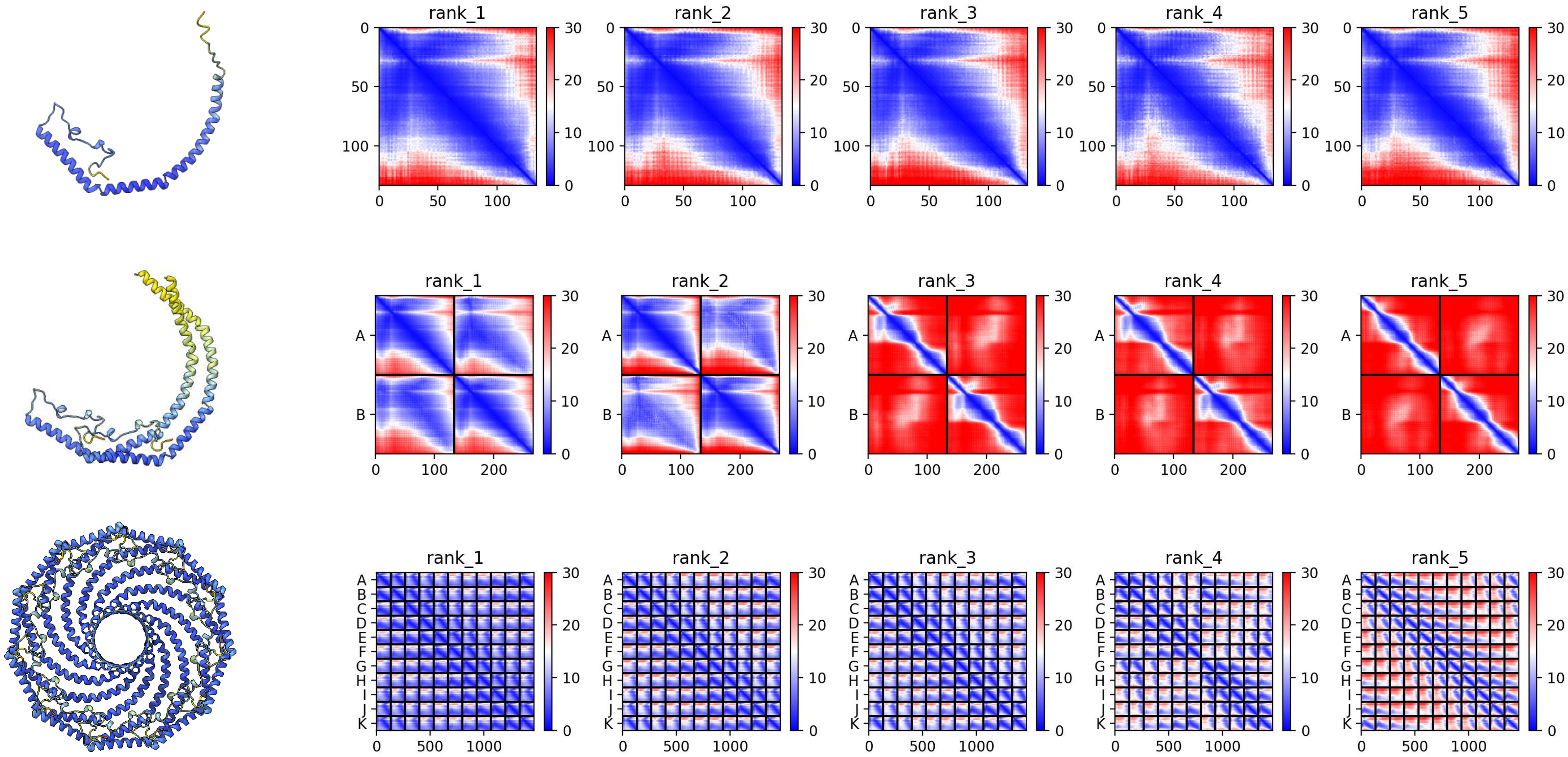

### XP_052778958.1 *M. arenaria*

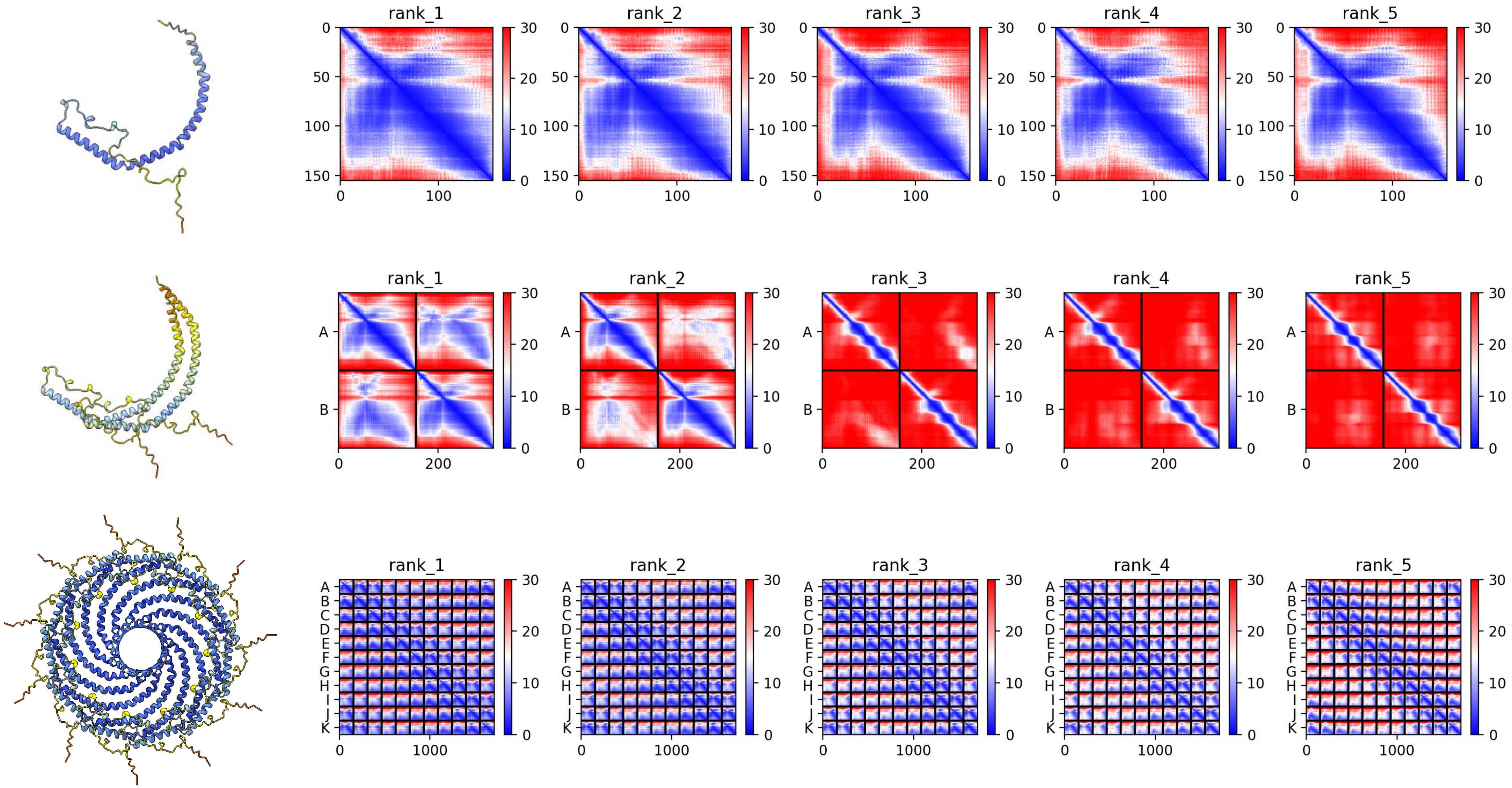

### XP_052780907.1 *M. arenaria*

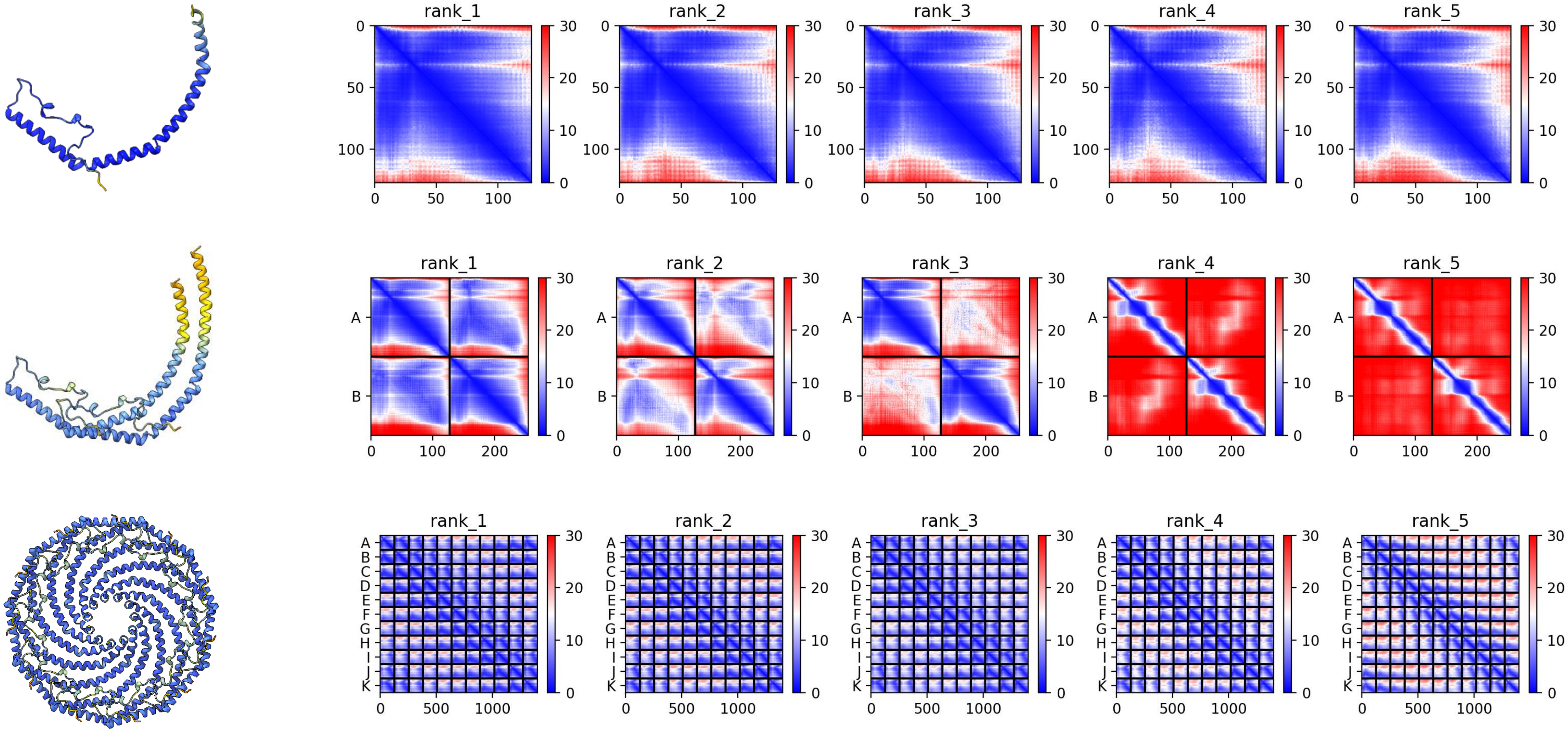

### XP_052815466.1 *M. arenaria*

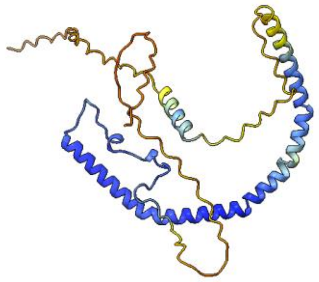

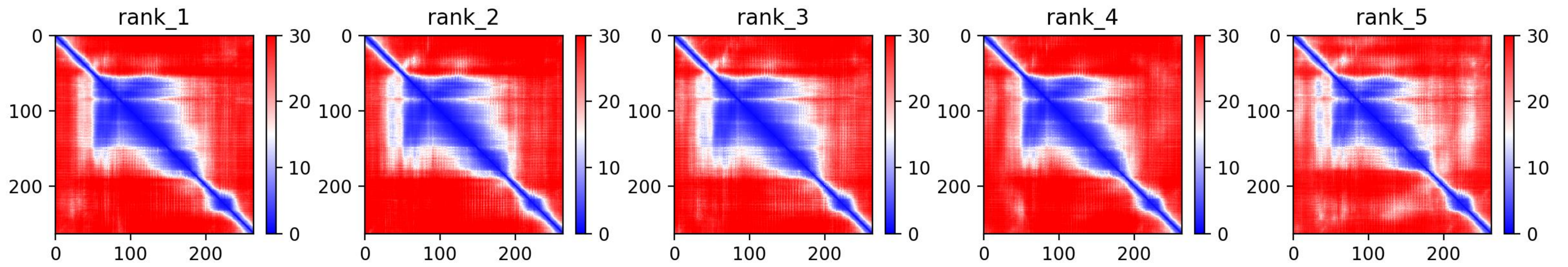


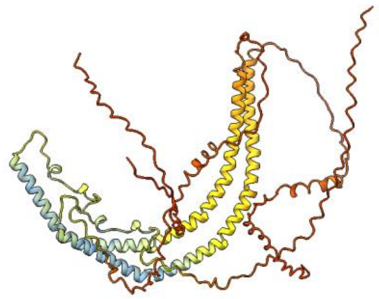

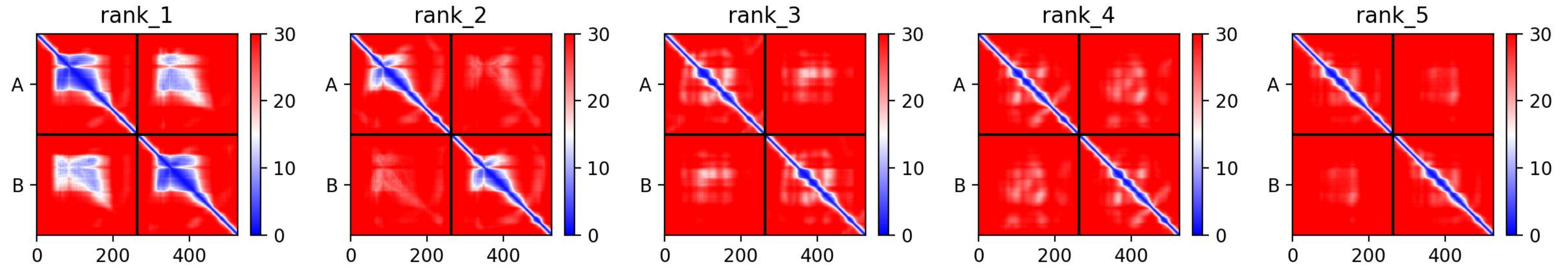


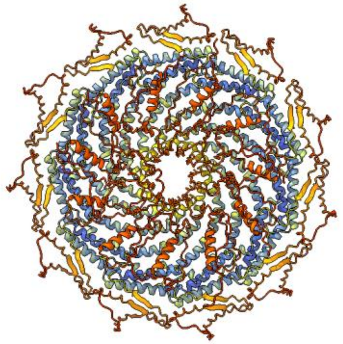

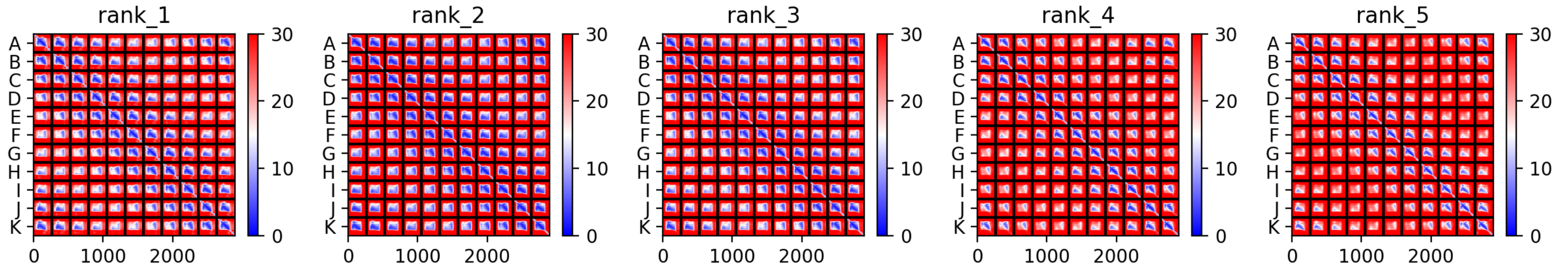

### XP_052820175.1 *M. arenaria*

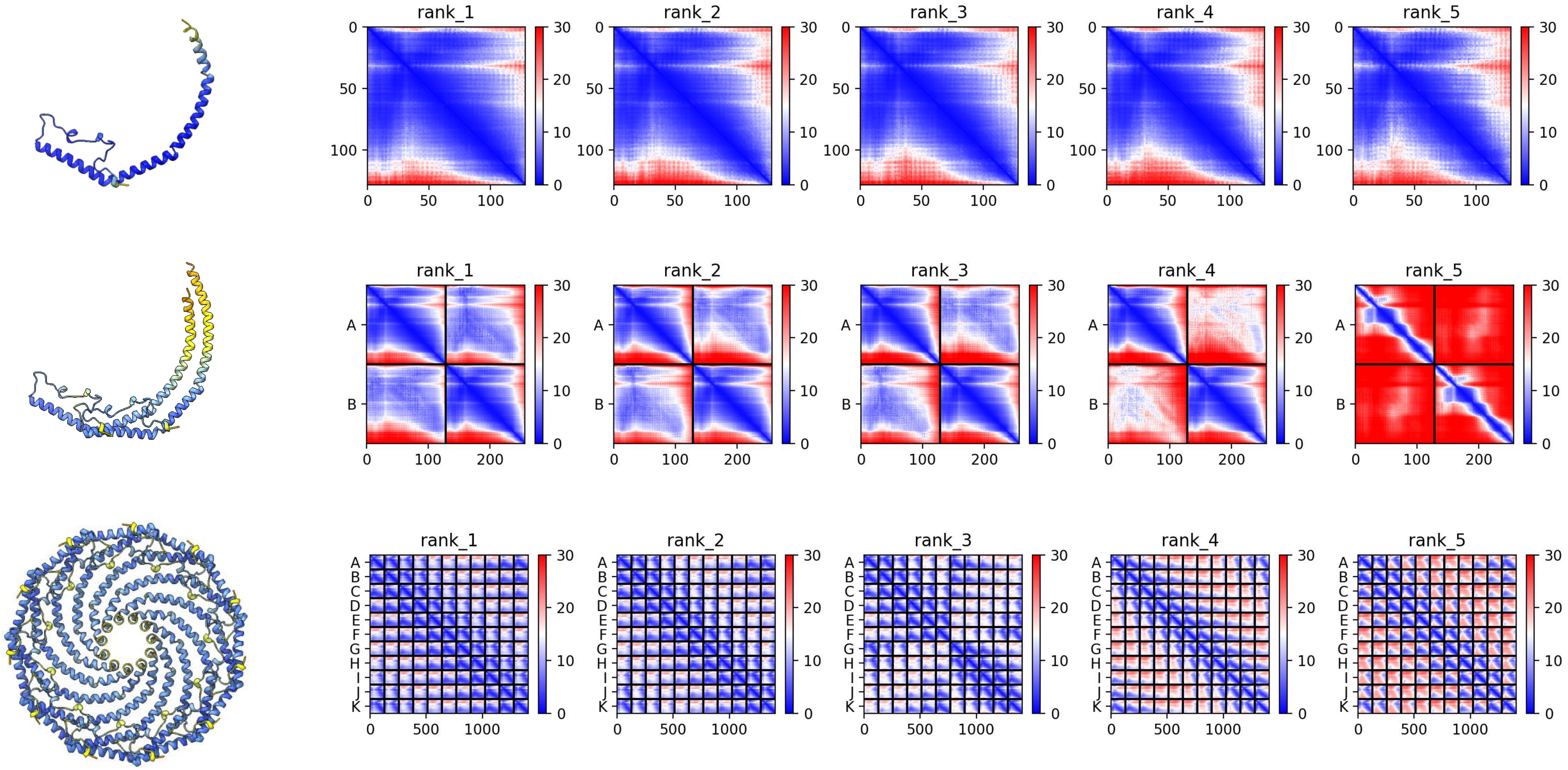

### XP_052820174.1 *M. arenaria*

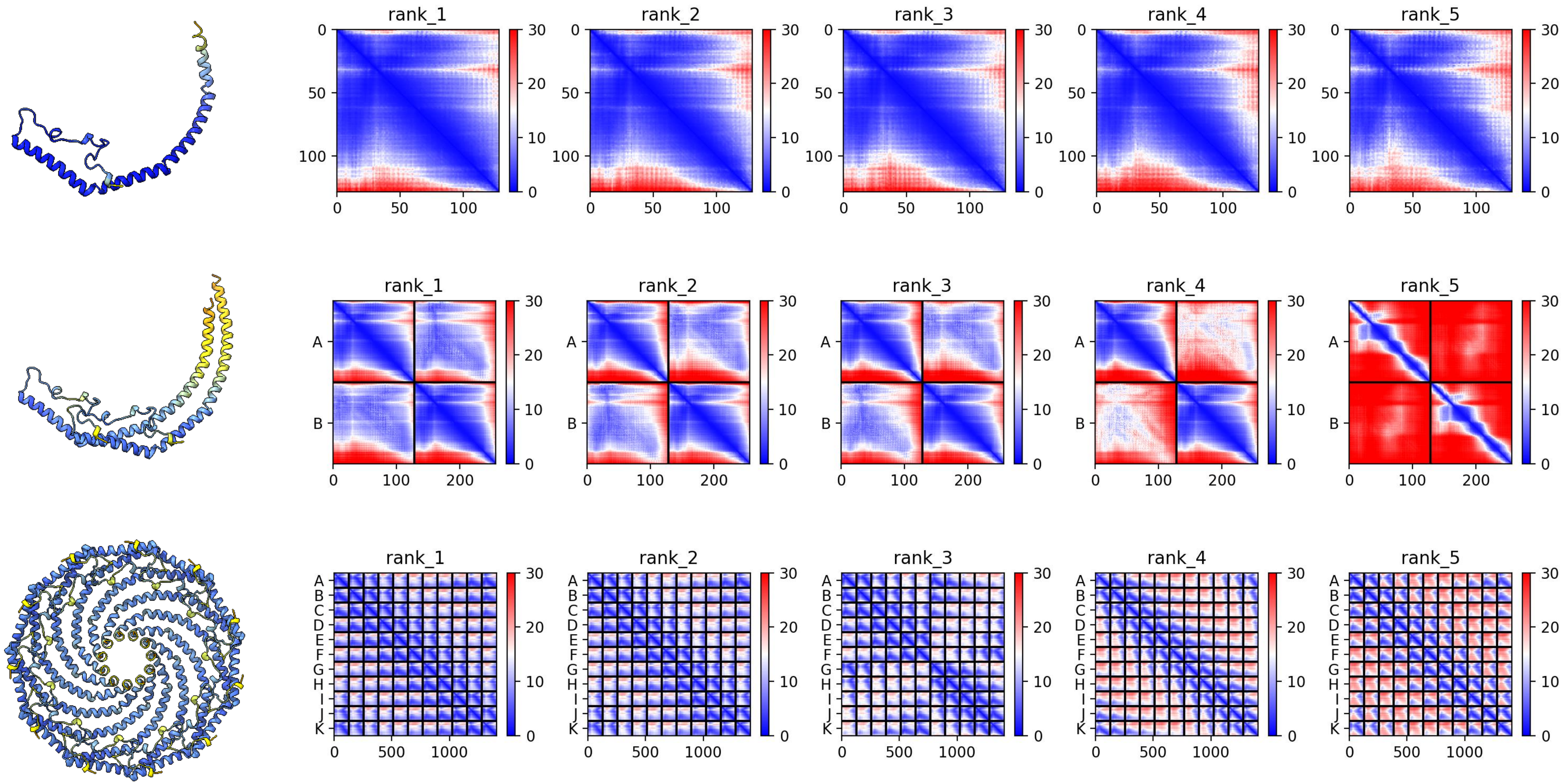

#### XP_052788808.1 *M. arenaria*

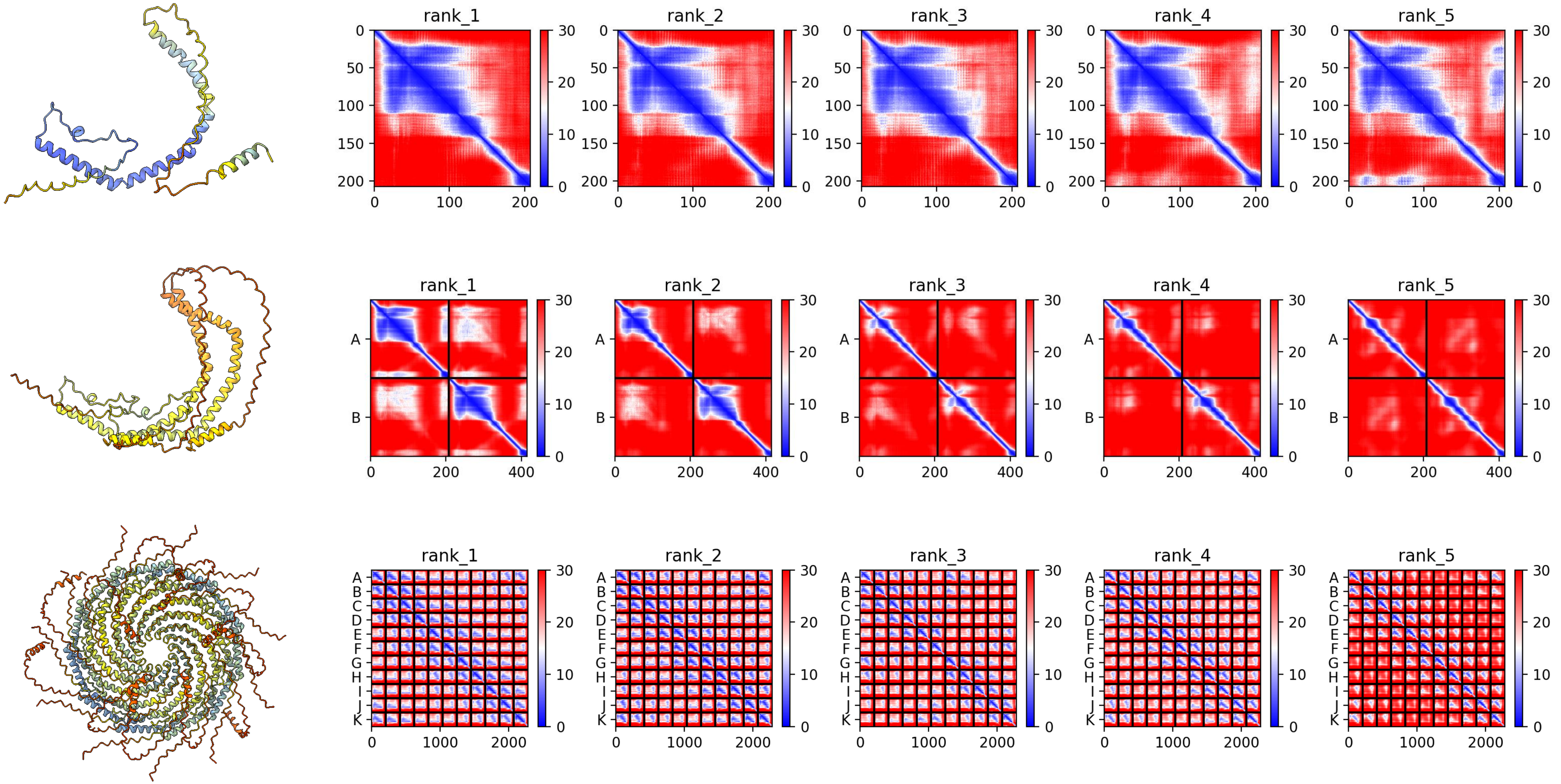

### XP_052784964.1 *M. arenaria*

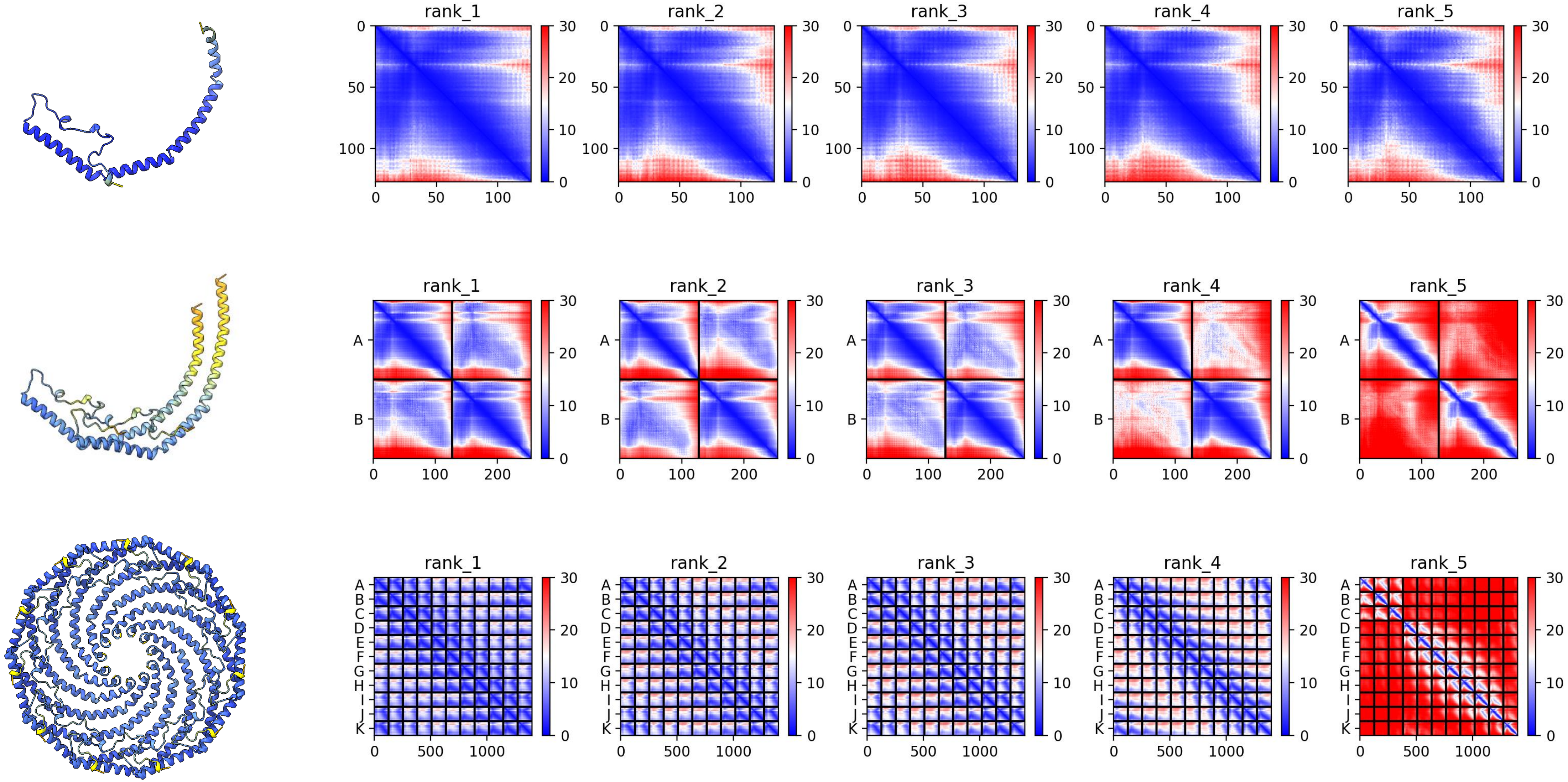

### XP_053396032.1 *M. mercenaria*

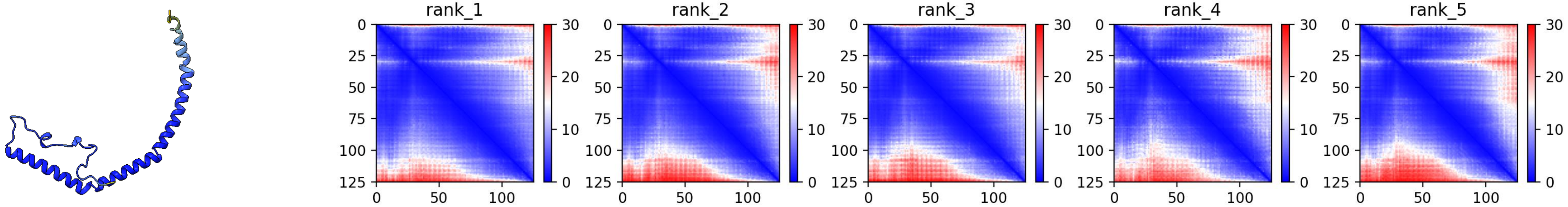


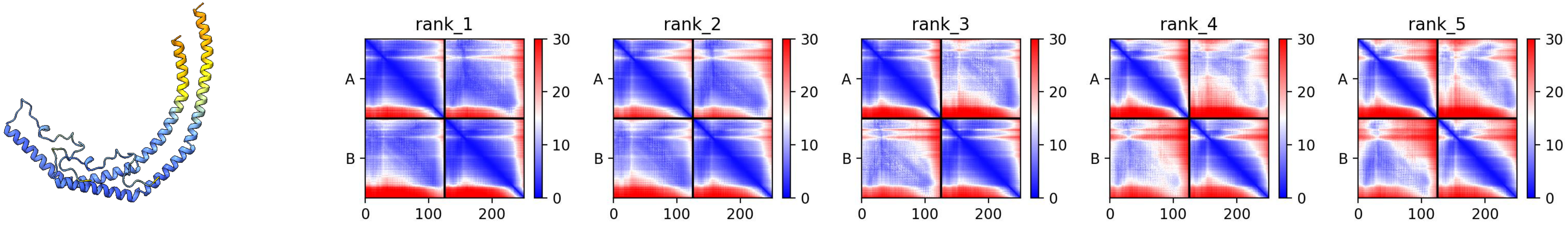


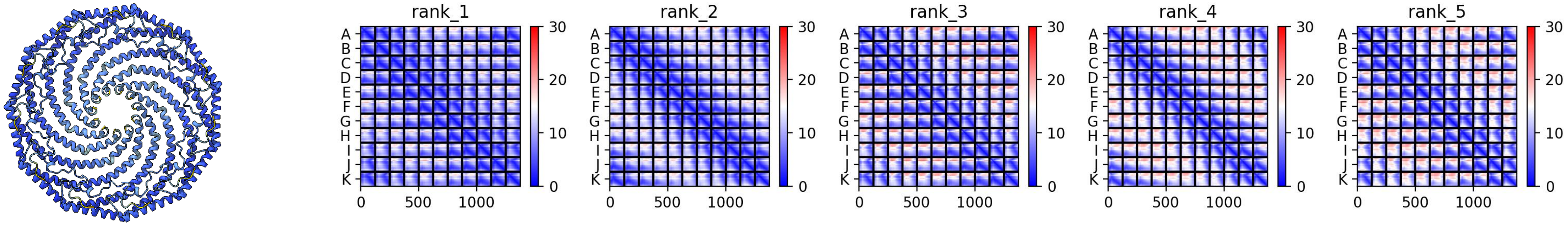

### XP_045164732.2 *M. mercenaria*

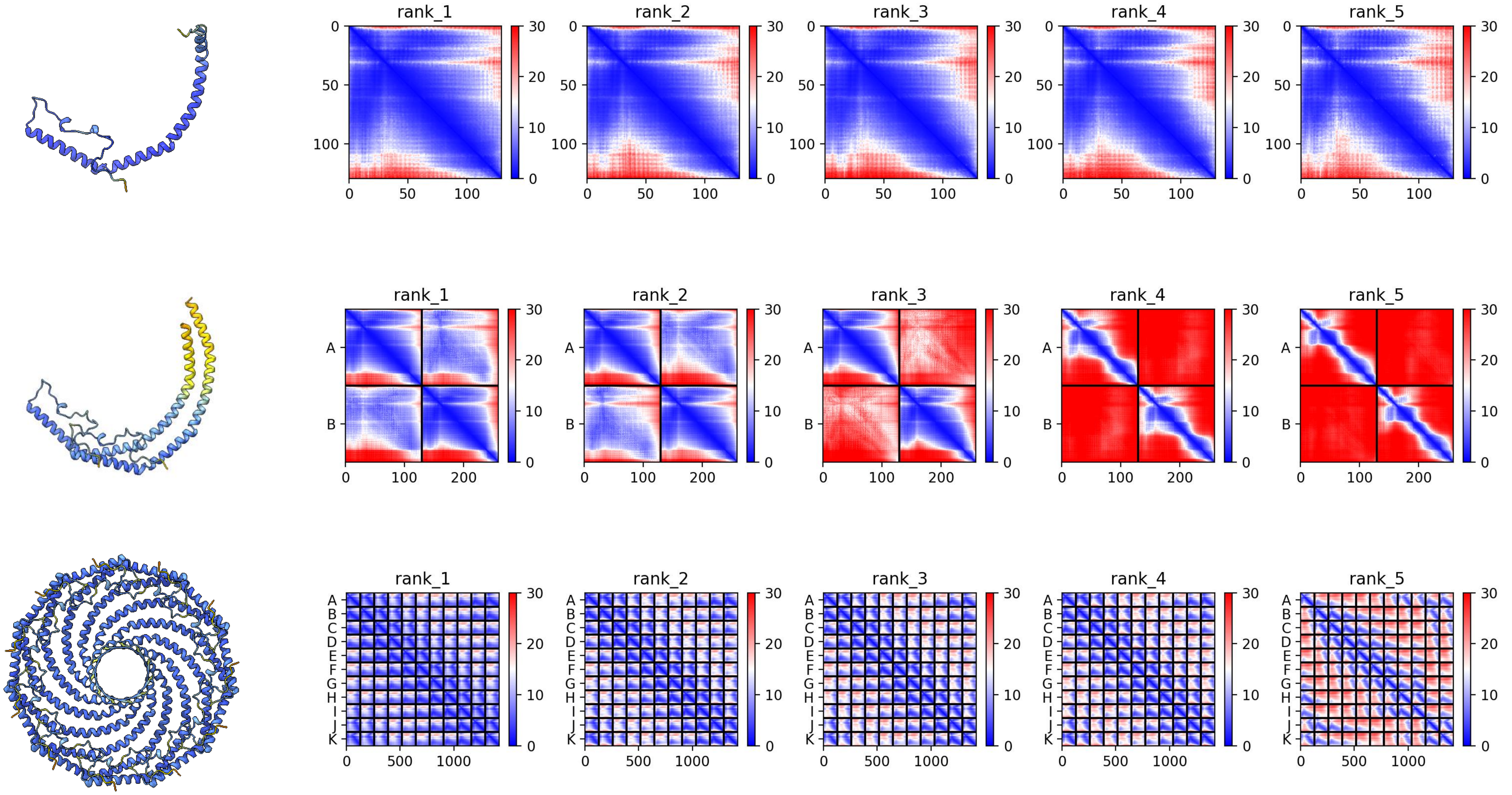

### XP_053377544.1 *M. mercenaria*

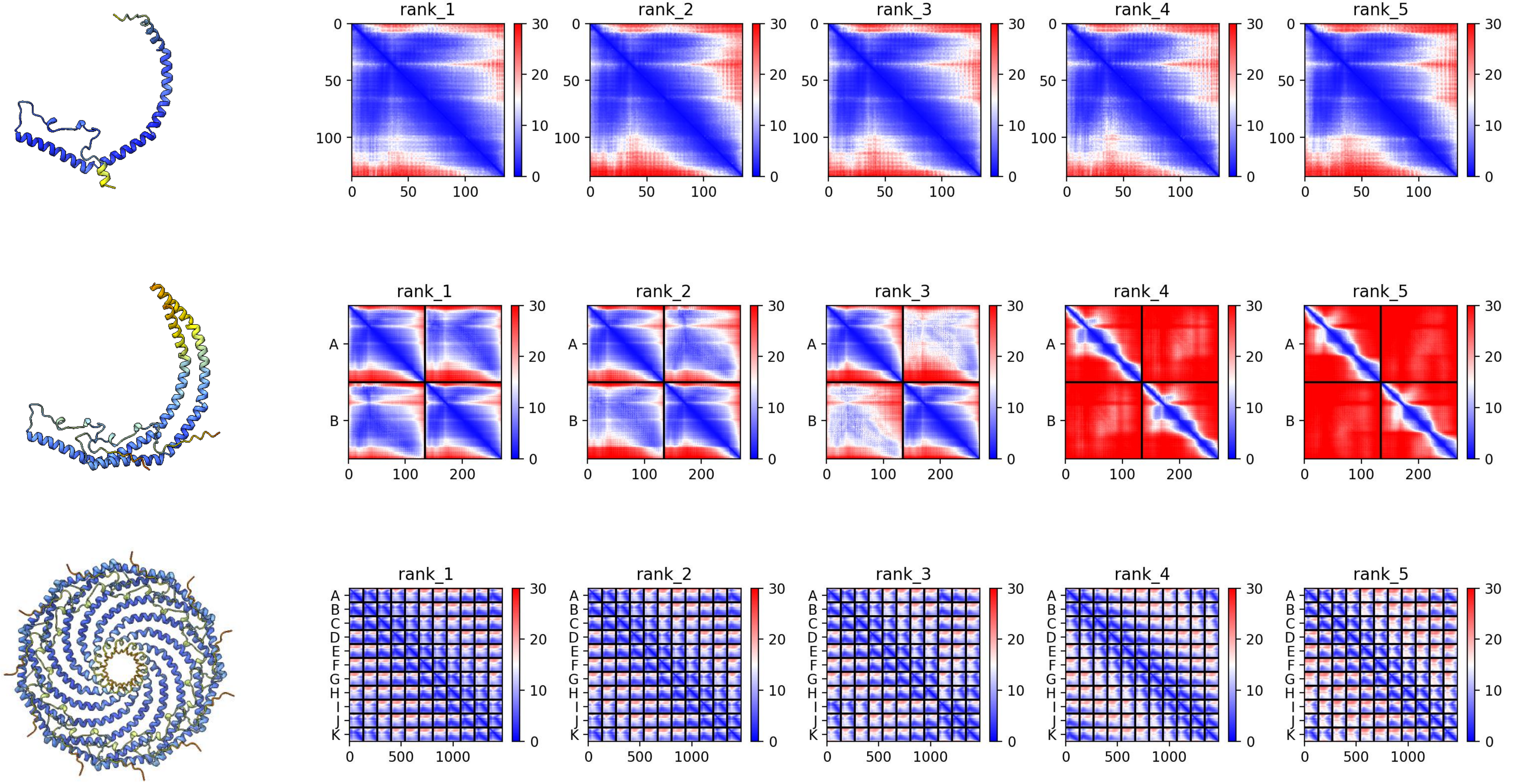

### XP_045174338.2 *M. mercenaria*

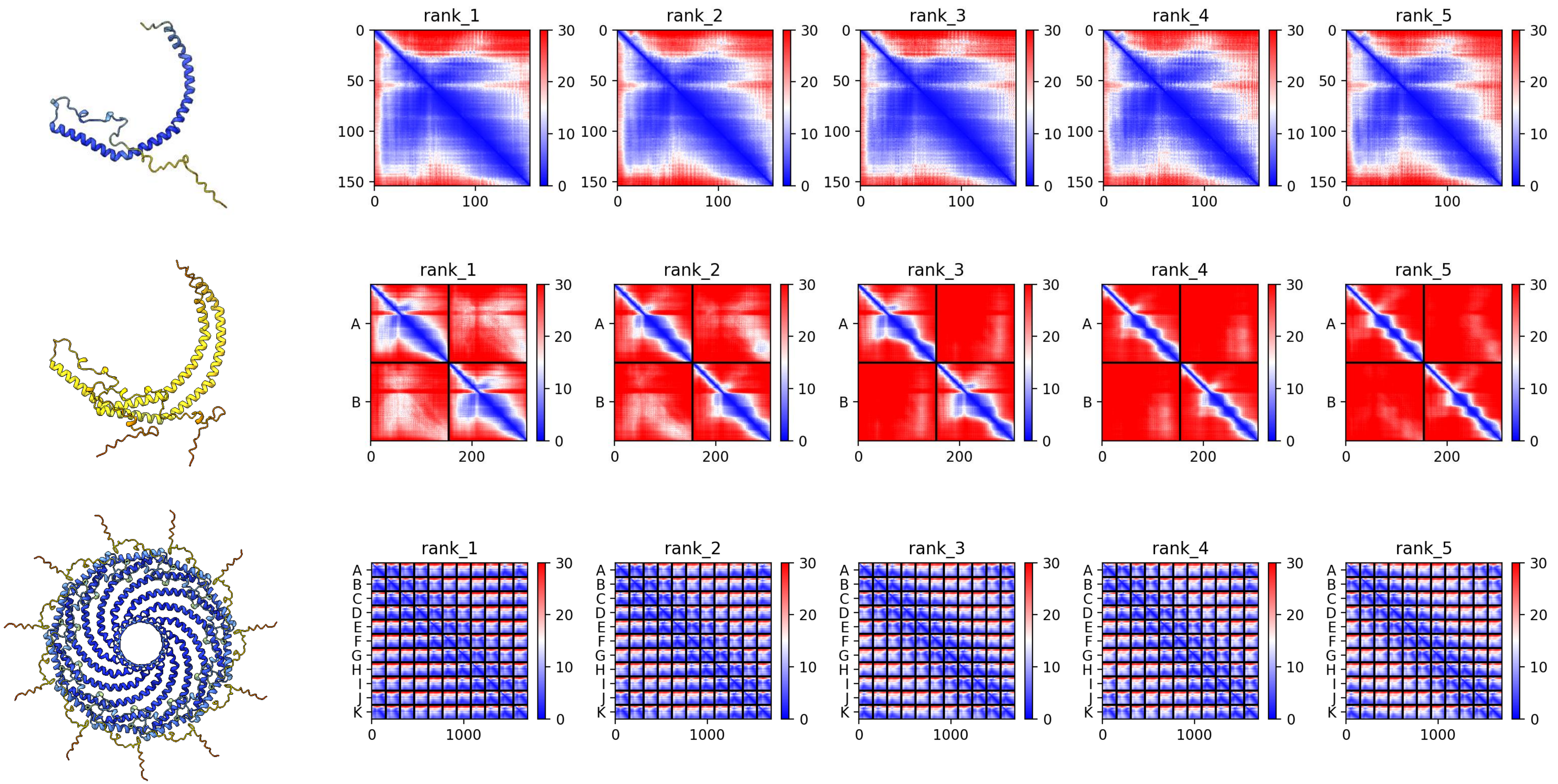

### XP_033630427.1 *A. rubens*

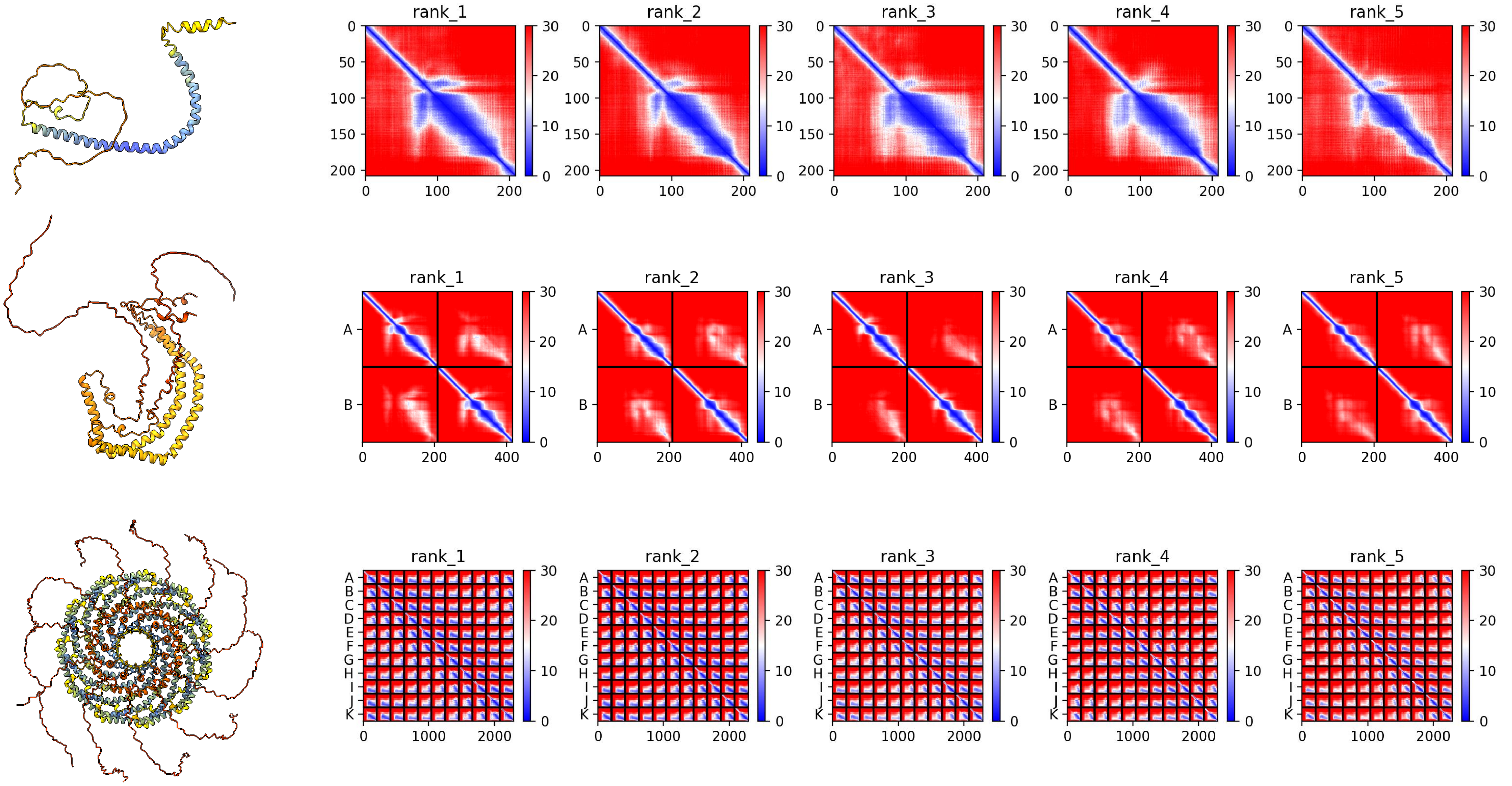

### XP_033636910.1 *A. rubens*

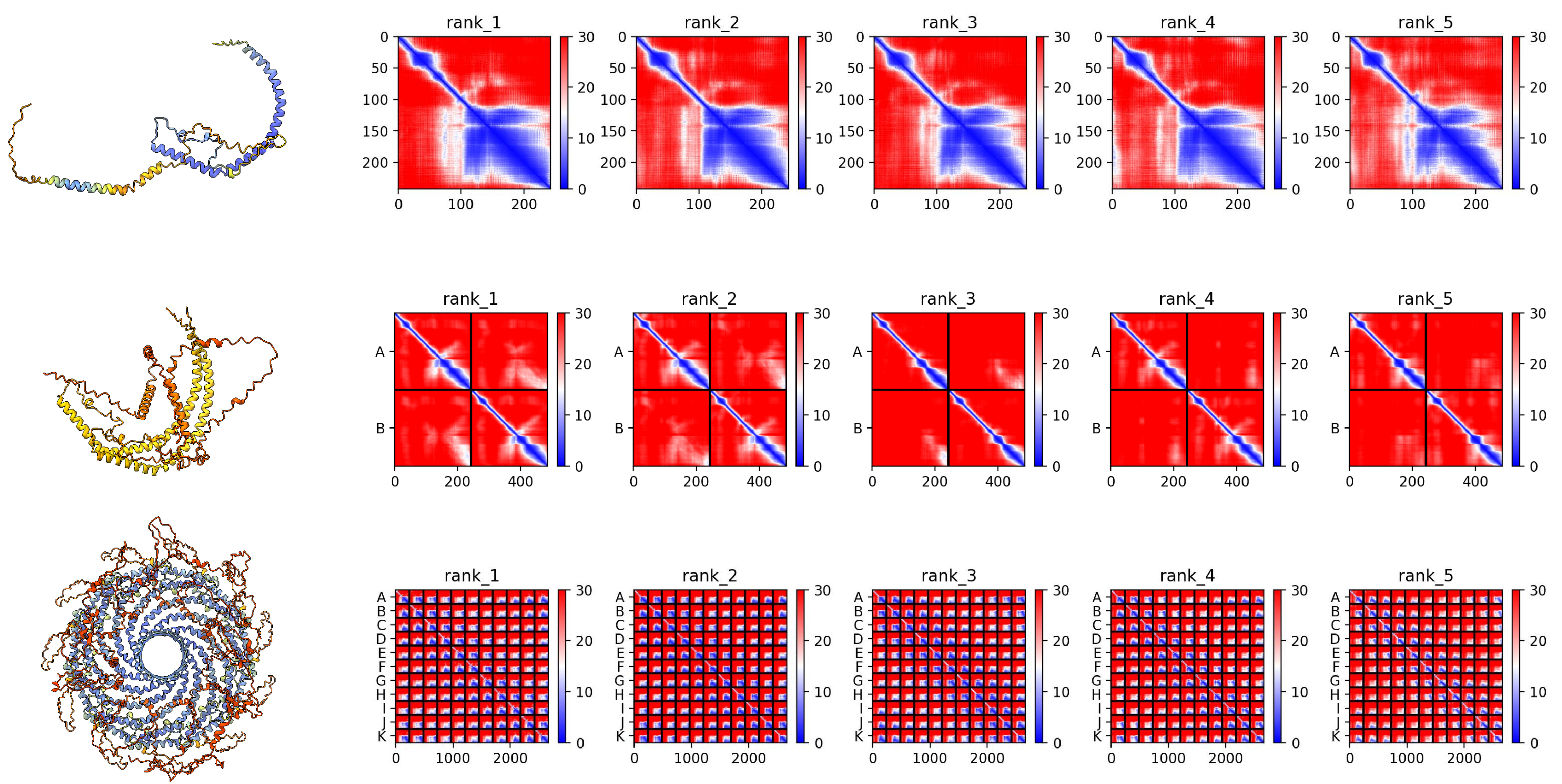

### XP_041476235.1 *L. variegatus*

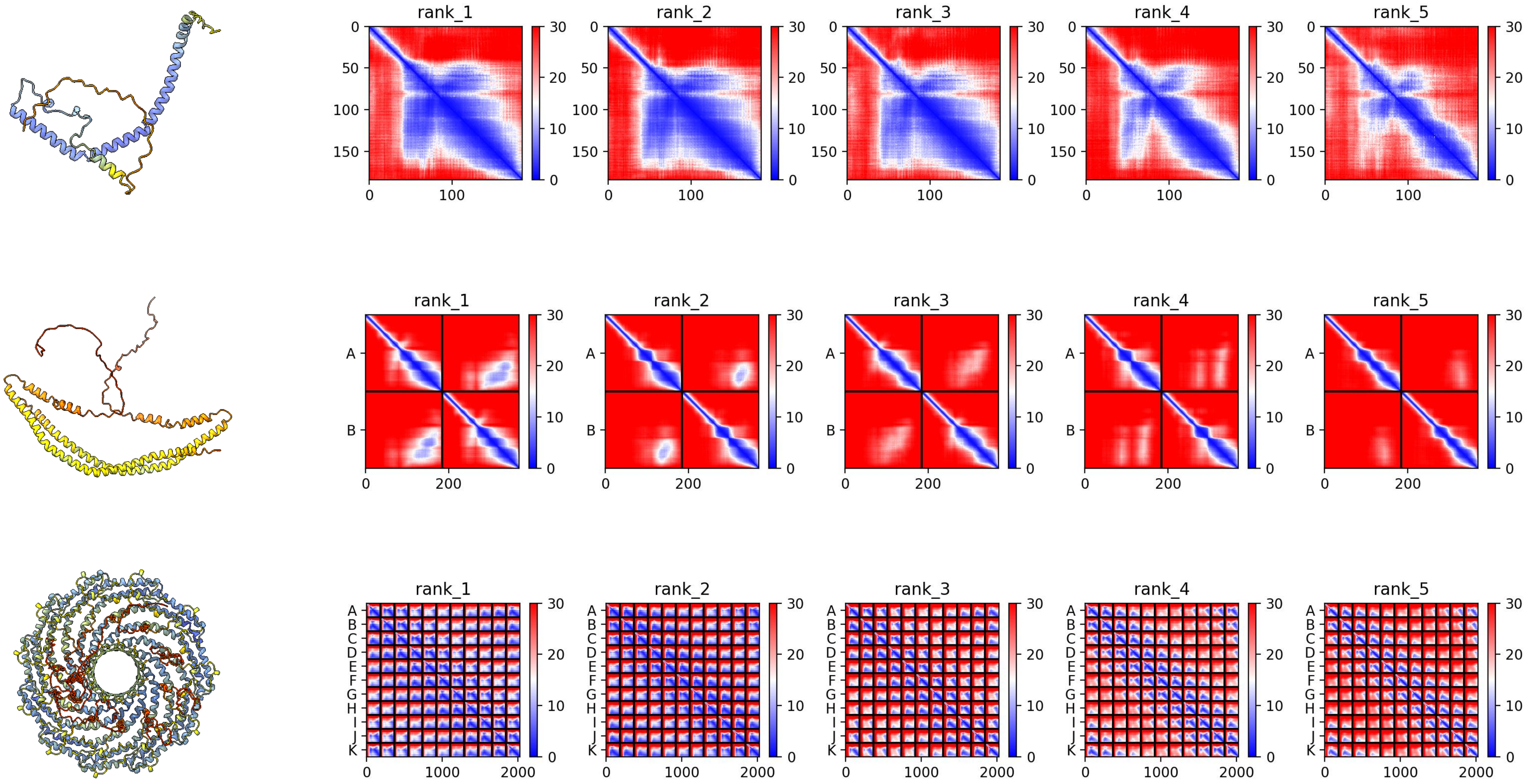

### *XP_041469030.1 L. variegatus*

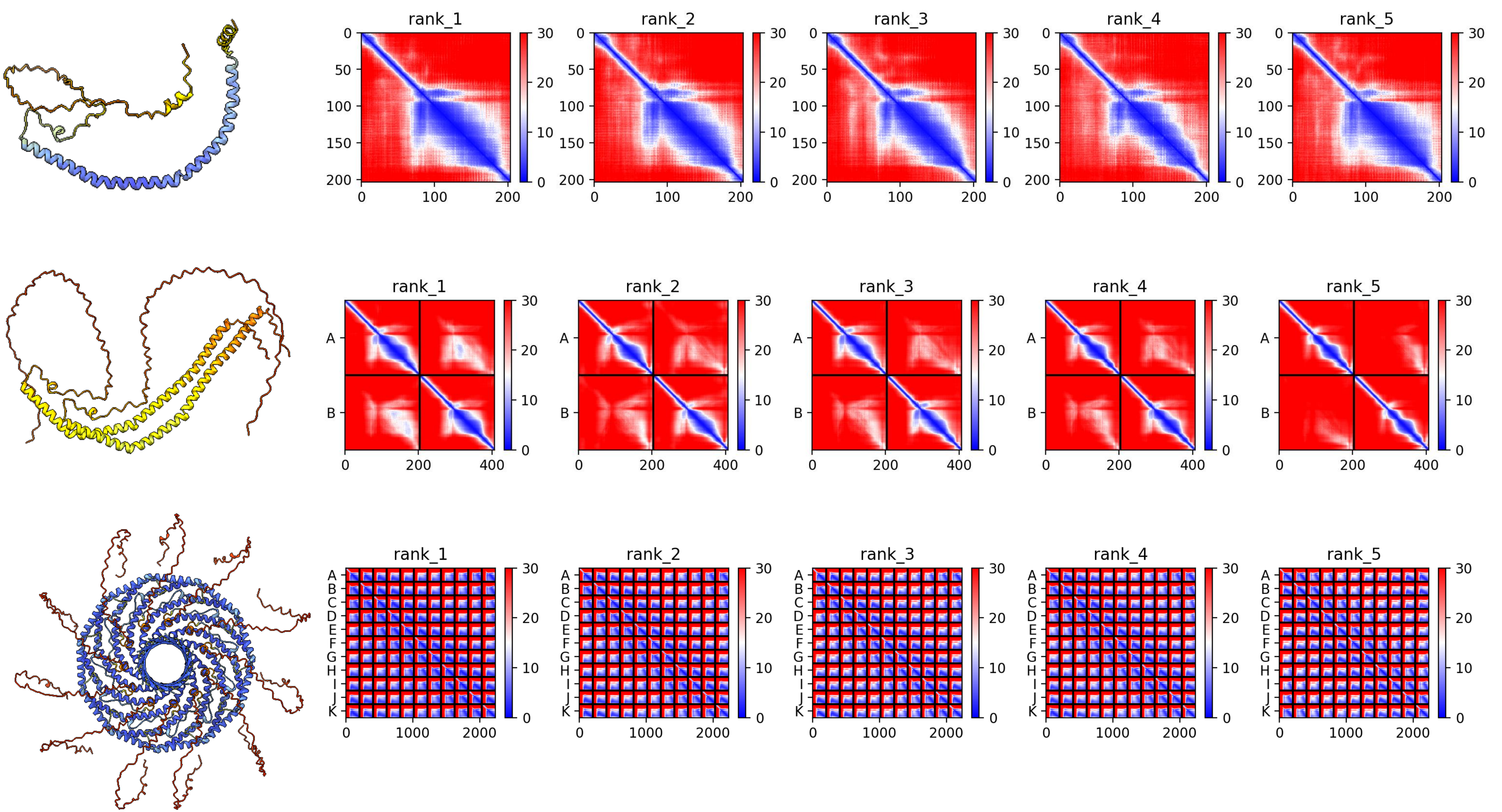

***XP_041454781.1 L. variegatus***

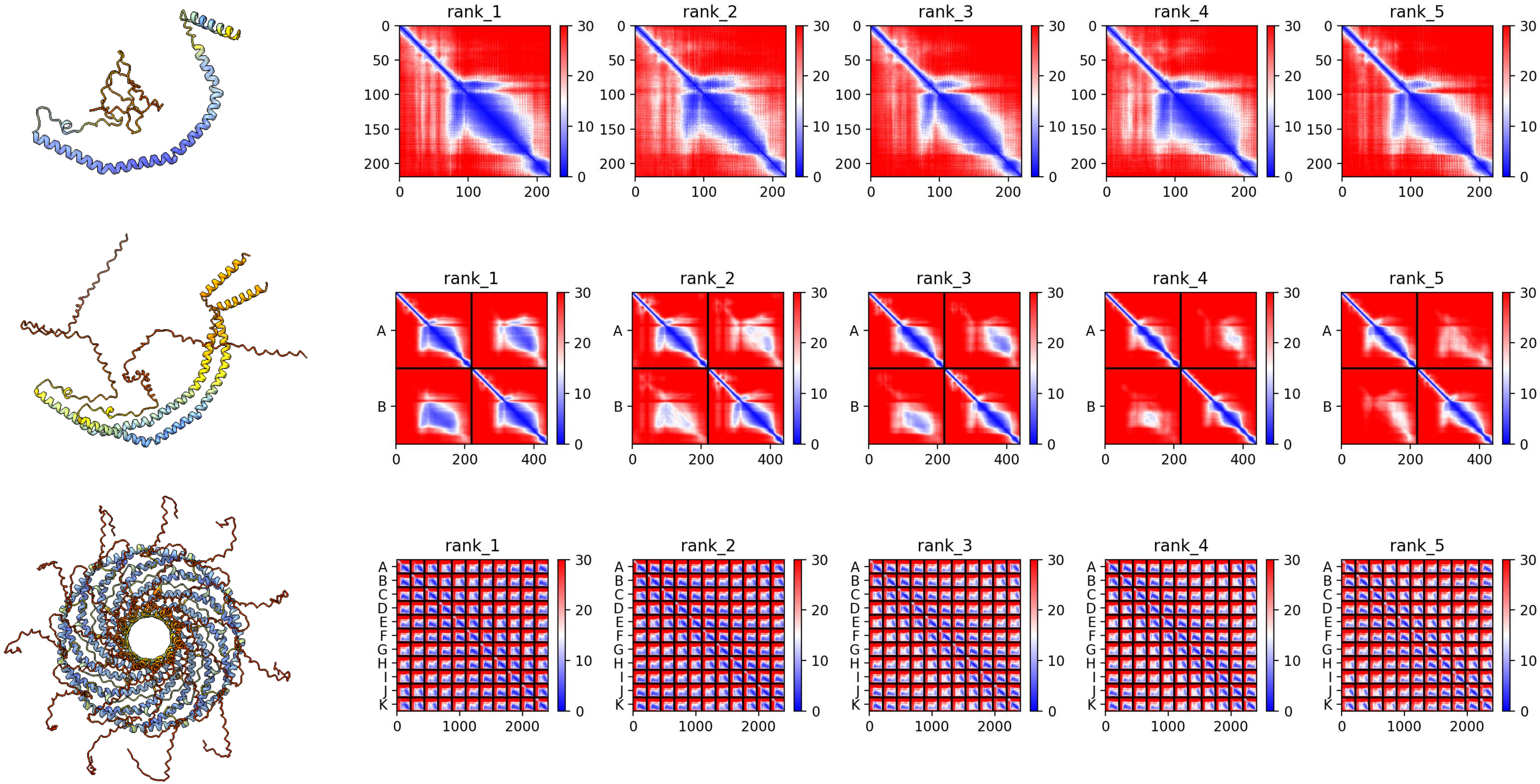

### *XP_041460678.1 L. variegatus*

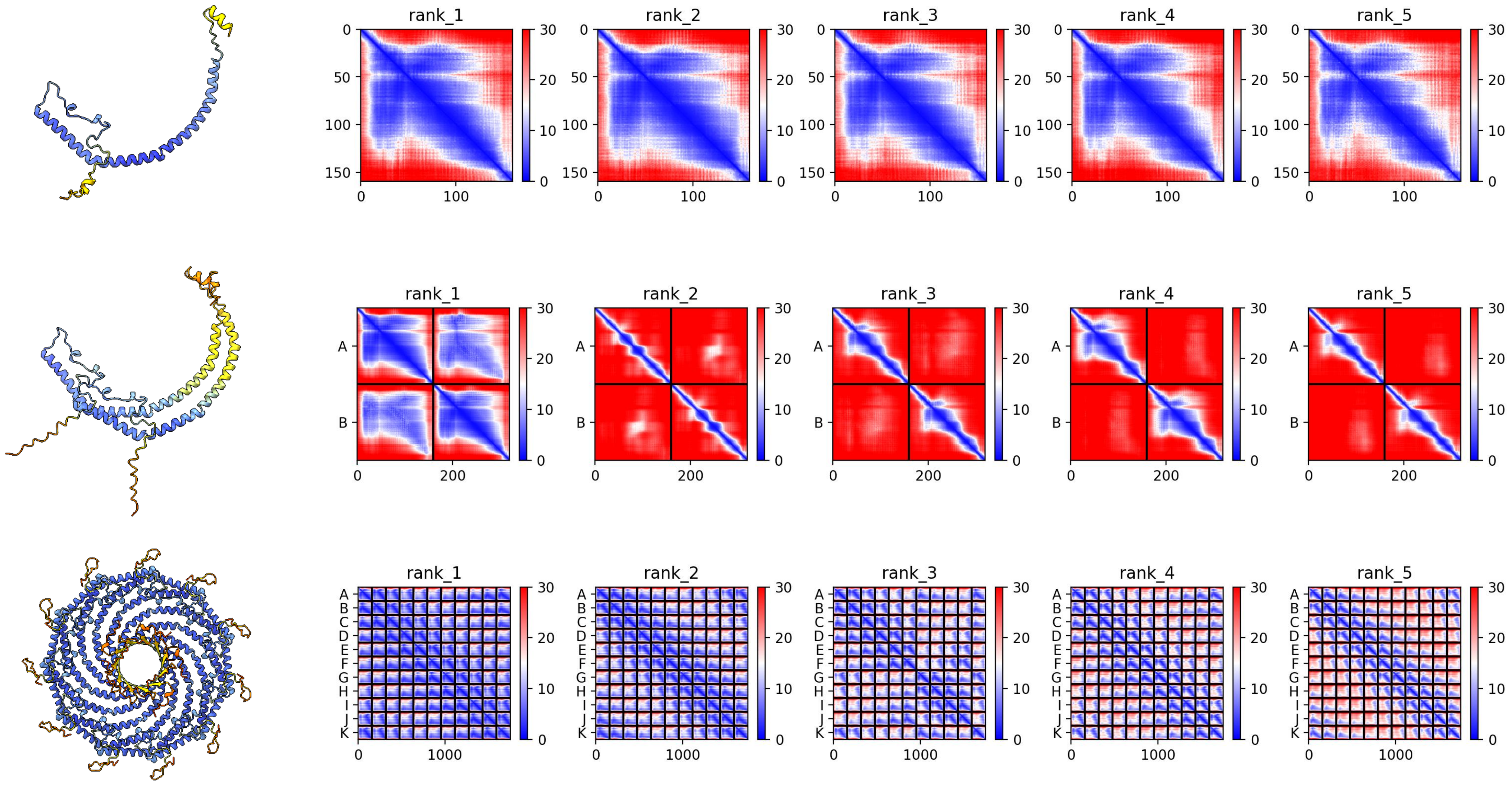

### *XP_054764459.1 L. pictus*

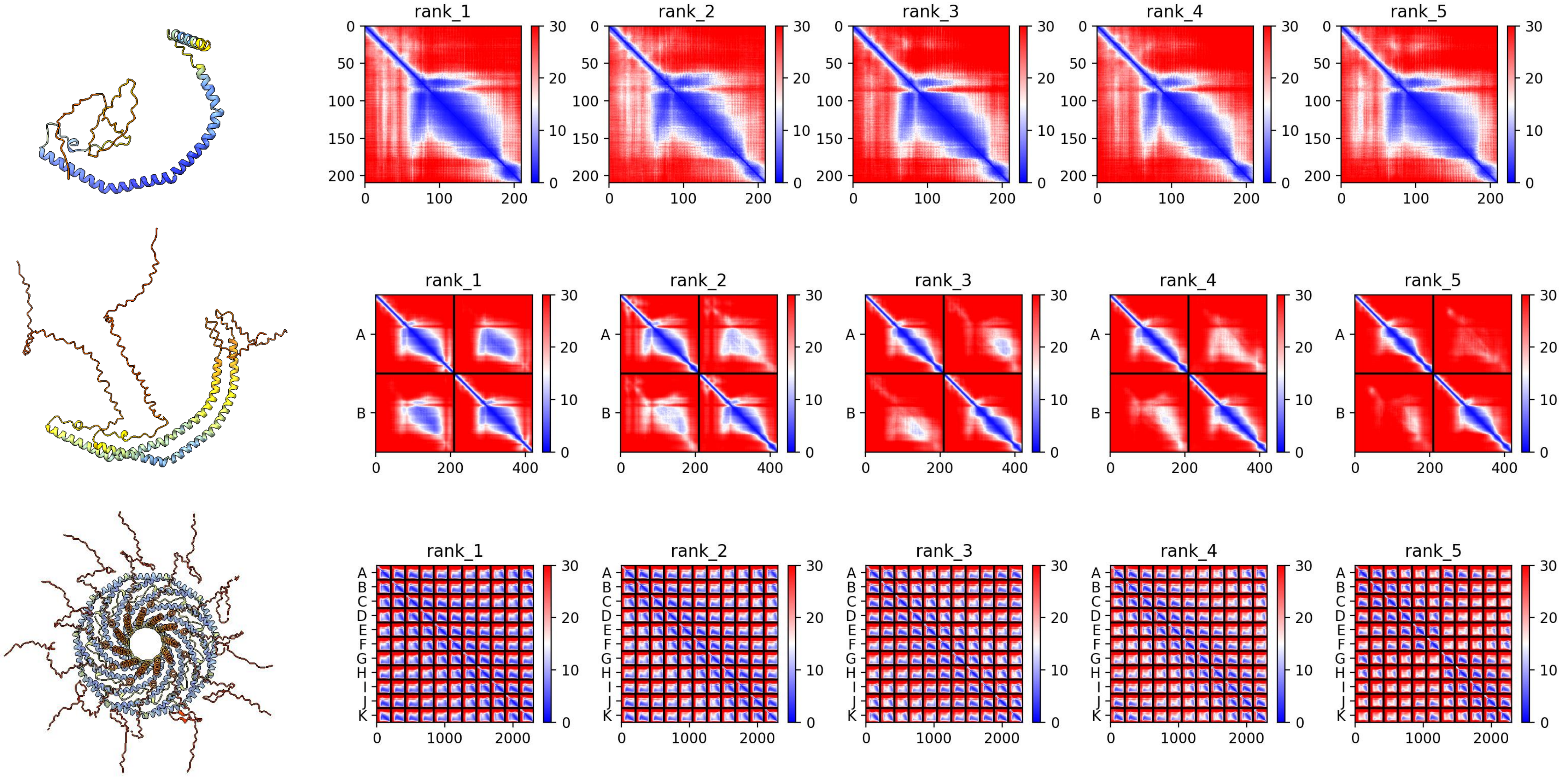

### *XP_054752263.1 L. pictus*

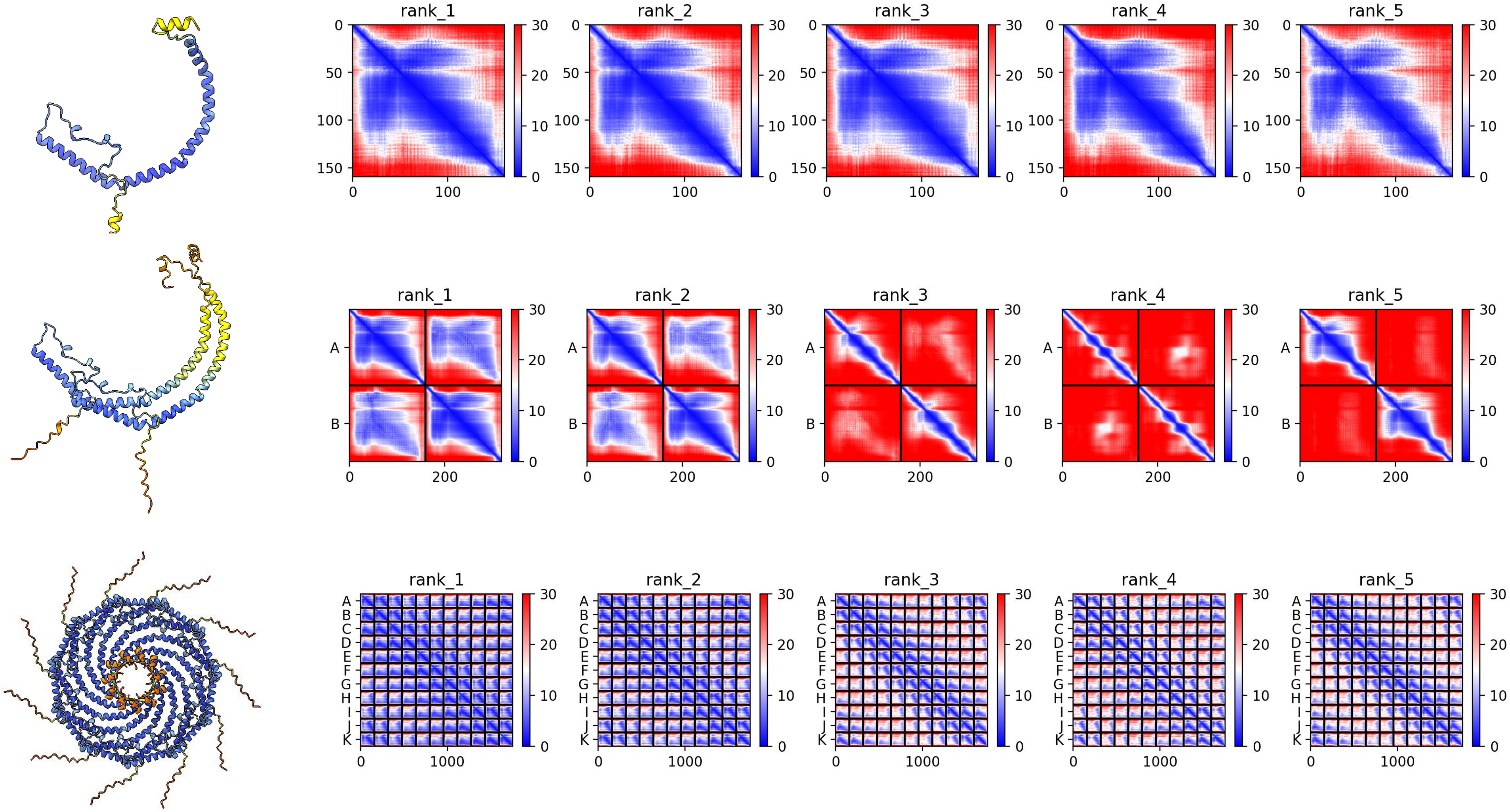

### *XP_054769514.1 L. pictus*

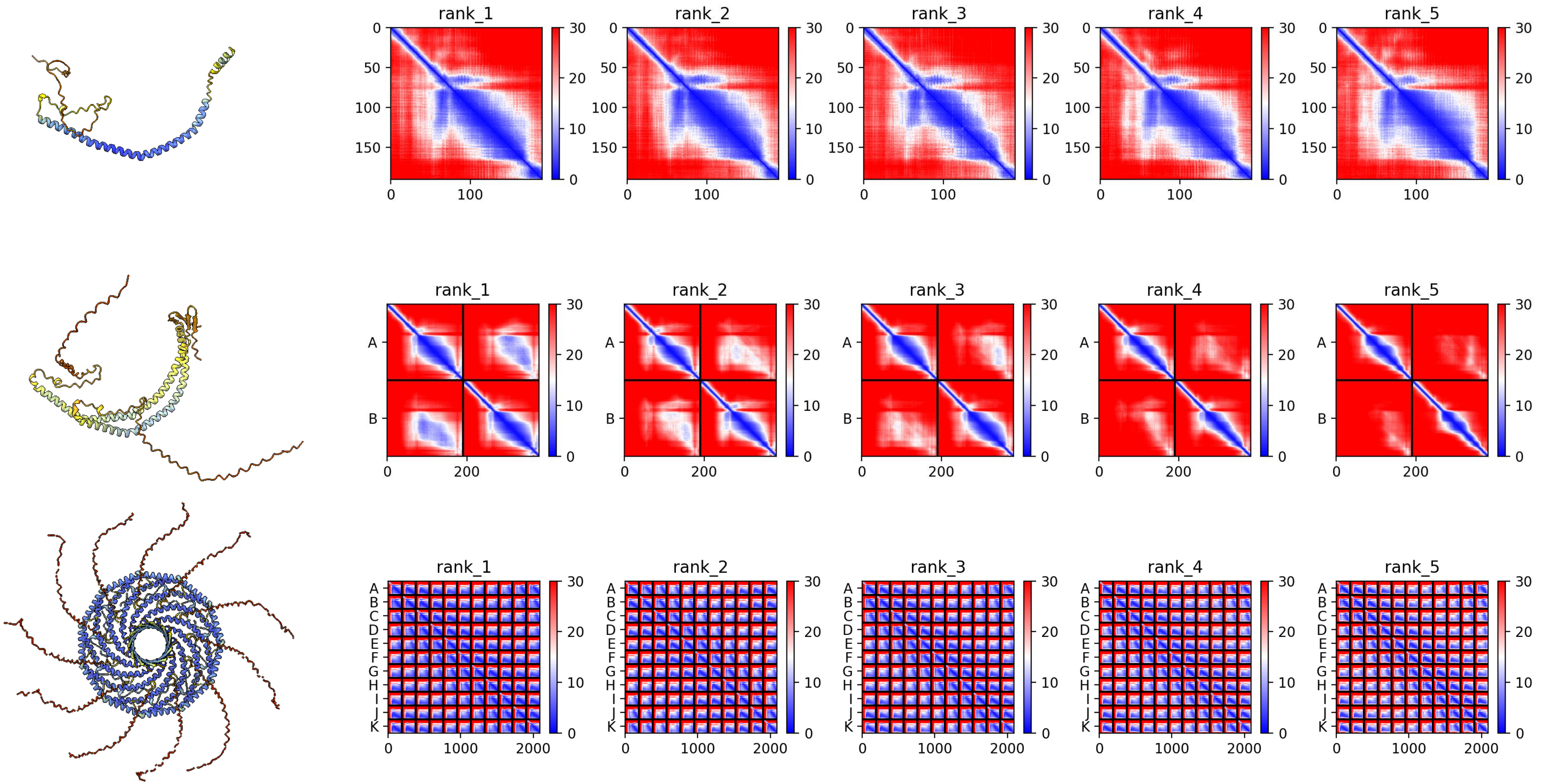

#### *XP_054764941.1 L. pictus*

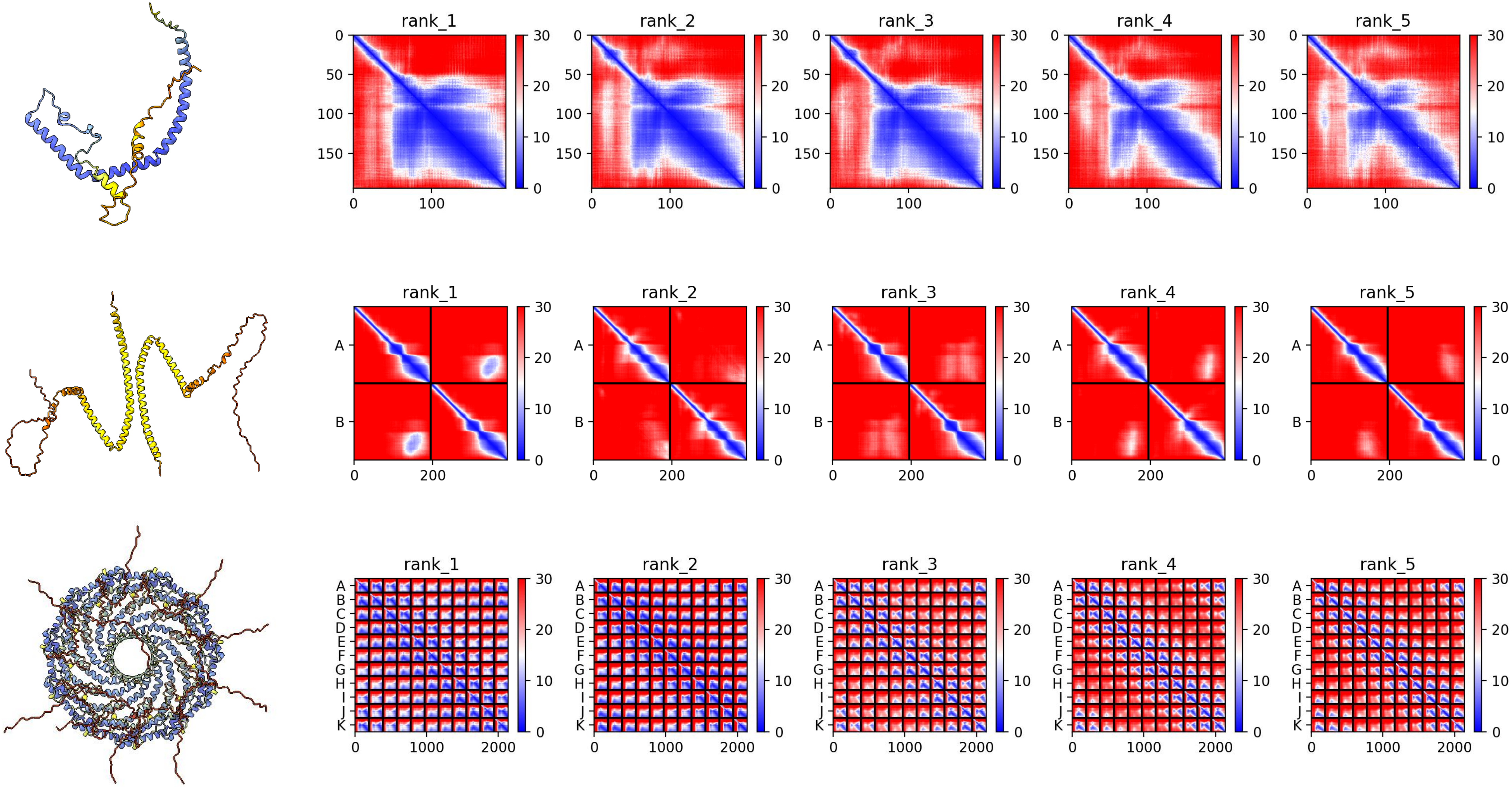

### *KAJ8040503.1 H. leucospilota*

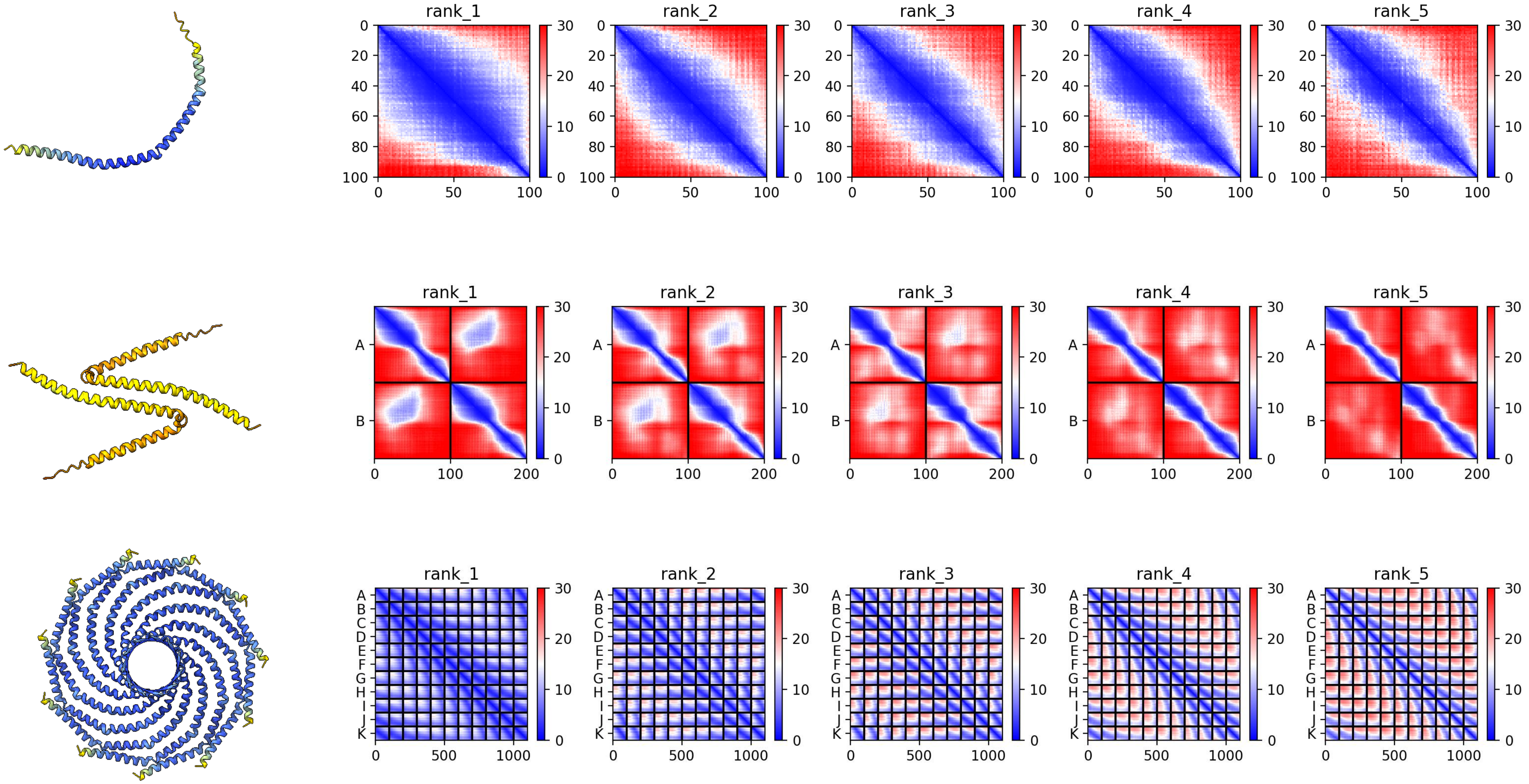

**KAJ8046320.1 *H. leucospilota***

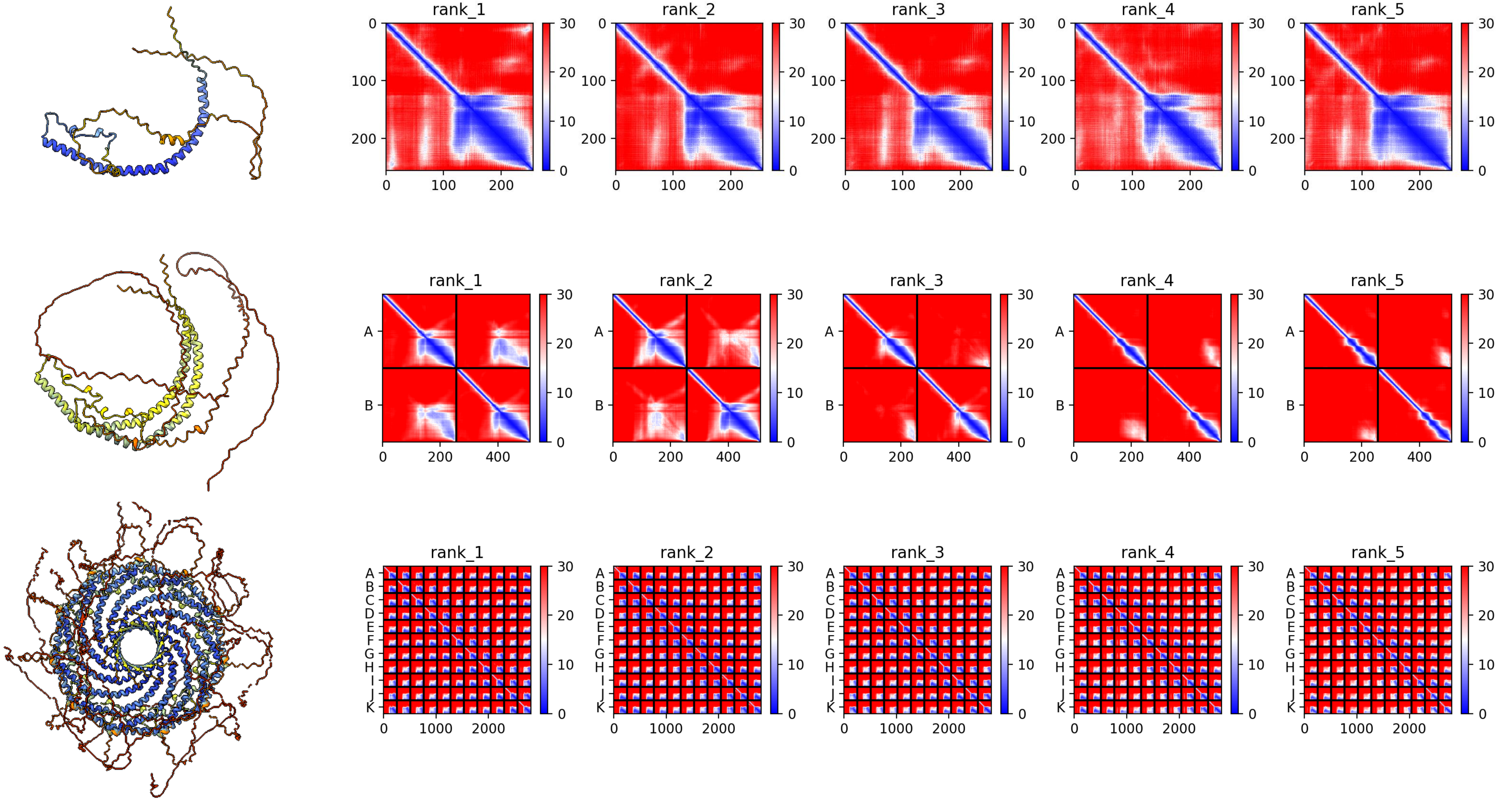

### KAJ8022872.1 *H. leucospilota*

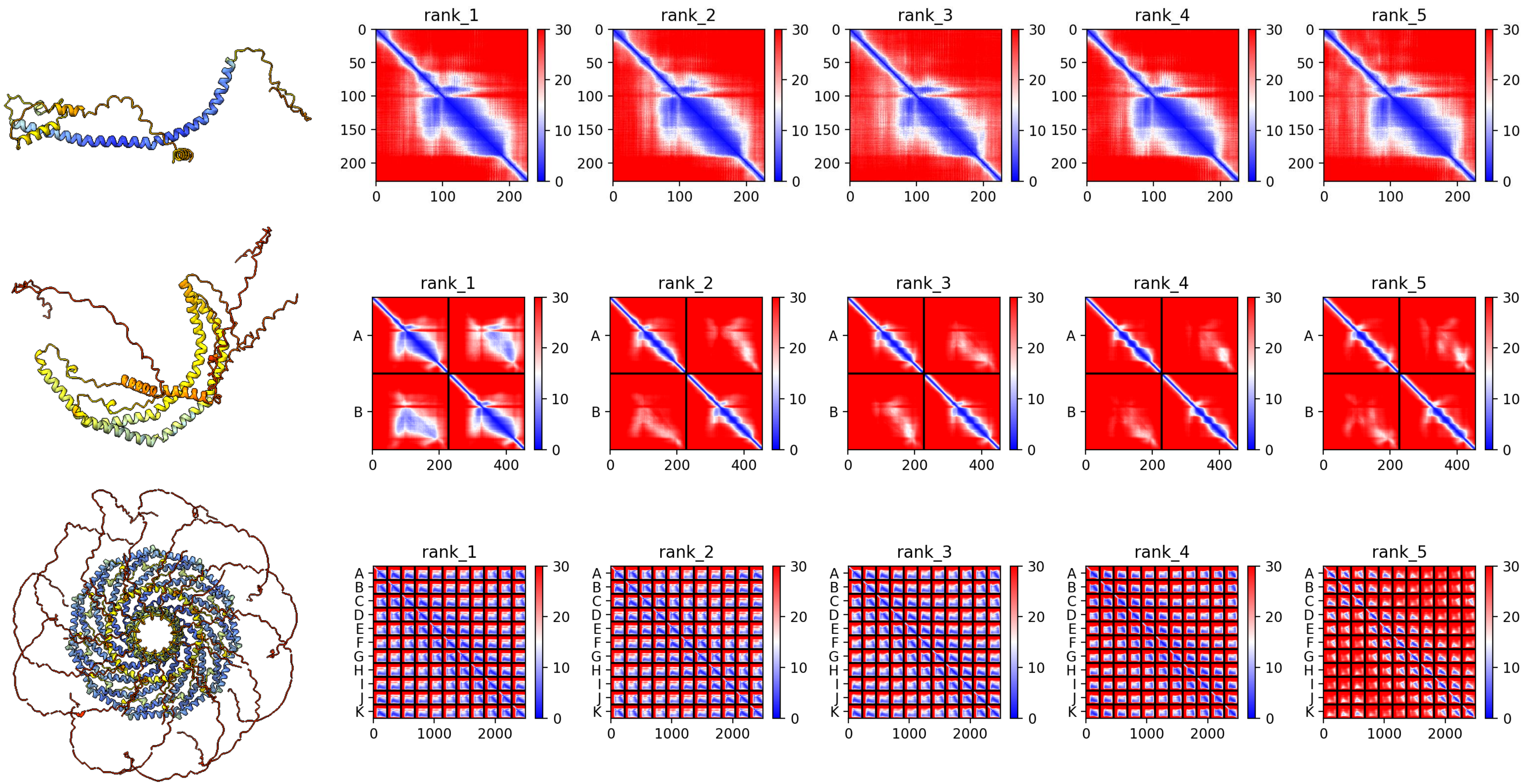

### KAJ8020606.1 *H. leucospilota*

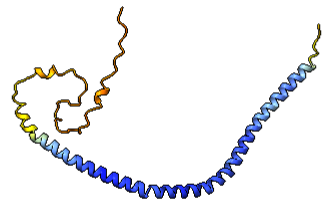

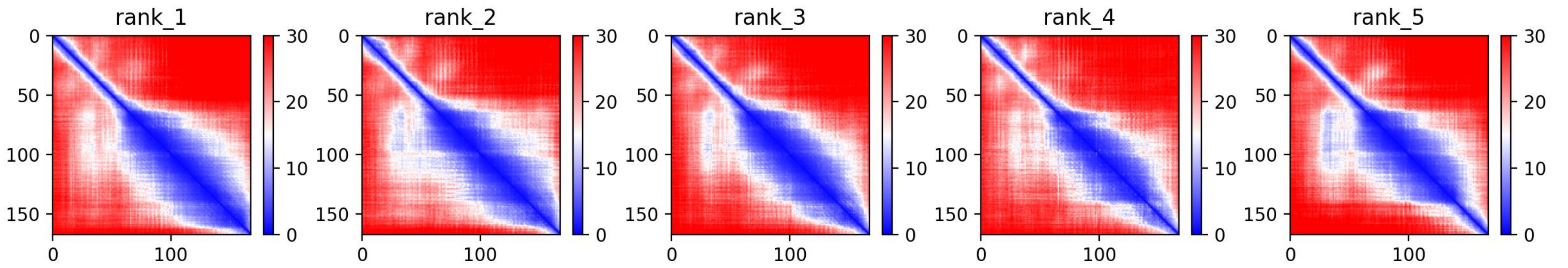


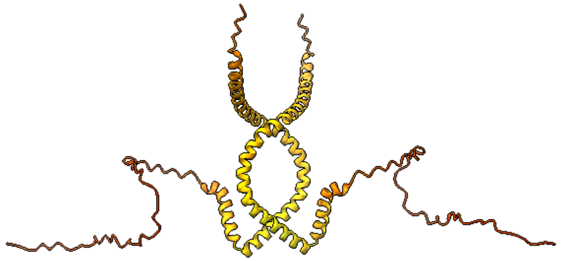

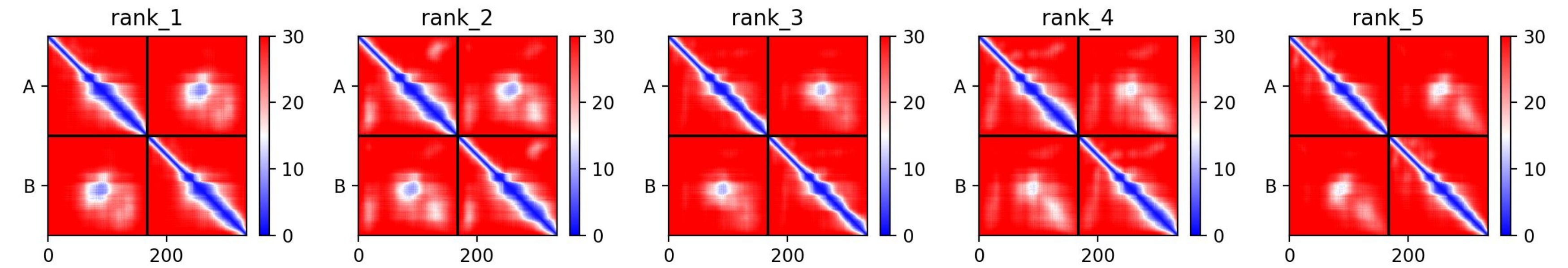


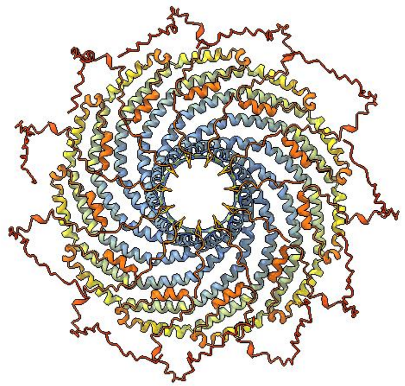

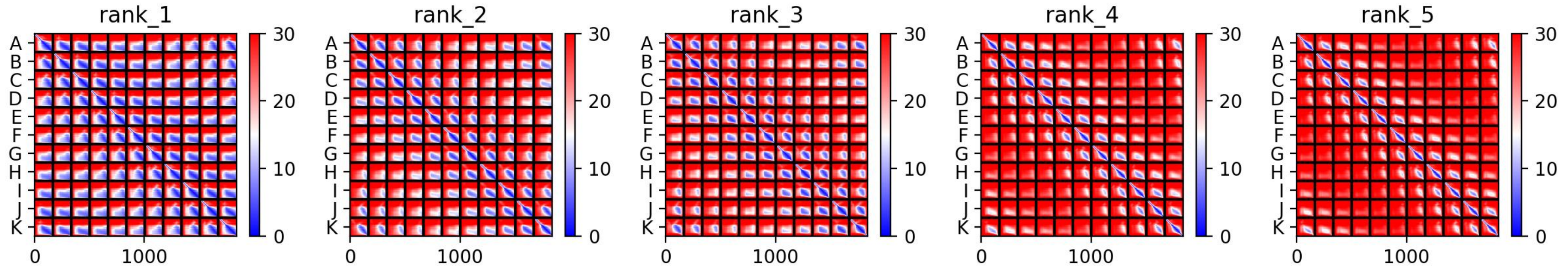

### XP_061423086.1 *L. reissneri*

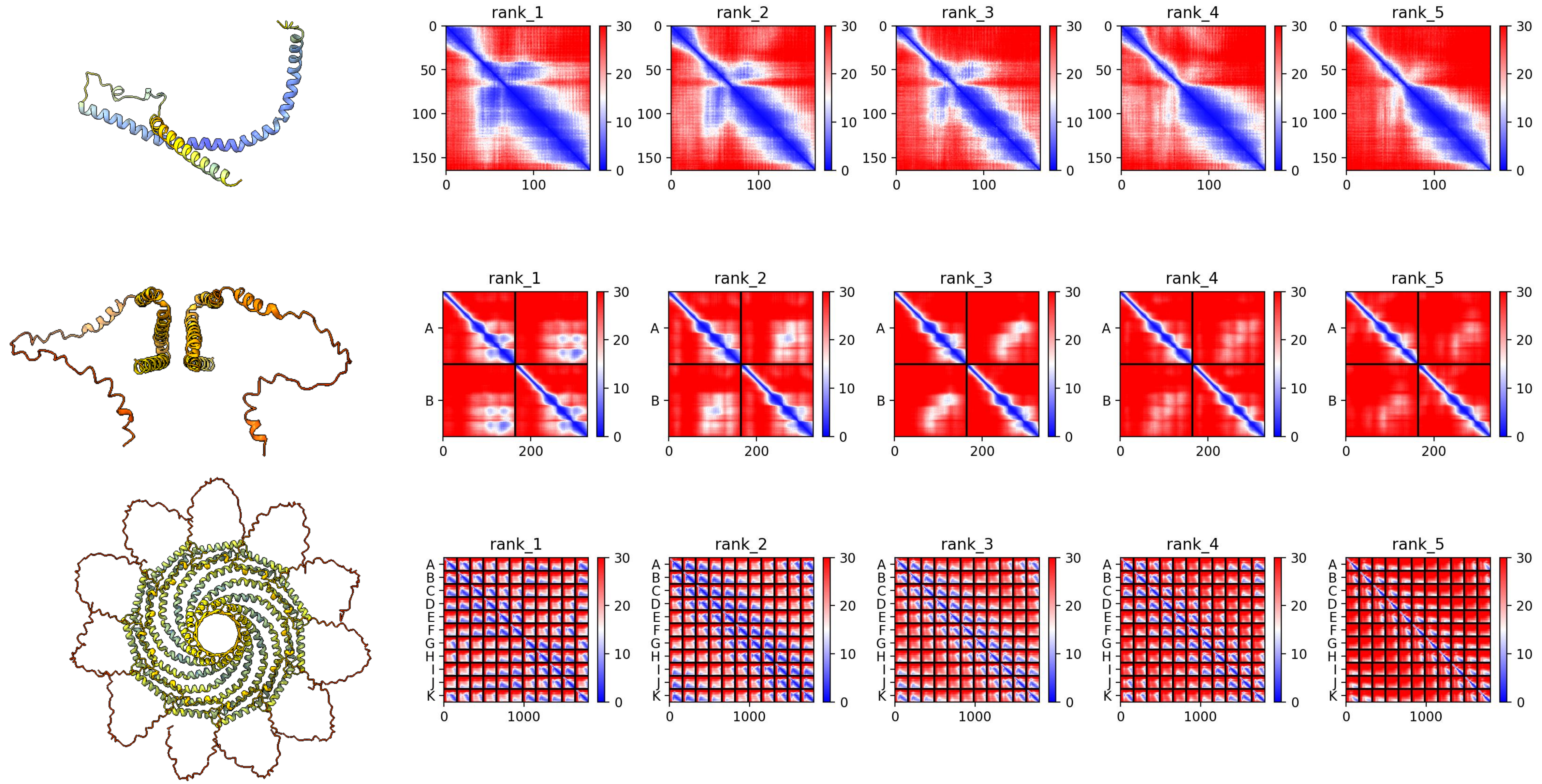

### XP_061428890.1 *L. reissneri*

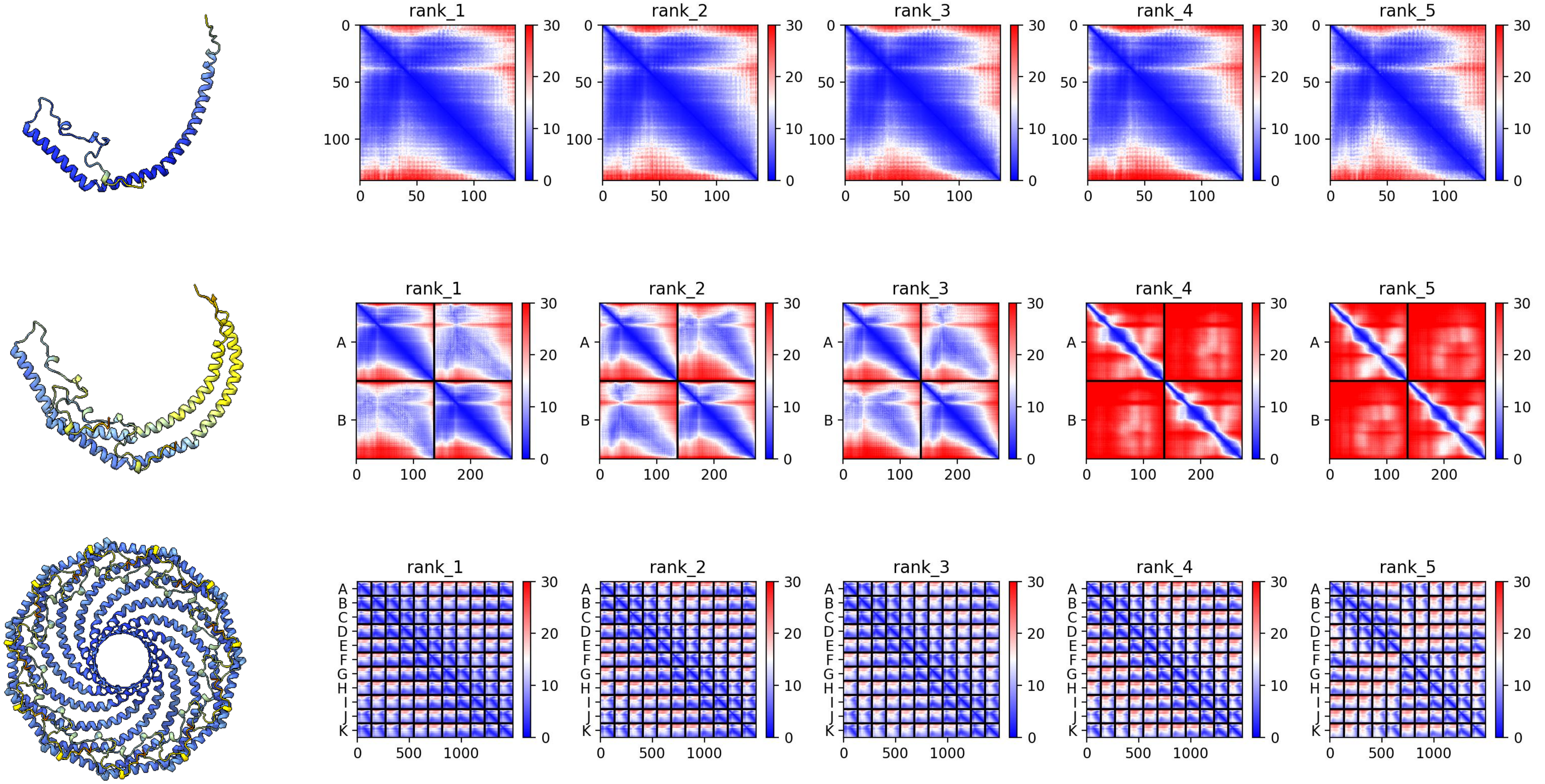

### XP_061428788.1 *L. reissneri*

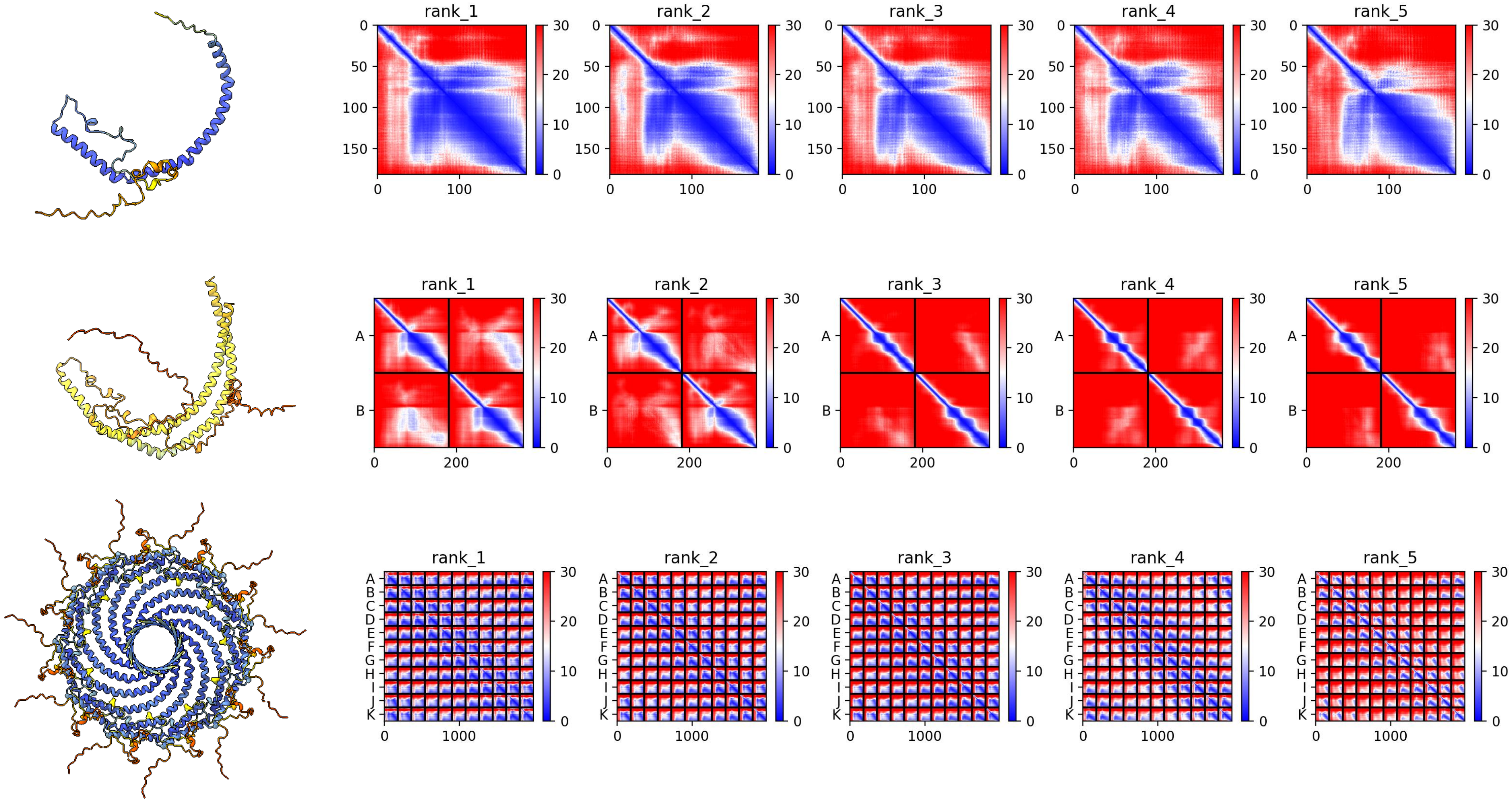

### XP_032827412.1 *P. marinus*

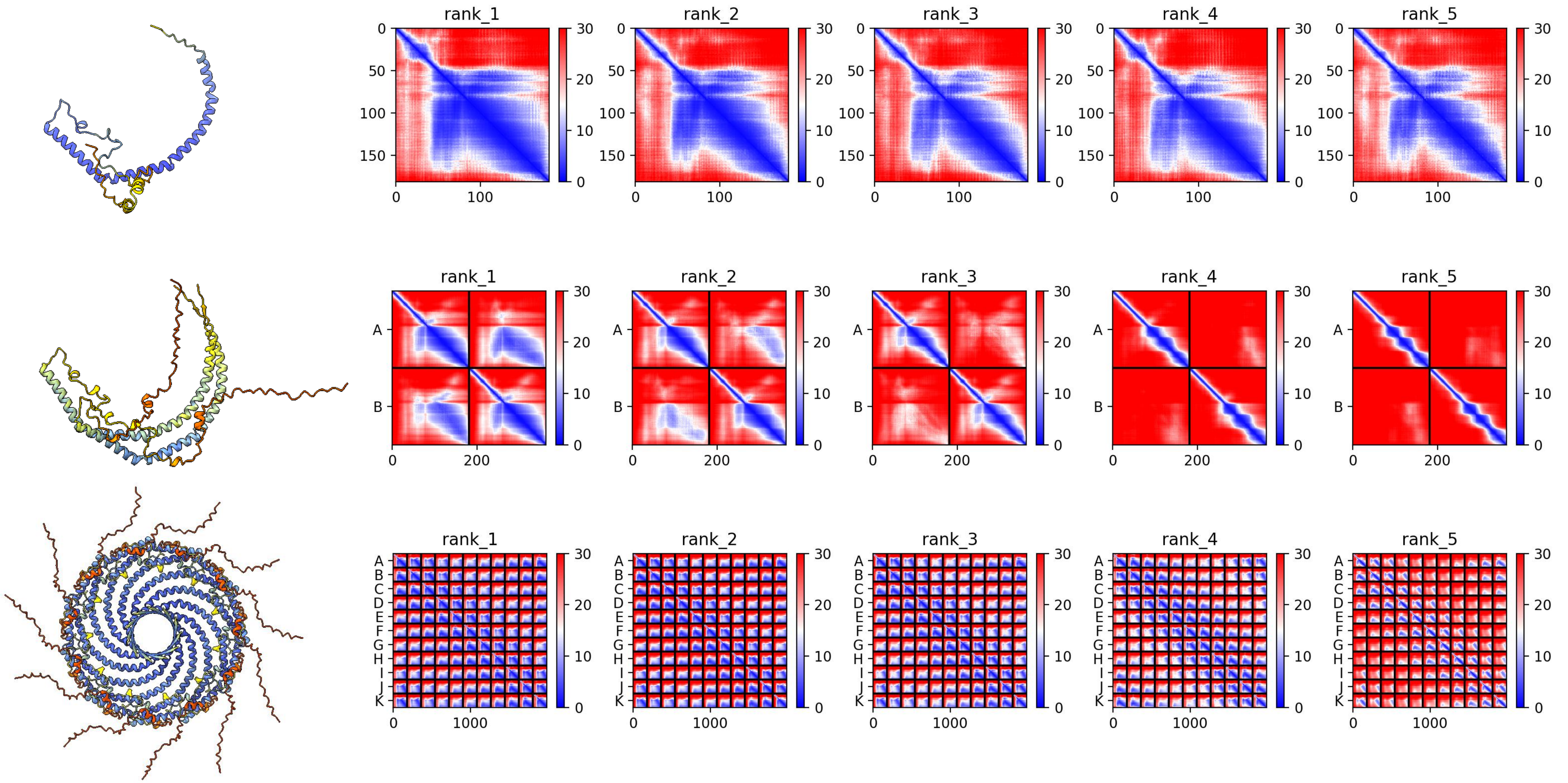

### XP_032819264.1 *P. marinus*

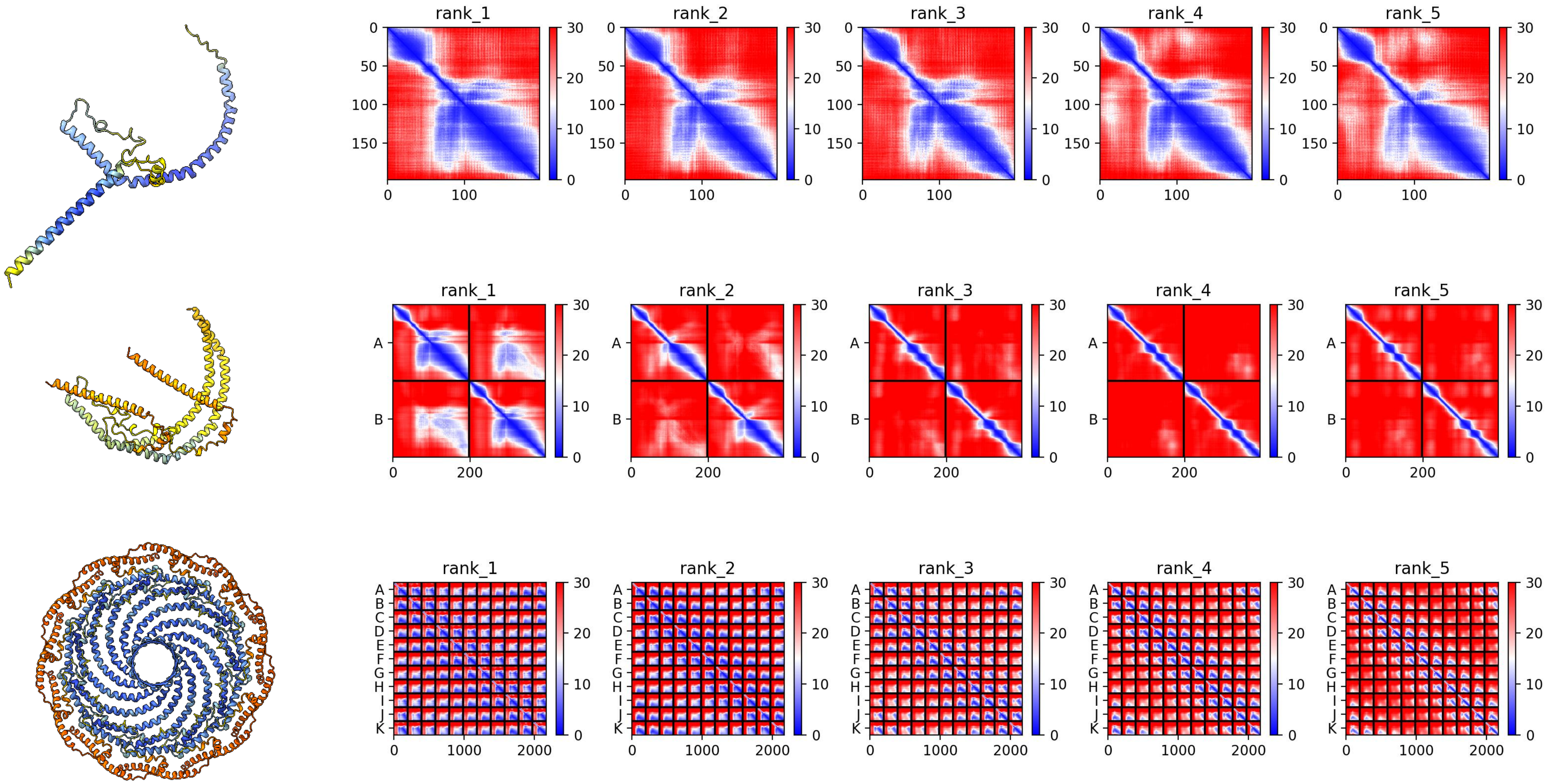

### XP_032819263.1 *P. marinus*

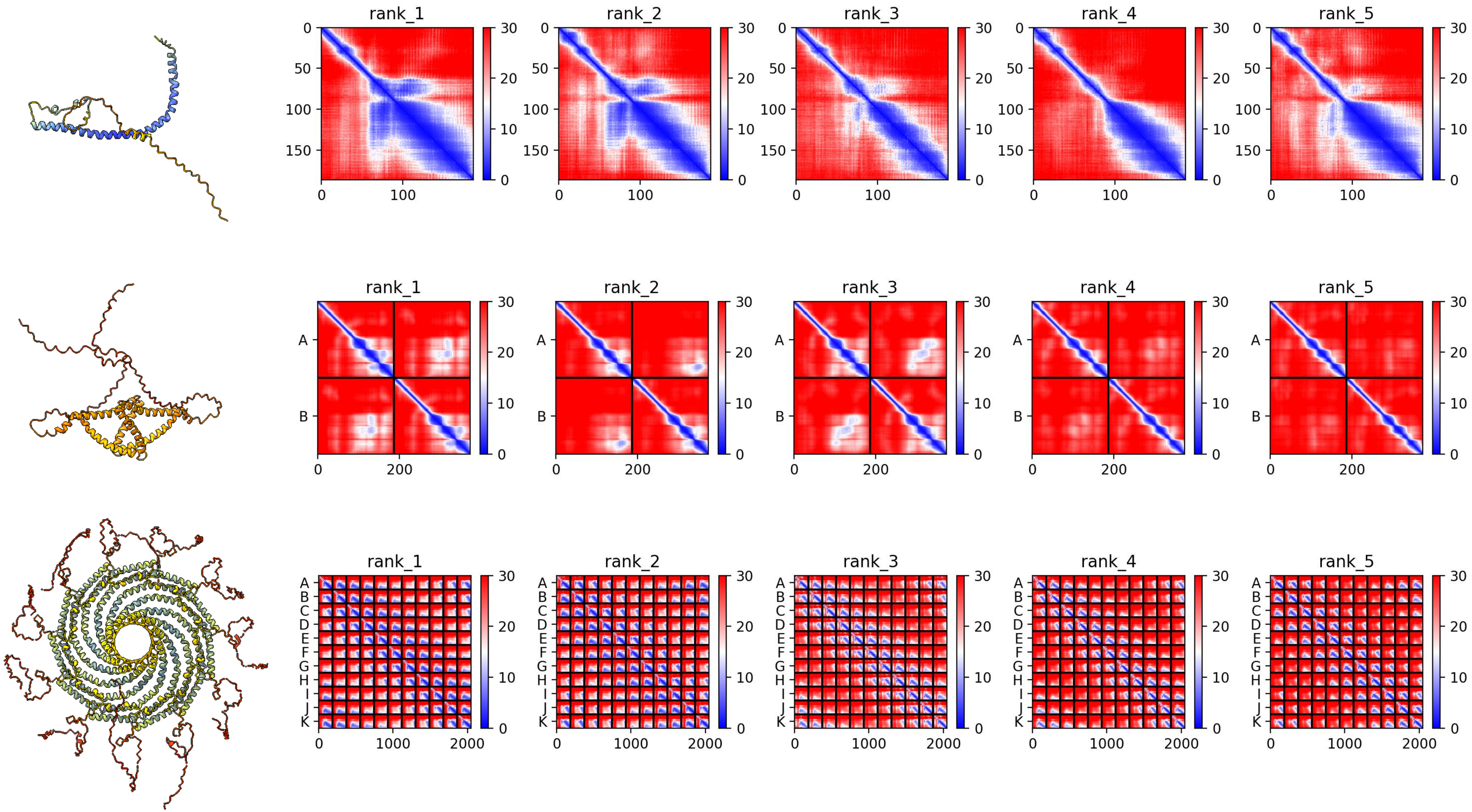

### XP_032827409.1 *P. marinus*

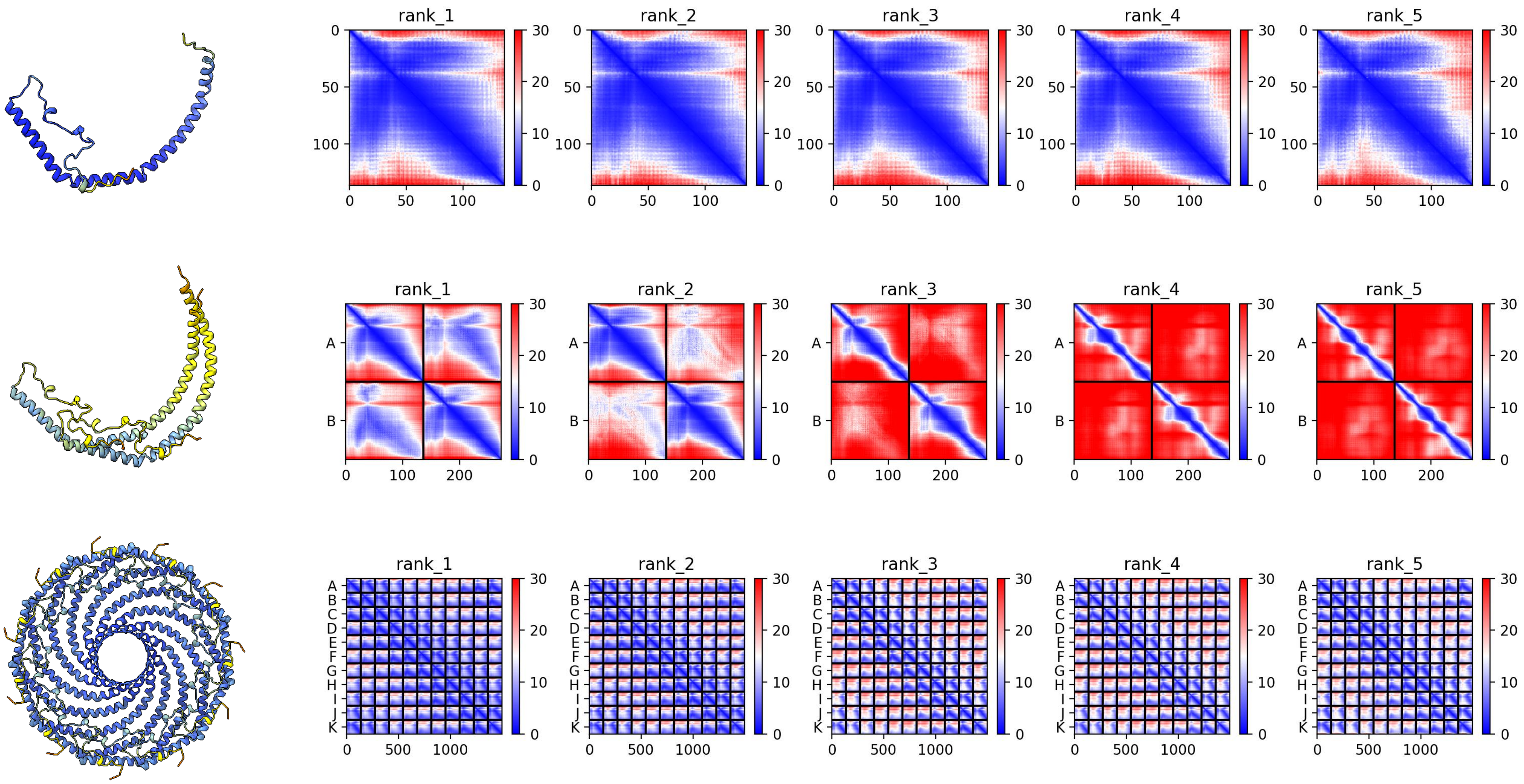

### XP_032827408.1 *P. marinus*

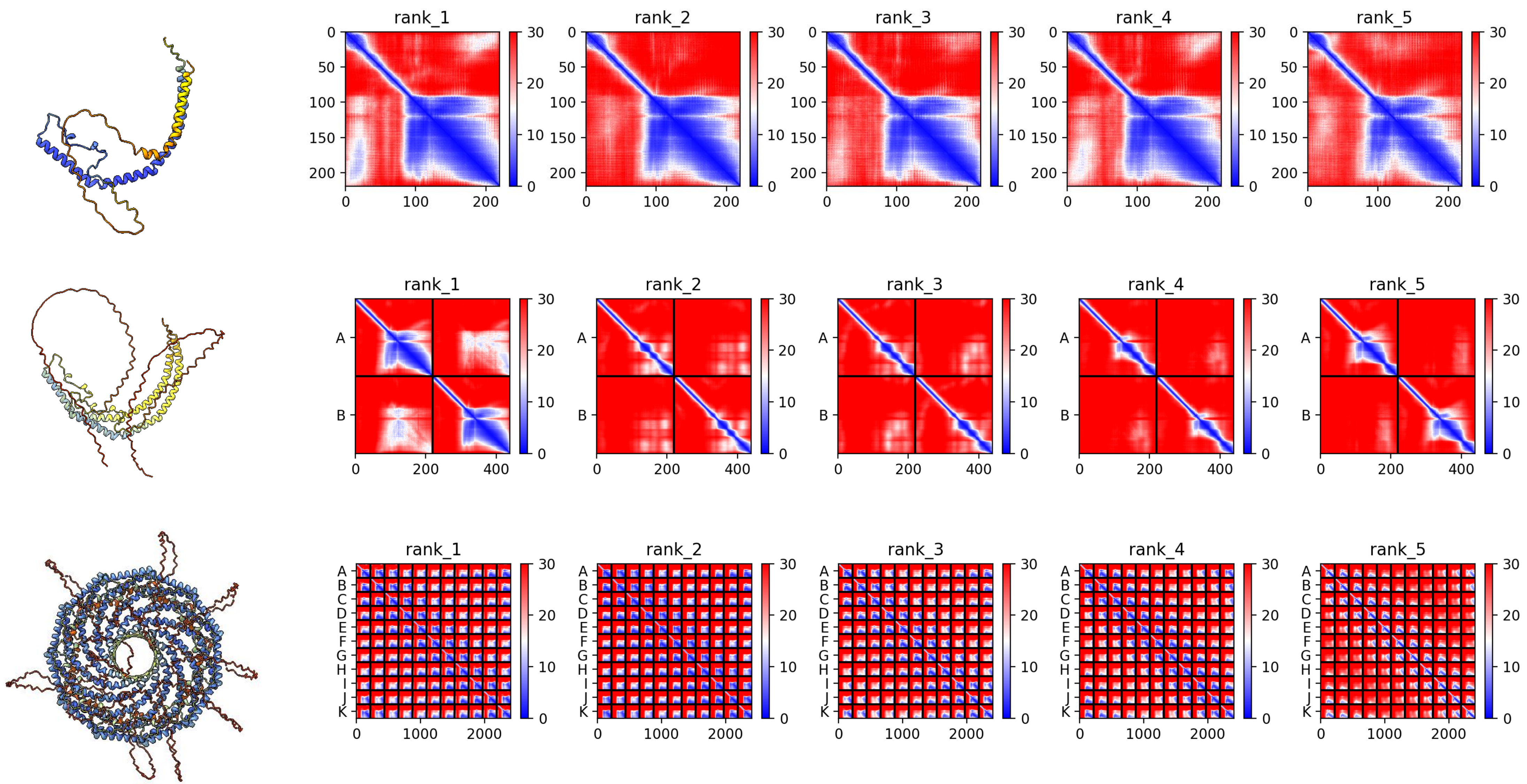

#### *XP_035693613.1 B. floridae*

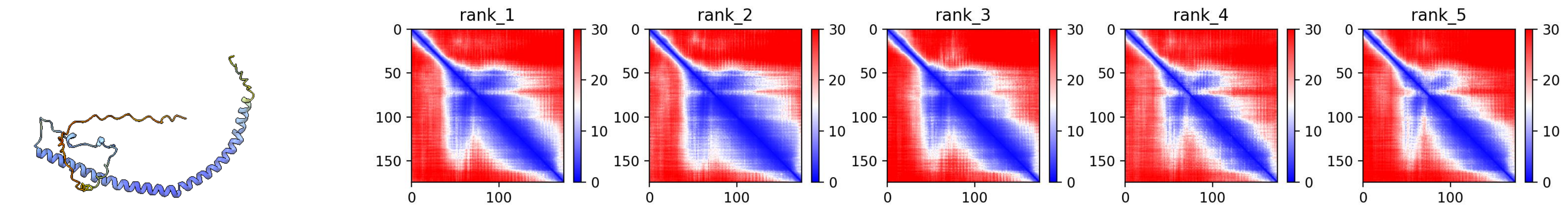


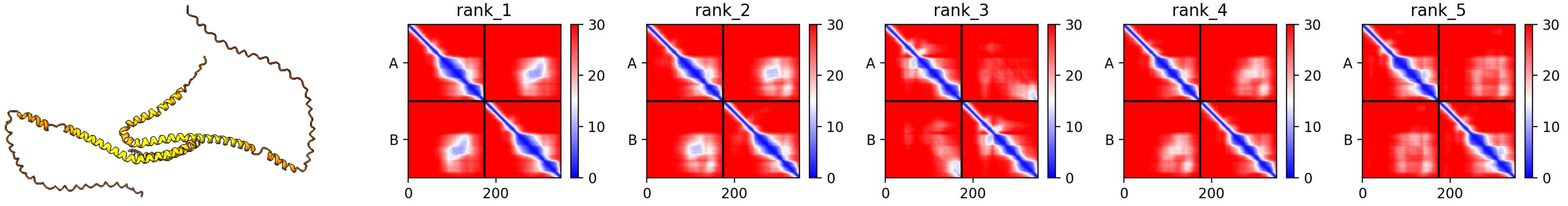


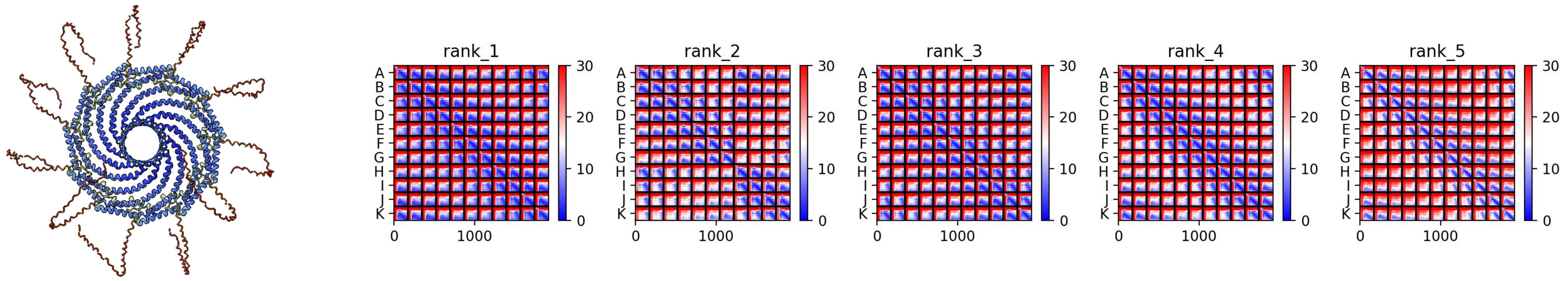

#### *XP_035698192.1 B. floridae*

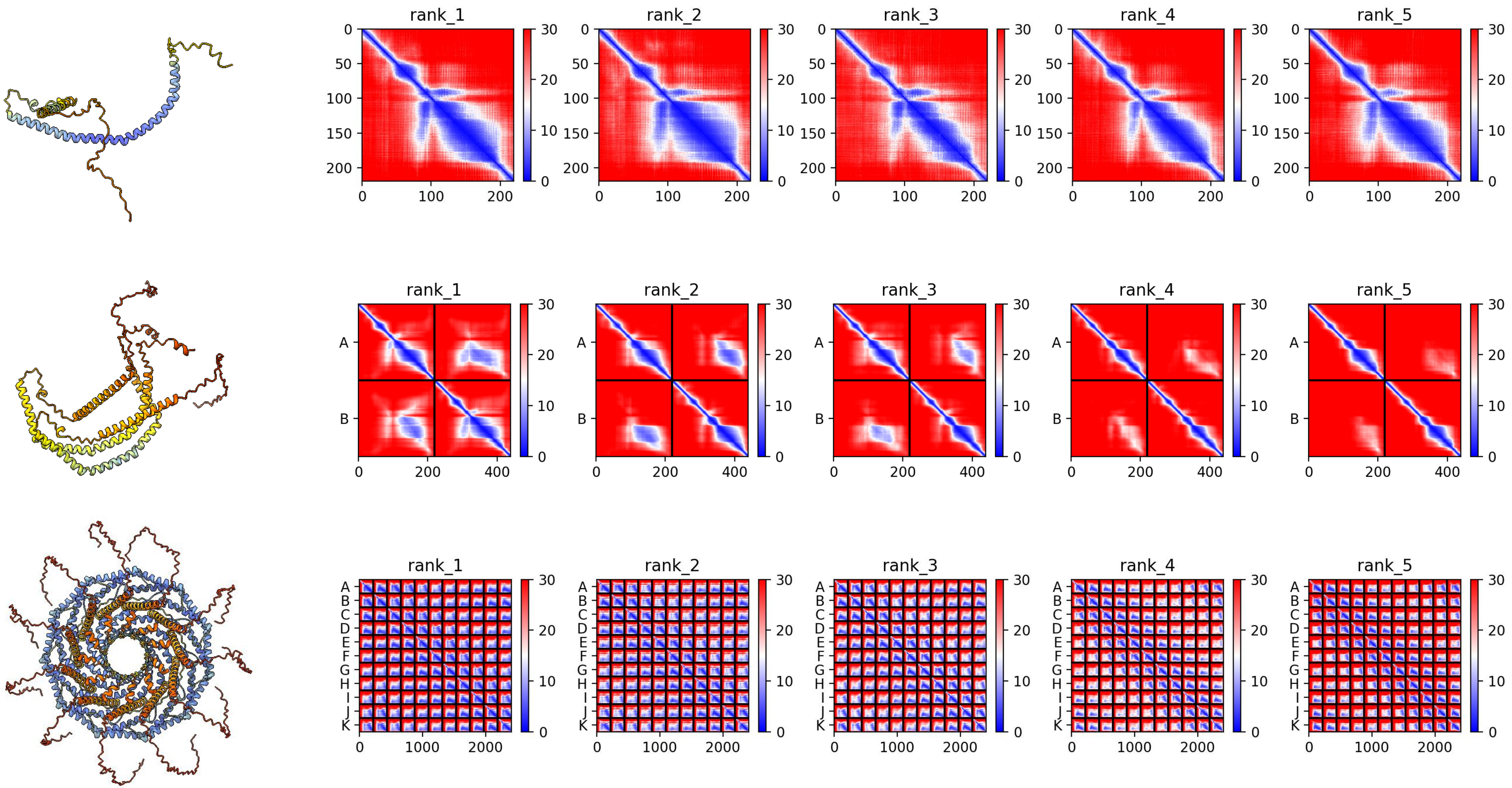

### *XP_035680487.1 B. floridae*

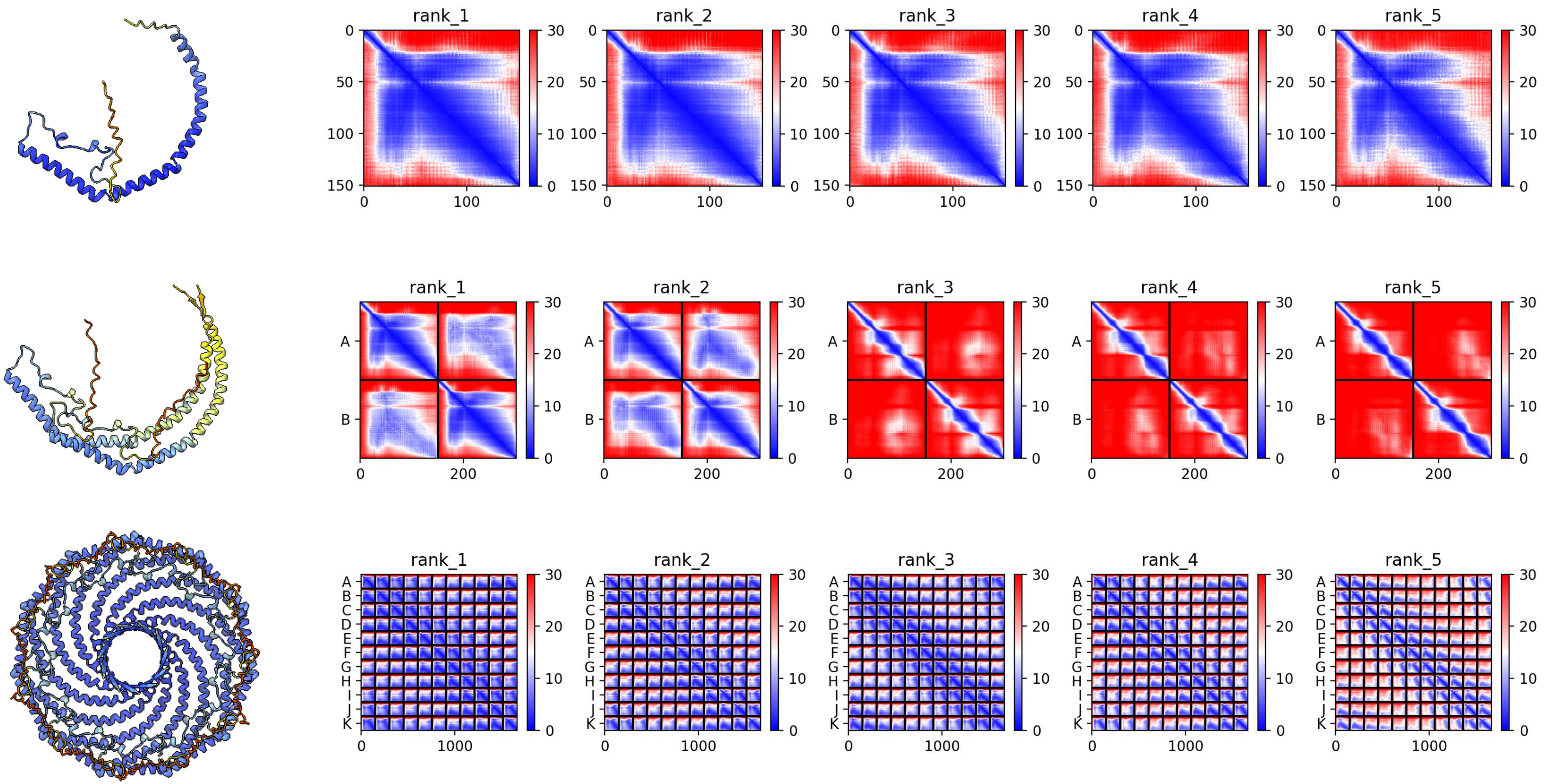

#### *XP_035680488.1 B. floridae*

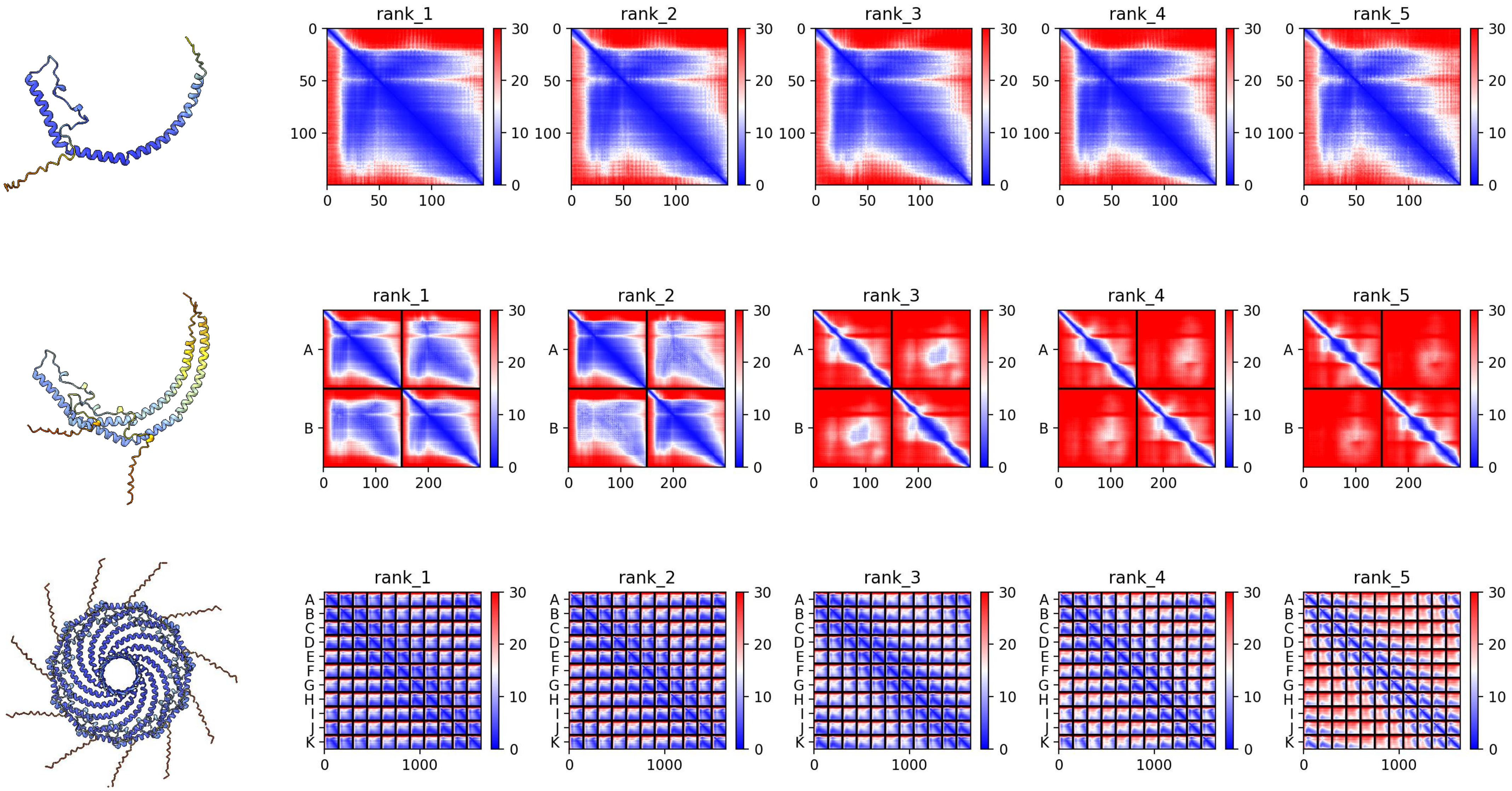

### *CF919476 B. floridae*

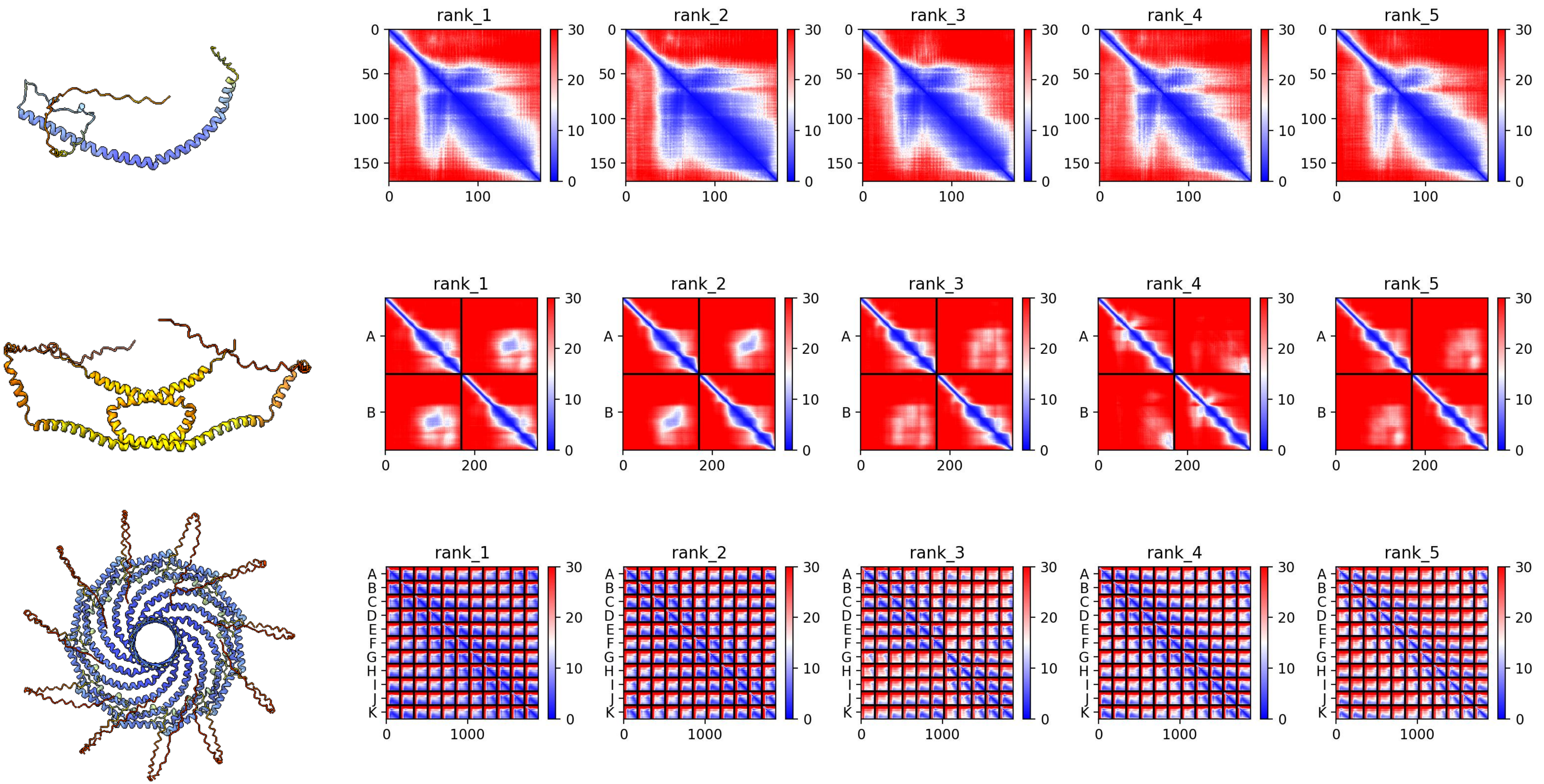

### NP_001099134.2 G. gallus

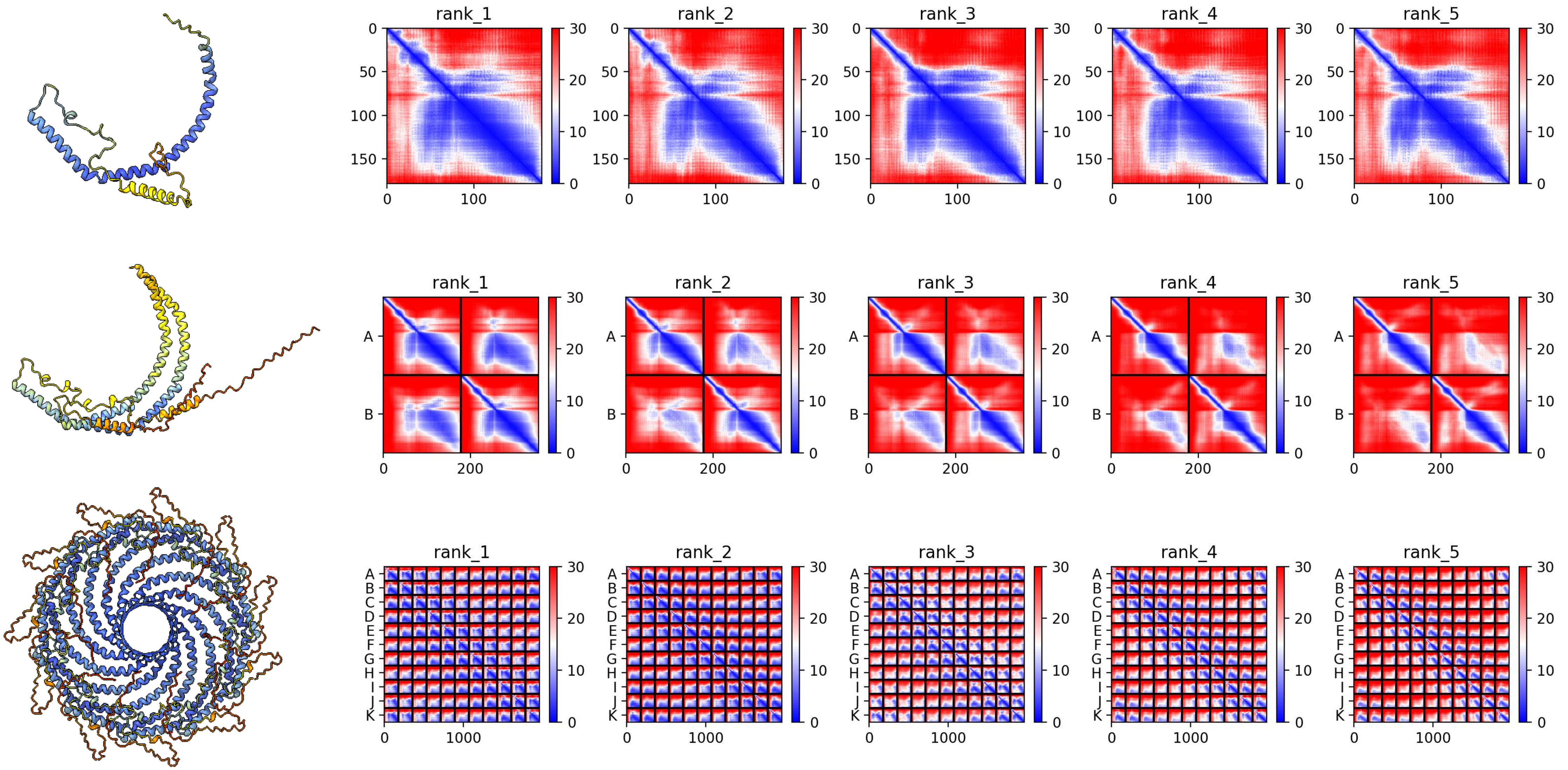

### *NP_001007087.2 G. gallus*

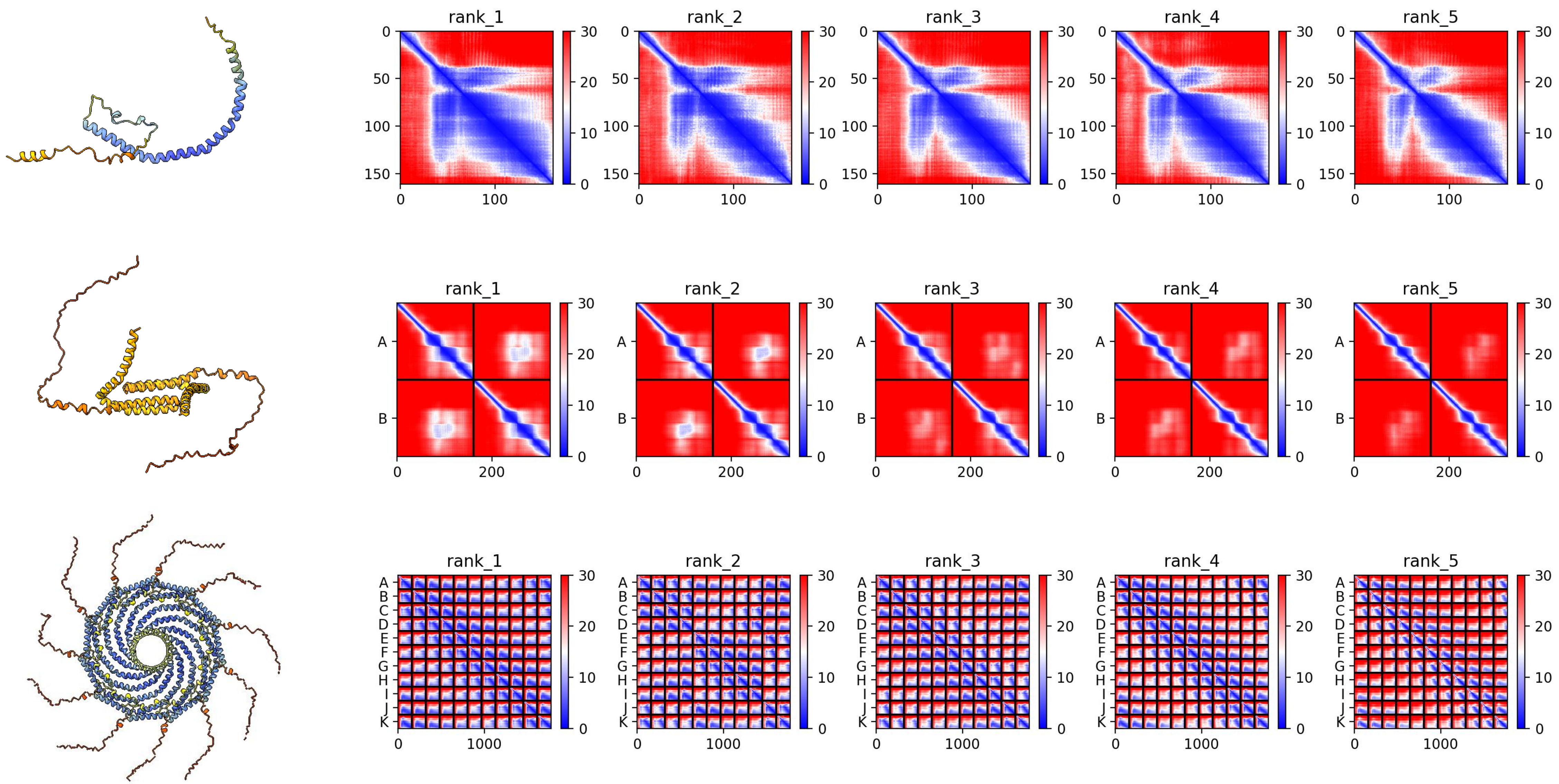

### *NP_989701.2 G. gallus*

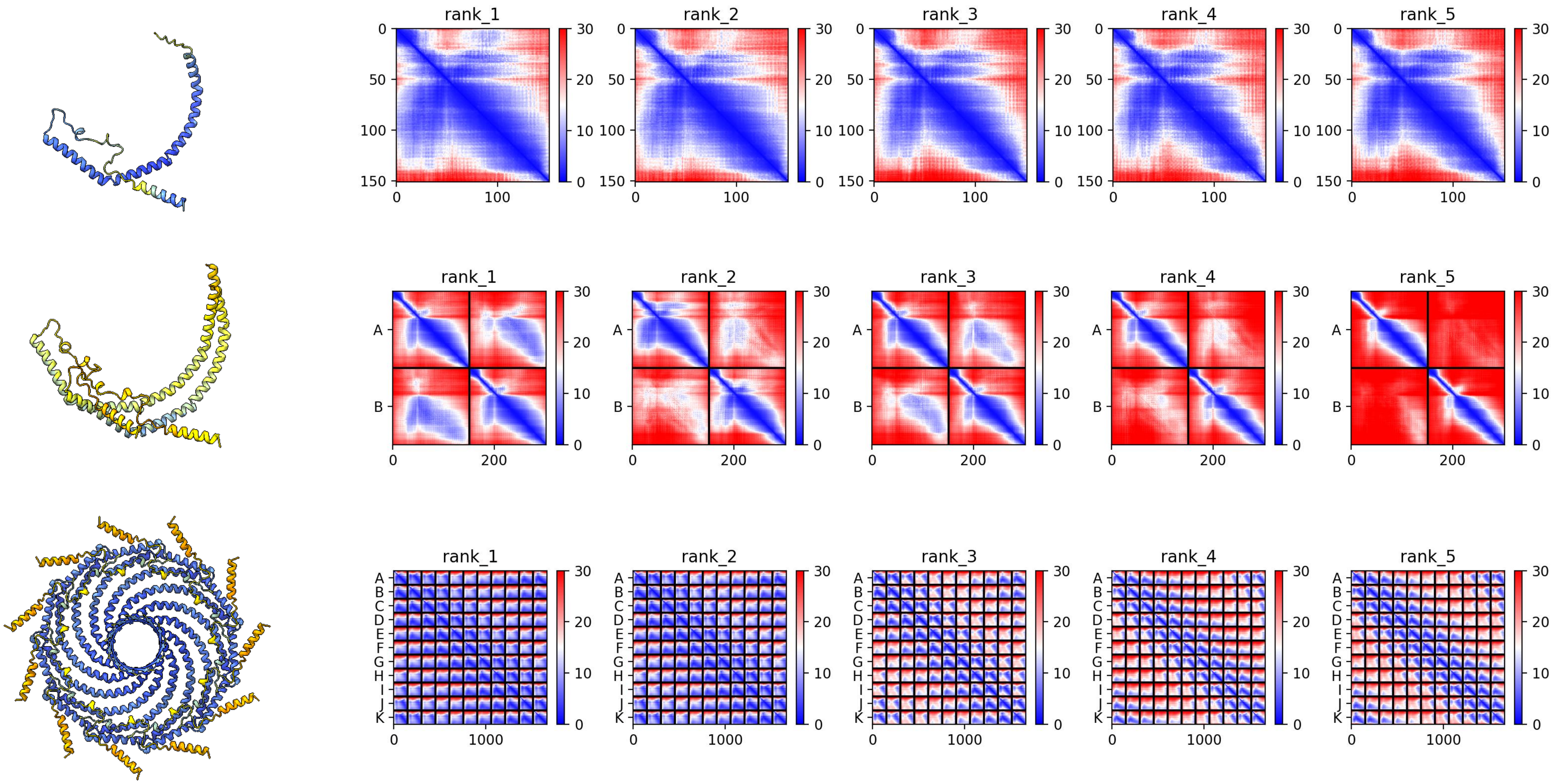
