## Supplementary figures and images for "Evolutionarily diverse caveolins share a common structural framework built around amphipathic discs"

### Table S1

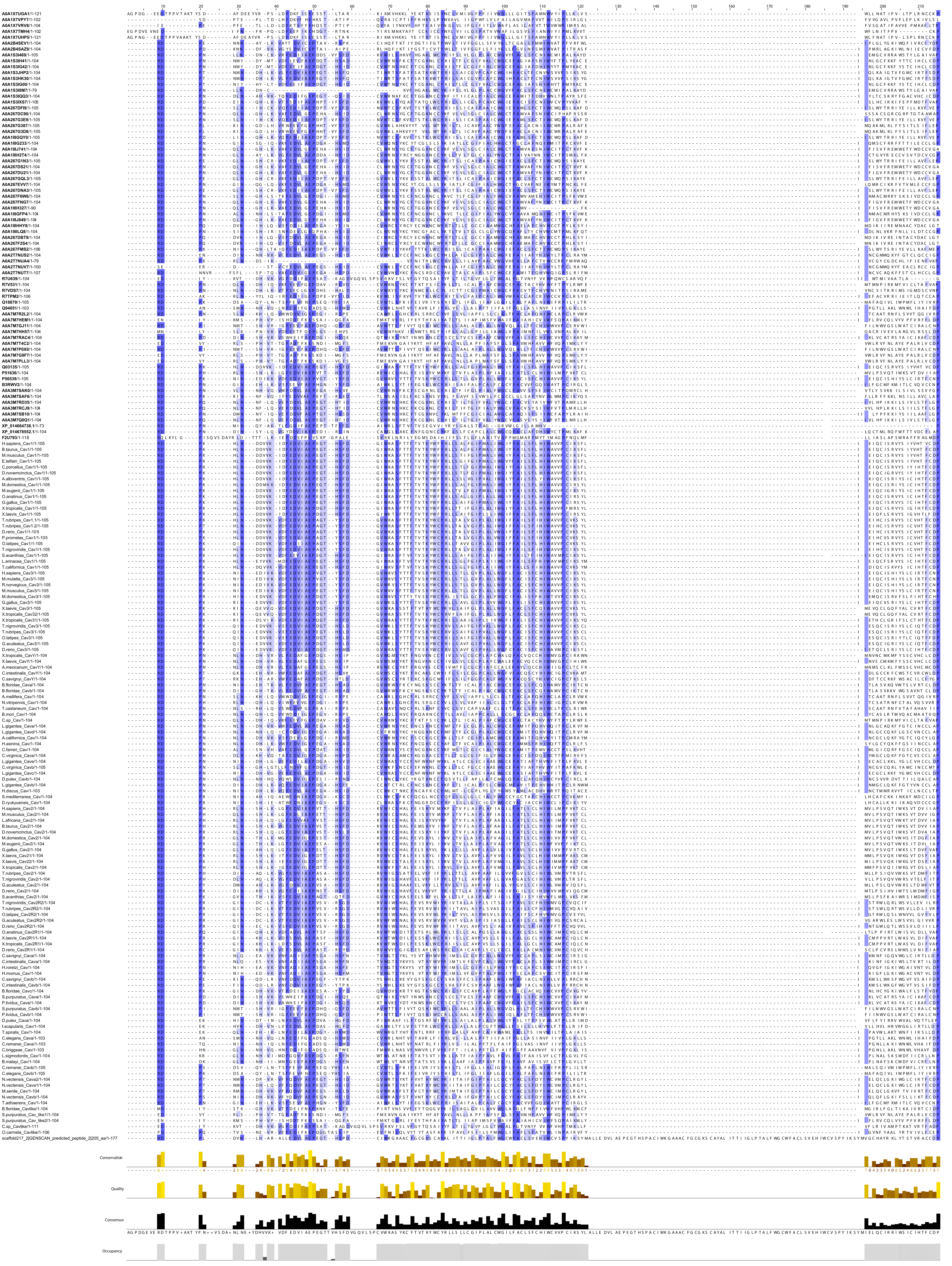

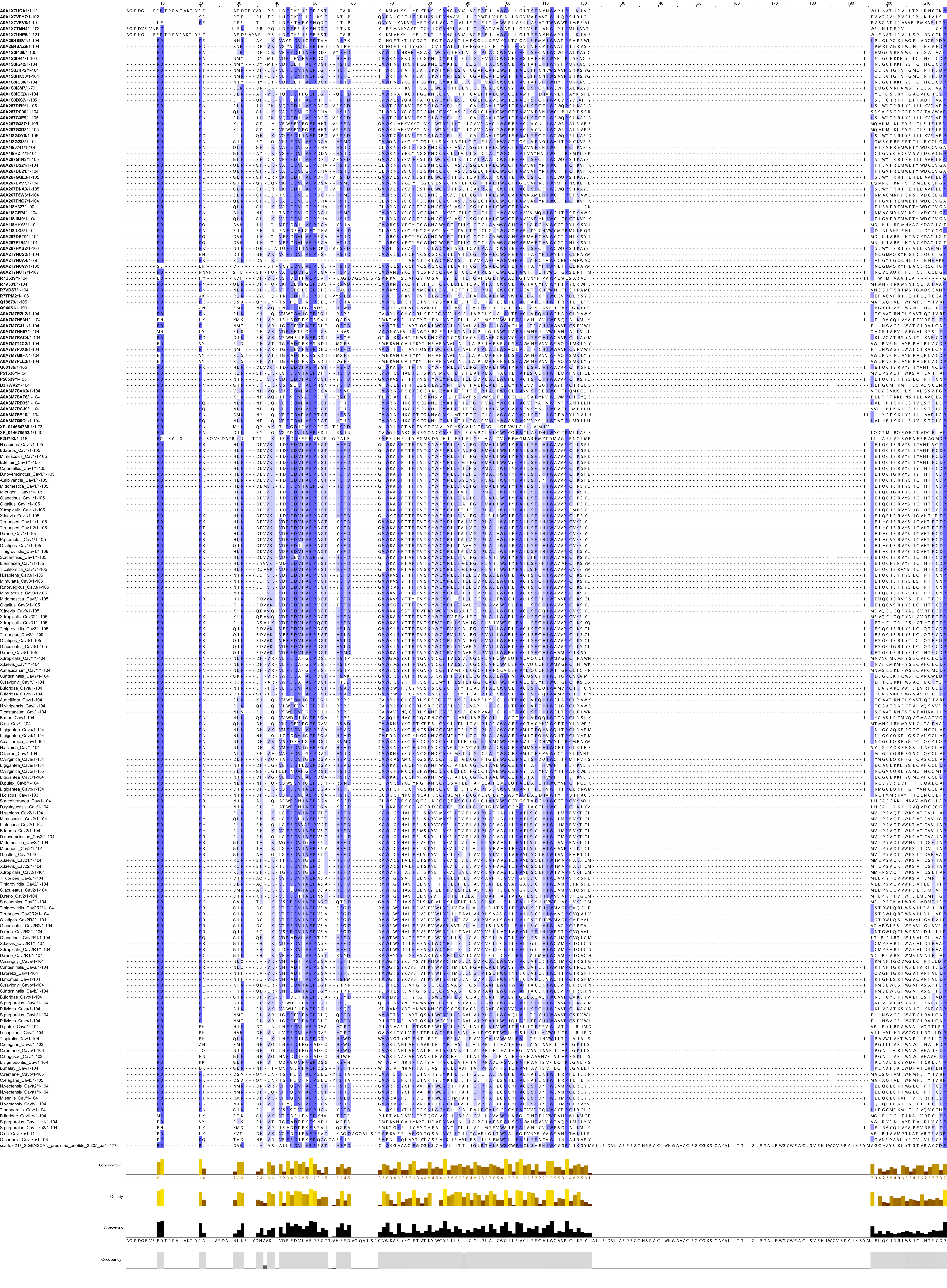

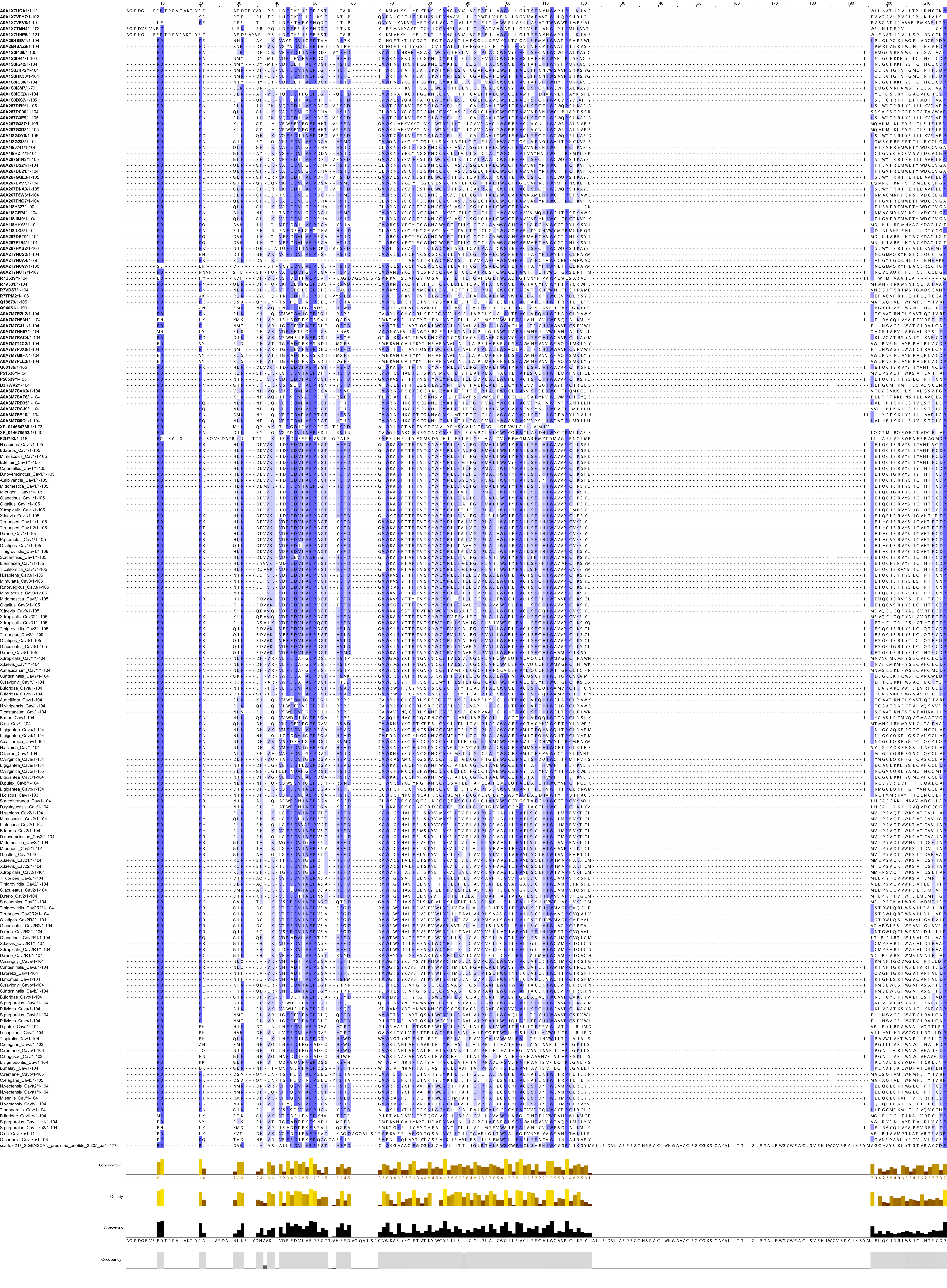
